# Beyond analytic solution: analysis of FRAP experiments by spatial simulation of the forward problem

**DOI:** 10.1101/2023.03.05.531160

**Authors:** Ann E. Cowan, Leslie M. Loew

## Abstract

Fluorescence redistribution after photobleaching (FRAP) is a commonly used method to understand the dynamic behavior of molecules within cells. Analytic solutions have been developed for specific, well-defined models of dynamic behavior in idealized geometries, but these solutions are inaccurate in complex geometries or when complex binding and diffusion behaviors exist. We demonstrate the use of numerical reaction-diffusion simulation approaches using the easily accessible Virtual Cell (VCell) software, to establish methods for analyzing photobleaching data. We show how multiple simulations employing parameter scans and varying bleaching locations and sizes can help to bracket diffusion coefficients and kinetic rate constants. This approach is applied to problems in membrane surface diffusion, diffusion and binding in cytosolic volumes in complex cell geometries, and analysis of diffusion and binding in intracellular liquid droplets.

**Statement of Significance:** Fluorescence Redistribution After Photobleaching (FRAP) is a widely used experimental method that can reveal important parameters for reaction/diffusion events within cells. However, analytic methods to analyze FRAP experiments are limited to specific geometries and conditions. We demonstrate how spatial numerical simulation methods using the freely available software Virtual Cell can be used to obtain parameter information from FRAP experiments in situations that are not amenable to analytic solutions and that are accessible to most bench biologists.

## Introduction

Fluorescence Redistribution After Photobleaching (FRAP), was pioneered by Ken Jacobson and several other renowned biophysicists in a series of seminal papers in 1976(1–4). It involves the bleaching of a fluorescent label with a brief but intense illumination in a small region of a cell; the recovery of the fluorescence is then monitored to assess the kinetics of diffusion and/or binding of the labeled molecule. With the advent of laser scanning confocal microscopes, FRAP is now routine in many laboratories, and the development of new labeling methods allows fluorescence probing of increasingly diverse events within cells. Extracting accurate biophysical parameters from such data, however, remains a daunting task. The complex interplay of reaction-diffusion events with each cell’s specific geometry, as well as distortions introduced by imaging systems, preclude the use of available image analysis methods; simple analytic approaches to determine parameter values such as on and off rates, diffusion coefficients, or velocities require assumptions that are rarely met in real experimental situations. Few software tools are available that can aid microscopists in obtaining these parameters. In part this is because the characteristic time of the response in a reaction-diffusion system is highly dependent on the geometry of both the cell and the experimental perturbation. In addition, distortions introduced by the imaging system are not easily accounted for. However, analytic methods have been successfully developed to analyze specific experimental designs ((5–14); reviewed in (15)). Most of these methods are mathematically complex, cannot be generalized and are difficult for cell biologists to apply. These analytic methods necessarily each apply only to a specific class of idealized cell geometries (e.g. sphere, disc, or cylinder) and contain simplifying assumptions that can significantly reduce the accuracy of the kinetic constants determined. More importantly, analytic solutions such as these use only a small subset of the information in the image, e.g. the bleached area of a FRAP experiment, but there is useful information within the entire spatiotemporal dataset that can be used to refine the accuracy of the parameters or obtain additional information about the system.

Approaches for analyzing photobleaching and photoactivation experiments based on numerical solutions of reaction-diffusion in 2D have been published (5, 9, 16, 17), and in several cases have been integrated into software tools. One of the first such software offerings was Tropical (18), which used a sequential optimization method for 2-D reaction-diffusion problems. More recent solutions include PyFRAP (19), provided as an ImageJ plugin, that is also restricted to 2 dimensional simulations. A stochastic software tool has also been developed for 3D simulations ((20)) and was applied to analyze FRAP experiments in bacterial cells.

The Virtual Cell (VCell) modeling environment is uniquely suited to simulate dynamic fluorescence imaging experiments because it is designed to solve reaction-diffusion equations within any given geometry(21–23). Designed as a general modeling tool specifically for use by experimental biologists, VCell is capable of simulating any reaction-diffusion-advection system defined within the volumes and on the surfaces of arbitrarily shaped 2D or 3D geometry, including geometry obtained from experimental images. VCell is used by hundreds of researchers around the world, to whom it provides a powerful tool for collaborative research and modeling in computational cell biology.

Many VCell models have been created specifically to simulate experimental fluorescence-based imaging experiments to derive the quantitative biological details of the system being studied. These include models of uncaging reactions (24, 25), photobleaching (26–29), calcium indicator probes (24), translocation of a fluorescent binding protein (30) and fluorescent biosensors (31, 32). VCell models are stored within a centralized database, and many of these models have been made available to the public from within VCell.

In this work, we created a set of public VCell models that can serve as templates for the analysis of complex photobleaching experiments. To demonstrate and validate the utility of using VCell to simulate FRAP experiments, we applied analytical methods to VCell simulations using idealized geometries and diffusion conditions. We then created a series of VCell models that demonstrate simulation of photobleaching experiments for cytoplasmic or membrane species using various non-ideal, experimentally relevant bleaching geometries and cellular geometries. These models can also incorporate binding kinetics and we employed a range of rate constants and diffusion coefficients to bracket commonly observed experimental values.

## Methods

All numerical simulations were performed using the Virtual Cell (VCell) software freely available at http://vcell.org. Two Biomodels were created for simulations, both are available to the public from the VCell database under username “les”, one with Biomodel Name: FRAP_Membrane_Rel and one with Biomodel Name: FRAP_cyt. Planar and spherical geometries for spatial simulations were specified analytically; the 3-dimensional cellular geometry was reused from a publicly available VCell Tutorial model and was originally created from a confocal image stack of a differentiated N1E neuroblastoma cell.

The BioModel FRAP_Membrane_Rel includes multiple applications for each of the different types of simulations described in the Results, with species and bleaching reactions placed in a membrane compartment “PM”. The Model FRAP_cyt also includes multiple applications and places species and bleaching and binding reactions within a volume compartment “Cyt” (see Figure 1). Where binding interactions are included, initial (i.e. pre-bleach) steady state conditions were established based on the *kD* and used as initial concentrations for binding species. Default parameter values and species initial conditions are shown in Table 1; changes made to the default parameter values are specified in the text and figure legends.

**Figure 1.**
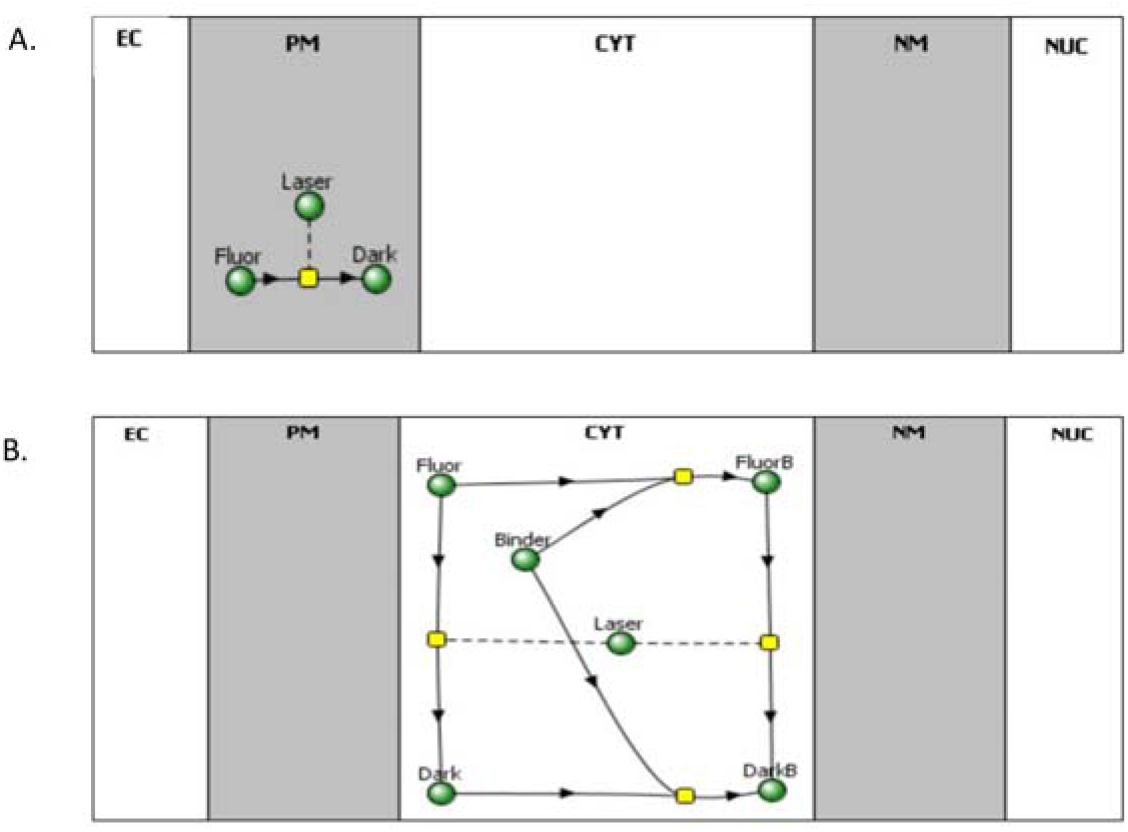
Reaction Diagram for VCell models. A. FRAP_Membrane._Rel B. Frap_Cyt. Gray regions represent 2D surfaces (i.e. membranes) and white regions represent volumes. The green spheres represent species (i.e. variables); the yellow squares represent reactions. An arrow emanating from a species indicates that the species is consumed when the reaction rate is positive; an arrow pointing to a species indicates that species is produced when the reaction rate is positive. Dashed lines indicates the corresponding species (“Laser”) modulates the reaction rate, but is neither produced nor consumed.

**Table 1.** Default Parameter values and initial conditions for simulations.

| <u>Biomodel FRAP Membrane</u> |  |  |
| --- | --- | --- |
| Parameter | Value | Units |
| duration | 1 | s |
| BleachRadius | 2 | um |
| sigmalateral | 0.5 | um |
| start | 1 | s |
| sigmaaxial | 1.5 | um |
| Kr | 0 | s-1 |
| kbleach | 10 | um<sup>2</sup>.s-1.molecules-1 |
| Species | Value | Units |
| Dark | 0 | molecules.um-2 |
| Laser |  | molecules.um-2 |
| Fluor | 10 | molecules.um-2 |
| Fluor Diffusion Coefficient | 0.1 | um<sup>2</sup>.s-1. |
| <u>Biomodel FRAP cyt</u> |  |  |
| Parameter | Value | Units |
| duration | 1 | s |
| BleachRadius | 2 | um |
| sigmalateral | 0.5 | um |
| start | 1 | s |
| sigmaaxial | 1.5 | um |
| kbleach | 1 | s-1.uM-1 |
| Species | Value | Units |
| Dark | 0 | uM |
| Laser |  | uM |
| Fluor | 10 | uM |
| Flour Diffusion Coefficient | 25 | um-2.s-1 |
| Binder | 0 | uM |
| Binder Diffusion Coefficient | 0 | um-2.s-1 |

The default Fully-Implicit Finite Volume, Regular Grid (Variable Time Step) solver was used for all spatial simulations, using default Error Tolerance values. For FRAP_cytosol simulations that use VCell “Microscope Measurement Protocols” the Z projection function was selected that convolved FluorB and Fluor into a single “fluorescence” value. Specific details of each simulation can be obtained either directly from the publically available VCell models or from the VCell generated simulation reports in supplemental data.

Fluorescence curves analyzing binding to a uniformly distributed immobile binding component in the cytosol, were generated by exporting the Z projection “fluorescence” data as a .nrrd file; this data was then analyzed in ImageJ to collect average intensity within either the entire 4 μm diameter bleach region or in a 4 μm circle centered within the bleach region when half the cell was bleached. Data were normalized as ((I_t_ – I_0_)/(I_0_ – I_i_))/fb where I_t_ is intensity at time t, I_0_ is the intensity immediately postbleach, I_i_ is the prebleach intensity and fb is the fraction of total intensity remaining after the bleach. Fluorescence curves analyzing binding within a localized droplet were data obtained from a central slice of the 3-dimensional simulation (thus representing a confocal slice) at a single pixel in the center of the bleach region, using the internal VCell simulation analysis features; kymographs were also generated directly within VCell.

## Results

### Setting up bleaching and binding reactions in VCell

For consistency, all VCell models used in our analyses describe 5 cellular compartments: extracellular (EC), plasma membrane (PM), cytosol (CYT), nuclear membrane (NM) and nucleus (NUC) (Note that the NUC and NM compartments are only required to simulate cytosolic diffusion, which is excluded from the nuclear region; however, they are included in the membrane bleach simulations so the same experimental geometry could be applied to both types of bleaching experiments). The basic photobleaching reaction is modeled using three species: a fluorescent molecule (Fluor), the dark form of the molecule after photobleaching (Dark) and a species (Laser) used to describe the bleaching intensity as a function of time and space. To model photobleaching of membrane components, the species are placed in the PM compartment. To model photobleaching of cytosolic components, the species are placed in the CYT compartment. A single rate expression (r0) describes the bleaching reaction that transforms Fluor into Dark using a rate law of:

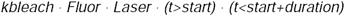

Where *kbleach, start* and *duration* are defined as parameters so they can be easily varied to match a particular experiment. *kbleach* defines the strength of the bleaching beam, *Laser* is a species that can be mapped to any arbitrary region in the geometry, and the Boolean expressions describe the start and end time (=*start + duration*) of the bleaching reaction. Because photobleaching is, generally, irreversible, the reverse rate constant for the reaction is 0; note that a reverse reaction could be included to model the case of dark state to light state transitions during the experiment. This simple model accounts for all reactions needed to model a simple photobleaching experiment with only diffusion.

For models that include binding of the labeled molecule to an unlabeled binding partner, we also include a pair of identical mass action reactions between a species called “*Binder*” and, respectively, *Fluor* and *Dark*; the products of these reactions are respectively, *FluorB* and *DarkB*. The rates of these reactions are given by:

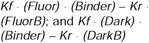

If we are to only consider diffusion, we can either set the forward and reverse rate constants, *Kf* and *Kr*, to 0, or we can set the initial concentrations of *Binder, FluorB* and *DarkB* to 0. If we do wish to consider the effect of binding, we set the initial concentrations of the species *Fluor, Binder* and *FluorB* to their steady state equilibrated values, so that the bleaching event is the only trigger for the subsequent dynamics (*Dark* and *DarkB* are, of course, 0 before the bleaching event).

Importantly, we also need to include a bleach reaction that converts *FluorB* to *DarkB* under the influence of *Laser*, using the same rate law described above. The diagram in Figure 1 shows the relationships of the variables (i.e. species) in this kinetic scheme. When analyzing the simulation results, we often compute the *TotalF* (=*Fluor* + *FluorB*), as this is the experimental observable.

### VCell simulations of ideal 2D diffusion are well fit by theory

Historically, analytical theories developed for FRAP were idealized to solve for small bleaching spots in an infinitely large 2D surface. The diffusion analysis described in Kang et al, 2012 (33) was developed to more accurately analyze photobleaching experiments using a circular bleach on a scanning confocal microscope. This analytic solution uses the radius of the originally prescribed bleached region and the Gaussian radius of the bleached region at some later time to solve for the diffusion coefficient using the equation:

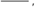

where r_e_ is the effective radius of the bleached area determined using a Gaussian fit at some time t after bleaching, r_n_ is the nominal experimentally defined radius of the bleach region, and τ_1/2_ is defined as the half time of recovery from time t, measuring the average fluorescence over the entire initial bleach surface area. We examined how well this analytic approach was able to recover the diffusion coefficient defined in our VCell simulations.

As an initial test we created simulations in which the fluorescent molecules were uniformly distributed in a 100 × 100 μm 2D membrane sheet (PM) with value boundary conditions to provide an infinite source of Fluor (biomodel “FRAP_Membrane_Rel”, Application: “100um planar membrane, value BC”). This condition closely approximates the assumption of an infinite source of fluorophore that is used to obtain the analytic solution developed by Kang et al (2012) (33). In the simulations the bleached region is defined by placing the immobile Laser species that catalyzes the bleaching reaction within a circle of given radius centered at x = 50, y = 50. Results of simulations with a fixed value boundary condition for Fluor, with a diffusion coefficient set to 1 μm^2^/s and an initial bleach (r_n_) of 4 μm diameter are shown in Figure 2. We found the earliest postbleach profile that was well fit by a Gaussian profile occurred at 5s in the simulation (3s following termination of the Bleach) and yielded a value for r_e_ of 6.825μm and τ_1/2_ of 9.02s, resulting in a calculated D of 0.0.995 μm^2^/s. Similar analyses with a bleach radius (r_n_) of 1 and 3 μm yielded diffusion coefficients of 0.989 and 1.005 μm^2^/s, respectively. These are all gratifyingly close to the input value of 1μm^2^/s. As an additional check, we also used 0 flux boundary conditions and achieved virtually identical results, indicating that the 100×100 μm^2^ was sufficiently large that boundary effects were insignificant. We conclude that VCell simulations accurately reproduce diffusive behavior and the Kang et al., 2012 (33) analysis is highly accurate in appropriate experimental settings.

**Figure 2.**
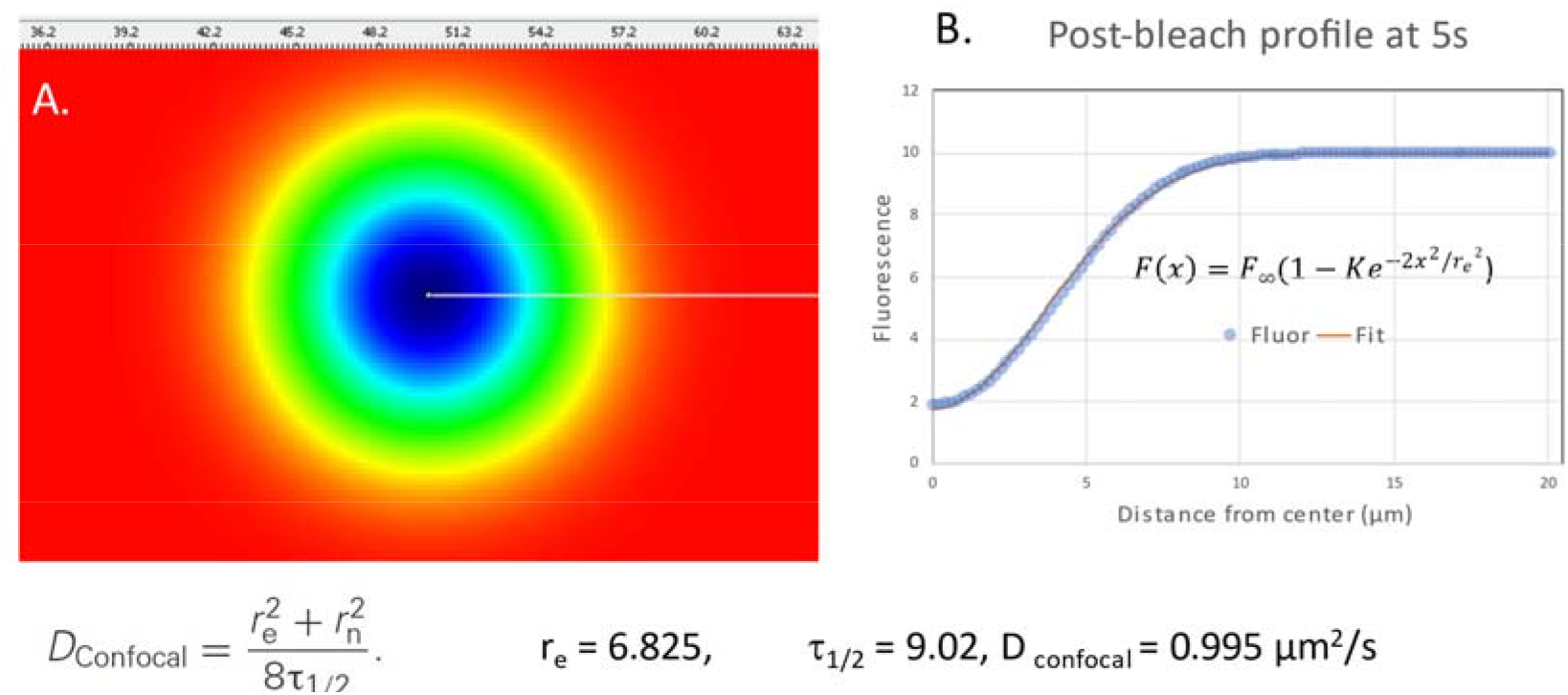
Simulations of FRAP within a 100 μm X 100 μm plane, analyzed by the methods of Kang et al. (33). A fluorescent species “fluor” with *D*=1 μm^2^/s was placed in the surface with boundary conditions clamped at a value of 10 μM. A 4 μm radius bleach area was defined in the center of the membrane plane (duration 1 s at *t*=1 s). A. Simulation snapshot at *t*=5 s (3 s post-bleach; see Supplemental Movie 1 for the full time course). B. Post-bleach profile along the line indicated in A. at *t*=5 s fit to a Gaussian to obtain r_e_. t_1/2_ was determined by finding the 50% recovery time from the 5s time point, using the mean fluorescence within the bleached 4μm radius bleached area.

To more closely represent an idealized cell membrane, we then created simulations using a 3D spherical surface of 20 μm diameter using analytic expressions to describe the geometry; it was placed within a cubic simulation domain of 22μm on each side. Bleach areas were defined using a Boolean expression to restrict the position of the “*Laser*” species to a circle of the chosen radius centered at x =11, y=11 and a Gaussian distribution in Z depending on the effective numerical aperture of the microscope optics.

The parameter *sigmaaxial* is used to specify the characteristic length of the Gaussian distribution in z, which is centered at the top of the spherical membrane (z=1); the parameter *BleachRadius* defines the radius of a circular scan region in x and y. Simulations were then run varying *D, duration* and *BleachRadius* under conditions where the bleach is highly focused (*sigmaaxial =* 1.5 μm) or when the bleaching beam can be approximated as a column in z (*sigmaaxial =* 30 μm). The former might be considered equivalent to a 2-photon bleaching profile, while the latter is a more conventional confocal or widefield profile and would bleach both the top and bottom surfaces of the cell. The VCell public model is “FRAP_Membrane_Rel” by user “les”; the Application within this model that simulates the spherical membrane is “Spherical_Cell_Circular_Bleach”. As the radius of the bleach region expands, one would expect that the analytic solution will diverge from the prescribed diffusion coefficient of 1μm^2^/s because the condition of an infinite source of fluorescent molecule is violated. As shown in Fig 3, we find that this is indeed the case; whereas with a *BleachRadius* of 1 μm the calculated diffusion coefficient is1.27 μm^2^/s; using a *BleachRadius* of 4 μm the, “D_Confocal_” is calculated to be 1.83μm^2^/s. Thus, for the larger bleach area, the assumption that the surface of the sphere can be approximately taken as infinite leads to less accurate estimation of *D*. This is our first indication that more accurate approximations may be obtained by iteratively matching multiple simulations with varying *D* to the experimental data, rather than attempting to solve the problem analytically.

**Figure 3.**
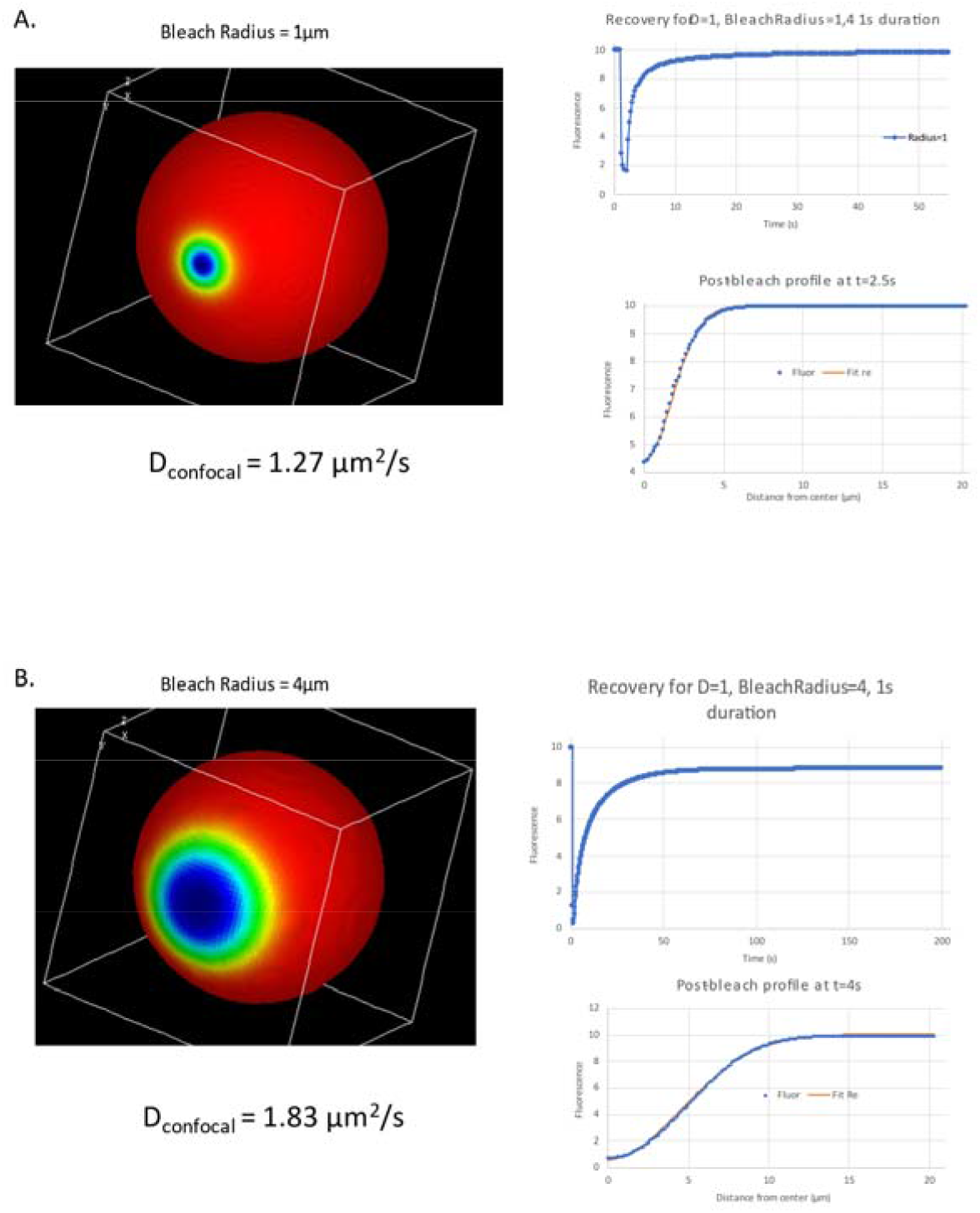
Analysis of FRAP on a spherical membrane surface with 1μm (A.) and 4μm (B.) bleach radius. The prescribed D for the Fluor species was 1μm^2^/s. Analysis of the recovery was performed as in Fig. 2, with the Gaussian fit to the post-bleach profiles performed at 2.5s (0.5s post-bleach) and 4s (2s post-bleach) for A. and B. respectively. See Supplemental Movie 2 for the time course of the simulation of FRAP with a 1 um bleach radius (A.) and Supplemental

### Numerical simulations of membrane FRAP experiments in cells with complex geometries

As shown above, the ideal analytical solutions of FRAP experiments becomes less accurate in situations where a significant fraction of the fluorescent molecules are bleached. We now provide an example of how the complex geometry of a real cell will further complicate the situation. Figure 4 shows a simulated photobleaching experiment based on a 3D confocal image of neuroblastoma cell. VCell includes tools for the import of image stacks and their segmentation to enable their use as geometries for simulations. For this work, the geometry was obtained from the VCell database of public models. Simulated bleaching of a circle in the center of cell, over the nucleus, reveals dynamics in the bleach region are largely the same whether only one membrane is bleached (i.e. as if using a bleaching beam in TIRF optics) or when both the top and bottom membrane of the cell are bleached (e.g. using a lower NA lens to approximate a bleaching area as a column through the cell). Applying the analysis of Kang (33) produces D of 1.34 and 1.21μm^2^/s for these 2 experimental approaches, compared to the input D of 1μm^2^/s. However, bleaching an equivalent area of a thin process leads to vastly slower redistribution curves in the bleach region, owing to the fact that fluorescent species from the bulk of the cell is restricted by having to enter the thin region that is small compared to the bleached area (34). In such cases the time course of redistribution can have profound effects on reaction kinetics within the process, as for example demonstrated by Brown et al., 2008 (35) in their analysis of PIP2 diffusion in dendritic spines. In this situation, using numerical simulation to match the time course for fluorescence redistribution is the most accurate way of both assessing diffusion and determining how different geometries affect reaction kinetics.

**Figure 4.**
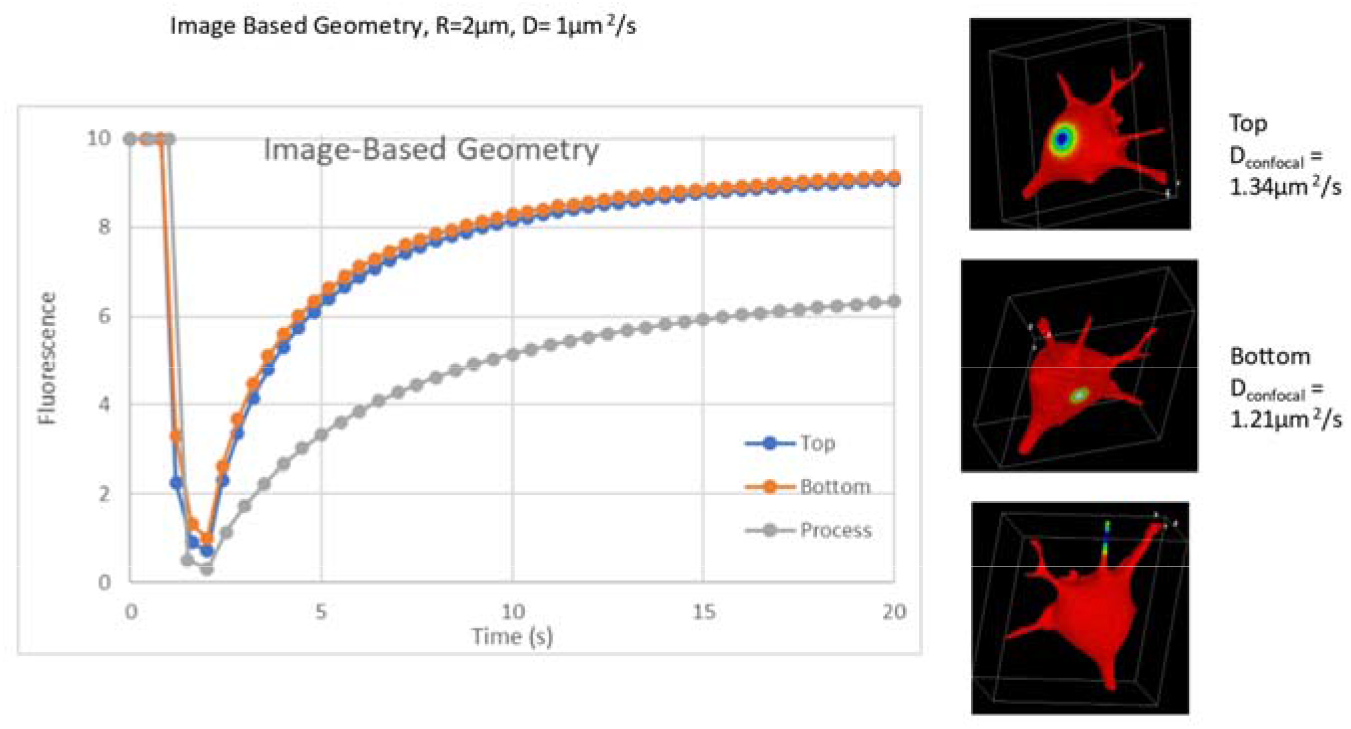
Recovery curves for membrane FRAP in 3 regions of an image-based geometry of a differentiating N1E neuroblastoma cell. The radius of of the “Laser” bleaching region is 2μm and the bleach starts at t=1s and ends at t=2s. D=1μm^2^/s. The top and bottom bleached region were analyzed as in Fig. 2. See Supplemental Movie 4-6 for the full time course of the top, bottom and process bleached region.

### Using numerical simulation to analyze diffusion and reaction in a volume

As noted by Kang et al., (33) analyzing diffusion within the cytosol using analytic approaches presents additional difficulties. Compared to the high viscosity membrane environment, free diffusion is much faster in the cytosol, and thus requires very fast collection of images and larger bleaching areas to allow one to capture the appropriate dynamics. To collect images rapidly, photobleaching experiments involving cytosolic components are often collected using widefield optics such that fluorescence from the entire cell is convolved in a single 2D image. However, the widefield results do not adhere to a 2-dimensional diffusion model, especially in the case of complex cellular geometries. Using the full cellular geometry (e.g. from confocal image stacks collected after collecting FRAP data in widefield) for simulation of the forward problem can be performed, and the simulation data flattened or convolved for direct comparison to experimental images or to Z-projections of confocal stacks. Using a larger geometry of the bleach region also increases the error from application of analytical solutions to derive D; and large organelles such as the nucleus can distort the free space available for diffusion. Numerical simulation of the forward problem can be used in these cases as well.

Additionally, binding to immobile or slowly moving components further complicates the analysis, as described previously (7, 9, 12, 36). Analytic solutions for assessing binding reactions using fluorescence photobleaching generally utilize different methods depending on the relative contributions of binding kinetics and diffusion time to the fluorescence redistribution(12, 9, 36). When binding is fast relative to diffusion time, fluorescence redistribution curves are dominated by diffusion – a diffusion-limited regime. At the other extreme, high affinity binding with slow off rates are “reaction-dominant” and yield curves that are well fit to a single exponential expression directly related to the off-rate. The real difficulty lies in cases where binding and diffusion time scales are similar. Again, solving the “forward problem” with parameter scans can be applied instead.

To illustrate the value of numerical simulation methods to analyze different experimental protocols and assess how parameters affect potential results, we created models for photobleaching of cytosolic components using the VCell Biomodel “FRAP_Cyt” by user “les”. These simulations utilized the same 3-dimensional cellular geometry used in the plasma membrane applications. The bleach region was specified by restricting the *Laser* as described for membrane bleaching; using a small “*sigmaaxial”* value can be used to simulate photobleaching using 2-photon excitation, and a larger “*sigmaaxial”* to approximate the much longer pattern of 1-photon excitation.

We focus here on 1-photon excitation, as would be common in a widefield or confocal microscope (additional simulations with “2-photon” Laser bleaching can be found in the VCell Biomodel “FRAP_Cyt”). The effect of different geometries and bleaching configurations on the dynamics of fluorescence intensity measured in the bleach region was assessed in both the absence and presence of binding reactions where one of the binding components is immobile. VCell provides the functionality to perform simulations in a 3-dimensional geometry, and then flatten all of the fluorescent species into a 2 dimensional distribution of “fluorescence” by either summing through z or convolution with a point spread function. Figure 5. shows such images of simulated fluorescence at the time point immediately post-bleach for two different bleaching protocols, either a small circular region 4 μm in diameter (Fig. 5A. Top), or bleaching an entire half of the cell (Fig. 5B. Top). The corresponding time courses for recovery are shown below each of these images. In these simulations, the total concentration of both binding sites, *Binder*, and fluorescent molecules, *Fluor*, are 10 μM, distributed between bound and free forms as determined by the *K*_*D*_; *Fluor* was given a diffusion coefficient of 25 μm^2^/s, while the binding sites (*Binder*) and complex (*FluorB*) were immobile. *k*_*off*_ for the binding reaction was varied from 0.01 s^-1^ to 10 s^-1^ and *k*_*on*_ was held constant at 1 μM ^-1^ s^-1^. Bleaching a 4 μm circle yields much different kinetics when *k*_*off*_ is slow, (0.1 or 0.01 s^-1^), but is indistinguishable from no binding when *k*_*off*_ is fast (1 and 10 s^-1^) (Figure 5A. Bottom). However, when half the cell is bleached, the time course for fluorescence redistribution is slower than with no binding even when *k*_*off*_ is fast (Figure 5B. Bottom). Thus, a larger bleach region is more effective at identifying lower affinity binding reactions in the presence of free diffusion, and can be effectively analyzed by numerical simulations using appropriate models. Other scenarios, such as holding the *K*_*D*_ constant while synchronously varying *k*_*on*_ *and k*_*off*_ or varying *D*, may be found in BioModel “FRAP_Cyt”.

**Figure 5.**
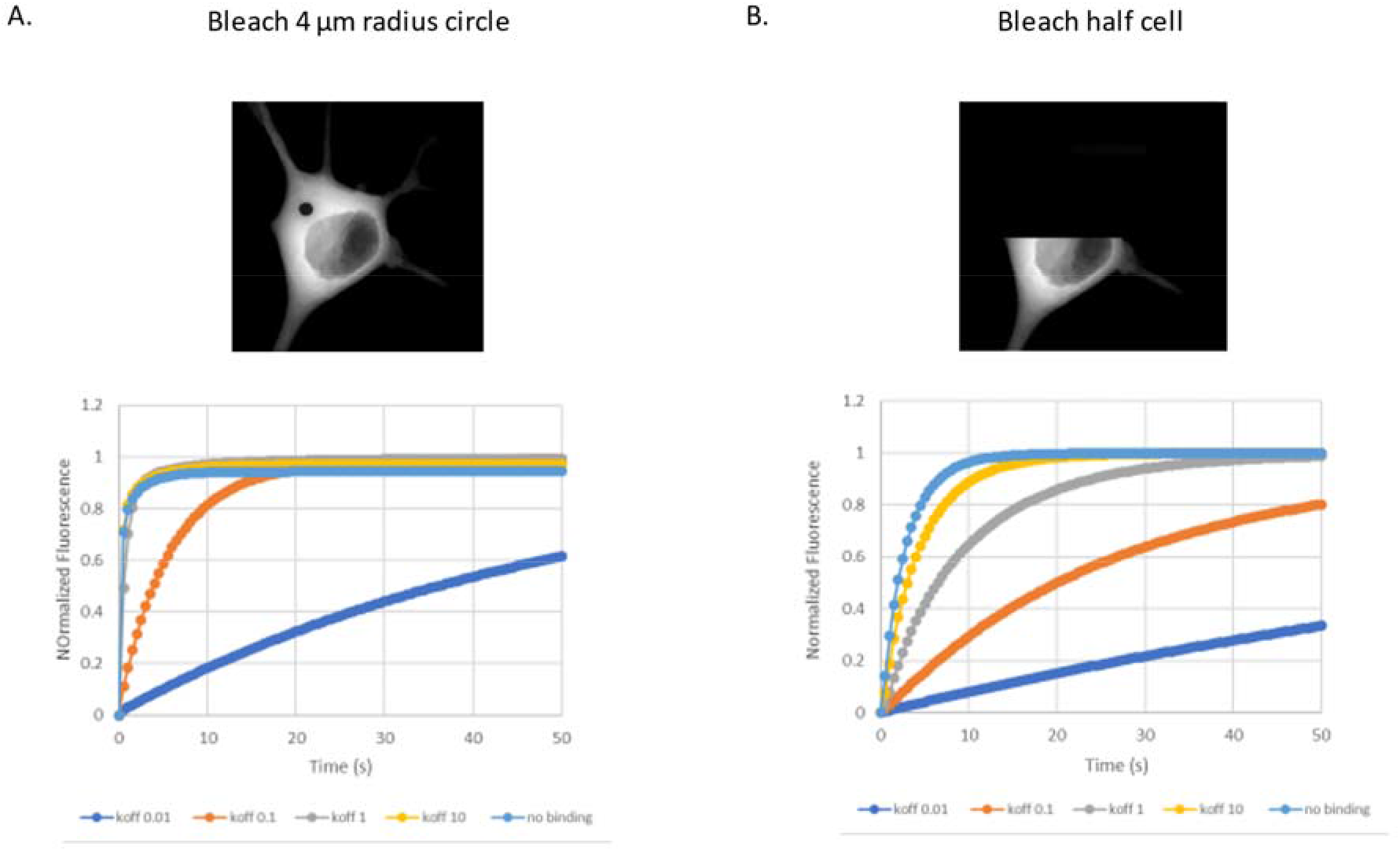
Simulated photobleaching of cytosolic species with immobile binding partners and varying *K*_*D*_. A. Top. “simulated fluorescence” created from a z-projection of convolved fluorescent species taken immediately after bleaching a 4 μm diameter region in the cytosol. The image is from a simulation where KD = 1 μM (Koff = 1 s-1). Simulated fluorescence can be created using a convolution function to approximate the experimental distribution of fluorescence molecules predicted from a simulation. See Supplemental Movie 7 for the full time course. Bottom shows normalized fluorescence curves from 5 different simulations where *K*_*D*_ is varied by changing *K*_*off*_ and measured within the entire initially bleached 4μm diameter region. B. Top, “simulated fluorescence” immediately after photobleaching the top half of the cell. See Supplemental Movie 8 for the full time course. Bottom, normalized fluorescence curves from 5 different simulations where *K*_*D*_ is varied by changing *K*_*off*_ while holding *K*_*on*_ constant. Average “fluorescence” in a 4 μm diameter circle within the bleached region was measured. Bleaching a small circular bleach makes it difficult to identify low affinity binding interactions; bleaching a much larger region (e.g. half the cell) reveals very different time courses even with fast off rates. All simulations are in VCell BioModel “FRAP_Cyt”; The Applications corresponding to A. and B. are, respectively: “ProjectZ Convolved Image-based center circular bleach vary Kd” and “ProjectZ Convolved Image-based half bleach vary Kd”

### Diffusion and binding in liquid droplets

Biomolecular condensates, also known as liquid droplets or membraneless organelles, are sub-cellular aggregates of weakly binding proteins and/or nucleic acids. The biology of liquid droplets has developed explosively because they are so ubiquitous and appear to organize many cellular functions (37–39). Their biophysical properties can be understood within the thermodynamic framework of phase separation as well as polymer physics (40, 41) Experimentally, FRAP has been a valuable tool to elucidate the dynamics of liquid droplets. Indeed, in a classic paper (42), FRAP was critical to demonstrate that “P granules” are liquid droplets, showing that they localize to one end of a germ-line cell by dissolving and recondensing. However, the complexity of the multiple processes that govern the results of a FRAP experiment (binding, diffusion in the cytosol, diffusion within the droplet) makes interpretation difficult and can even lead to faulty conclusions (43). In this section, we exemplify how modeling and simulation of virtual FRAP experiments can help to dissect the key biophysical properties of liquid droplets.

We use the same simple FRAP with binding reaction scheme in the cytosol as described in Fig. 6. However, instead of having the “Binder” evenly distributed within the cytosol, we confine it to a spherical “droplet” of 4μm diameter situated just outside the nucleus (Fig. 6A. top, center); the total concentration of binding sites is 10μM within the droplet leading to a steady state pre-bleach concentration of *Fluor* = *Binder* = 2.70 μM and *FluorB* = 7.30 μM. The Z-projection images shown in the Figure appears to show the droplet within the nucleus, but it is actually well above the nucleus in the axial direction). The simulated experiment in Fig. 6A. models a 1s bleach duration at t = 1 s with the *Laser* positioned to exactly encompass the entire droplet. We display the resulting recovery curves for 2 parameter sets designed to bracket a broad range of on and off rates for the binding of the free fluorophore to binding sites in the droplet (maintaining a Kd of 1 μM); we also use diffusion coefficients for the free fluorophore designed to bracket values for very large and small proteins in cytosol (*D*=3 μm^2^/s and 25 μm^2^/s, respectively). The 4 combinations of these parameters produce distinctive recovery curves when measured from within droplet, while the recovery in an adjacent region (Fig. 6A. inset) is primarily sensitive to the diffusion of the free fluorophore. Monitoring the recovery at a greater distance from the droplet (data not shown, but accessible from the public VCell model) would produce a recovery curve that is even less influenced by the binding rate, albeit with a smaller dynamic range. Thus it should be possible to use parameter scans to find a correspondence with experimental results to determine both binding kinetics and free fluorophore diffusion. These simulation results and additional examples of parameter combinations can be found in the VCell model “FRAP_Cyt”, Application: “ProjectZ Image-based binding to liquid droplet”.

**Figure 6.**
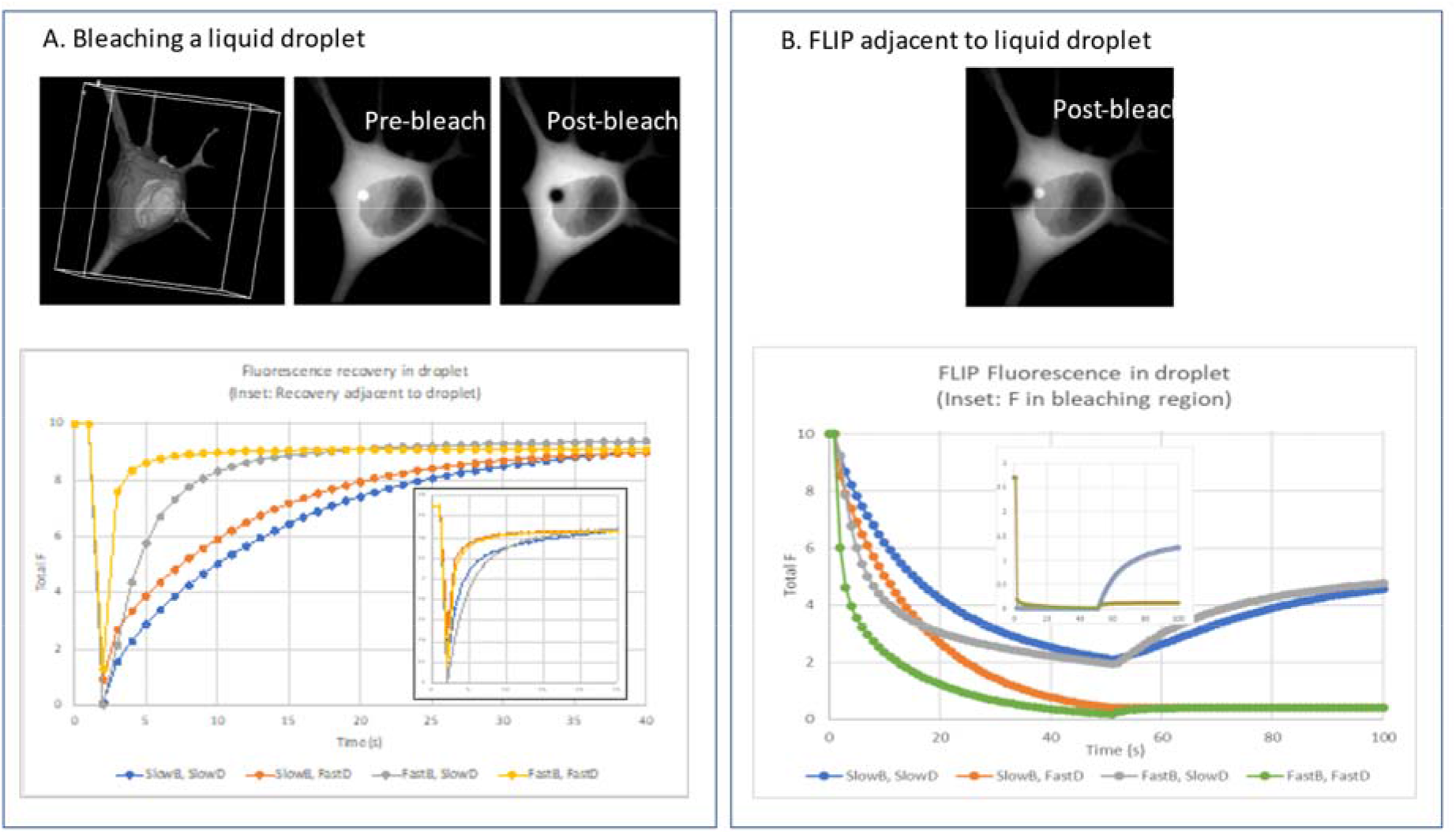
Photobleaching simulations with binding to a liquid droplet. A. Bleaching over the liquid droplet. The 3D image-based geometry of the cell is shown at the upper left as a surface rendering, with a translucent plasma membrane to allow visualization of the nucleus. Upper center and right images are Z-projections of the total fluorescence species (sum of free and bound), before and immediately after the bleach; The 4μm diameter spherical liquid droplet is positioned above the nucleus; the Laser is modeled as a 4μm diameter cylinder positioned through the liquid droplet. The plots at the bottom correspond to different combinations of slow (Kr= 0.1 s^-1^, Kf = 0.1 μM^-1^s^-1^) and fast (Kr= 100 s^-1^, Kf = 100 μM^-1^s^-1^) binding kinetics, with slow (D=3 μm^2^/s) and fast (D=25 μm^2^/s) diffusion of the unbound Fluor and Dark species. The primary plot is the change in TotalF (free + bound) at the center of the liquid droplet; the inset shows the Fluor species 1 μm to the left of the liquid droplet. See Supplemental Movies 9 - 12 for the full simulations, respectively. B. Flip adjacent to the liquid droplet. The Laser species is a cylinder of diameter 8μm placed just outside the spherical liquid droplet, as illustrated in the first post-bleach Z-projection at the top. The bleaching reaction is maintained between t=1 s and t=50 s, with recovery allowed between t=50 s and t=100 s. The large plot shows TotalF in the center of the liquid droplet, with the 4 parameter sets defined as in A.; the inset shows the correspond change in the free Fluor species in the center of the bleached area. See Supplemental Movies 13-16 for the full simulation of each parameter set.

Fig. 6B. is based on simulations in “FRAP_Cyt”, Application: “ProjectZ Image-based binding to liquid droplet, FRAP adjacent to droplet”. In these virtual experiments, we bleached an 8 μm diameter cylindrical column immediately adjacent to the droplet for 49 s while measuring the fluorescence both inside the droplet and (Fig. 6B. inset) in the center of the bleached volume for 100 s. These simulations model a “Fluorescence loss in photobleaching” (FLIP) experiment, where the depletion of fluorescent label outside the area of interest probes the off rate from the droplet. We explore the same combination of slow and fast rates and slow and fast diffusion parameters as in Fig. 6A. As can be seen in the inset to Fig. 6B., the fluorescence species within the bleaching region remain entirely depleted throughout the 49 s bleaching time.

However, the *TotalF* (free plus bound fluorescent species) within the droplet is depleted at a slower rate, with the 2 conditions corresponding to the D= /s plateauing at around 2 μM right before bleaching is terminated, followed by a recovery; for D=25 μm^2^/s, 49 s is sufficient to almost completely deplete all the fluorescent species in the cell, allowing for little recovery. Within the bleached region (inset), the behavior is completely determined by diffusion: the gray and blue traces are superimposed as are the green and orange traces. Thus, these virtual FLIP experiments point to how diffusion and binding kinetics can be experimentally dissected by comparing experiments to simulations.

Diffusion within the droplet is a key hallmark of its liquid nature. An established experiment to probe for diffusion within the droplet is to FRAP half of it (42, 43). The fastest and most accurate measurement technique is to acquire confocal line scans through the middle of the droplet to build a kymograph. However, as explored in Fig. 7., the recovery is governed by both diffusion from the unbleached half and the kinetics of exchange with the free fluorescent species. We compared simulated kymographs where the bound species are immobile (*Db* = 0) and with a diffusion coefficient corresponding to a large protein in a viscous medium (*Db* = 1 μm^2^/s); in all cases, we kept the diffusion coefficient of the free fluorophore at *Df* = 3 μm^2^/s, but these simulations are relatively insensitive to *Df*. When the binding kinetics are sufficiently slow (Fig 7B.) the effect of diffusion within the droplet is apparent in the kymographs: when *Db* = 0, the boundary between bleached and unbleached regions is maintained, with recovery in the bleached region completely dependent on the kinetics of exchange with free fluorophore; when *Db* = 1 μm^2^/s, the boundary is erased within ∼1 s following the bleach, and there is depletion of fluorescence from the unbleached half of the droplet. Importantly, while *Db* is relatively low at 1 μm^2^/s, the small size of the droplet allows for rapid diffusional equilibration. If an experiment and a corresponding simulation were run on larger droplets, a perceptible boundary would be maintained for longer periods. Turning to the case of fast exchange kinetics in Fig. 7C., we see that the effect of diffusion within the droplet is much more subtle. While the kymographs are almost indistinguishable, there is a difference that can be detected at individual time points close to the end of the bleach. The images are XY slices through the center of the droplet at time 2.4 s, i.e. 400 ms post bleach; the corresponding line scans are shown at the bottom of Fig. 7C. While the differences are small, the diffusion rates should be determinable if the kinetic constants for exchange are known. The latter may be determined by results of the analysis described above for Fig 6.

**Figure 7.**
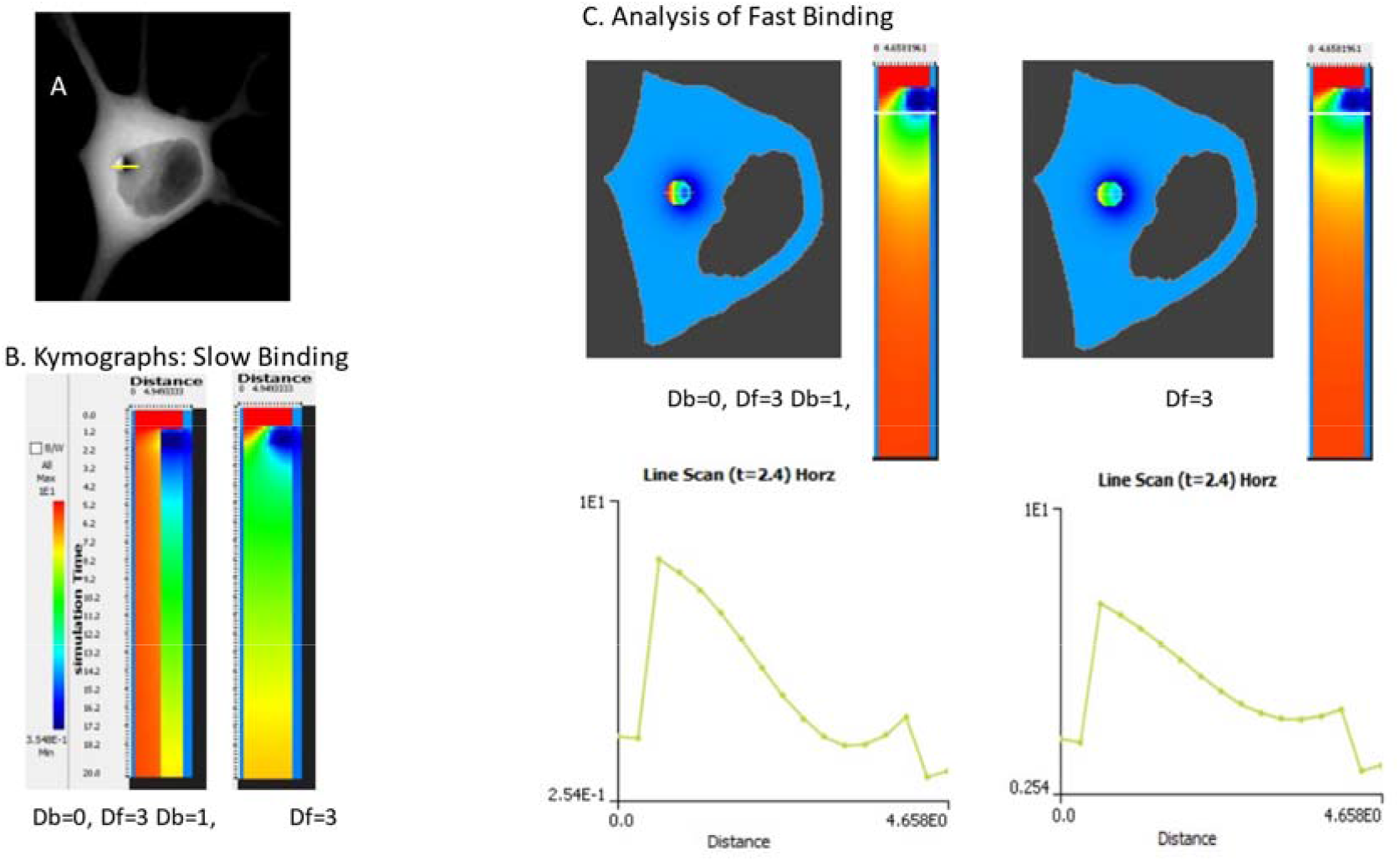
Analysis of diffusion with the liquid droplet by half-bleach. A. Z-projection of the first post-bleach simulation point for half-bleach of the liquid droplet with the Laser modeled as a hemi-cylinder positioned over the spherical droplet; a line through the droplet indicates where the kymograph analysis is generated. B. Kymographs for a line scan spanning the center of the liquid droplet and lasting 20s with slow binding kinetics (Kr= 0.1s^-1^, Kf = 0.1μM^-1^s^-1^). The Kymograph on the left shows results with 0.0 D within the droplet, and D=3μm^2^/s for the free unbound species. The kymograph on the right shows the pattern when D within the liquid droplet is 1 μm^2^/s. See Supplemental Movies 17 and 18, respectively, for the full time course. C. More subtle differences are observed when binding is fast (Kr= 100 s^-1^, Kf = 100 μM^-1^s^-1^). Results for liquid droplet diffusion of Db= 0 on the left and Db=1 μm^2^/s on the right. The slices at the top and the line scans on the bottom are all at 400 ms post-bleach (t=2.4 s). See Supplemental Movies 19 and 20, respectively for the full time course.

## Discussion

In this work, we show that numerical solutions to partial differential equations (PDEs) can help guide the analysis of the reaction-diffusion dynamics revealed by FRAP experiments. We were motivated to create these mathematical models because analytical solutions or numerical parameter estimation algorithms for derivation of diffusion coefficients apply only to idealized cell and bleached region geometries. If binding kinetics are also involved, there must additionally be a good separation between the timescales of diffusion and reaction rates. On the other hand, numerical solutions of PDEs can be obtained using experimentally derived cell geometries and bleaching parameters that match the FRAP experiment. They can also include models of any binding reaction kinetics involving the fluorescently labeled molecule.

Determining the values of the diffusion and kinetic constants can then be accomplished by running parameter scans and comparing the simulation results to the experiments. Our results were presented for a wide bracket of parameter values; in practice, these can be iteratively narrowed to find the parameter set to best match the experimental results. Of course, if the experiments cannot be made to fit the simulations, the assumptions underlying the kinetic model may be incorrect. Such an outcome could actually produce new insights into the underlying biophysical mechanisms.

We examined membrane diffusion, diffusion in a volume, diffusion with binding and FRAP experiments on liquid droplets. We first (Fig. 2) used a simple model of 2D diffusion in a plane to show that analysis of the simulation results with a well-established fitting method (33). can precisely recover the value of D that was input into the model, even when using different sized bleaching areas. However, when using more realistic finite geometries, the fitting method tends to somewhat overestimate D (Figs. 3 and 4), while it is not at all applicable to determining diffusion on a thin process (Fig. 4). Turning to diffusion with binding (Fig. 5), we showed how the model can actually help to design the best FRAP experimental protocol to best differentiate diffusional rates from off rates when the Kd is also unknown. For the case of liquid droplets (Figs. 6 and 7), the additional process of diffusion within the droplet must be dissected from binding and diffusion of the free fluorophore; we show that this can be accomplished by the use of 3 protocols: FRAP of the entire droplet, FLIP on a region adjacent to the droplet, and the analysis of kymographs from line scans across a half-bleached droplet. Other simulations and models exploring various geometries and parameter combinations can be found in the VCell public models “FRAP_Cyt” and “FRAP_Membrane_Rel”.

While these various FRAP protocols are not at all exhaustive, they can serve as a guide for the construction of any model of a FRAP experiment. Additionally, there are issues associated with fluorescence microscopy that could be readily added to more accurately simulate an experiment. It would be straightforward, for example, to add a constant slow bleach process to these models to simulate “bleach while monitoring”, or to add a process to account for unquenching occurring during the bleach. There are also other features of VCell that may be exploited to produce realistic models of FRAP experiments. For example, the distribution of binding sites may be non-uniform; or in the case of liquid droplets, there may be multiple droplets with variable size and shape throughout the cell. Images of these can be used in a VCell feature called “Field Data” to define the spatially heterogenous distribution of initial concentrations and their diffusion coefficients. Fig. 5 used ImageJ to help analyze simulation results; VCell has some integration with ImageJ to help both automated incorporation of experimental data into models and more advanced analysis of simulation results. Finally, it is important to emphasize that the modeling techniques described here for FRAP and FLIP may readily be applied to other photochemistry-based microscope experiments, including photoactivation of fluorescence (44), uncaging of signaling molecules (45), photoablation of labeled proteins (46), and optogenetic activation of signaling proteins (47).

## Supporting information

Full details of VCell BioModel "FRAP_Cyt"

Full details of VCell BioModel "FRAP_Membrane_Rel"

XML-encoded VCell BioModel "FRAP_Cyt"

XML-encoded VCell BioModel "FRAP_Membrane_Rel"

## Author Contributions

Both LML and AEC conceived the research, built the models, performed simulations and wrote and edited the manuscript.

## Acknowledgements

The Virtual Cell software is supported by National Institutes of Health Grant R24-GM137787 from the National Institute of General Medical Sciences. We thank the VCell team, especially Boris Slepchenko, for helpful discussions. We also wish to acknowledge the contributions of our dear friend Ken Jacobson, who was a valued collaborator and served for over 20 years as a key member of the Scientific Advisory Committee for the Virtual Cell Project; this paper is dedicated to his memory.

## Supplemental Materials

FRAP_Cyt.pdf and FRAP_Membane_Rel.pdf Complete reports on the details of the VCell models used in this work.

FRAP_Cyt.vcml and FRAP_Membane_Rel.vcml Complete readable code for the 2 VCell BioModels used in this work. These models are accessible together will all the simulations in the VCell database.

## Supplemental Movies

Supplemental Movie 1. Simulation of a 100 × 100 Planar Membrane with a 4μm diameter bleach (Figure 2).

Supplemental Movie 2. Spherical membrane based simulation of FRAP with a 1 μm bleach radius (Figure 3A.).

Supplemental Movie 3. Spherical membrane based simulation of FRAP with a 4 μm bleach radius time course (Figure 3B.).

Supplemental Movie 4. Image based simulation membrane FRAP circular bleach top membrane (Figure 4)

Supplemental Movie 5. Image based simulation membrane FRAP circular bleach bottom membrane (Figure 4).

Supplemental Movie 6. Image based simulation membrane FRAP circular process membrane (Figure 4).

Supplemental Movie 7. “Simulated fluorescence” time course for cytosolic fluorophore binding to an immobile partner with kD = 1: 4 μm diameter bleach (Figure 5A.).

Supplemental Movie 8. “Simulated fluorescence” time course for cytosolic fluorophore binding to an immobile partner with kD = 1: half cell bleach (Figure 5B.).

Supplemental Movie 9. Single z section simulation of bleach liquid droplet D=3 slow binding (Figure 6A.)

Supplemental Movie 10. Single z section simulation of bleach liquid droplet D=3 fast binding (Figure 6A.)

Supplemental Movie 11. Single z section simulation of bleach liquid droplet D=25 slow binding (Figure 6A.)

Supplemental Movie 12. Single z section simulation of bleach liquid droplet D=25 fast binding (Figure 6A.)

Supplemental Movie 13. Single z section simulation of bleach near liquid droplet D=3 slow binding (Figure 6B.)

Supplemental Movie 14. Single z section simulation of bleach near liquid droplet D=3 fast binding (Figure 6B.)

Supplemental Movie 15. Single z section simulation of bleach near liquid droplet D=25 slow binding (Figure 6B.)

Supplemental Movie 16. Single z section simulation of bleach near liquid droplet D=25 fast binding (Figure 6B.)

Supplemental Movie 17. Single z section simulation of bleach half liquid droplet D=0 in droplet slow binding (Figure 7B.)

Supplemental Movie 18. Single z section simulation of bleach half liquid droplet D=1 in droplet slow binding (Figure 7B.)

Supplemental Movie 19. Single z section simulation of bleach half liquid droplet D=0 in droplet fast binding (Figure 7C.)

Supplemental Movie 20. Single z section simulation of bleach half liquid droplet D=1 in droplet fast binding (Figure 7C.)

