## Supplementary material for "Beyond analytic solution: analysis of FRAP experiments by spatial simulation of the forward problem": Full details of VCell BioModel "FRAP_Cyt"

### 1. Physiology For FRAP_Cyt

#### 1.1. General Info

| |
|---|
| **BioModel Name: FRAP_Cyt** |
| **Owner: les** |

#### 1.2. Structures and Reactions Diagram

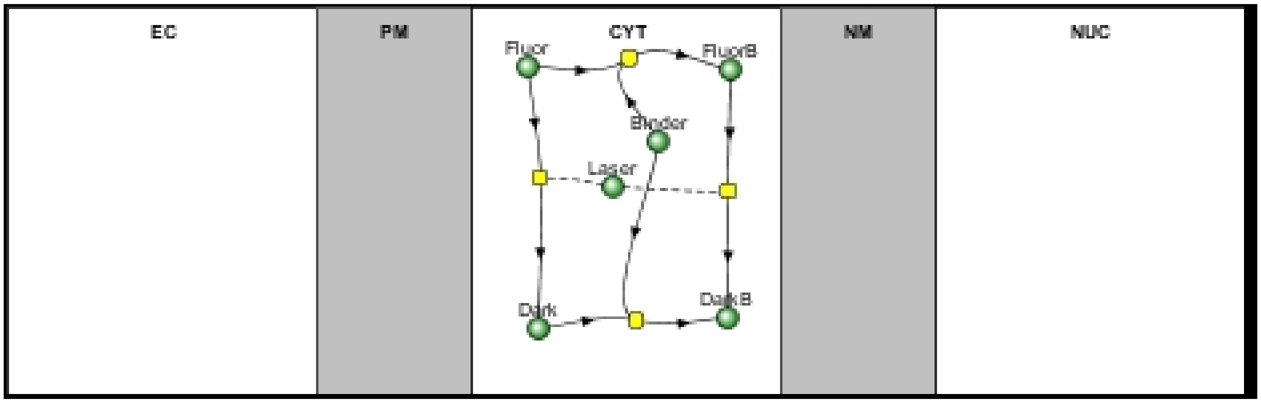

#### 1.3. Reaction(s) in CYT

##### 1.3.1. Reaction BindD

**Dark + Binder ⟷ DarkB**

| Kinetics Parameters | | | |
|---|---|---|---|
| **Name** | **Expression** | **Role** | **Unit** |
| J | (((Kf * Dark) * Binder) - (Kr * DarkB)) | reaction rate | $M.s^{1}$ |
| Kf | 1.0 | forward rate constant | $s^{1}.M^{1}$ |
| Kr | 1.0 | reverse rate constant | $s^{1}$ |

##### 1.3.2. Reaction BindF

**Fluor + Binder ⟷ FluorB**

| Kinetics Parameters | | | |
|---|---|---|---|
| **Name** | **Expression** | **Role** | **Unit** |
| J | (((Kf * Fluor) * Binder) - (Kr * FluorB)) | reaction rate | M.s$^1$ |
| Kf | 1.0 | forward rate constant | s$^1$.M$^1$ |
| Kr | 1.0 | reverse rate constant | s$^1$ |

#### 1.3.3. Reaction BleachFree

**Fluor** ⟷ **Dark**

| Modifiers List |
|---|
| Laser |

| Kinetics Parameters | | | |
|---|---|---|---|
| **Name** | **Expression** | **Role** | **Unit** |
| J | ((Kf * Fluor) - (Kr * Dark)) | reaction rate | M.s$^1$ |
| Kf | ((kbleach * Laser) * (t > start) * (t < (start + duration))) | forward rate constant | s$^1$ |
| Kr | 0.0 | reverse rate constant | s$^1$ |
| kbleach | 1.0 | user defined | s$^1$.M$^1$ |

#### 1.3.4. Reaction BleachBound

**FluorB** ⟷ **DarkB**

| Modifiers List |
|---|
| Laser |

| Kinetics Parameters | | | |
|---|---|---|---|
| **Name** | **Expression** | **Role** | **Unit** |

| Kinetics Parameters | | | |
|---|---|---|---|
| **Name** | **Expression** | **Role** | **Unit** |
| J | ((Kf * FluorB) - (Kr * DarkB)) | reaction rate | $M.s^{1}$ |
| Kf | ((kbleach * Laser) * (t > start) * (t < (start + duration))) | forward rate constant | $s^{1}$ |
| Kr | 0.0 | reverse rate constant | $s^{1}$ |
| kbleach | 1.0 | user defined | $s^{1}.M^{1}$ |

### 2. Applications For FRAP_Cyt

#### 2.1. Application: Spherical_Cell_Gaussian_Bleach

| Application Name: Spherical_Cell_Gaussian_Bleach |
|---|

##### 2.1.1. Structure Mapping For Spherical_Cell_Gaussian_Bleach

| Structure Mapping | | | | |
|---|---|---|---|---|
| **Structure** | **Subdomain** | **Resolved (T/F)** | **Surf/Vol** | **VolFract** |
| EC | subdomain0 | F | | |
| NUC | subdomain1 | F | | |
| CYT | subdomain1 | F | | |

##### 2.1.2. Reaction Mapping For Spherical_Cell_Gaussian_Bleach

| Reaction Mapping | | | |
|---|---|---|---|
| **Name** | **Type** | **Enabled (T/F)** | **Fast (T/F)** |
| BindD | Reaction | T | F |
| BindF | Reaction | T | F |
| BleachFree | Reaction | T | F |
| BleachBound | Reaction | T | F |

| Initial Conditions | | | | |
|---|---|---|---|---|
| **Species** | **Structure** | **Initial Conc.** | **Diffusion Const.** | **Fixed (T/F)** |
| s0 | CYT | 0.0 M | 0.0 $m^2.s^1$ | F |
| s1 | CYT | 0.0 M | 0.0 $m^2.s^1$ | F |
| s2 | CYT | 0.0 M | 0.0 $m^2.s^1$ | F |

| Initial Conditions | | | | |
|---|---|---|---|---|
| **Species** | **Structure** | **Initial Conc.** | **Diffusion Const.** | **Fixed (T/F)** |
| s3 | CYT | 0.0 M | 0.0 $m^{2}.s^{1}$ | F |
| s4 | CYT | 0.0 M | 0.0 $m^{2}.s^{1}$ | F |
| s5 | CYT | 0.0 M | 0.0 $m^{2}.s^{1}$ | F |

#### 2.1.3. Membrane Mapping For Spherical_Cell_Gaussian_Bleach

| Electrical Mapping - Membrane Potential | | | |
|---|---|---|---|
| **Membrane** | **Calculate V (T/F)** | **V initial** | **Specific Capacitance** |
| PM | F | 0.0 mV | 1.0 $pF.m^{2}$ |
| NM | F | 0.0 mV | 1.0 $pF.m^{2}$ |

| |
|---|
| Temperature: 300.0 K |

#### 2.1.4. Geometry: Geometry343793035

| | |
|---|---|
| Size | (22.0, 22.0, 22.0) |
| Origin | (0.0, 0.0, 0.0) |

#### 2.1.5. Math Description: Spherical_Cell_Gaussian_Bleach_generated

##### 2.1.5.1. Constants

| Constant Name | Expression |
|---|---|
| _F_ | 96485.3321 |
| _F_nmol_ | 9.64853321E-5 |
| _K_GHK_ | 1.0E-9 |
| _N_pmol_ | 6.02214179E11 |
| _PI_ | 3.141592653589793 |

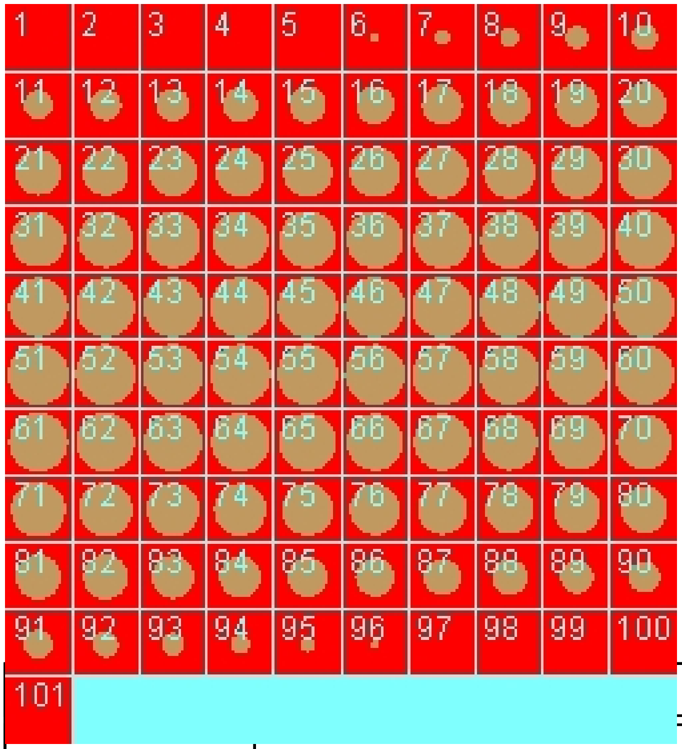

| Constant Name | Expression |
|---|---|
| _R_ | 8314.46261815 |
| _T_ | 300.0 |
| AreaPerUnitArea_PM | 1.0 |
| AreaPerUnitVolume_NM | 1.2 |
| Binder_init_uM | 0.0 |
| BleachRadius | 2.0 |
| Dark_init_uM | 0.0 |
| DarkB_init_uM | 0.0 |
| duration | 1.0 |
| Fluor_init_uM | 0.0 |
| FluorB_init_uM | 0.0 |
| K_millivolts_per_volt | 1000.0 |
| kbleach_BleachBound | 1.0 |
| kbleach_BleachFree | 1.0 |

| Constant Name | Expression |
|---|---|
| Kf_BindD | 1.0 |
| Kf_BindF | 1.0 |
| KMOLE | 0.001660538783162726 |
| Kr_BindD | 1.0 |
| Kr_BindF | 1.0 |
| Kr_BleachBound | 0.0 |
| Kr_BleachFree | 0.0 |
| Laser_init_uM | 0.0 |
| sigmaaxial | 1.5 |
| sigmalateral | 0.5 |
| start | 1.0 |
| Voltage_NM | 0.0 |
| Voltage_PM | 0.0 |
| VolumePerUnitVolume_CYT | 1.0 |
| VolumePerUnitVolume_EC | 1.0 |
| VolumePerUnitVolume_NUC | 65.45 |

##### 2.1.5.2. Functions

| Function Name | Expression |
|---|---|
| J_BindD | (((Kf_BindD * Dark) * Binder) - (Kr_BindD * DarkB)) |
| J_BindF | (((Kf_BindF * Fluor) * Binder) - (Kr_BindF * FluorB)) |
| J_BleachBound | ((Kf_BleachBound * FluorB) - (Kr_BleachBound * DarkB)) |
| J_BleachFree | ((Kf_BleachFree * Fluor) - (Kr_BleachFree * Dark)) |
| Kf_BleachBound | ((kbleach_BleachBound * Laser) * (t > start) * (t < (start + duration))) |
| Kf_BleachFree | ((kbleach_BleachFree * Laser) * (t > start) * (t < (start + duration))) |
| Size_CYT | (VolumePerUnitVolume_CYT * vcRegionVolume('subdomain1')) |

| Function Name | Expression |
|---|---|
| Size_EC | (VolumePerUnitVolume_EC * vcRegionVolume('subdomain0')) |
| Size_NM | (AreaPerUnitVolume_NM * vcRegionVolume('subdomain1')) |
| Size_NUC | (VolumePerUnitVolume_NUC * vcRegionVolume('subdomain1')) |
| Size_PM | (AreaPerUnitArea_PM * vcRegionArea('subdomain0_subdomain1_membrane')) |
| sobj_subdomain11_subdomain00_size | vcRegionArea('subdomain0_subdomain1_membrane') |
| vobj_subdomain00_size | vcRegionVolume('subdomain0') |
| vobj_subdomain11_size | vcRegionVolume('subdomain1') |

#### 2.1.5.3. Volume Domains

##### 2.1.5.3.1. subdomain1

| **OdeEquation Fluor** | |
|---|---|
| Rate | ( - J_BindF - J_BleachFree) |
| Initial | Fluor_init_uM |

| **OdeEquation Dark** | |
|---|---|
| Rate | ( - J_BindD + J_BleachFree) |
| Initial | Dark_init_uM |

| **OdeEquation Binder** | |
|---|---|
| Rate | ( - J_BindD - J_BindF) |
| Initial | Binder_init_uM |

| **OdeEquation FluorB** | |
|---|---|
| Rate | (J_BindF - J_BleachBound) |
| Initial | FluorB_init_uM |

| OdeEquation Laser | |
|---|---|
| Rate | 0.0 |
| Initial | Laser_init_uM |

| OdeEquation DarkB | |
|---|---|
| Rate | (J_BindD + J_BleachBound) |
| Initial | DarkB_init_uM |

###### 2.1.5.3.2. subdomain0

##### 2.1.5.4. Membrane Domains

###### 2.1.5.4.1. subdomain0_subdomain1_membrane

##### 2.1.6. Simulation(s)

#### 2.2. Application: TIRF_center_bleach

| **Application Name: TIRF_center_bleach** |
|---|
| **Application Description: (copied from Spherical_Cell_Gaussian_Bleach)** |

##### 2.2.1. Structure Mapping For TIRF_center_bleach

| **Structure Mapping** | | | | |
|---|---|---|---|---|
| **Structure** | **Subdomain** | **Resolved (T/F)** | **Surf/Vol** | **VolFract** |
| EC | EC | F | | |
| NUC | IC | F | | |
| CYT | IC | F | | |

##### 2.2.2. Reaction Mapping For TIRF_center_bleach

| Reaction Mapping | | | |
|---|---|---|---|
| **Name** | **Type** | **Enabled (T/F)** | **Fast (T/F)** |
| BindD | Reaction | T | F |
| BindF | Reaction | T | F |
| BleachFree | Reaction | T | F |
| BleachBound | Reaction | T | F |

| Initial Conditions | | | | |
|---|---|---|---|---|
| **Species** | **Structure** | **Initial Conc.** | **Diffusion Const.** | **Fixed (T/F)** |
| s0 | CYT | 0.0 M | 0.0 $m^2.s^1$ | F |
| s1 | CYT | 0.0 M | 0.0 $m^2.s^1$ | F |
| s2 | CYT | 0.0 M | 0.0 $m^2.s^1$ | F |
| s3 | CYT | 0.0 M | 0.0 $m^2.s^1$ | F |
| s4 | CYT | 0.0 M | 0.0 $m^2.s^1$ | F |
| s5 | CYT | 0.0 M | 0.0 $m^2.s^1$ | F |

#### 2.2.3. Membrane Mapping For TIRF_center_bleach

| Electrical Mapping - Membrane Potential | | | |
|---|---|---|---|
| **Membrane** | **Calculate V (T/F)** | **V initial** | **Specific Capacitance** |
| PM | F | 0.0 mV | 1.0 $pF.m^2$ |
| NM | F | 0.0 mV | 1.0 $pF.m^2$ |

| |
|---|
| Temperature: 300.0 K |

#### 2.2.4. Geometry: Geometry7

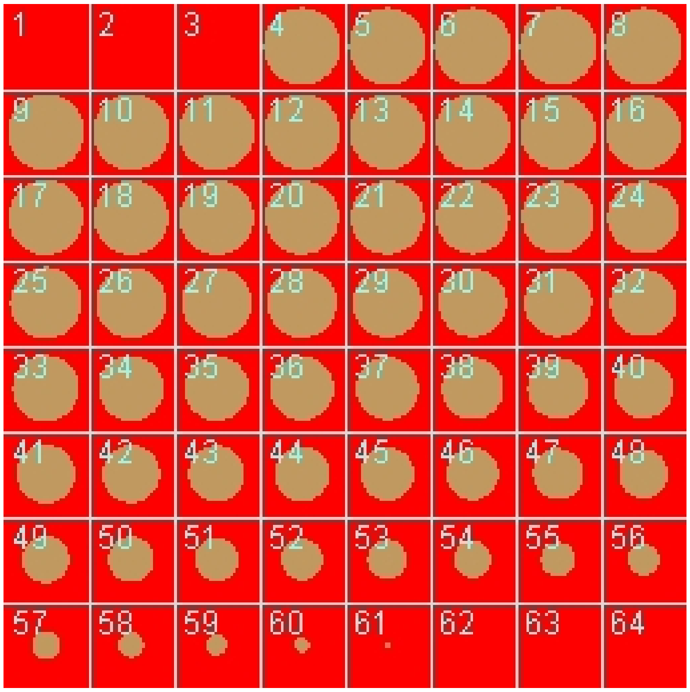

| | |
|---|---|
| Size | (22.0, 22.0, 11.0) |
| Origin | (-11.0, -11.0, -0.5) |

#### 2.2.5. Math Description: Copy of Spherical_Cell_Gaussian_Bleach_generated

##### 2.2.5.1. Constants

| **Constant Name** | **Expression** |
|---|---|
| _F_ | 96485.3321 |
| _F_nmol_ | 9.64853321E-5 |
| _K_GHK_ | 1.0E-9 |
| _N_pmol_ | 6.02214179E11 |
| _PI_ | 3.141592653589793 |
| _R_ | 8314.46261815 |
| _T_ | 300.0 |
| AreaPerUnitArea_PM | 1.0 |

| Constant Name | Expression |
|---|---|
| AreaPerUnitVolume_NM | 1.2 |
| Binder_init_uM | 0.0 |
| BleachRadius | 2.0 |
| Dark_init_uM | 0.0 |
| DarkB_init_uM | 0.0 |
| duration | 1.0 |
| Fluor_init_uM | 0.0 |
| FluorB_init_uM | 0.0 |
| K_millivolts_per_volt | 1000.0 |
| kbleach_BleachBound | 1.0 |
| kbleach_BleachFree | 1.0 |
| Kf_BindD | 1.0 |
| Kf_BindF | 1.0 |
| KMOLE | 0.001660538783162726 |
| Kr_BindD | 1.0 |
| Kr_BindF | 1.0 |
| Kr_BleachBound | 0.0 |
| Kr_BleachFree | 0.0 |
| Laser_init_uM | 0.0 |
| sigmaaxial | 1.5 |
| sigmalateral | 0.5 |
| start | 1.0 |
| Voltage_NM | 0.0 |
| Voltage_PM | 0.0 |
| VolumePerUnitVolume_CYT | 1.0 |
| VolumePerUnitVolume_EC | 1.0 |

| Constant Name | Expression |
|---|---|
| VolumePerUnitVolume_NUC | 65.45 |

#### 2.2.5.2. Functions

| Function Name | Expression |
|---|---|
| J_BindD | (((Kf_BindD * Dark) * Binder) - (Kr_BindD * DarkB)) |
| J_BindF | (((Kf_BindF * Fluor) * Binder) - (Kr_BindF * FluorB)) |
| J_BleachBound | ((Kf_BleachBound * FluorB) - (Kr_BleachBound * DarkB)) |
| J_BleachFree | ((Kf_BleachFree * Fluor) - (Kr_BleachFree * Dark)) |
| Kf_BleachBound | ((kbleach_BleachBound * Laser) * (t > start) * (t < (start + duration))) |
| Kf_BleachFree | ((kbleach_BleachFree * Laser) * (t > start) * (t < (start + duration))) |
| Size_CYT | (VolumePerUnitVolume_CYT * vcRegionVolume('IC')) |
| Size_EC | (VolumePerUnitVolume_EC * vcRegionVolume('EC')) |
| Size_NM | (AreaPerUnitVolume_NM * vcRegionVolume('IC')) |
| Size_NUC | (VolumePerUnitVolume_NUC * vcRegionVolume('IC')) |
| Size_PM | (AreaPerUnitArea_PM * vcRegionArea('EC_IC_membrane')) |
| sobj_IC1_EC0_size | vcRegionArea('EC_IC_membrane') |
| vobj_EC0_size | vcRegionVolume('EC') |
| vobj_IC1_size | vcRegionVolume('IC') |

#### 2.2.5.3. Volume Domains

### 2.2.5.3.1. IC

| OdeEquation Fluor | |
|---|---|
| Rate | ( - J_BindF - J_BleachFree) |
| Initial | Fluor_init_uM |

| OdeEquation Dark | |
|---|---|
| Rate | ( - J_BindD + J_BleachFree) |

| OdeEquation Dark | |
|---|---|
| Initial | Dark_init_uM |

| OdeEquation Binder | |
|---|---|
| Rate | ( - J_BindD - J_BindF) |
| Initial | Binder_init_uM |

| OdeEquation FluorB | |
|---|---|
| Rate | (J_BindF - J_BleachBound) |
| Initial | FluorB_init_uM |

| OdeEquation Laser | |
|---|---|
| Rate | 0.0 |
| Initial | Laser_init_uM |

| OdeEquation DarkB | |
|---|---|
| Rate | (J_BindD + J_BleachBound) |
| Initial | DarkB_init_uM |

#### 2.2.5.3.2. EC

##### 2.2.5.4. Membrane Domains

###### 2.2.5.4.1. EC_IC_membrane

##### 2.2.6. Simulation(s)

#### 2.3. Application: TIRF_off_center_bleach

| **Application Name: TIRF_off_center_bleach** |
|---|
| **Application Description: (copied from TIRF_center_bleach) (copied from Spherical_Cell_Gaussian_Bleach)** |

#### 2.3.1. Structure Mapping For TIRF_off_center_bleach

| Structure Mapping | | | | |
|---|---|---|---|---|
| **Structure** | **Subdomain** | **Resolved (T/F)** | **Surf/Vol** | **VolFract** |
| EC | EC | F | | |
| NUC | IC | F | | |
| CYT | IC | F | | |

#### 2.3.2. Reaction Mapping For TIRF_off_center_bleach

| Reaction Mapping | | | |
|---|---|---|---|
| **Name** | **Type** | **Enabled (T/F)** | **Fast (T/F)** |
| BindD | Reaction | T | F |
| BindF | Reaction | T | F |
| BleachFree | Reaction | T | F |
| BleachBound | Reaction | T | F |

| Initial Conditions | | | | |
|---|---|---|---|---|
| **Species** | **Structure** | **Initial Conc.** | **Diffusion Const.** | **Fixed (T/F)** |
| s0 | CYT | 0.0 M | 0.0 $m^2.s^1$ | F |
| s1 | CYT | 0.0 M | 0.0 $m^2.s^1$ | F |
| s2 | CYT | 0.0 M | 0.0 $m^2.s^1$ | F |
| s3 | CYT | 0.0 M | 0.0 $m^2.s^1$ | F |
| s4 | CYT | 0.0 M | 0.0 $m^2.s^1$ | F |
| s5 | CYT | 0.0 M | 0.0 $m^2.s^1$ | F |

#### 2.3.3. Membrane Mapping For TIRF_off_center_bleach

| Electrical Mapping - Membrane Potential | | | |
|---|---|---|---|
| **Membrane** | **Calculate V (T/F)** | **V initial** | **Specific Capacitance** |
| PM | F | 0.0 mV | 1.0 pF.m² |
| NM | F | 0.0 mV | 1.0 pF.m² |

| |
|---|
| Temperature: 300.0 K |

#### 2.3.4. Geometry: Geometry7

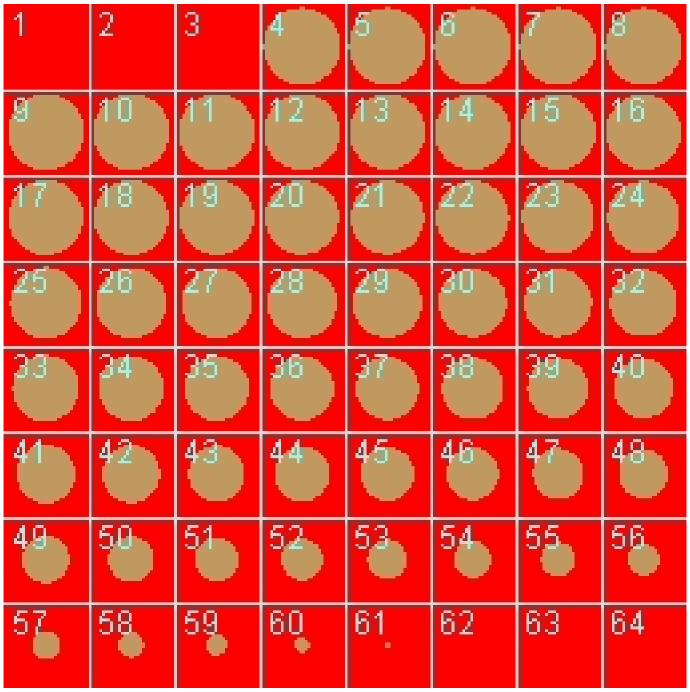

| | |
|---|---|
| Size | (22.0, 22.0, 11.0) |
| Origin | (-11.0, -11.0, -0.5) |

#### 2.3.5. Math Description: Copy of TIRF_center_bleach_generated

##### 2.3.5.1. Constants

| Constant Name | Expression |
| --- | --- |
| _F_ | 96485.3321 |
| _F_nmol_ | 9.64853321E-5 |
| _K_GHK_ | 1.0E-9 |
| _N_pmol_ | 6.02214179E11 |
| _PI_ | 3.141592653589793 |
| _R_ | 8314.46261815 |
| _T_ | 300.0 |
| AreaPerUnitArea_PM | 1.0 |
| AreaPerUnitVolume_NM | 1.2 |
| Binder_init_uM | 0.0 |
| BleachRadius | 2.0 |
| Dark_init_uM | 0.0 |
| DarkB_init_uM | 0.0 |
| duration | 1.0 |
| Fluor_init_uM | 0.0 |
| FluorB_init_uM | 0.0 |
| K_millivolts_per_volt | 1000.0 |
| kbleach_BleachBound | 1.0 |
| kbleach_BleachFree | 1.0 |
| Kf_BindD | 1.0 |
| Kf_BindF | 1.0 |
| KMOLE | 0.001660538783162726 |
| Kr_BindD | 1.0 |
| Kr_BindF | 1.0 |
| Kr_BleachBound | 0.0 |
| Kr_BleachFree | 0.0 |
| Laser_init_uM | 0.0 |

| Constant Name | Expression |
|---|---|
| sigmaaxial | 1.5 |
| sigmalateral | 0.5 |
| start | 1.0 |
| Voltage_NM | 0.0 |
| Voltage_PM | 0.0 |
| VolumePerUnitVolume_CYT | 1.0 |
| VolumePerUnitVolume_EC | 1.0 |
| VolumePerUnitVolume_NUC | 65.45 |

#### 2.3.5.2. Functions

| Function Name | Expression |
|---|---|
| J_BindD | (((Kf_BindD * Dark) * Binder) - (Kr_BindD * DarkB)) |
| J_BindF | (((Kf_BindF * Fluor) * Binder) - (Kr_BindF * FluorB)) |
| J_BleachBound | ((Kf_BleachBound * FluorB) - (Kr_BleachBound * DarkB)) |
| J_BleachFree | ((Kf_BleachFree * Fluor) - (Kr_BleachFree * Dark)) |
| Kf_BleachBound | ((kbleach_BleachBound * Laser) * (t > start) * (t < (start + duration))) |
| Kf_BleachFree | ((kbleach_BleachFree * Laser) * (t > start) * (t < (start + duration))) |
| Size_CYT | (VolumePerUnitVolume_CYT * vcRegionVolume('IC')) |
| Size_EC | (VolumePerUnitVolume_EC * vcRegionVolume('EC')) |
| Size_NM | (AreaPerUnitVolume_NM * vcRegionVolume('IC')) |
| Size_NUC | (VolumePerUnitVolume_NUC * vcRegionVolume('IC')) |
| Size_PM | (AreaPerUnitArea_PM * vcRegionArea('EC_IC_membrane')) |
| sobj_IC1_EC0_size | vcRegionArea('EC_IC_membrane') |
| vobj_EC0_size | vcRegionVolume('EC') |
| vobj_IC1_size | vcRegionVolume('IC') |

#### 2.3.5.3. Volume Domains

### 2.3.5.3.1. IC

| OdeEquation Fluor | |
|---|---|
| Rate | ( - J_BindF - J_BleachFree) |
| Initial | Fluor_init_uM |

| OdeEquation Dark | |
|---|---|
| Rate | ( - J_BindD + J_BleachFree) |
| Initial | Dark_init_uM |

| OdeEquation Binder | |
|---|---|
| Rate | ( - J_BindD - J_BindF) |
| Initial | Binder_init_uM |

| OdeEquation FluorB | |
|---|---|
| Rate | (J_BindF - J_BleachBound) |
| Initial | FluorB_init_uM |

| OdeEquation Laser | |
|---|---|
| Rate | 0.0 |
| Initial | Laser_init_uM |

| OdeEquation DarkB | |
|---|---|
| Rate | (J_BindD + J_BleachBound) |
| Initial | DarkB_init_uM |

### 2.3.5.3.2. EC

###### 2.3.5.4. Membrane Domains

###### 2.3.5.4.1. EC_IC_membrane

##### 2.3.6. Simulation(s)

#### 2.4. Application: Image-based center bleach

| Application Name: Image-based center bleach |
|---|
| Application Description: (copied from Spherical_Cell_Gaussian_Bleach) |

##### 2.4.1. Structure Mapping For Image-based center bleach

| Structure Mapping | | | | |
|---|---|---|---|---|
| **Structure** | **Subdomain** | **Resolved (T/F)** | **Surf/Vol** | **VolFract** |
| EC | ec | F | | |
| NUC | Nucleus | F | | |
| CYT | cytosol | F | | |

##### 2.4.2. Reaction Mapping For Image-based center bleach

| Reaction Mapping | | | |
|---|---|---|---|
| **Name** | **Type** | **Enabled (T/F)** | **Fast (T/F)** |
| BindD | Reaction | T | F |
| BindF | Reaction | T | F |
| BleachFree | Reaction | T | F |
| BleachBound | Reaction | T | F |

| Initial Conditions | | | | |
|---|---|---|---|---|
| **Species** | **Structure** | **Initial Conc.** | **Diffusion Const.** | **Fixed** |

| Initial Conditions | | | | |
|---|---|---|---|---|
| **Species** | **Structure** | **Initial Conc.** | **Diffusion Const.** | **Fixed (T/F)** |
| s0 | CYT | 10.0 M | 10.0 $m^2.s^1$ | F |
| s1 | CYT | 0.0 M | 10.0 $m^2.s^1$ | F |
| s2 | CYT | 0.0 M | 0.0 $m^2.s^1$ | F |
| s3 | CYT | 0.0 M | 0.0 $m^2.s^1$ | F |
| s4 | CYT | exp( - ((((x - 23.0) ^ 2.0) / (2.0 * (sigmalateral ^ 2.0))) + (((y - 39.0) ^ 2.0) / (2.0 * (sigmalateral ^ 2.0))) + (((z - 13.0) ^ 2.0) / (2.0 * (sigmaaxial ^ 2.0))))) M | 0.0 $m^2.s^1$ | F |
| s5 | CYT | 0.0 M | 0.0 $m^2.s^1$ | F |

#### 2.4.3. Membrane Mapping For Image-based center bleach

| Electrical Mapping - Membrane Potential | | | |
|---|---|---|---|
| **Membrane** | **Calculate V (T/F)** | **V initial** | **Specific Capacitance** |
| PM | F | 0.0 mV | 1.0 $pF.m^2$ |
| NM | F | 0.0 mV | 1.0 $pF.m^2$ |

| |
|---|
| Temperature: 300.0 K |

#### 2.4.4. Geometry: Site visit _Application0_20111127_695607844

| | |
|---|---|
| Size | (74.24, 74.24, 26.0) |
| Origin | (0.0, 0.0, 0.0) |

#### 2.4.5. Math Description: Copy of Spherical_Cell_Gaussian_Bleach_generated

##### 2.4.5.1. Constants

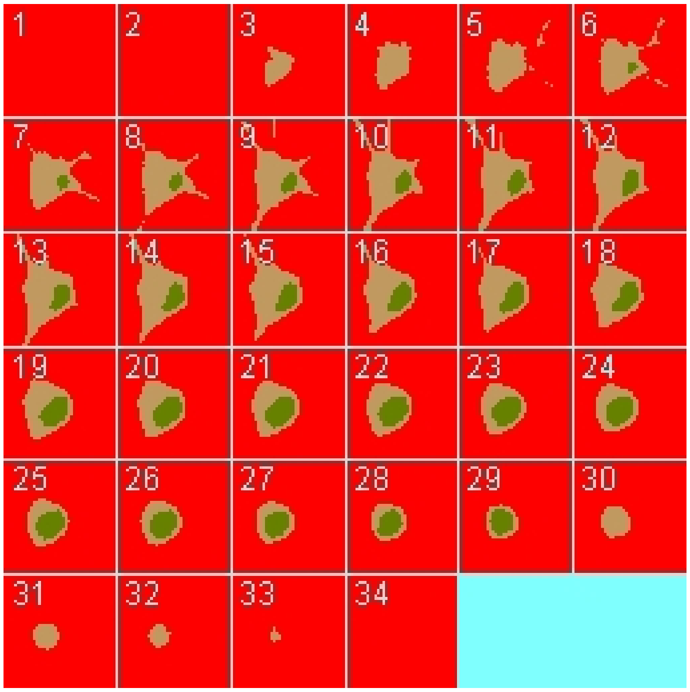

| Constant Name | Expression |
|---|---|
| _F_ | 96485.3321 |
| _F_nmol_ | 9.64853321E-5 |
| _K_GHK_ | 1.0E-9 |
| _N_pmol_ | 6.02214179E11 |
| _PI_ | 3.141592653589793 |
| _R_ | 8314.46261815 |
| _T_ | 300.0 |
| AreaPerUnitArea_NM | 1.0 |
| AreaPerUnitArea_PM | 1.0 |
| Binder_init_uM | 0.0 |
| BleachRadius | 2.0 |
| Dark_diffusionRate | 10.0 |
| Dark_init_uM | 0.0 |
| DarkB_init_uM | 0.0 |

| Constant Name | Expression |
| --- | --- |
| duration | 1.0 |
| Fluor_diffusionRate | 10.0 |
| Fluor_init_uM | 10.0 |
| FluorB_init_uM | 0.0 |
| K_millivolts_per_volt | 1000.0 |
| kbleach_BleachBound | 1.0 |
| kbleach_BleachFree | 1.0 |
| Kf_BindD | 1.0 |
| Kf_BindF | 1.0 |
| KMOLE | 0.001660538783162726 |
| Kr_BindD | 1.0 |
| Kr_BindF | 1.0 |
| Kr_BleachBound | 0.0 |
| Kr_BleachFree | 0.0 |
| sigmaaxial | 1.5 |
| sigmalateral | 0.5 |
| start | 1.0 |
| Voltage_NM | 0.0 |
| Voltage_PM | 0.0 |
| VolumePerUnitVolume_CYT | 1.0 |
| VolumePerUnitVolume_EC | 1.0 |
| VolumePerUnitVolume_NUC | 1.0 |

#### 2.4.5.2. Functions

| Function Name | Expression |
| --- | --- |
| J_BindD | (((Kf_BindD * Dark) * Binder) - (Kr_BindD * DarkB)) |

| Function Name | Expression |
|---|---|
| J_BindF | (((Kf_BindF * Fluor) * Binder) - (Kr_BindF * FluorB)) |
| J_BleachBound | ((Kf_BleachBound * FluorB) - (Kr_BleachBound * DarkB)) |
| J_BleachFree | ((Kf_BleachFree * Fluor) - (Kr_BleachFree * Dark)) |
| Kf_BleachBound | ((kbleach_BleachBound * Laser) * (t > start) * (t < (start + duration))) |
| Kf_BleachFree | ((kbleach_BleachFree * Laser) * (t > start) * (t < (start + duration))) |
| Laser_init_uM | exp( - ((((x - 23.0) ^ 2.0) / (2.0 * (sigmalateral ^ 2.0))) + (((y - 39.0) ^ 2.0) / (2.0 * (sigmalateral ^ 2.0))) + (((z - 13.0) ^ 2.0) / (2.0 * (sigmaaxial ^ 2.0))))) |
| Size_CYT | (VolumePerUnitVolume_CYT * vcRegionVolume('cytosol')) |
| Size_EC | (VolumePerUnitVolume_EC * vcRegionVolume('ec')) |
| Size_NM | (AreaPerUnitArea_NM * vcRegionArea('Nucleus_cytosol_membrane')) |
| Size_NUC | (VolumePerUnitVolume_NUC * vcRegionVolume('Nucleus')) |
| Size_PM | (AreaPerUnitArea_PM * vcRegionArea('cytosol_ec_membrane')) |
| sobj_cytosol1_ec0_size | vcRegionArea('cytosol_ec_membrane') |
| sobj_Nucleus2_cytosol1_size | vcRegionArea('Nucleus_cytosol_membrane') |
| vobj_cytosol1_size | vcRegionVolume('cytosol') |
| vobj_ec0_size | vcRegionVolume('ec') |
| vobj_Nucleus2_size | vcRegionVolume('Nucleus') |

#### 2.4.5.3. Volume Domains

### 2.4.5.3.1. ec

##### 2.4.5.3.2. cytosol

| PdeEquation Fluor | |
|---|---|
| Rate | ( - J_BindF - J_BleachFree) |
| Diffusion | Fluor_diffusionRate |
| Initial | Fluor_init_uM |

| PdeEquation Dark | |
|---|---|
| Rate | ( - J_BindD + J_BleachFree) |
| Diffusion | Dark_diffusionRate |
| Initial | Dark_init_uM |

| OdeEquation Binder | |
|---|---|
| Rate | ( - J_BindD - J_BindF) |
| Initial | Binder_init_uM |

| OdeEquation FluorB | |
|---|---|
| Rate | (J_BindF - J_BleachBound) |
| Initial | FluorB_init_uM |

| OdeEquation Laser | |
|---|---|
| Rate | 0.0 |
| Initial | Laser_init_uM |

| OdeEquation DarkB | |
|---|---|
| Rate | (J_BindD + J_BleachBound) |
| Initial | DarkB_init_uM |

###### 2.4.5.3.3. Nucleus

##### 2.4.5.4. Membrane Domains

###### 2.4.5.4.1. cytosol_ec_membrane

| JumpCondition Fluor | |
|---|---|
| InFlux | 0.0 |
| OutFlux | 0.0 |

| JumpCondition Dark | |
|---|---|
| InFlux | 0.0 |
| OutFlux | 0.0 |

##### 2.4.5.4.2. Nucleus_cytosol_membrane

| JumpCondition Fluor | |
|---|---|
| InFlux | 0.0 |
| OutFlux | 0.0 |

| JumpCondition Dark | |
|---|---|
| InFlux | 0.0 |
| OutFlux | 0.0 |

#### 2.4.6. Simulation(s)

##### 2.4.6.1. 2Photon Bleach

| **Simulation Name: 2Photon Bleach** |
|---|

| Overriden Parameters | | |
|---|---|---|
| **Name** | **Actual Value** | **Default Value** |
| Fluor_diffusionRate | "1.0", "10.0" | 10.0 |
| kbleach_BleachFree | 10.0 | 1.0 |
| duration | "0.3", "3.0" | 1.0 |
| kbleach_BleachBound | 10.0 | 1.0 |
| Dark_diffusionRate | Fluor_diffusionRate | 10.0 |

| Geometry Setting |
|---|

| | |
|---|---|
| Geometry Size (um) | (74.24, 74.24, 26.0) |
| Mesh Size (elements) | (151, 151, 54) |

| Advanced Settings | |
|---|---|
| Solver Name | Fully-Implicit Finite Volume, Regular Grid (Variable Time Step) |
| Time Bounds - Starting | 0.0 |
| Time Bounds - Ending | 20.0 |
| Time Step - Min | 0.0 |
| Time Step - Default | 0.05 |
| Time Step - Max | 0.1 |
| Error Tolerance - Absolute | 1.0E-9 |
| Error Tolerance - Relative | 1.0E-7 |
| Output Time Step | 0.5 |
| Use Symbolic Jacobian (T/F) | F |

#### 2.4.6.2. Widefield Bleach

| **Simulation Name: Widefield Bleach** |
|---|

| **Overriden Parameters** | | |
|---|---|---|
| **Name** | **Actual Value** | **Default Value** |
| Fluor_diffusionRate | "1.0", "10.0" | 10.0 |
| kbleach_BleachFree | 10.0 | 1.0 |
| sigmaaxial | 30.0 | 1.5 |
| duration | "0.3", "3.0" | 1.0 |
| kbleach_BleachBound | 10.0 | 1.0 |

| | | |
|---|---|---|
| Dark_diffusionRate | Fluor_diffusionRate | 10.0 |

| Geometry Setting | |
|---|---|
| Geometry Size (um) | (74.24, 74.24, 26.0) |
| Mesh Size (elements) | (151, 151, 54) |

| Advanced Settings | |
|---|---|
| Solver Name | Fully-Implicit Finite Volume, Regular Grid (Variable Time Step) |
| Time Bounds - Starting | 0.0 |
| Time Bounds - Ending | 20.0 |
| Time Step - Min | 0.0 |
| Time Step - Default | 0.05 |
| Time Step - Max | 0.1 |
| Error Tolerance - Absolute | 1.0E-9 |
| Error Tolerance - Relative | 1.0E-7 |
| Output Time Step | 0.5 |
| Use Symbolic Jacobian (T/F) | F |

#### 2.4.6.3. 2Photon Bleach w binding

| Simulation Name: 2Photon Bleach w binding |
|---|

| Overriden Parameters | | |
|---|---|---|
| **Name** | **Actual Value** | **Default Value** |
| Fluor_diffusionRate | "1.0", "10.0" | 10.0 |
| Kf_BindF | 100.0 | 1.0 |
| Binder_init_uM | 5.0 | 0.0 |

| | | |
|---|---|---|
| Kr_BindD | 100.0 | 1.0 |
| kbleach_BleachFree | 10.0 | 1.0 |
| kbleach_BleachBound | 10.0 | 1.0 |
| Dark_diffusionRate | Fluor_diffusionRate | 10.0 |

| Geometry Setting | |
|---|---|
| Geometry Size (um) | (74.24, 74.24, 26.0) |
| Mesh Size (elements) | (151, 151, 54) |

| Advanced Settings | |
|---|---|
| Solver Name | Fully-Implicit Finite Volume, Regular Grid (Variable Time Step) |
| Time Bounds - Starting | 0.0 |
| Time Bounds - Ending | 20.0 |
| Time Step - Min | 0.0 |
| Time Step - Default | 0.05 |
| Time Step - Max | 0.1 |
| Error Tolerance - Absolute | 1.0E-9 |
| Error Tolerance - Relative | 1.0E-7 |
| Output Time Step | 0.5 |
| Use Symbolic Jacobian (T/F) | F |

##### 2.4.6.4. Widefield Bleach w binding

**Simulation Name: Widefield Bleach w binding**

| Overriden Parameters | | |
|---|---|---|
| **Name** | **Actual Value** | **Default Value** |

| | | |
|---|---|---|
| kbleach_BleachFree | 10.0 | 1.0 |
| kbleach_BleachBound | 10.0 | 1.0 |
| duration | "0.3", "3.0" | 1.0 |
| Kr_BindD | 100.0 | 1.0 |
| Fluor_diffusionRate | "1.0", "10.0" | 10.0 |
| Dark_diffusionRate | Fluor_diffusionRate | 10.0 |
| sigmaaxial | 30.0 | 1.5 |
| Binder_init_uM | 5.0 | 0.0 |
| Kf_BindF | 100.0 | 1.0 |

| **Geometry Setting** | |
|---|---|
| Geometry Size (um) | (74.24, 74.24, 26.0) |
| Mesh Size (elements) | (151, 151, 54) |

| **Advanced Settings** | |
|---|---|
| Solver Name | Fully-Implicit Finite Volume, Regular Grid (Variable Time Step) |
| Time Bounds - Starting | 0.0 |
| Time Bounds - Ending | 20.0 |
| Time Step - Min | 0.0 |
| Time Step - Default | 0.05 |
| Time Step - Max | 0.1 |
| Error Tolerance - Absolute | 1.0E-9 |
| Error Tolerance - Relative | 1.0E-7 |
| Output Time Step | 0.5 |
| Use Symbolic Jacobian (T/F) | F |

#### 2.5. Application: Image-based process bleach

| **Application Name: Image-based process bleach** |
| --- |
| **Application Description: (copied from Image-based center bleach) (copied from Spherical_Cell_Gaussian_Bleach)** |

##### 2.5.1. Structure Mapping For Image-based process bleach

| **Structure Mapping** | | | | |
| --- | --- | --- | --- | --- |
| **Structure** | **Subdomain** | **Resolved (T/F)** | **Surf/Vol** | **VolFract** |
| EC | ec | F | | |
| NUC | Nucleus | F | | |
| CYT | cytosol | F | | |

##### 2.5.2. Reaction Mapping For Image-based process bleach

| **Reaction Mapping** | | | |
| --- | --- | --- | --- |
| **Name** | **Type** | **Enabled (T/F)** | **Fast (T/F)** |
| BindD | Reaction | T | F |
| BindF | Reaction | T | F |
| BleachFree | Reaction | T | F |
| BleachBound | Reaction | T | F |

| **Initial Conditions** | | | | |
| --- | --- | --- | --- | --- |
| **Species** | **Structure** | **Initial Conc.** | **Diffusion Const.** | **Fixed (T/F)** |
| s0 | CYT | 10.0 M | 10.0 $m^2.s^1$ | F |
| s1 | CYT | 0.0 M | 10.0 $m^2.s^1$ | F |
| s2 | CYT | 0.0 M | 0.0 $m^2.s^1$ | F |

| Initial Conditions | | | | |
|---|---|---|---|---|
| **Species** | **Structure** | **Initial Conc.** | **Diffusion Const.** | **Fixed (T/F)** |
| s3 | CYT | 0.0 M | 0.0 $m^2.s^1$ | F |
| s4 | CYT | exp( - ((((x - 60.0) ^ 2.0) / (2.0 * (sigmalateral ^ 2.0))) + (((y - 53.0) ^ 2.0) / (2.0 * (sigmalateral ^ 2.0))) + (((z - 4.0) ^ 2.0) / (2.0 * (sigmaaxial ^ 2.0))))) M | 0.0 $m^2.s^1$ | F |
| s5 | CYT | 0.0 M | 0.0 $m^2.s^1$ | F |

#### 2.5.3. Membrane Mapping For Image-based process bleach

| Electrical Mapping - Membrane Potential | | | |
|---|---|---|---|
| **Membrane** | **Calculate V (T/F)** | **V initial** | **Specific Capacitance** |
| PM | F | 0.0 mV | 1.0 $pF.m^2$ |
| NM | F | 0.0 mV | 1.0 $pF.m^2$ |

| |
|---|
| Temperature: 300.0 K |

#### 2.5.4. Geometry: Site visit _Application0_20111127_695607844

| | |
|---|---|
| Size | (74.24, 74.24, 26.0) |
| Origin | (0.0, 0.0, 0.0) |

#### 2.5.5. Math Description: Copy of Image-based center bleach_generated

##### 2.5.5.1. Constants

| Constant Name | Expression |
|---|---|
| _F_ | 96485.3321 |
| _F_nmol_ | 9.64853321E-5 |
| _K_GHK_ | 1.0E-9 |

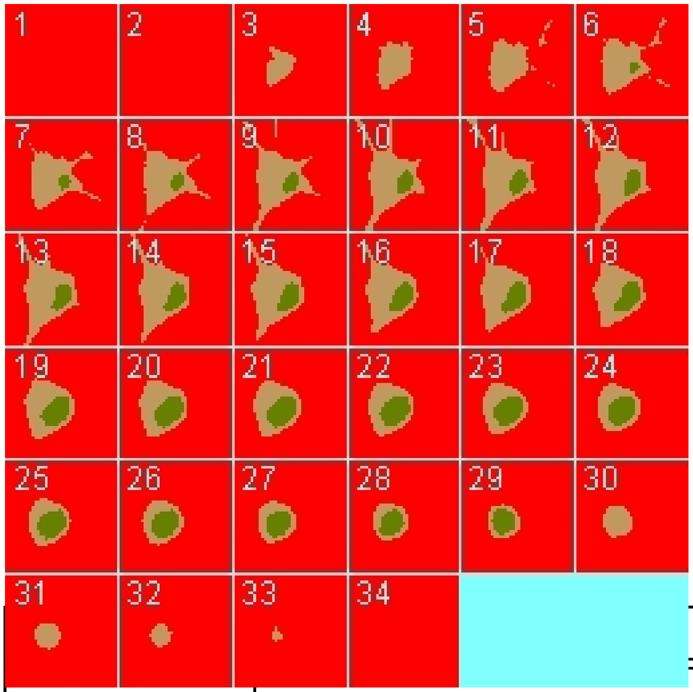

| Constant Name | Expression |
| --- | --- |
| _N_pmol_ | 6.02214179E11 |
| _PI_ | 3.141592653589793 |
| _R_ | 8314.46261815 |
| _T_ | 300.0 |
| AreaPerUnitArea_NM | 1.0 |
| AreaPerUnitArea_PM | 1.0 |
| Binder_init_uM | 0.0 |
| BleachRadius | 2.0 |
| Dark_diffusionRate | 10.0 |
| Dark_init_uM | 0.0 |
| DarkB_init_uM | 0.0 |
| duration | 1.0 |
| Fluor_diffusionRate | 10.0 |
| Fluor_init_uM | 10.0 |
| FluorB_init_uM | 0.0 |
| K_millivolts_per_volt | 1000.0 |

| Constant Name | Expression |
| --- | --- |
| kbleach_BleachBound | 1.0 |
| kbleach_BleachFree | 1.0 |
| Kf_BindD | 1.0 |
| Kf_BindF | 1.0 |
| KMOLE | 0.001660538783162726 |
| Kr_BindD | 1.0 |
| Kr_BindF | 1.0 |
| Kr_BleachBound | 0.0 |
| Kr_BleachFree | 0.0 |
| sigmaaxial | 1.5 |
| sigmalateral | 0.5 |
| start | 1.0 |
| Voltage_NM | 0.0 |
| Voltage_PM | 0.0 |
| VolumePerUnitVolume_CYT | 1.0 |
| VolumePerUnitVolume_EC | 1.0 |
| VolumePerUnitVolume_NUC | 1.0 |

#### 2.5.5.2. Functions

| Function Name | Expression |
| --- | --- |
| J_BindD | (((Kf_BindD * Dark) * Binder) - (Kr_BindD * DarkB)) |
| J_BindF | (((Kf_BindF * Fluor) * Binder) - (Kr_BindF * FluorB)) |
| J_BleachBound | ((Kf_BleachBound * FluorB) - (Kr_BleachBound * DarkB)) |
| J_BleachFree | ((Kf_BleachFree * Fluor) - (Kr_BleachFree * Dark)) |
| Kf_BleachBound | ((kbleach_BleachBound * Laser) * (t > start) * (t < (start + duration))) |
| Kf_BleachFree | ((kbleach_BleachFree * Laser) * (t > start) * (t < (start + duration))) |

| Function Name | Expression |
|---|---|
| Laser_init_uM | exp( - ((((x - 60.0) ^ 2.0) / (2.0 * (sigmalateral ^ 2.0))) + (((y - 53.0) ^ 2.0) / (2.0 * (sigmalateral ^ 2.0))) + (((z - 4.0) ^ 2.0) / (2.0 * (sigmaaxial ^ 2.0))))) |
| Size_CYT | (VolumePerUnitVolume_CYT * vcRegionVolume('cytosol')) |
| Size_EC | (VolumePerUnitVolume_EC * vcRegionVolume('ec')) |
| Size_NM | (AreaPerUnitArea_NM * vcRegionArea('Nucleus_cytosol_membrane')) |
| Size_NUC | (VolumePerUnitVolume_NUC * vcRegionVolume('Nucleus')) |
| Size_PM | (AreaPerUnitArea_PM * vcRegionArea('cytosol_ec_membrane')) |
| sobj_cytosol1_ec0_size | vcRegionArea('cytosol_ec_membrane') |
| sobj_Nucleus2_cytosol1_size | vcRegionArea('Nucleus_cytosol_membrane') |
| vobj_cytosol1_size | vcRegionVolume('cytosol') |
| vobj_ec0_size | vcRegionVolume('ec') |
| vobj_Nucleus2_size | vcRegionVolume('Nucleus') |

#### 2.5.5.3. Volume Domains

### 2.5.5.3.1. ec

##### 2.5.5.3.2. cytosol

| PdeEquation Fluor | |
|---|---|
| Rate | ( - J_BindF - J_BleachFree) |
| Diffusion | Fluor_diffusionRate |
| Initial | Fluor_init_uM |

| PdeEquation Dark | |
|---|---|
| Rate | ( - J_BindD + J_BleachFree) |
| Diffusion | Dark_diffusionRate |
| Initial | Dark_init_uM |

| OdeEquation Binder | |
| --- | --- |
| Rate | ( - J_BindD - J_BindF) |
| Initial | Binder_init_uM |

| OdeEquation FluorB | |
| --- | --- |
| Rate | (J_BindF - J_BleachBound) |
| Initial | FluorB_init_uM |

| OdeEquation Laser | |
| --- | --- |
| Rate | 0.0 |
| Initial | Laser_init_uM |

| OdeEquation DarkB | |
| --- | --- |
| Rate | (J_BindD + J_BleachBound) |
| Initial | DarkB_init_uM |

#### 2.5.5.3.3. Nucleus

#### 2.5.5.4. Membrane Domains

#### 2.5.5.4.1. cytosol_ec_membrane

| JumpCondition Fluor | |
| --- | --- |
| InFlux | 0.0 |
| OutFlux | 0.0 |

| JumpCondition Dark | |
| --- | --- |
| InFlux | 0.0 |
| OutFlux | 0.0 |

#### 2.5.5.4.2. Nucleus_cytosol_membrane

| JumpCondition Fluor | |
|---|---|
| InFlux | 0.0 |
| OutFlux | 0.0 |

| JumpCondition Dark | |
|---|---|
| InFlux | 0.0 |
| OutFlux | 0.0 |

#### 2.5.6. Simulation(s)

##### 2.5.6.1. 2P Bleach Process

| Simulation Name: 2P Bleach Process |
|---|

| Overriden Parameters | | |
|---|---|---|
| **Name** | **Actual Value** | **Default Value** |
| Fluor_diffusionRate | "1.0", "10.0" | 10.0 |
| kbleach_BleachFree | 10.0 | 1.0 |
| duration | "0.3", "3.0" | 1.0 |
| kbleach_BleachBound | 10.0 | 1.0 |
| Dark_diffusionRate | Fluor_diffusionRate | 10.0 |

| Geometry Setting | |
|---|---|
| Geometry Size (um) | (74.24, 74.24, 26.0) |
| Mesh Size (elements) | (151, 151, 54) |

| Advanced Settings |
|---|

| | |
|---|---|
| Solver Name | Fully-Implicit Finite Volume, Regular Grid (Variable Time Step) |
| Time Bounds - Starting | 0.0 |
| Time Bounds - Ending | 20.0 |
| Time Step - Min | 0.0 |
| Time Step - Default | 0.05 |
| Time Step - Max | 0.1 |
| Error Tolerance - Absolute | 1.0E-9 |
| Error Tolerance - Relative | 1.0E-7 |
| Output Time Step | 0.5 |
| Use Symbolic Jacobian (T/F) | F |

##### 2.5.6.2. Widefield Bleach Process

| **Simulation Name: Widefield Bleach Process** |
|---|

| **Overriden Parameters** | | |
|---|---|---|
| **Name** | **Actual Value** | **Default Value** |
| Fluor_diffusionRate | "1.0", "10.0" | 10.0 |
| kbleach_BleachFree | 10.0 | 1.0 |
| sigmaaxial | 30.0 | 1.5 |
| duration | "0.3", "3.0" | 1.0 |
| kbleach_BleachBound | 10.0 | 1.0 |
| Dark_diffusionRate | Fluor_diffusionRate | 10.0 |

| **Geometry Setting** | |
|---|---|
| Geometry Size (um) | (74.24, 74.24, 26.0) |

| | |
|---|---|
| Mesh Size (elements) | (151, 151, 54) |

| Advanced Settings | |
|---|---|
| Solver Name | Fully-Implicit Finite Volume, Regular Grid (Variable Time Step) |
| Time Bounds - Starting | 0.0 |
| Time Bounds - Ending | 20.0 |
| Time Step - Min | 0.0 |
| Time Step - Default | 0.05 |
| Time Step - Max | 0.1 |
| Error Tolerance - Absolute | 1.0E-9 |
| Error Tolerance - Relative | 1.0E-7 |
| Output Time Step | 0.5 |
| Use Symbolic Jacobian (T/F) | F |

#### 2.6. Application: Image-based below nucleus bleach

| **Application Name: Image-based below nucleus bleach** |
|---|
| **Application Description: (copied from Image-based process bleach) (copied from Image-based center bleach) (copied from Spherical_Cell_Gaussian_Bleach)** |

##### 2.6.1. Structure Mapping For Image-based below nucleus bleach

| **Structure Mapping** | | | | |
|---|---|---|---|---|
| **Structure** | **Subdomain** | **Resolved (T/F)** | **Surf/Vol** | **VolFract** |
| EC | ec | F | | |
| NUC | Nucleus | F | | |
| CYT | cytosol | F | | |

#### 2.6.2. Reaction Mapping For Image-based below nucleus bleach

| Reaction Mapping | | | |
|---|---|---|---|
| **Name** | **Type** | **Enabled (T/F)** | **Fast (T/F)** |
| BindD | Reaction | T | F |
| BindF | Reaction | T | F |
| BleachFree | Reaction | T | F |
| BleachBound | Reaction | T | F |

| Initial Conditions | | | | |
|---|---|---|---|---|
| **Species** | **Structure** | **Initial Conc.** | **Diffusion Const.** | **Fixed (T/F)** |
| s0 | CYT | 10.0 M | 10.0 $m^2.s^1$ | F |
| s1 | CYT | 0.0 M | 10.0 $m^2.s^1$ | F |
| s2 | CYT | 0.0 M | 0.0 $m^2.s^1$ | F |
| s3 | CYT | 0.0 M | 0.0 $m^2.s^1$ | F |
| s4 | CYT | exp( - ((((x - 28.0) ^ 2.0) / (2.0 * (sigmalateral ^ 2.0))) + (((y - 41.0) ^ 2.0) / (2.0 * (sigmalateral ^ 2.0))) + (((z - 24.4) ^ 2.0) / (2.0 * (sigmaaxial ^ 2.0))))) M | 0.0 $m^2.s^1$ | F |
| s5 | CYT | 0.0 M | 0.0 $m^2.s^1$ | F |

#### 2.6.3. Membrane Mapping For Image-based below nucleus bleach

| Electrical Mapping - Membrane Potential | | | |
|---|---|---|---|
| **Membrane** | **Calculate V (T/F)** | **V initial** | **Specific Capacitance** |
| PM | F | 0.0 mV | 1.0 $pF.m^2$ |
| NM | F | 0.0 mV | 1.0 $pF.m^2$ |

| |
|---|
| Temperature: 300.0 K |

#### 2.6.4. Geometry: Site visit _Application0_20111127_695607844

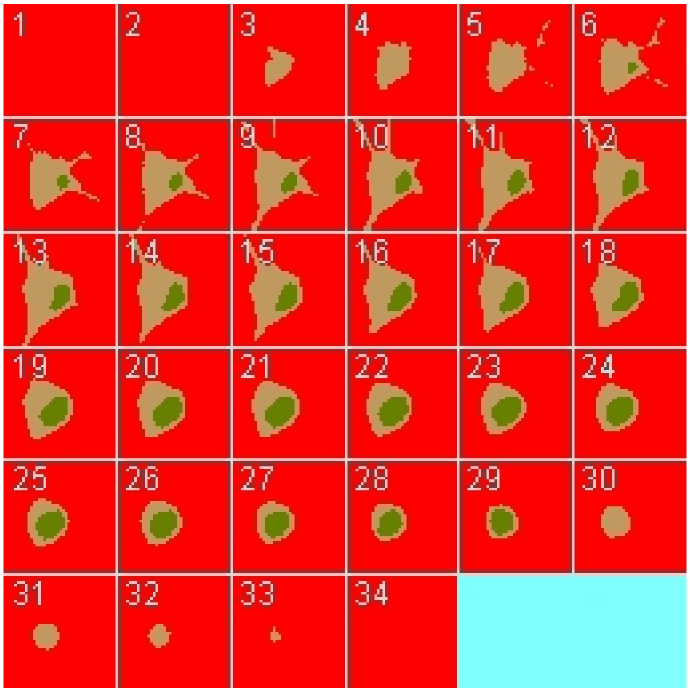

| | |
|---|---|
| Size | (74.24, 74.24, 26.0) |
| Origin | (0.0, 0.0, 0.0) |

#### 2.6.5. Math Description: Copy of Image-based process bleach_generated

##### 2.6.5.1. Constants

| Constant Name | Expression |
|---|---|
| _F_ | 96485.3321 |
| _F_nmol_ | 9.64853321E-5 |
| _K_GHK_ | 1.0E-9 |
| _N_pmol_ | 6.02214179E11 |
| _PI_ | 3.141592653589793 |

| Constant Name | Expression |
|---|---|
| _R_ | 8314.46261815 |
| _T_ | 300.0 |
| AreaPerUnitArea_NM | 1.0 |
| AreaPerUnitArea_PM | 1.0 |
| Binder_init_uM | 0.0 |
| BleachRadius | 2.0 |
| Dark_diffusionRate | 10.0 |
| Dark_init_uM | 0.0 |
| DarkB_init_uM | 0.0 |
| duration | 1.0 |
| Fluor_diffusionRate | 10.0 |
| Fluor_init_uM | 10.0 |
| FluorB_init_uM | 0.0 |
| K_millivolts_per_volt | 1000.0 |
| kbleach_BleachBound | 1.0 |
| kbleach_BleachFree | 1.0 |
| Kf_BindD | 1.0 |
| Kf_BindF | 1.0 |
| KMOLE | 0.001660538783162726 |
| Kr_BindD | 1.0 |
| Kr_BindF | 1.0 |
| Kr_BleachBound | 0.0 |
| Kr_BleachFree | 0.0 |
| sigmaaxial | 1.5 |
| sigmalateral | 0.5 |
| start | 1.0 |
| Voltage_NM | 0.0 |
| Voltage_PM | 0.0 |

| Constant Name | Expression |
| --- | --- |
| VolumePerUnitVolume_CYT | 1.0 |
| VolumePerUnitVolume_EC | 1.0 |
| VolumePerUnitVolume_NUC | 1.0 |

#### 2.6.5.2. Functions

| Function Name | Expression |
| --- | --- |
| J_BindD | (((Kf_BindD * Dark) * Binder) - (Kr_BindD * DarkB)) |
| J_BindF | (((Kf_BindF * Fluor) * Binder) - (Kr_BindF * FluorB)) |
| J_BleachBound | ((Kf_BleachBound * FluorB) - (Kr_BleachBound * DarkB)) |
| J_BleachFree | ((Kf_BleachFree * Fluor) - (Kr_BleachFree * Dark)) |
| Kf_BleachBound | ((kbleach_BleachBound * Laser) * (t > start) * (t < (start + duration))) |
| Kf_BleachFree | ((kbleach_BleachFree * Laser) * (t > start) * (t < (start + duration))) |
| Laser_init_uM | exp( - ((((x - 28.0) ^ 2.0) / (2.0 * (sigmalateral ^ 2.0))) + (((y - 41.0) ^ 2.0) / (2.0 * (sigmalateral ^ 2.0))) + (((z - 24.4) ^ 2.0) / (2.0 * (sigmaaxial ^ 2.0))))) |
| Size_CYT | (VolumePerUnitVolume_CYT * vcRegionVolume('cytosol')) |
| Size_EC | (VolumePerUnitVolume_EC * vcRegionVolume('ec')) |
| Size_NM | (AreaPerUnitArea_NM * vcRegionArea('Nucleus_cytosol_membrane')) |
| Size_NUC | (VolumePerUnitVolume_NUC * vcRegionVolume('Nucleus')) |
| Size_PM | (AreaPerUnitArea_PM * vcRegionArea('cytosol_ec_membrane')) |
| sobj_cytosol1_ec0_size | vcRegionArea('cytosol_ec_membrane') |
| sobj_Nucleus2_cytosol1_size | vcRegionArea('Nucleus_cytosol_membrane') |
| vobj_cytosol1_size | vcRegionVolume('cytosol') |
| vobj_ec0_size | vcRegionVolume('ec') |
| vobj_Nucleus2_size | vcRegionVolume('Nucleus') |

##### 2.6.5.3. Volume Domains

#### 2.6.5.3.1. ec

###### 2.6.5.3.2. cytosol

| PdeEquation Fluor | |
|---|---|
| Rate | ( - J_BindF - J_BleachFree) |
| Diffusion | Fluor_diffusionRate |
| Initial | Fluor_init_uM |

| PdeEquation Dark | |
|---|---|
| Rate | ( - J_BindD + J_BleachFree) |
| Diffusion | Dark_diffusionRate |
| Initial | Dark_init_uM |

| OdeEquation Binder | |
|---|---|
| Rate | ( - J_BindD - J_BindF) |
| Initial | Binder_init_uM |

| OdeEquation FluorB | |
|---|---|
| Rate | (J_BindF - J_BleachBound) |
| Initial | FluorB_init_uM |

| OdeEquation Laser | |
|---|---|
| Rate | 0.0 |
| Initial | Laser_init_uM |

| OdeEquation DarkB | |
|---|---|
| Rate | (J_BindD + J_BleachBound) |

| OdeEquation DarkB | |
|---|---|
| Initial | DarkB_init_uM |

#### 2.6.5.3.3. Nucleus

#### 2.6.5.4. Membrane Domains

#### 2.6.5.4.1. cytosol_ec_membrane

| JumpCondition Fluor | |
|---|---|
| InFlux | 0.0 |
| OutFlux | 0.0 |

| JumpCondition Dark | |
|---|---|
| InFlux | 0.0 |
| OutFlux | 0.0 |

#### 2.6.5.4.2. Nucleus_cytosol_membrane

| JumpCondition Fluor | |
|---|---|
| InFlux | 0.0 |
| OutFlux | 0.0 |

| JumpCondition Dark | |
|---|---|
| InFlux | 0.0 |
| OutFlux | 0.0 |

#### 2.6.6. Simulation(s)

#### 2.6.6.1. 2P Bleach Process_1

| Simulation Name: 2P Bleach Process_1 |
|---|

| Overriden Parameters | | |
|---|---|---|
| **Name** | **Actual Value** | **Default Value** |
| Fluor_diffusionRate | "1.0", "10.0" | 10.0 |
| kbleach_BleachFree | 10.0 | 1.0 |
| duration | "0.3", "3.0" | 1.0 |
| kbleach_BleachBound | 10.0 | 1.0 |
| Dark_diffusionRate | Fluor_diffusionRate | 10.0 |

| Geometry Setting | |
|---|---|
| Geometry Size (um) | (74.24, 74.24, 26.0) |
| Mesh Size (elements) | (151, 151, 54) |

| Advanced Settings | |
|---|---|
| Solver Name | Fully-Implicit Finite Volume, Regular Grid (Variable Time Step) |
| Time Bounds - Starting | 0.0 |
| Time Bounds - Ending | 20.0 |
| Time Step - Min | 0.0 |
| Time Step - Default | 0.05 |
| Time Step - Max | 0.1 |
| Error Tolerance - Absolute | 1.0E-9 |
| Error Tolerance - Relative | 1.0E-7 |
| Output Time Step | 0.5 |
| Use Symbolic Jacobian (T/F) | F |

#### 2.6.6.2. Widefield Bleach Process_1

| **Simulation Name: Widefield Bleach Process_1** |
|---|

| **Overriden Parameters** | | |
|---|---|---|
| **Name** | **Actual Value** | **Default Value** |
| Fluor_diffusionRate | "1.0", "10.0" | 10.0 |
| sigmaaxial | 30.0 | 1.5 |
| kbleach_BleachFree | 10.0 | 1.0 |
| duration | "0.3", "3.0" | 1.0 |
| kbleach_BleachBound | 10.0 | 1.0 |
| Dark_diffusionRate | Fluor_diffusionRate | 10.0 |

| **Geometry Setting** | |
|---|---|
| Geometry Size (um) | (74.24, 74.24, 26.0) |
| Mesh Size (elements) | (151, 151, 54) |

| **Advanced Settings** | |
|---|---|
| Solver Name | Fully-Implicit Finite Volume, Regular Grid (Variable Time Step) |
| Time Bounds - Starting | 0.0 |
| Time Bounds - Ending | 20.0 |
| Time Step - Min | 0.0 |
| Time Step - Default | 0.05 |
| Time Step - Max | 0.1 |
| Error Tolerance - Absolute | 1.0E-9 |
| Error Tolerance - Relative | 1.0E-7 |

| | |
|---|---|
| Output Time Step | 0.5 |
| Use Symbolic Jacobian (T/F) | F |

#### 2.7. Application: Image-based center circular bleach

| **Application Name: Image-based center circular bleach** |
|---|
| **Application Description: (copied from Image-based center bleach) (copied from Spherical_Cell_Gaussian_Bleach)** |

##### 2.7.1. Structure Mapping For Image-based center circular bleach

| **Structure Mapping** | | | | |
|---|---|---|---|---|
| **Structure** | **Subdomain** | **Resolved (T/F)** | **Surf/Vol** | **VolFract** |
| EC | ec | F | | |
| NUC | Nucleus | F | | |
| CYT | cytosol | F | | |

##### 2.7.2. Reaction Mapping For Image-based center circular bleach

| **Reaction Mapping** | | | |
|---|---|---|---|
| **Name** | **Type** | **Enabled (T/F)** | **Fast (T/F)** |
| BindD | Reaction | T | F |
| BindF | Reaction | T | F |
| BleachFree | Reaction | T | F |
| BleachBound | Reaction | T | F |

| **Initial Conditions** | | | | |
|---|---|---|---|---|
| **Species** | **Structure** | **Initial Conc.** | **Diffusion Const.** | **Fixed (T/F)** |
| s0 | CYT | 2.701562118716424 M | 10.0 $m^2.s^1$ | F |

| Initial Conditions | | | | |
|---|---|---|---|---|
| **Species** | **Structure** | **Initial Conc.** | **Diffusion Const.** | **Fixed (T/F)** |
| s1 | CYT | 0.0 M | 10.0 $m^2.s^1$ | F |
| s2 | CYT | 2.701562118716425 M | 0.0 $m^2.s^1$ | F |
| s3 | CYT | 7.2984378812835775 M | 0.0 $m^2.s^1$ | F |
| s4 | CYT | (exp( - (((z - 13.0) ^ 2.0) / (2.0 * (sigmaaxial ^ 2.0)))) * ((((x - 23.0) ^ 2.0) + ((y - 39.0) ^ 2.0)) < 4.0)) M | 0.0 $m^2.s^1$ | F |
| s5 | CYT | 0.0 M | 0.0 $m^2.s^1$ | F |

#### 2.7.3. Membrane Mapping For Image-based center circular bleach

| Electrical Mapping - Membrane Potential | | | |
|---|---|---|---|
| **Membrane** | **Calculate V (T/F)** | **V initial** | **Specific Capacitance** |
| PM | F | 0.0 mV | 1.0 $pF.m^2$ |
| NM | F | 0.0 mV | 1.0 $pF.m^2$ |

| |
|---|
| Temperature: 300.0 K |

#### 2.7.4. Geometry: Site visit _Application0_20111127_695607844

| | |
|---|---|
| Size | (74.24, 74.24, 26.0) |
| Origin | (0.0, 0.0, 0.0) |

#### 2.7.5. Math Description: Copy of Image-based center bleach_generated

##### 2.7.5.1. Constants

| Constant Name | Expression |
|---|---|
| _F_ | 96485.3321 |

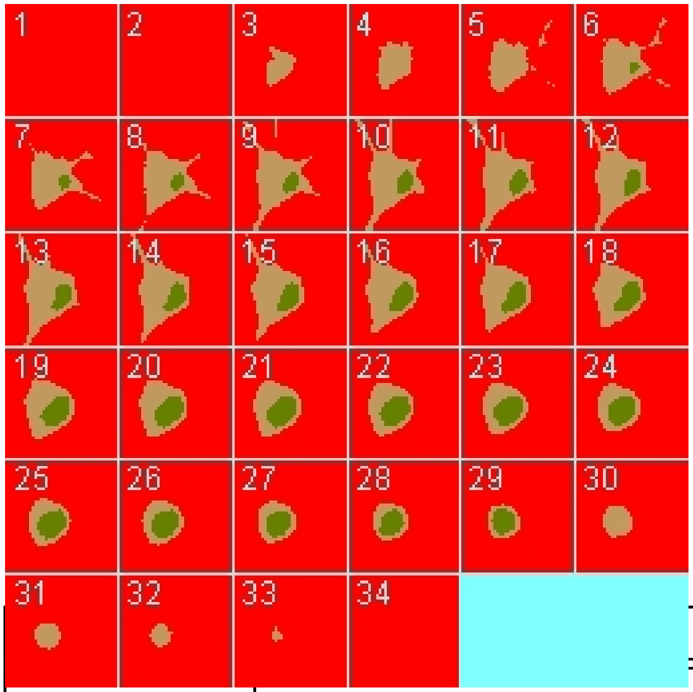

| Constant Name | Expression |
|---|---|
| _F_nmol_ | 9.64853321E-5 |
| _K_GHK_ | 1.0E-9 |
| _N_pmol_ | 6.02214179E11 |
| _PI_ | 3.141592653589793 |
| _R_ | 8314.46261815 |
| _T_ | 300.0 |
| AreaPerUnitArea_NM | 1.0 |
| AreaPerUnitArea_PM | 1.0 |
| Binder_init_uM | 2.701562118716425 |
| BleachRadius | 2.0 |
| Dark_diffusionRate | 10.0 |
| Dark_init_uM | 0.0 |
| DarkB_init_uM | 0.0 |
| duration | 1.0 |
| Fluor_diffusionRate | 10.0 |
| Fluor_init_uM | 2.701562118716424 |

| Constant Name | Expression |
| --- | --- |
| FluorB_init_uM | 7.2984378812835775 |
| K_millivolts_per_volt | 1000.0 |
| kbleach_BleachBound | 1.0 |
| kbleach_BleachFree | 1.0 |
| Kf_BindD | 1.0 |
| Kf_BindF | 1.0 |
| KMOLE | 0.001660538783162726 |
| Kr_BindD | 1.0 |
| Kr_BindF | 1.0 |
| Kr_BleachBound | 0.0 |
| Kr_BleachFree | 0.0 |
| sigmaaxial | 1.5 |
| sigmalateral | 0.5 |
| start | 1.0 |
| Voltage_NM | 0.0 |
| Voltage_PM | 0.0 |
| VolumePerUnitVolume_CYT | 1.0 |
| VolumePerUnitVolume_EC | 1.0 |
| VolumePerUnitVolume_NUC | 1.0 |

#### 2.7.5.2. Functions

| Function Name | Expression |
| --- | --- |
| J_BindD | (((Kf_BindD * Dark) * Binder) - (Kr_BindD * DarkB)) |
| J_BindF | (((Kf_BindF * Fluor) * Binder) - (Kr_BindF * FluorB)) |
| J_BleachBound | ((Kf_BleachBound * FluorB) - (Kr_BleachBound * DarkB)) |
| J_BleachFree | ((Kf_BleachFree * Fluor) - (Kr_BleachFree * Dark)) |

| Function Name | Expression |
|---|---|
| Kf_BleachBound | ((kbleach_BleachBound * Laser) * (t > start) * (t < (start + duration))) |
| Kf_BleachFree | ((kbleach_BleachFree * Laser) * (t > start) * (t < (start + duration))) |
| Laser_init_uM | (exp( - (((z - 13.0) ^ 2.0) / (2.0 * (sigmaaxial ^ 2.0)))) * ((((x - 23.0) ^ 2.0) + ((y - 39.0) ^ 2.0)) < 4.0)) |
| Size_CYT | (VolumePerUnitVolume_CYT * vcRegionVolume('cytosol')) |
| Size_EC | (VolumePerUnitVolume_EC * vcRegionVolume('ec')) |
| Size_NM | (AreaPerUnitArea_NM * vcRegionArea('Nucleus_cytosol_membrane')) |
| Size_NUC | (VolumePerUnitVolume_NUC * vcRegionVolume('Nucleus')) |
| Size_PM | (AreaPerUnitArea_PM * vcRegionArea('cytosol_ec_membrane')) |
| sobj_cytosol1_ec0_size | vcRegionArea('cytosol_ec_membrane') |
| sobj_Nucleus2_cytosol1_size | vcRegionArea('Nucleus_cytosol_membrane') |
| vobj_cytosol1_size | vcRegionVolume('cytosol') |
| vobj_ec0_size | vcRegionVolume('ec') |
| vobj_Nucleus2_size | vcRegionVolume('Nucleus') |

#### 2.7.5.3. Volume Domains

### 2.7.5.3.1. ec

##### 2.7.5.3.2. cytosol

| PdeEquation Fluor | |
|---|---|
| Rate | ( - J_BindF - J_BleachFree) |
| Diffusion | Fluor_diffusionRate |
| Initial | Fluor_init_uM |

| PdeEquation Dark | |
|---|---|
| Rate | ( - J_BindD + J_BleachFree) |

| PdeEquation Dark | |
|---|---|
| Diffusion | Dark_diffusionRate |
| Initial | Dark_init_uM |

| OdeEquation Binder | |
|---|---|
| Rate | ( - J_BindD - J_BindF) |
| Initial | Binder_init_uM |

| OdeEquation FluorB | |
|---|---|
| Rate | (J_BindF - J_BleachBound) |
| Initial | FluorB_init_uM |

| OdeEquation Laser | |
|---|---|
| Rate | 0.0 |
| Initial | Laser_init_uM |

| OdeEquation DarkB | |
|---|---|
| Rate | (J_BindD + J_BleachBound) |
| Initial | DarkB_init_uM |

###### 2.7.5.3.3. Nucleus

##### 2.7.5.4. Membrane Domains

###### 2.7.5.4.1. cytosol_ec_membrane

| JumpCondition Fluor | |
|---|---|
| InFlux | 0.0 |
| OutFlux | 0.0 |

| JumpCondition Dark |
|---|

| JumpCondition Dark | |
|---|---|
| InFlux | 0.0 |
| OutFlux | 0.0 |

##### 2.7.5.4.2. Nucleus_cytosol_membrane

| JumpCondition Fluor | |
|---|---|
| InFlux | 0.0 |
| OutFlux | 0.0 |

| JumpCondition Dark | |
|---|---|
| InFlux | 0.0 |
| OutFlux | 0.0 |

#### 2.7.6. Simulation(s)

##### 2.7.6.1. 2Photon Bleach no binding

| **Simulation Name: 2Photon Bleach no binding** |
|---|
| **Simulation Description: cloned from '2Photon Bleach_1' owned by user temp** |

| Overriden Parameters | | |
|---|---|---|
| **Name** | **Actual Value** | **Default Value** |
| Fluor_diffusionRate | "1.0", "10.0" | 10.0 |
| Binder_init_uM | 0.0 | 2.701562118716425 |
| kbleach_BleachFree | 10.0 | 1.0 |
| duration | "0.3", "3.0" | 1.0 |
| kbleach_BleachBound | 10.0 | 1.0 |
| Dark_diffusionRate | Fluor_diffusionRate | 10.0 |
| Fluor_init_uM | 10.0 | 2.701562118716424 |

| | | |
|---|---|---|
| FluorB_init_uM | 0.0 | 7.2984378812835775 |

| Geometry Setting | |
|---|---|
| Geometry Size (um) | (74.24, 74.24, 26.0) |
| Mesh Size (elements) | (256, 256, 90) |

| Advanced Settings | |
|---|---|
| Solver Name | Fully-Implicit Finite Volume, Regular Grid (Variable Time Step) |
| Time Bounds - Starting | 0.0 |
| Time Bounds - Ending | 20.0 |
| Time Step - Min | 0.0 |
| Time Step - Default | 0.05 |
| Time Step - Max | 0.1 |
| Error Tolerance - Absolute | 1.0E-9 |
| Error Tolerance - Relative | 1.0E-7 |
| Output Time Step | 0.5 |
| Use Symbolic Jacobian (T/F) | F |

#### 2.7.6.2. Widefield Bleach no binding

| Simulation Name: Widefield Bleach no binding |
|---|
| **Simulation Description: cloned from 'Widefield Bleach_1' owned by user temp** |

| Overriden Parameters | | |
|---|---|---|
| **Name** | **Actual Value** | **Default Value** |
| kbleach_BleachFree | 10.0 | 1.0 |
| kbleach_BleachBound | 10.0 | 1.0 |

| | | |
|---|---|---|
| duration | "0.3", "3.0" | 1.0 |
| Fluor_init_uM | 10.0 | 2.701562118716424 |
| Fluor_diffusionRate | "1.0", "10.0" | 10.0 |
| Dark_diffusionRate | Fluor_diffusionRate | 10.0 |
| sigmaaxial | 30.0 | 1.5 |
| Binder_init_uM | 0.0 | 2.701562118716425 |
| FluorB_init_uM | 0.0 | 7.2984378812835775 |

| **Geometry Setting** | |
|---|---|
| Geometry Size (um) | (74.24, 74.24, 26.0) |
| Mesh Size (elements) | (256, 256, 90) |

| **Advanced Settings** | |
|---|---|
| Solver Name | Fully-Implicit Finite Volume, Regular Grid (Variable Time Step) |
| Time Bounds - Starting | 0.0 |
| Time Bounds - Ending | 20.0 |
| Time Step - Min | 0.0 |
| Time Step - Default | 0.05 |
| Time Step - Max | 0.1 |
| Error Tolerance - Absolute | 1.0E-9 |
| Error Tolerance - Relative | 1.0E-7 |
| Output Time Step | 0.5 |
| Use Symbolic Jacobian (T/F) | F |

##### 2.7.6.3. 2Photon Bleach w fast binding

| |
|---|
| **Simulation Name: 2Photon Bleach w fast binding** |
| **Simulation Description: cloned from '2Photon Bleach w binding_1' owned by user temp** |

| **Overriden Parameters** | | |
|---|---|---|
| **Name** | **Actual Value** | **Default Value** |
| Fluor_diffusionRate | "1.0", "10.0" | 10.0 |
| Kf_BindF | 100.0 | 1.0 |
| Kr_BindF | 100.0 | 1.0 |
| Kf_BindD | 100.0 | 1.0 |
| Kr_BindD | 100.0 | 1.0 |
| kbleach_BleachFree | 10.0 | 1.0 |
| kbleach_BleachBound | 10.0 | 1.0 |
| Dark_diffusionRate | Fluor_diffusionRate | 10.0 |

| **Geometry Setting** | |
|---|---|
| Geometry Size (um) | (74.24, 74.24, 26.0) |
| Mesh Size (elements) | (256, 256, 90) |

| **Advanced Settings** | |
|---|---|
| Solver Name | Fully-Implicit Finite Volume, Regular Grid (Variable Time Step) |
| Time Bounds - Starting | 0.0 |
| Time Bounds - Ending | 20.0 |
| Time Step - Min | 0.0 |
| Time Step - Default | 0.05 |

| | |
|---|---|
| Time Step - Max | 0.1 |
| Error Tolerance - Absolute | 1.0E-9 |
| Error Tolerance - Relative | 1.0E-7 |
| Output Time Step | 0.5 |
| Use Symbolic Jacobian (T/F) | F |

##### 2.7.6.4. Widefield Bleach w fast binding

| |
|---|
| **Simulation Name: Widefield Bleach w fast binding** |
| **Simulation Description: cloned from 'Widefield Bleach w binding_1' owned by user temp** |

| **Overriden Parameters** | | |
|---|---|---|
| **Name** | **Actual Value** | **Default Value** |
| kbleach_BleachFree | 10.0 | 1.0 |
| Kf_BindD | 100.0 | 1.0 |
| kbleach_BleachBound | 10.0 | 1.0 |
| duration | "0.3", "3.0" | 1.0 |
| Kr_BindF | 100.0 | 1.0 |
| Kr_BindD | 100.0 | 1.0 |
| Fluor_diffusionRate | "1.0", "10.0" | 10.0 |
| Dark_diffusionRate | Fluor_diffusionRate | 10.0 |
| sigmaaxial | 30.0 | 1.5 |
| Kf_BindF | 100.0 | 1.0 |

| **Geometry Setting** | |
|---|---|
| Geometry Size (um) | (74.24, 74.24, 26.0) |
| Mesh Size (elements) | (256, 256, 90) |

| Advanced Settings | |
|---|---|
| Solver Name | Fully-Implicit Finite Volume, Regular Grid (Variable Time Step) |
| Time Bounds - Starting | 0.0 |
| Time Bounds - Ending | 20.0 |
| Time Step - Min | 0.0 |
| Time Step - Default | 0.05 |
| Time Step - Max | 0.1 |
| Error Tolerance - Absolute | 1.0E-9 |
| Error Tolerance - Relative | 1.0E-7 |
| Output Time Step | 0.5 |
| Use Symbolic Jacobian (T/F) | F |

##### 2.7.6.5. 2Photon Bleach w default binding

| **Simulation Name: 2Photon Bleach w default binding** |
|---|
| **Simulation Description: cloned from '2Photon Bleach w slower binding' owned by user temp cloned from '2Photon Bleach w binding_1' owned by user temp** |

| Overriden Parameters | | |
|---|---|---|
| **Name** | **Actual Value** | **Default Value** |
| Fluor_diffusionRate | "1.0", "10.0" | 10.0 |
| kbleach_BleachFree | 10.0 | 1.0 |
| kbleach_BleachBound | 10.0 | 1.0 |
| Dark_diffusionRate | Fluor_diffusionRate | 10.0 |

| Geometry Setting |
|---|

| | |
|---|---|
| Geometry Size (um) | (74.24, 74.24, 26.0) |
| Mesh Size (elements) | (256, 256, 90) |

| Advanced Settings | |
|---|---|
| Solver Name | Fully-Implicit Finite Volume, Regular Grid (Variable Time Step) |
| Time Bounds - Starting | 0.0 |
| Time Bounds - Ending | 20.0 |
| Time Step - Min | 0.0 |
| Time Step - Default | 0.05 |
| Time Step - Max | 0.1 |
| Error Tolerance - Absolute | 1.0E-9 |
| Error Tolerance - Relative | 1.0E-7 |
| Output Time Step | 0.5 |
| Use Symbolic Jacobian (T/F) | F |

#### 2.7.6.6. Widefield Bleach w defalt binding

| Simulation Name: Widefield Bleach w defalt binding |
|---|
| Simulation Description: cloned from 'Widefield Bleach w slower binding' owned by user temp<br>cloned from 'Widefield Bleach w binding_1' owned by user temp |

| Overriden Parameters | | |
|---|---|---|
| **Name** | **Actual Value** | **Default Value** |
| Fluor_diffusionRate | "1.0", "10.0" | 10.0 |
| sigmaaxial | 30.0 | 1.5 |
| kbleach_BleachFree | 10.0 | 1.0 |
| duration | "0.3", "3.0" | 1.0 |

| | | |
|---|---|---|
| kbleach_BleachBound | 10.0 | 1.0 |
| Dark_diffusionRate | Fluor_diffusionRate | 10.0 |

| Geometry Setting | |
|---|---|
| Geometry Size (um) | (74.24, 74.24, 26.0) |
| Mesh Size (elements) | (256, 256, 90) |

| Advanced Settings | |
|---|---|
| Solver Name | Fully-Implicit Finite Volume, Regular Grid (Variable Time Step) |
| Time Bounds - Starting | 0.0 |
| Time Bounds - Ending | 20.0 |
| Time Step - Min | 0.0 |
| Time Step - Default | 0.05 |
| Time Step - Max | 0.1 |
| Error Tolerance - Absolute | 1.0E-9 |
| Error Tolerance - Relative | 1.0E-7 |
| Output Time Step | 0.5 |
| Use Symbolic Jacobian (T/F) | F |

##### 2.7.6.7. 2Photon Bleach w slow binding

| **Simulation Name: 2Photon Bleach w slow binding** |
|---|
| **Simulation Description: cloned from 'Copy of 2Photon Bleach w slow binding' owned by user temp<br>cloned from '2Photon Bleach w slower binding' owned by user temp<br>cloned from '2Photon Bleach w binding_1' owned by user temp** |

| Overriden Parameters |
|---|

| Name | Actual Value | Default Value |
|---|---|---|
| Fluor_diffusionRate | "1.0", "10.0" | 10.0 |
| Kf_BindF | 0.1 | 1.0 |
| Kr_BindF | 0.1 | 1.0 |
| Kf_BindD | 0.1 | 1.0 |
| Kr_BindD | 0.1 | 1.0 |
| kbleach_BleachFree | 10.0 | 1.0 |
| kbleach_BleachBound | 10.0 | 1.0 |
| Dark_diffusionRate | Fluor_diffusionRate | 10.0 |

| Geometry Setting | |
|---|---|
| Geometry Size (um) | (74.24, 74.24, 26.0) |
| Mesh Size (elements) | (256, 256, 90) |

| Advanced Settings | |
|---|---|
| Solver Name | Fully-Implicit Finite Volume, Regular Grid (Variable Time Step) |
| Time Bounds - Starting | 0.0 |
| Time Bounds - Ending | 20.0 |
| Time Step - Min | 0.0 |
| Time Step - Default | 0.05 |
| Time Step - Max | 0.1 |
| Error Tolerance - Absolute | 1.0E-9 |
| Error Tolerance - Relative | 1.0E-7 |
| Output Time Step | 0.5 |
| Use Symbolic Jacobian (T/F) | F |

#### 2.7.6.8. Widefield Bleach w slow binding

| **Simulation Name: Widefield Bleach w slow binding** |
|---|
| **Simulation Description: cloned from 'Copy of Widefield Bleach w slow binding' owned by user temp<br>cloned from 'Widefield Bleach w slower binding' owned by user temp<br>cloned from 'Widefield Bleach w binding_1' owned by user temp** |

| **Overriden Parameters** | | |
|---|---|---|
| **Name** | **Actual Value** | **Default Value** |
| kbleach_BleachFree | 10.0 | 1.0 |
| Kf_BindD | 0.1 | 1.0 |
| kbleach_BleachBound | 10.0 | 1.0 |
| duration | "0.3", "3.0" | 1.0 |
| Kr_BindF | 0.1 | 1.0 |
| Kr_BindD | 0.1 | 1.0 |
| Fluor_diffusionRate | "1.0", "10.0" | 10.0 |
| Dark_diffusionRate | Fluor_diffusionRate | 10.0 |
| sigmaaxial | 30.0 | 1.5 |
| Kf_BindF | 0.1 | 1.0 |

| **Geometry Setting** | |
|---|---|
| Geometry Size (um) | (74.24, 74.24, 26.0) |
| Mesh Size (elements) | (256, 256, 90) |

| **Advanced Settings** | |
|---|---|
| Solver Name | Fully-Implicit Finite Volume, Regular Grid (Variable Time Step) |

| | |
|---|---|
| Time Bounds - Starting | 0.0 |
| Time Bounds - Ending | 20.0 |
| Time Step - Min | 0.0 |
| Time Step - Default | 0.05 |
| Time Step - Max | 0.1 |
| Error Tolerance - Absolute | 1.0E-9 |
| Error Tolerance - Relative | 1.0E-7 |
| Output Time Step | 0.5 |
| Use Symbolic Jacobian (T/F) | F |

#### 2.8. Application: Non spatial to determine SS

| **Application Name: Non spatial to determine SS** |
|---|
| **Application Description: (copied from Image-based center circular bleach) (copied from Image-based center bleach) (copied from Spherical_Cell_Gaussian_Bleach)** |

##### 2.8.1. Structure Mapping For Non spatial to determine SS

| **Structure Mapping** | | | | |
|---|---|---|---|---|
| **Structure** | **Subdomain** | **Resolved (T/F)** | **Surf/Vol** | **VolFract** |
| EC | Compartment | F | | |
| NUC | Compartment | F | | |
| CYT | Compartment | F | | |

##### 2.8.2. Reaction Mapping For Non spatial to determine SS

| **Reaction Mapping** | | | |
|---|---|---|---|
| **Name** | **Type** | **Enabled (T/F)** | **Fast (T/F)** |
| BindD | Reaction | T | F |
| BindF | Reaction | T | F |

| Reaction Mapping | | | |
|---|---|---|---|
| **Name** | **Type** | **Enabled (T/F)** | **Fast (T/F)** |
| BleachFree | Reaction | T | F |
| BleachBound | Reaction | T | F |

| Initial Conditions | | | | |
|---|---|---|---|---|
| **Species** | **Structure** | **Initial Conc.** | **Diffusion Const.** | **Fixed (T/F)** |
| s0 | CYT | 10.0 M | 0.0 $m^2.s^1$ | F |
| s1 | CYT | 0.0 M | 0.0 $m^2.s^1$ | F |
| s2 | CYT | 10.0 M | 0.0 $m^2.s^1$ | F |
| s3 | CYT | 0.0 M | 0.0 $m^2.s^1$ | F |
| s4 | CYT | 0.0 M | 0.0 $m^2.s^1$ | F |
| s5 | CYT | 0.0 M | 0.0 $m^2.s^1$ | F |

#### 2.8.3. Membrane Mapping For Non spatial to determine SS

| Electrical Mapping - Membrane Potential | | | |
|---|---|---|---|
| **Membrane** | **Calculate V (T/F)** | **V initial** | **Specific Capacitance** |
| PM | F | 0.0 mV | 1.0 $pF.m^2$ |
| NM | F | 0.0 mV | 1.0 $pF.m^2$ |

| |
|---|
| Temperature: 300.0 K |

#### 2.8.4. Geometry: nonspatial1898625483

Non spatial geometry.

#### 2.8.5. Math Description: Non spatial to determine SS_generated

#### 2.8.5.1. Constants

| Constant Name | Expression |
|---|---|
| _F_ | 96485.3321 |
| _F_nmol_ | 9.64853321E-5 |
| _K_GHK_ | 1.0E-9 |
| _N_pmol_ | 6.02214179E11 |
| _PI_ | 3.141592653589793 |
| _R_ | 8314.46261815 |
| _T_ | 300.0 |
| Binder_init_uM | 10.0 |
| BleachRadius | 2.0 |
| Dark_init_uM | 0.0 |
| DarkB_init_uM | 0.0 |
| duration | 1.0 |
| Fluor_init_uM | 10.0 |
| FluorB_init_uM | 0.0 |
| K_millivolts_per_volt | 1000.0 |
| kbleach_BleachBound | 1.0 |
| kbleach_BleachFree | 1.0 |
| Kf_BindD | 1.0 |
| Kf_BindF | 1.0 |
| KMOLE | 0.001660538783162726 |
| Kr_BindD | 1.0 |
| Kr_BindF | 1.0 |
| Kr_BleachBound | 0.0 |
| Kr_BleachFree | 0.0 |
| Laser_init_uM | 0.0 |
| sigmaaxial | 1.5 |

| Constant Name | Expression |
|---|---|
| sigmalateral | 0.5 |
| start | 1.0 |
| Voltage_NM | 0.0 |
| Voltage_PM | 0.0 |

#### 2.8.5.2. Functions

| Function Name | Expression |
|---|---|
| J_BindD | (((Kf_BindD * Dark) * Binder) - (Kr_BindD * DarkB)) |
| J_BindF | (((Kf_BindF * Fluor) * Binder) - (Kr_BindF * FluorB)) |
| J_BleachBound | ((Kf_BleachBound * FluorB) - (Kr_BleachBound * DarkB)) |
| J_BleachFree | ((Kf_BleachFree * Fluor) - (Kr_BleachFree * Dark)) |
| Kf_BleachBound | ((kbleach_BleachBound * Laser) * (t > start) * (t < (start + duration))) |
| Kf_BleachFree | ((kbleach_BleachFree * Laser) * (t > start) * (t < (start + duration))) |
| Size_CYT | (1.0 * 14891.899581611733) |
| Size_EC | (1.0 * 124712.10435961554) |
| Size_NM | (1.0 * 1406.7733692487282) |
| Size_NUC | (1.0 * 3697.013658772733) |
| Size_PM | (1.0 * 4738.640600365477) |

#### 2.8.5.3. Volume Domains

##### 2.8.5.3.1. Compartment

| OdeEquation Fluor | |
|---|---|
| Rate | ( - J_BindF - J_BleachFree) |
| Initial | Fluor_init_uM |

| OdeEquation Dark | |
|---|---|
| Rate | ( - J_BindD + J_BleachFree) |
| Initial | Dark_init_uM |

| OdeEquation Binder | |
|---|---|
| Rate | ( - J_BindD - J_BindF) |
| Initial | Binder_init_uM |

| OdeEquation FluorB | |
|---|---|
| Rate | (J_BindF - J_BleachBound) |
| Initial | FluorB_init_uM |

| OdeEquation Laser | |
|---|---|
| Rate | 0.0 |
| Initial | Laser_init_uM |

| OdeEquation DarkB | |
|---|---|
| Rate | (J_BindD + J_BleachBound) |
| Initial | DarkB_init_uM |

#### 2.8.6. Simulation(s)

##### 2.8.6.1. Simulation0

| Simulation Name: Simulation0 |
|---|
| Simulation Description: cloned from 'Simulation0' owned by user temp |

| Advanced Settings | |
|---|---|
| Solver Name | Combined Stiff Solver (IDA/CVODE) |
| Time Bounds - Starting | 0.0 |
| Time Bounds - Ending | 20.0 |
| Time Step - Min | 1.0E-8 |
| Time Step - Default | 0.1 |

| | |
|---|---|
| Time Step - Max | 1.0 |
| Error Tolerance - Absolute | 1.0E-9 |
| Error Tolerance - Relative | 1.0E-9 |
| Keep Every | 1 |
| Keep At Most | 1000 |
| Use Symbolic Jacobian (T/F) | F |

#### 2.9. Application: ProjectZ Image-based binding to liquid droplet

| |
|---|
| **Application Name: ProjectZ Image-based binding to liquid droplet** |
| **Application Description: (copied from Image-based center circular bleach) (copied from Image-based center bleach) (copied from Spherical_Cell_Gaussian_Bleach)** |

##### 2.9.1. Structure Mapping For ProjectZ Image-based binding to liquid droplet

| **Structure Mapping** | | | | |
|---|---|---|---|---|
| **Structure** | **Subdomain** | **Resolved (T/F)** | **Surf/Vol** | **VolFract** |
| EC | ec | F | | |
| NUC | Nucleus | F | | |
| CYT | cytosol | F | | |

##### 2.9.2. Reaction Mapping For ProjectZ Image-based binding to liquid droplet

| **Reaction Mapping** | | | |
|---|---|---|---|
| **Name** | **Type** | **Enabled (T/F)** | **Fast (T/F)** |
| BindD | Reaction | T | F |
| BindF | Reaction | T | F |
| BleachFree | Reaction | T | F |
| BleachBound | Reaction | T | F |

| Initial Conditions | | | | |
|---|---|---|---|---|
| **Species** | **Structure** | **Initial Conc.** | **Diffusion Const.** | **Fixed (T/F)** |
| s0 | CYT | 2.701562118716424 M | 10.0 $m^{2}.s^{1}$ | F |
| s1 | CYT | 0.0 M | 10.0 $m^{2}.s^{1}$ | F |
| s2 | CYT | (2.701562118716425 * ((((x - 23.0) ^ 2.0) + ((y - 39.0) ^ 2.0) + ((z - 13.0) ^ 2.0)) < 4.0)) M | (((((x - 23.0) ^ 2.0) + ((y - 39.0) ^ 2.0) + ((z - 13.0) ^ 2.0)) < 4.0) * BinderD) $m^{2}.s^{1}$ | F |
| s3 | CYT | (7.2984378812835775 * ((((x - 23.0) ^ 2.0) + ((y - 39.0) ^ 2.0) + ((z - 13.0) ^ 2.0)) < 4.0)) M | (((((x - 23.0) ^ 2.0) + ((y - 39.0) ^ 2.0) + ((z - 13.0) ^ 2.0)) < 4.0) * FBinderD) $m^{2}.s^{1}$ | F |
| s4 | CYT | ((exp( - (((z - 13.0) ^ 2.0) / (2.0 * (sigmaaxial ^ 2.0)))) * ((((x - 23.0) ^ 2.0) + ((y - 39.0) ^ 2.0)) < 4.0)) * (x > BleachX)) M | 0.0 $m^{2}.s^{1}$ | F |
| s5 | CYT | 0.0 M | (((((x - 23.0) ^ 2.0) + ((y - 39.0) ^ 2.0) + ((z - 13.0) ^ 2.0)) < 4.0) * FBinderD) $m^{2}.s^{1}$ | F |

#### 2.9.3. Membrane Mapping For ProjectZ Image-based binding to liquid droplet

| Electrical Mapping - Membrane Potential | | | |
|---|---|---|---|
| **Membrane** | **Calculate V (T/F)** | **V initial** | **Specific Capacitance** |
| PM | F | 0.0 mV | 1.0 $pF.m^{2}$ |
| NM | F | 0.0 mV | 1.0 $pF.m^{2}$ |

| |
|---|
| Temperature: 300.0 K |

#### 2.9.4. Geometry: Site visit _Application0_20111127_695607844

| | |
|---|---|
| Size | (74.24, 74.24, 26.0) |
| Origin | (0.0, 0.0, 0.0) |

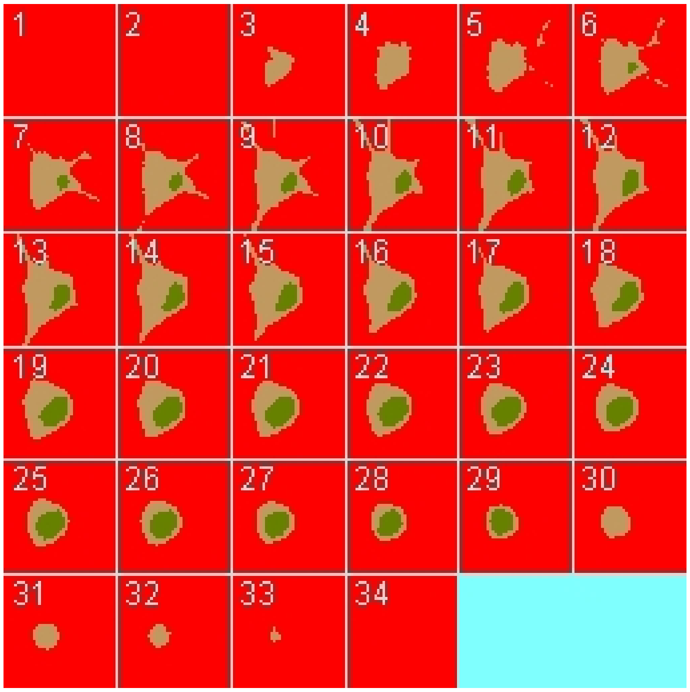

#### 2.9.5. Math Description: Copy of Image-based center circular bleach_generated

##### 2.9.5.1. Constants

| Constant Name | Expression |
|---|---|
| _F_ | 96485.3321 |
| _F_nmol_ | 9.64853321E-5 |
| _K_GHK_ | 1.0E-9 |
| _N_pmol_ | 6.02214179E11 |
| _PI_ | 3.141592653589793 |
| _R_ | 8314.46261815 |
| _T_ | 300.0 |
| AreaPerUnitArea_NM | 1.0 |
| AreaPerUnitArea_PM | 1.0 |
| BinderD | 0.1 |
| BleachRadius | 2.0 |

| Constant Name | Expression |
|---|---|
| BleachX | 0.0 |
| Dark_diffusionRate | 10.0 |
| Dark_init_uM | 0.0 |
| DarkB_init_uM | 0.0 |
| duration | 1.0 |
| FBinderD | 0.1 |
| Fluor_diffusionRate | 10.0 |
| Fluor_init_uM | 2.701562118716424 |
| K_millivolts_per_volt | 1000.0 |
| kbleach_BleachBound | 1.0 |
| kbleach_BleachFree | 1.0 |
| Kf_BindD | 1.0 |
| Kf_BindF | 1.0 |
| KMOLE | 0.001660538783162726 |
| Kr_BindD | 1.0 |
| Kr_BindF | 1.0 |
| Kr_BleachBound | 0.0 |
| Kr_BleachFree | 0.0 |
| sigmaaxial | 1.5 |
| sigmalateral | 0.5 |
| start | 1.0 |
| UnitFactor_molecules_um_neg_3_uM_neg_1 | (1.0 * pow(KMOLE,-1.0)) |
| Voltage_NM | 0.0 |
| Voltage_PM | 0.0 |
| VolumePerUnitVolume_CYT | 1.0 |
| VolumePerUnitVolum | 1.0 |

| Constant Name | Expression |
|---|---|
| e_EC | |
| VolumePerUnitVolume_NUC | 1.0 |

#### 2.9.5.2. Functions

| Function Name | Expression |
|---|---|
| Binder_diffusionRate | (((((x - 23.0) ^ 2.0) + ((y - 39.0) ^ 2.0) + ((z - 13.0) ^ 2.0)) < 4.0) * BinderD) |
| Binder_init_uM | (2.701562118716425 * ((((x - 23.0) ^ 2.0) + ((y - 39.0) ^ 2.0) + ((z - 13.0) ^ 2.0)) < 4.0)) |
| DarkB_diffusionRate | (((((x - 23.0) ^ 2.0) + ((y - 39.0) ^ 2.0) + ((z - 13.0) ^ 2.0)) < 4.0) * FBinderD) |
| FluorB_diffusionRate | (((((x - 23.0) ^ 2.0) + ((y - 39.0) ^ 2.0) + ((z - 13.0) ^ 2.0)) < 4.0) * FBinderD) |
| FluorB_init_uM | (7.2984378812835775 * ((((x - 23.0) ^ 2.0) + ((y - 39.0) ^ 2.0) + ((z - 13.0) ^ 2.0)) < 4.0)) |
| J_BindD | (((Kf_BindD * Dark) * Binder) - (Kr_BindD * DarkB)) |
| J_BindF | (((Kf_BindF * Fluor) * Binder) - (Kr_BindF * FluorB)) |
| J_BleachBound | ((Kf_BleachBound * FluorB) - (Kr_BleachBound * DarkB)) |
| J_BleachFree | ((Kf_BleachFree * Fluor) - (Kr_BleachFree * Dark)) |
| Kf_BleachBound | ((kbleach_BleachBound * Laser) * (t > start) * (t < (start + duration))) |
| Kf_BleachFree | ((kbleach_BleachFree * Laser) * (t > start) * (t < (start + duration))) |
| Laser_init_uM | ((exp( - (((z - 13.0) ^ 2.0) / (2.0 * (sigmaaxial ^ 2.0)))) * ((((x - 23.0) ^ 2.0) + ((y - 39.0) ^ 2.0)) < 4.0)) * (x > BleachX)) |
| Size_CYT | (VolumePerUnitVolume_CYT * vcRegionVolume('cytosol')) |
| Size_EC | (VolumePerUnitVolume_EC * vcRegionVolume('ec')) |
| Size_NM | (AreaPerUnitArea_NM * vcRegionArea('Nucleus_cytosol_membrane')) |
| Size_NUC | (VolumePerUnitVolume_NUC * vcRegionVolume('Nucleus')) |
| Size_PM | (AreaPerUnitArea_PM * vcRegionArea('cytosol_ec_membrane')) |
| sobj_cytosol1_ec0_size | vcRegionArea('cytosol_ec_membrane') |
| sobj_Nucleus2_cytosol1_size | vcRegionArea('Nucleus_cytosol_membrane') |
| vobj_cytosol1_size | vcRegionVolume('cytosol') |

| Function Name | Expression |
| --- | --- |
| vobj_ec0_size | vcRegionVolume('ec') |
| vobj_Nucleus2_size | vcRegionVolume('Nucleus') |

#### 2.9.5.3. Volume Domains

### 2.9.5.3.1. ec

##### 2.9.5.3.2. cytosol

| PdeEquation Fluor | |
| --- | --- |
| Rate | ( - J_BindF - J_BleachFree) |
| Diffusion | Fluor_diffusionRate |
| Initial | Fluor_init_uM |

| PdeEquation Dark | |
| --- | --- |
| Rate | ( - J_BindD + J_BleachFree) |
| Diffusion | Dark_diffusionRate |
| Initial | Dark_init_uM |

| PdeEquation Binder | |
| --- | --- |
| Rate | ( - J_BindD - J_BindF) |
| Diffusion | Binder_diffusionRate |
| Initial | Binder_init_uM |

| PdeEquation FluorB | |
| --- | --- |
| Rate | (J_BindF - J_BleachBound) |
| Diffusion | FluorB_diffusionRate |
| Initial | FluorB_init_uM |

| OdeEquation Laser |
| --- |

| OdeEquation Laser | |
|---|---|
| Rate | 0.0 |
| Initial | Laser_init_uM |

| PdeEquation DarkB | |
|---|---|
| Rate | (J_BindD + J_BleachBound) |
| Diffusion | DarkB_diffusionRate |
| Initial | DarkB_init_uM |

###### 2.9.5.3.3. Nucleus

##### 2.9.5.4. Membrane Domains

###### 2.9.5.4.1. cytosol_ec_membrane

| JumpCondition Fluor | |
|---|---|
| InFlux | 0.0 |
| OutFlux | 0.0 |

| JumpCondition Dark | |
|---|---|
| InFlux | 0.0 |
| OutFlux | 0.0 |

| JumpCondition Binder | |
|---|---|
| InFlux | 0.0 |
| OutFlux | 0.0 |

| JumpCondition FluorB | |
|---|---|
| InFlux | 0.0 |
| OutFlux | 0.0 |

| JumpCondition DarkB | |
|---|---|
| InFlux | 0.0 |
| OutFlux | 0.0 |

##### 2.9.5.4.2. Nucleus_cytosol_membrane

| JumpCondition Fluor | |
|---|---|
| InFlux | 0.0 |
| OutFlux | 0.0 |

| JumpCondition Dark | |
|---|---|
| InFlux | 0.0 |
| OutFlux | 0.0 |

| JumpCondition Binder | |
|---|---|
| InFlux | 0.0 |
| OutFlux | 0.0 |

| JumpCondition FluorB | |
|---|---|
| InFlux | 0.0 |
| OutFlux | 0.0 |

| JumpCondition DarkB | |
|---|---|
| InFlux | 0.0 |
| OutFlux | 0.0 |

#### 2.9.6. Simulation(s)

##### 2.9.6.1. 2Photon Bleach w fast binding_1

| Simulation Name: 2Photon Bleach w fast binding_1 |
|---|

**Simulation Description: cloned from '2Photon Bleach w fast binding_1' owned by user temp**
**cloned from '2Photon Bleach w fast binding_1' owned by user temp**
**cloned from '2Photon Bleach w fast binding_1' owned by user temp**
**cloned from '2Photon Bleach w binding_1' owned by user temp**

| Overriden Parameters | | |
|---|---|---|
| **Name** | **Actual Value** | **Default Value** |
| kbleach_BleachFree | 10.0 | 1.0 |
| Kf_BindD | 100.0 | 1.0 |
| FBinderD | "0.0", "0.1", "1.0" | 0.1 |
| kbleach_BleachBound | 10.0 | 1.0 |
| Kr_BindF | 100.0 | 1.0 |
| Kr_BindD | 100.0 | 1.0 |
| BinderD | FBinderD | 0.1 |
| Fluor_diffusionRate | "3.0", "25.0" | 10.0 |
| Dark_diffusionRate | Fluor_diffusionRate | 10.0 |
| Kf_BindF | 100.0 | 1.0 |

| Geometry Setting | |
|---|---|
| Geometry Size (um) | (74.24, 74.24, 26.0) |
| Mesh Size (elements) | (257, 257, 35) |

| Advanced Settings | |
|---|---|
| Solver Name | Fully-Implicit Finite Volume, Regular Grid (Variable Time Step) |
| Time Bounds - Starting | 0.0 |
| Time Bounds - Ending | 100.0 |

| | |
|---|---|
| Time Step - Min | 0.0 |
| Time Step - Default | 0.05 |
| Time Step - Max | 0.1 |
| Error Tolerance - Absolute | 1.0E-9 |
| Error Tolerance - Relative | 1.0E-7 |
| Output Time Step | 1.0 |
| Use Symbolic Jacobian (T/F) | F |

#### 2.9.6.2. Widefield Bleach w fast binding_1

| **Simulation Name: Widefield Bleach w fast binding_1** |
|---|
| **Simulation Description: cloned from 'Widefield Bleach w fast binding_1' owned by user temp<br>cloned from 'Widefield Bleach w fast binding_1' owned by user temp<br>cloned from 'Widefield Bleach w fast binding_1' owned by user temp<br>cloned from 'Widefield Bleach w binding_1' owned by user temp** |

| **Overriden Parameters** | | |
|---|---|---|
| **Name** | **Actual Value** | **Default Value** |
| kbleach_BleachFree | 10.0 | 1.0 |
| Kf_BindD | 100.0 | 1.0 |
| FBinderD | "0.0", "0.1", "1.0" | 0.1 |
| kbleach_BleachBound | 10.0 | 1.0 |
| Kr_BindF | 100.0 | 1.0 |
| Kr_BindD | 100.0 | 1.0 |
| BinderD | FBinderD | 0.1 |
| Fluor_diffusionRate | "3.0", "25.0" | 10.0 |
| Dark_diffusionRate | Fluor_diffusionRate | 10.0 |
| sigmaaxial | 30.0 | 1.5 |

| | | |
|---|---|---|
| Kf_BindF | 100.0 | 1.0 |

| Geometry Setting | |
|---|---|
| Geometry Size (um) | (74.24, 74.24, 26.0) |
| Mesh Size (elements) | (256, 256, 35) |

| Advanced Settings | |
|---|---|
| Solver Name | Fully-Implicit Finite Volume, Regular Grid (Variable Time Step) |
| Time Bounds - Starting | 0.0 |
| Time Bounds - Ending | 100.0 |
| Time Step - Min | 0.0 |
| Time Step - Default | 0.05 |
| Time Step - Max | 0.1 |
| Error Tolerance - Absolute | 1.0E-9 |
| Error Tolerance - Relative | 1.0E-7 |
| Output Time Step | 1.0 |
| Use Symbolic Jacobian (T/F) | F |

##### 2.9.6.3. 2Photon Bleach w default binding_1

| **Simulation Name: 2Photon Bleach w default binding_1** |
|---|
| **Simulation Description: cloned from '2Photon Bleach w default binding_1' owned by user temp<br>cloned from '2Photon Bleach w default binding_1' owned by user temp<br>cloned from '2Photon Bleach w default binding_1' owned by user temp<br>cloned from '2Photon Bleach w slower binding' owned by user temp<br>cloned from '2Photon Bleach w binding_1' owned by user temp** |

| Overriden Parameters |
|---|

| Name | Actual Value | Default Value |
|---|---|---|
| BinderD | FBinderD | 0.1 |
| Fluor_diffusionRate | "3.0", "25.0" | 10.0 |
| FBinderD | "0.0", "0.1", "1.0" | 0.1 |
| kbleach_BleachFree | 10.0 | 1.0 |
| kbleach_BleachBound | 10.0 | 1.0 |
| Dark_diffusionRate | Fluor_diffusionRate | 10.0 |

| Geometry Setting | |
|---|---|
| Geometry Size (um) | (74.24, 74.24, 26.0) |
| Mesh Size (elements) | (256, 256, 35) |

| Advanced Settings | |
|---|---|
| Solver Name | Fully-Implicit Finite Volume, Regular Grid (Variable Time Step) |
| Time Bounds - Starting | 0.0 |
| Time Bounds - Ending | 100.0 |
| Time Step - Min | 0.0 |
| Time Step - Default | 0.05 |
| Time Step - Max | 0.1 |
| Error Tolerance - Absolute | 1.0E-9 |
| Error Tolerance - Relative | 1.0E-7 |
| Output Time Step | 1.0 |
| Use Symbolic Jacobian (T/F) | F |

##### 2.9.6.4. Widefield Bleach w defalt binding_1

| **Simulation Name: Widefield Bleach w defalt binding_1** |
|---|
| **Simulation Description: cloned from 'Widefield Bleach w defalt binding_1' owned by user temp<br>cloned from 'Widefield Bleach w defalt binding_1' owned by user temp<br>cloned from 'Widefield Bleach w defalt binding_1' owned by user temp<br>cloned from 'Widefield Bleach w slower binding' owned by user temp<br>cloned from 'Widefield Bleach w binding_1' owned by user temp** |

| **Overriden Parameters** | | |
|---|---|---|
| **Name** | **Actual Value** | **Default Value** |
| BinderD | FBinderD | 0.1 |
| Fluor_diffusionRate | "1.0", "10.0" | 10.0 |
| FBinderD | "0.0", "0.1", "1.0" | 0.1 |
| sigmaaxial | 30.0 | 1.5 |
| kbleach_BleachFree | 10.0 | 1.0 |
| kbleach_BleachBound | 10.0 | 1.0 |
| Dark_diffusionRate | Fluor_diffusionRate | 10.0 |

| **Geometry Setting** | |
|---|---|
| Geometry Size (um) | (74.24, 74.24, 26.0) |
| Mesh Size (elements) | (256, 256, 35) |

| **Advanced Settings** | |
|---|---|
| Solver Name | Fully-Implicit Finite Volume, Regular Grid (Variable Time Step) |
| Time Bounds - Starting | 0.0 |
| Time Bounds - Ending | 100.0 |
| Time Step - Min | 0.0 |

| | |
|---|---|
| Time Step - Default | 0.05 |
| Time Step - Max | 0.1 |
| Error Tolerance - Absolute | 1.0E-9 |
| Error Tolerance - Relative | 1.0E-7 |
| Output Time Step | 1.0 |
| Use Symbolic Jacobian (T/F) | F |

#### 2.9.6.5. 2Photon Bleach w slow binding_1

| **Simulation Name: 2Photon Bleach w slow binding_1** |
|---|
| **Simulation Description: cloned from '2Photon Bleach w slow binding_1' owned by user temp<br>cloned from '2Photon Bleach w slow binding_1' owned by user temp<br>cloned from '2Photon Bleach w slow binding_1' owned by user temp<br>cloned from 'Copy of 2Photon Bleach w slow binding' owned by user temp<br>cloned from '2Photon Bleach w slower binding' owned by user temp<br>cloned from '2Photon Bleach w binding_1' owned by user temp** |

| **Overriden Parameters** | | |
|---|---|---|
| **Name** | **Actual Value** | **Default Value** |
| kbleach_BleachFree | 10.0 | 1.0 |
| Kf_BindD | 0.1 | 1.0 |
| FBinderD | "0.0", "0.1", "1.0" | 0.1 |
| kbleach_BleachBound | 10.0 | 1.0 |
| Kr_BindF | 0.1 | 1.0 |
| Kr_BindD | 0.1 | 1.0 |
| BinderD | FBinderD | 0.1 |
| Fluor_diffusionRate | "3.0", "25.0" | 10.0 |
| Dark_diffusionRate | Fluor_diffusionRate | 10.0 |
| Kf_BindF | 0.1 | 1.0 |

| Geometry Setting | |
|---|---|
| Geometry Size (um) | (74.24, 74.24, 26.0) |
| Mesh Size (elements) | (256, 256, 35) |

| Advanced Settings | |
|---|---|
| Solver Name | Fully-Implicit Finite Volume, Regular Grid (Variable Time Step) |
| Time Bounds - Starting | 0.0 |
| Time Bounds - Ending | 100.0 |
| Time Step - Min | 0.0 |
| Time Step - Default | 0.05 |
| Time Step - Max | 0.1 |
| Error Tolerance - Absolute | 1.0E-9 |
| Error Tolerance - Relative | 1.0E-7 |
| Output Time Step | 1.0 |
| Use Symbolic Jacobian (T/F) | F |

#### 2.9.6.6. Widefield Bleach w slow binding_1

| **Simulation Name: Widefield Bleach w slow binding_1** |
|---|
| **Simulation Description: cloned from 'Widefield Bleach w slow binding_1' owned by user temp<br>cloned from 'Widefield Bleach w slow binding_1' owned by user temp<br>cloned from 'Widefield Bleach w slow binding_1' owned by user temp<br>cloned from 'Copy of Widefield Bleach w slow binding' owned by user temp<br>cloned from 'Widefield Bleach w slower binding' owned by user temp<br>cloned from 'Widefield Bleach w binding_1' owned by user temp** |

| **Overriden Parameters** | | |
|---|---|---|
| **Name** | **Actual Value** | **Default Value** |

| | | |
|---|---|---|
| kbleach_BleachFree | 10.0 | 1.0 |
| Kf_BindD | 0.1 | 1.0 |
| FBinderD | "0.0", "0.1", "1.0" | 0.1 |
| kbleach_BleachBound | 10.0 | 1.0 |
| Kr_BindF | 0.1 | 1.0 |
| Kr_BindD | 0.1 | 1.0 |
| BinderD | FBinderD | 0.1 |
| Fluor_diffusionRate | "3.0", "25.0" | 10.0 |
| Dark_diffusionRate | Fluor_diffusionRate | 10.0 |
| sigmaaxial | 30.0 | 1.5 |
| Kf_BindF | 0.1 | 1.0 |

| **Geometry Setting** | |
|---|---|
| Geometry Size (um) | (74.24, 74.24, 26.0) |
| Mesh Size (elements) | (256, 256, 35) |

| **Advanced Settings** | |
|---|---|
| Solver Name | Fully-Implicit Finite Volume, Regular Grid (Variable Time Step) |
| Time Bounds - Starting | 0.0 |
| Time Bounds - Ending | 100.0 |
| Time Step - Min | 0.0 |
| Time Step - Default | 0.05 |
| Time Step - Max | 0.1 |
| Error Tolerance - Absolute | 1.0E-9 |
| Error Tolerance - Relative | 1.0E-7 |

| | |
|---|---|
| Output Time Step | 1.0 |
| Use Symbolic Jacobian (T/F) | F |

#### 2.9.6.7. Half Widefield Bleach w fast  binding_1

| **Simulation Name: Half Widefield Bleach w fast binding_1** |
|---|
| **Simulation Description: cloned from 'Half Widefield Bleach w fast binding_1' owned by user temp<br>cloned from 'Half Widefield Bleach w fast binding_1' owned by user temp<br>cloned from 'Half Widefield Bleach w fast binding_1' owned by user temp<br>cloned from 'Widefield Bleach w fast binding_1' owned by user temp<br>cloned from 'Widefield Bleach w binding_1' owned by user temp** |

| **Overriden Parameters** | | |
|---|---|---|
| **Name** | **Actual Value** | **Default Value** |
| kbleach_BleachFree | 10.0 | 1.0 |
| Kf_BindD | 100.0 | 1.0 |
| FBinderD | "0.0", "0.1", "1.0" | 0.1 |
| kbleach_BleachBound | 10.0 | 1.0 |
| BleachX | 23.0 | 0.0 |
| Kr_BindF | 100.0 | 1.0 |
| Kr_BindD | 100.0 | 1.0 |
| BinderD | FBinderD | 0.1 |
| Fluor_diffusionRate | "3.0", "25.0" | 10.0 |
| sigmaaxial | 30.0 | 1.5 |
| Dark_diffusionRate | Fluor_diffusionRate | 10.0 |
| Kf_BindF | 100.0 | 1.0 |

| **Geometry Setting** | |
|---|---|
| Geometry Size (um) | (74.24, 74.24, 26.0) |

| | |
|---|---|
| Mesh Size (elements) | (256, 256, 35) |

| Advanced Settings | |
|---|---|
| Solver Name | Fully-Implicit Finite Volume, Regular Grid (Variable Time Step) |
| Time Bounds - Starting | 0.0 |
| Time Bounds - Ending | 100.0 |
| Time Step - Min | 0.0 |
| Time Step - Default | 0.05 |
| Time Step - Max | 0.1 |
| Error Tolerance - Absolute | 1.0E-9 |
| Error Tolerance - Relative | 1.0E-7 |
| Output Time Step | 1.0 |
| Use Symbolic Jacobian (T/F) | F |

##### 2.9.6.8. Half Widefield Bleach w slow binding_1

| Simulation Name: Half Widefield Bleach w slow binding_1 |
|---|
| **Simulation Description: cloned from 'Half Widefield Bleach w slow binding_1' owned by user temp<br>cloned from 'Half Widefield Bleach w slow binding_1' owned by user temp<br>cloned from 'Half Widefield Bleach w slow binding_1' owned by user temp<br>cloned from 'Widefield Bleach w slow binding_1' owned by user temp<br>cloned from 'Copy of Widefield Bleach w slow binding' owned by user temp<br>cloned from 'Widefield Bleach w slower binding' owned by user temp<br>cloned from 'Widefield Bleach w binding_1' owned by user temp** |

| Overriden Parameters | | |
|---|---|---|
| **Name** | **Actual Value** | **Default Value** |
| kbleach_BleachFree | 10.0 | 1.0 |
| Kf_BindD | 0.1 | 1.0 |

| | | |
|---|---|---|
| FBinderD | "0.0", "0.1", "1.0" | 0.1 |
| kbleach_BleachBound | 10.0 | 1.0 |
| BleachX | 23.0 | 0.0 |
| Kr_BindF | 0.1 | 1.0 |
| Kr_BindD | 0.1 | 1.0 |
| BinderD | FBinderD | 0.1 |
| Fluor_diffusionRate | "3.0", "25.0" | 10.0 |
| sigmaaxial | 30.0 | 1.5 |
| Dark_diffusionRate | Fluor_diffusionRate | 10.0 |
| Kf_BindF | 0.1 | 1.0 |

| **Geometry Setting** | |
|---|---|
| Geometry Size (um) | (74.24, 74.24, 26.0) |
| Mesh Size (elements) | (256, 256, 35) |

| **Advanced Settings** | |
|---|---|
| Solver Name | Fully-Implicit Finite Volume, Regular Grid (Variable Time Step) |
| Time Bounds - Starting | 0.0 |
| Time Bounds - Ending | 100.0 |
| Time Step - Min | 0.0 |
| Time Step - Default | 0.05 |
| Time Step - Max | 0.1 |
| Error Tolerance - Absolute | 1.0E-9 |
| Error Tolerance - Relative | 1.0E-7 |
| Output Time Step | 1.0 |

| | |
|---|---|
| Use Symbolic Jacobian (T/F) | F |

#### 2.9.6.9. Half Widefield Bleach w slow binding_short

| **Simulation Name: Half Widefield Bleach w slow binding_short** |
|---|
| **Simulation Description: cloned from 'Copy of Half Widefield Bleach w slow binding_1' owned by user temp<br>cloned from 'Copy of Half Widefield Bleach w slow binding_1' owned by user temp<br>cloned from 'Copy of Half Widefield Bleach w slow binding_1' owned by user temp<br>cloned from 'Half Widefield Bleach w slow binding_1' owned by user temp<br>cloned from 'Widefield Bleach w slow binding_1' owned by user temp<br>cloned from 'Copy of Widefield Bleach w slow binding' owned by user temp<br>cloned from 'Widefield Bleach w slower binding' owned by user temp<br>cloned from 'Widefield Bleach w binding_1' owned by user temp** |

| **Overriden Parameters** | | |
|---|---|---|
| **Name** | **Actual Value** | **Default Value** |
| kbleach_BleachFree | 10.0 | 1.0 |
| Kf_BindD | 0.1 | 1.0 |
| FBinderD | "0.0", "0.1", "1.0" | 0.1 |
| kbleach_BleachBound | 10.0 | 1.0 |
| BleachX | 23.0 | 0.0 |
| Kr_BindF | 0.1 | 1.0 |
| Kr_BindD | 0.1 | 1.0 |
| BinderD | FBinderD | 0.1 |
| Fluor_diffusionRate | "3.0", "25.0" | 10.0 |
| sigmaaxial | 30.0 | 1.5 |
| Dark_diffusionRate | Fluor_diffusionRate | 10.0 |
| Kf_BindF | 0.1 | 1.0 |

| **Geometry Setting** |
|---|

| | |
|---|---|
| Geometry Size (um) | (74.24, 74.24, 26.0) |
| Mesh Size (elements) | (256, 256, 35) |

| **Advanced Settings** | |
|---|---|
| Solver Name | Fully-Implicit Finite Volume, Regular Grid (Variable Time Step) |
| Time Bounds - Starting | 0.0 |
| Time Bounds - Ending | 20.0 |
| Time Step - Min | 0.0 |
| Time Step - Default | 0.05 |
| Time Step - Max | 0.1 |
| Error Tolerance - Absolute | 1.0E-9 |
| Error Tolerance - Relative | 1.0E-7 |
| Output Time Step | 0.2 |
| Use Symbolic Jacobian (T/F) | F |

##### 2.9.6.10. Half Widefield Bleach w very slow binding_1 1

| **Simulation Name: Half Widefield Bleach w very slow binding_1 1** |
|---|
| **Simulation Description: cloned from 'Half Widefield Bleach w very slow binding_1 1' owned by user temp<br>cloned from 'Half Widefield Bleach w very slow binding_1 1' owned by user temp<br>cloned from 'Half Widefield Bleach w slow binding_1' owned by user temp<br>cloned from 'Half Widefield Bleach w slow binding_1' owned by user temp<br>cloned from 'Widefield Bleach w slow binding_1' owned by user temp<br>cloned from 'Copy of Widefield Bleach w slow binding' owned by user temp<br>cloned from 'Widefield Bleach w slower binding' owned by user temp<br>cloned from 'Widefield Bleach w binding_1' owned by user temp** |

| **Overriden Parameters** | | |
|---|---|---|
| **Name** | **Actual Value** | **Default Value** |

| | | |
|---|---|---|
| kbleach_BleachFree | 10.0 | 1.0 |
| Kf_BindD | 0.01 | 1.0 |
| FBinderD | "0.0", "0.1", "1.0" | 0.1 |
| kbleach_BleachBound | 10.0 | 1.0 |
| BleachX | 23.0 | 0.0 |
| Kr_BindF | 0.01 | 1.0 |
| Kr_BindD | 0.01 | 1.0 |
| BinderD | FBinderD | 0.1 |
| Fluor_diffusionRate | "3.0", "25.0" | 10.0 |
| sigmaaxial | 30.0 | 1.5 |
| Dark_diffusionRate | Fluor_diffusionRate | 10.0 |
| Kf_BindF | 0.01 | 1.0 |

| **Geometry Setting** | |
|---|---|
| Geometry Size (um) | (74.24, 74.24, 26.0) |
| Mesh Size (elements) | (256, 256, 35) |

| **Advanced Settings** | |
|---|---|
| Solver Name | Fully-Implicit Finite Volume, Regular Grid (Variable Time Step) |
| Time Bounds - Starting | 0.0 |
| Time Bounds - Ending | 100.0 |
| Time Step - Min | 0.0 |
| Time Step - Default | 0.05 |
| Time Step - Max | 0.1 |
| Error Tolerance - Absolute | 1.0E-9 |

| | |
|---|---|
| Error Tolerance - Relative | 1.0E-7 |
| Output Time Step | 1.0 |
| Use Symbolic Jacobian (T/F) | F |

#### 2.9.6.11. Half Widefield Bleach w fast binding_short

| **Simulation Name: Half Widefield Bleach w fast binding_short** |
|---|
| **Simulation Description: cloned from 'Copy of Half Widefield Bleach w fast binding_1' owned by user temp<br>cloned from 'Copy of Half Widefield Bleach w fast binding_1' owned by user temp<br>cloned from 'Half Widefield Bleach w fast binding_1' owned by user temp<br>cloned from 'Half Widefield Bleach w fast binding_1' owned by user temp<br>cloned from 'Widefield Bleach w fast binding_1' owned by user temp<br>cloned from 'Widefield Bleach w binding_1' owned by user temp** |

| **Overriden Parameters** | | |
|---|---|---|
| **Name** | **Actual Value** | **Default Value** |
| kbleach_BleachFree | 10.0 | 1.0 |
| Kf_BindD | 100.0 | 1.0 |
| FBinderD | "0.0", "0.1", "1.0" | 0.1 |
| kbleach_BleachBound | 10.0 | 1.0 |
| BleachX | 23.0 | 0.0 |
| Kr_BindF | 100.0 | 1.0 |
| Kr_BindD | 100.0 | 1.0 |
| BinderD | FBinderD | 0.1 |
| Fluor_diffusionRate | "3.0", "25.0" | 10.0 |
| sigmaaxial | 30.0 | 1.5 |
| Dark_diffusionRate | Fluor_diffusionRate | 10.0 |
| Kf_BindF | 100.0 | 1.0 |

| Geometry Setting | |
|---|---|
| Geometry Size (um) | (74.24, 74.24, 26.0) |
| Mesh Size (elements) | (256, 256, 35) |

| Advanced Settings | |
|---|---|
| Solver Name | Fully-Implicit Finite Volume, Regular Grid (Variable Time Step) |
| Time Bounds - Starting | 0.0 |
| Time Bounds - Ending | 20.0 |
| Time Step - Min | 0.0 |
| Time Step - Default | 0.05 |
| Time Step - Max | 0.1 |
| Error Tolerance - Absolute | 1.0E-9 |
| Error Tolerance - Relative | 1.0E-7 |
| Output Time Step | 0.2 |
| Use Symbolic Jacobian (T/F) | F |

##### 2.9.6.12. Half Widefield Bleach w very slow binding_short

| **Simulation Name: Half Widefield Bleach w very slow binding_short** |
|---|
| **Simulation Description: cloned from 'Half Widefield Bleach w very slow binding_short' owned by user temp<br>cloned from 'Half Widefield Bleach w very slow binding_1 1' owned by user temp<br>cloned from 'Half Widefield Bleach w very slow binding_1 1' owned by user temp<br>cloned from 'Half Widefield Bleach w slow binding_1' owned by user temp<br>cloned from 'Half Widefield Bleach w slow binding_1' owned by user temp<br>cloned from 'Widefield Bleach w slow binding_1' owned by user temp<br>cloned from 'Copy of Widefield Bleach w slow binding' owned by user temp<br>cloned from 'Widefield Bleach w slower binding' owned by user temp<br>cloned from 'Widefield Bleach w binding_1' owned by user temp** |

| Overriden Parameters | | |
|---|---|---|
| **Name** | **Actual Value** | **Default Value** |
| kbleach_BleachFree | 10.0 | 1.0 |
| Kf_BindD | 0.01 | 1.0 |
| FBinderD | "0.0", "0.1", "1.0" | 0.1 |
| kbleach_BleachBound | 10.0 | 1.0 |
| BleachX | 23.0 | 0.0 |
| Kr_BindF | 0.01 | 1.0 |
| Kr_BindD | 0.01 | 1.0 |
| BinderD | FBinderD | 0.1 |
| Fluor_diffusionRate | "3.0", "25.0" | 10.0 |
| Dark_diffusionRate | Fluor_diffusionRate | 10.0 |
| sigmaaxial | 30.0 | 1.5 |
| Kf_BindF | 0.01 | 1.0 |

| Geometry Setting | |
|---|---|
| Geometry Size (um) | (74.24, 74.24, 26.0) |
| Mesh Size (elements) | (256, 256, 35) |

| Advanced Settings | |
|---|---|
| Solver Name | Fully-Implicit Finite Volume, Regular Grid (Variable Time Step) |
| Time Bounds - Starting | 0.0 |
| Time Bounds - Ending | 20.0 |
| Time Step - Min | 0.0 |

| | |
|---|---|
| Time Step - Default | 0.05 |
| Time Step - Max | 0.1 |
| Error Tolerance - Absolute | 1.0E-9 |
| Error Tolerance - Relative | 1.0E-7 |
| Output Time Step | 0.2 |
| Use Symbolic Jacobian (T/F) | F |

##### 2.9.6.13. Widefield Bleach w very fast binding_1

| Simulation Name: Widefield Bleach w very fast binding_1 |
|---|
| **Simulation Description: cloned from 'Widefield Bleach w very fast binding_1' owned by user temp<br>cloned from 'Widefield Bleach w fast binding_1' owned by user temp<br>cloned from 'Widefield Bleach w fast binding_1' owned by user temp<br>cloned from 'Widefield Bleach w fast binding_1' owned by user temp<br>cloned from 'Widefield Bleach w binding_1' owned by user temp** |

| Overriden Parameters | | |
|---|---|---|
| **Name** | **Actual Value** | **Default Value** |
| kbleach_BleachFree | 10.0 | 1.0 |
| Kf_BindD | 1000.0 | 1.0 |
| FBinderD | "0.0", "0.1", "1.0" | 0.1 |
| kbleach_BleachBound | 10.0 | 1.0 |
| Kr_BindF | 1000.0 | 1.0 |
| Kr_BindD | 1000.0 | 1.0 |
| BinderD | FBinderD | 0.1 |
| Fluor_diffusionRate | "3.0", "25.0" | 10.0 |
| Dark_diffusionRate | Fluor_diffusionRate | 10.0 |
| sigmaaxial | 30.0 | 1.5 |

| | | |
|---|---|---|
| Kf_BindF | 1000.0 | 1.0 |

| Geometry Setting | |
|---|---|
| Geometry Size (um) | (74.24, 74.24, 26.0) |
| Mesh Size (elements) | (256, 256, 35) |

| Advanced Settings | |
|---|---|
| Solver Name | Fully-Implicit Finite Volume, Regular Grid (Variable Time Step) |
| Time Bounds - Starting | 0.0 |
| Time Bounds - Ending | 100.0 |
| Time Step - Min | 0.0 |
| Time Step - Default | 0.05 |
| Time Step - Max | 0.1 |
| Error Tolerance - Absolute | 1.0E-9 |
| Error Tolerance - Relative | 1.0E-7 |
| Output Time Step | 1.0 |
| Use Symbolic Jacobian (T/F) | F |

#### 2.10. Application: Image-based binding to liquid droplet, FRAP away from droplet

| **Application Name: Image-based binding to liquid droplet, FRAP away from droplet** |
|---|
| **Application Description: (copied from Image-based binding to liquid droplet) (copied from Image-based center circular bleach) (copied from Image-based center bleach) (copied from Spherical_Cell_Gaussian_Bleach)** |

##### 2.10.1. Structure Mapping For Image-based binding to liquid droplet, FRAP away from droplet

| Structure Mapping |
|---|

| Structure Mapping | | | | |
|---|---|---|---|---|
| **Structure** | **Subdomain** | **Resolved (T/F)** | **Surf/Vol** | **VolFract** |
| EC | ec | F | | |
| NUC | Nucleus | F | | |
| CYT | cytosol | F | | |

#### 2.10.2. Reaction Mapping For Image-based binding to liquid droplet, FRAP away from droplet

| Reaction Mapping | | | |
|---|---|---|---|
| **Name** | **Type** | **Enabled (T/F)** | **Fast (T/F)** |
| BindD | Reaction | T | F |
| BindF | Reaction | T | F |
| BleachFree | Reaction | T | F |
| BleachBound | Reaction | T | F |

| Initial Conditions | | | | |
|---|---|---|---|---|
| **Species** | **Structure** | **Initial Conc.** | **Diffusion Const.** | **Fixed (T/F)** |
| s0 | CYT | 2.701562118716424 M | 10.0 $m^2.s^1$ | F |
| s1 | CYT | 0.0 M | 10.0 $m^2.s^1$ | F |
| s2 | CYT | (2.701562118716425 * ((((x - 23.0) ^ 2.0) + ((y - 39.0) ^ 2.0) + ((z - 13.0) ^ 2.0)) < 4.0)) M | (((((x - 23.0) ^ 2.0) + ((y - 39.0) ^ 2.0) + ((z - 13.0) ^ 2.0)) < 4.0) * BinderD) $m^2.s^1$ | F |
| s3 | CYT | (7.2984378812835775 * ((((x - 23.0) ^ 2.0) + ((y - 39.0) ^ 2.0) + ((z - 13.0) ^ 2.0)) < 4.0)) M | (((((x - 23.0) ^ 2.0) + ((y - 39.0) ^ 2.0) + ((z - 13.0) ^ 2.0)) < 4.0) * FBinderD) $m^2.s^1$ | F |
| s4 | CYT | (exp( - (((z - 10.0) ^ 2.0) / (2.0 * (sigmaaxial ^ 2.0)))) * ((((x - 21.0) | 0.0 $m^2.s^1$ | F |

| Initial Conditions | | | | |
|---|---|---|---|---|
| Species | Structure | Initial Conc. | Diffusion Const. | Fixed (T/F) |
| | | ^ 2.0) + ((y - 30.0) ^ 2.0)) < (BleachRadius ^ 2.0))) M | | |
| s5 | CYT | 0.0 M | (((((x - 23.0) ^ 2.0) + ((y - 39.0) ^ 2.0) + ((z - 13.0) ^ 2.0)) < 4.0) * FBinderD) $m^2.s^1$ | F |

#### 2.10.3. Membrane Mapping For Image-based binding to liquid droplet, FRAP away from droplet

| Electrical Mapping - Membrane Potential | | | |
|---|---|---|---|
| Membrane | Calculate V (T/F) | V initial | Specific Capacitance |
| PM | F | 0.0 mV | 1.0 $pF.m^2$ |
| NM | F | 0.0 mV | 1.0 $pF.m^2$ |

| |
|---|
| Temperature: 300.0 K |

#### 2.10.4. Geometry: Site visit _Application0_20111127_695607844

| | |
|---|---|
| Size | (74.24, 74.24, 26.0) |
| Origin | (0.0, 0.0, 0.0) |

#### 2.10.5. Math Description: Copy of Image-based binding to liquid droplet_generated

##### 2.10.5.1. Constants

| Constant Name | Expression |
|---|---|
| _F_ | 96485.3321 |
| _F_nmol_ | 9.64853321E-5 |
| _K_GHK_ | 1.0E-9 |
| _N_pmol_ | 6.02214179E11 |

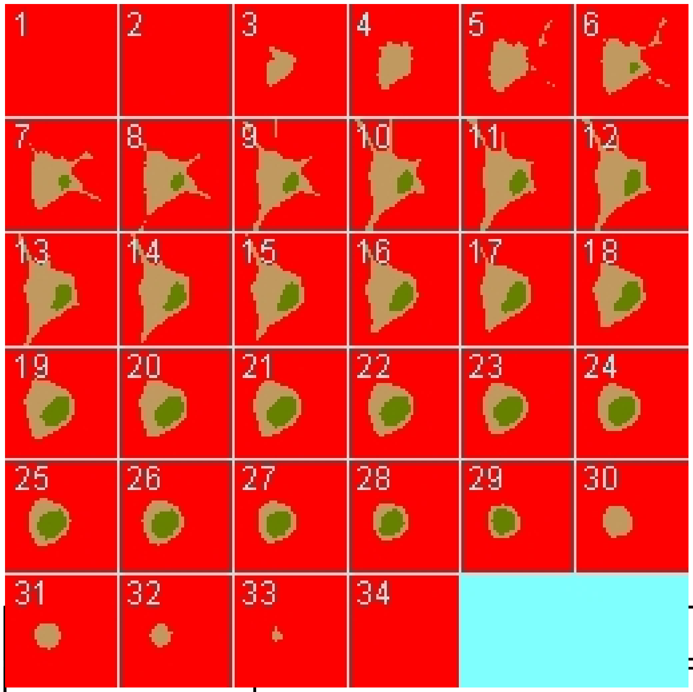

| Constant Name | Expression |
|---|---|
| _PI_ | 3.141592653589793 |
| _R_ | 8314.46261815 |
| _T_ | 300.0 |
| AreaPerUnitArea_NM | 1.0 |
| AreaPerUnitArea_PM | 1.0 |
| BinderD | 0.1 |
| BleachRadius | 2.0 |
| BleachX | 0.0 |
| Dark_diffusionRate | 10.0 |
| Dark_init_uM | 0.0 |
| DarkB_init_uM | 0.0 |
| duration | 1.0 |
| FBinderD | 0.1 |
| Fluor_diffusionRate | 10.0 |
| Fluor_init_uM | 2.701562118716424 |
| K_millivolts_per_volt | 1000.0 |

| Constant Name | Expression |
|---|---|
| kbleach_BleachBound | 1.0 |
| kbleach_BleachFree | 1.0 |
| Kf_BindD | 1.0 |
| Kf_BindF | 1.0 |
| KMOLE | 0.001660538783162726 |
| Kr_BindD | 1.0 |
| Kr_BindF | 1.0 |
| Kr_BleachBound | 0.0 |
| Kr_BleachFree | 0.0 |
| sigmaaxial | 1.5 |
| sigmalateral | 0.5 |
| start | 1.0 |
| Voltage_NM | 0.0 |
| Voltage_PM | 0.0 |
| VolumePerUnitVolume_CYT | 1.0 |
| VolumePerUnitVolume_EC | 1.0 |
| VolumePerUnitVolume_NUC | 1.0 |

#### 2.10.5.2. Functions

| Function Name | Expression |
|---|---|
| Binder_diffusionRate | (((((x - 23.0) ^ 2.0) + ((y - 39.0) ^ 2.0) + ((z - 13.0) ^ 2.0)) < 4.0) * BinderD) |
| Binder_init_uM | (2.701562118716425 * ((((x - 23.0) ^ 2.0) + ((y - 39.0) ^ 2.0) + ((z - 13.0) ^ 2.0)) < 4.0)) |
| DarkB_diffusionRate | (((((x - 23.0) ^ 2.0) + ((y - 39.0) ^ 2.0) + ((z - 13.0) ^ 2.0)) < 4.0) * FBinderD) |
| FluorB_diffusionRate | (((((x - 23.0) ^ 2.0) + ((y - 39.0) ^ 2.0) + ((z - 13.0) ^ 2.0)) < 4.0) * FBinderD) |
| FluorB_init_uM | (7.2984378812835775 * ((((x - 23.0) ^ 2.0) + ((y - 39.0) ^ 2.0) + ((z - 13.0) ^ 2.0)) < 4.0)) |
| J_BindD | (((Kf_BindD * Dark) * Binder) - (Kr_BindD * DarkB)) |

| Function Name | Expression |
|---|---|
| J_BindF | (((Kf_BindF * Fluor) * Binder) - (Kr_BindF * FluorB)) |
| J_BleachBound | ((Kf_BleachBound * FluorB) - (Kr_BleachBound * DarkB)) |
| J_BleachFree | ((Kf_BleachFree * Fluor) - (Kr_BleachFree * Dark)) |
| Kf_BleachBound | ((kbleach_BleachBound * Laser) * (t > start) * (t < (start + duration))) |
| Kf_BleachFree | ((kbleach_BleachFree * Laser) * (t > start) * (t < (start + duration))) |
| Laser_init_uM | (exp( - (((z - 10.0) ^ 2.0) / (2.0 * (sigmaaxial ^ 2.0)))) * ((((x - 21.0) ^ 2.0) + ((y - 30.0) ^ 2.0)) < (BleachRadius ^ 2.0))) |
| Size_CYT | (VolumePerUnitVolume_CYT * vcRegionVolume('cytosol')) |
| Size_EC | (VolumePerUnitVolume_EC * vcRegionVolume('ec')) |
| Size_NM | (AreaPerUnitArea_NM * vcRegionArea('Nucleus_cytosol_membrane')) |
| Size_NUC | (VolumePerUnitVolume_NUC * vcRegionVolume('Nucleus')) |
| Size_PM | (AreaPerUnitArea_PM * vcRegionArea('cytosol_ec_membrane')) |
| sobj_cytosol1_ec0_size | vcRegionArea('cytosol_ec_membrane') |
| sobj_Nucleus2_cytosol1_size | vcRegionArea('Nucleus_cytosol_membrane') |
| vobj_cytosol1_size | vcRegionVolume('cytosol') |
| vobj_ec0_size | vcRegionVolume('ec') |
| vobj_Nucleus2_size | vcRegionVolume('Nucleus') |

#### 2.10.5.3. Volume Domains

### 2.10.5.3.1. ec

##### 2.10.5.3.2. cytosol

| PdeEquation Fluor | |
|---|---|
| Rate | ( - J_BindF - J_BleachFree) |
| Diffusion | Fluor_diffusionRate |
| Initial | Fluor_init_uM |

| PdeEquation Dark | |
|---|---|
| Rate | ( - J_BindD + J_BleachFree) |
| Diffusion | Dark_diffusionRate |
| Initial | Dark_init_uM |

| PdeEquation Binder | |
|---|---|
| Rate | ( - J_BindD - J_BindF) |
| Diffusion | Binder_diffusionRate |
| Initial | Binder_init_uM |

| PdeEquation FluorB | |
|---|---|
| Rate | (J_BindF - J_BleachBound) |
| Diffusion | FluorB_diffusionRate |
| Initial | FluorB_init_uM |

| OdeEquation Laser | |
|---|---|
| Rate | 0.0 |
| Initial | Laser_init_uM |

| PdeEquation DarkB | |
|---|---|
| Rate | (J_BindD + J_BleachBound) |
| Diffusion | DarkB_diffusionRate |
| Initial | DarkB_init_uM |

##### 2.10.5.3.3. Nucleus

##### 2.10.5.4. Membrane Domains

##### 2.10.5.4.1. cytosol_ec_membrane

| JumpCondition Fluor | |
|---|---|
| InFlux | 0.0 |
| OutFlux | 0.0 |

| JumpCondition Dark | |
|---|---|
| InFlux | 0.0 |
| OutFlux | 0.0 |

| JumpCondition Binder | |
|---|---|
| InFlux | 0.0 |
| OutFlux | 0.0 |

| JumpCondition FluorB | |
|---|---|
| InFlux | 0.0 |
| OutFlux | 0.0 |

| JumpCondition DarkB | |
|---|---|
| InFlux | 0.0 |
| OutFlux | 0.0 |

##### 2.10.5.4.2. Nucleus_cytosol_membrane

| JumpCondition Fluor | |
|---|---|
| InFlux | 0.0 |
| OutFlux | 0.0 |

| JumpCondition Dark | |
|---|---|
| InFlux | 0.0 |
| OutFlux | 0.0 |

| JumpCondition Binder | |
|---|---|
| InFlux | 0.0 |
| OutFlux | 0.0 |

| JumpCondition FluorB | |
|---|---|
| InFlux | 0.0 |
| OutFlux | 0.0 |

| JumpCondition DarkB | |
|---|---|
| InFlux | 0.0 |
| OutFlux | 0.0 |

#### 2.10.6. Simulation(s)

##### 2.10.6.1. 2Photon Bleach w fast binding_1_1

| Simulation Name: 2Photon Bleach w fast binding_1_1 |
|---|
| Simulation Description: cloned from '2Photon Bleach w fast binding_1_1' owned by user temp<br>cloned from '2Photon Bleach w fast binding_1' owned by user temp<br>cloned from '2Photon Bleach w binding_1' owned by user temp |

| Overriden Parameters | | |
|---|---|---|
| **Name** | **Actual Value** | **Default Value** |
| Fluor_diffusionRate | "1.0", "10.0" | 10.0 |
| Kf_BindF | 100.0 | 1.0 |
| Kr_BindF | 100.0 | 1.0 |
| Kf_BindD | 100.0 | 1.0 |
| Kr_BindD | 100.0 | 1.0 |
| kbleach_BleachFree | 10.0 | 1.0 |

| | | |
|---|---|---|
| kbleach_BleachBound | 10.0 | 1.0 |
| Dark_diffusionRate | Fluor_diffusionRate | 10.0 |

| Geometry Setting | |
|---|---|
| Geometry Size (um) | (74.24, 74.24, 26.0) |
| Mesh Size (elements) | (257, 257, 35) |

| Advanced Settings | |
|---|---|
| Solver Name | Fully-Implicit Finite Volume, Regular Grid (Variable Time Step) |
| Time Bounds - Starting | 0.0 |
| Time Bounds - Ending | 100.0 |
| Time Step - Min | 0.0 |
| Time Step - Default | 0.05 |
| Time Step - Max | 0.1 |
| Error Tolerance - Absolute | 1.0E-9 |
| Error Tolerance - Relative | 1.0E-7 |
| Output Time Step | 1.0 |
| Use Symbolic Jacobian (T/F) | F |

#### 2.10.6.2. Widefield Bleach w fast binding_1_1

| **Simulation Name: Widefield Bleach w fast binding_1_1** |
|---|
| **Simulation Description: cloned from 'Widefield Bleach w fast binding_1_1' owned by user temp<br>cloned from 'Widefield Bleach w fast binding_1' owned by user temp<br>cloned from 'Widefield Bleach w binding_1' owned by user temp** |

| Overriden Parameters |
|---|

| Name | Actual Value | Default Value |
|---|---|---|
| kbleach_BleachFree | 10.0 | 1.0 |
| Kf_BindD | 100.0 | 1.0 |
| kbleach_BleachBound | 10.0 | 1.0 |
| Kr_BindF | 100.0 | 1.0 |
| Kr_BindD | 100.0 | 1.0 |
| Fluor_diffusionRate | "1.0", "10.0" | 10.0 |
| Dark_diffusionRate | Fluor_diffusionRate | 10.0 |
| sigmaaxial | 30.0 | 1.5 |
| Kf_BindF | 100.0 | 1.0 |

| Geometry Setting | |
|---|---|
| Geometry Size (um) | (74.24, 74.24, 26.0) |
| Mesh Size (elements) | (256, 256, 35) |

| Advanced Settings | |
|---|---|
| Solver Name | Fully-Implicit Finite Volume, Regular Grid (Variable Time Step) |
| Time Bounds - Starting | 0.0 |
| Time Bounds - Ending | 100.0 |
| Time Step - Min | 0.0 |
| Time Step - Default | 0.05 |
| Time Step - Max | 0.1 |
| Error Tolerance - Absolute | 1.0E-9 |
| Error Tolerance - Relative | 1.0E-7 |
| Output Time Step | 1.0 |

| | |
|---|---|
| Use Symbolic Jacobian (T/F) | F |

#### 2.10.6.3. 2Photon Bleach w default binding_1_1

| **Simulation Name: 2Photon Bleach w default binding_1_1** |
|---|
| **Simulation Description: cloned from '2Photon Bleach w default binding_1_1' owned by user temp<br>cloned from '2Photon Bleach w default binding_1' owned by user temp<br>cloned from '2Photon Bleach w slower binding' owned by user temp<br>cloned from '2Photon Bleach w binding_1' owned by user temp** |

| **Overriden Parameters** | | |
|---|---|---|
| **Name** | **Actual Value** | **Default Value** |
| Fluor_diffusionRate | "1.0", "10.0" | 10.0 |
| kbleach_BleachFree | 10.0 | 1.0 |
| kbleach_BleachBound | 10.0 | 1.0 |
| Dark_diffusionRate | Fluor_diffusionRate | 10.0 |

| **Geometry Setting** | |
|---|---|
| Geometry Size (um) | (74.24, 74.24, 26.0) |
| Mesh Size (elements) | (256, 256, 35) |

| **Advanced Settings** | |
|---|---|
| Solver Name | Fully-Implicit Finite Volume, Regular Grid (Variable Time Step) |
| Time Bounds - Starting | 0.0 |
| Time Bounds - Ending | 100.0 |
| Time Step - Min | 0.0 |
| Time Step - Default | 0.05 |
| Time Step - Max | 0.1 |

| | |
|---|---|
| Error Tolerance - Absolute | 1.0E-9 |
| Error Tolerance - Relative | 1.0E-7 |
| Output Time Step | 1.0 |
| Use Symbolic Jacobian (T/F) | F |

#### 2.10.6.4. Widefield Bleach w defalt binding_1_1

| **Simulation Name: Widefield Bleach w defalt binding_1_1** |
|---|
| **Simulation Description: cloned from 'Widefield Bleach w defalt binding_1_1' owned by user temp<br>cloned from 'Widefield Bleach w defalt binding_1' owned by user temp<br>cloned from 'Widefield Bleach w slower binding' owned by user temp<br>cloned from 'Widefield Bleach w binding_1' owned by user temp** |

| **Overriden Parameters** | | |
|---|---|---|
| **Name** | **Actual Value** | **Default Value** |
| Fluor_diffusionRate | "1.0", "10.0" | 10.0 |
| kbleach_BleachFree | 10.0 | 1.0 |
| sigmaaxial | 30.0 | 1.5 |
| kbleach_BleachBound | 10.0 | 1.0 |
| Dark_diffusionRate | Fluor_diffusionRate | 10.0 |

| **Geometry Setting** | |
|---|---|
| Geometry Size (um) | (74.24, 74.24, 26.0) |
| Mesh Size (elements) | (256, 256, 35) |

| **Advanced Settings** | |
|---|---|
| Solver Name | Fully-Implicit Finite Volume, Regular Grid (Variable Time Step) |
| Time Bounds - Starting | 0.0 |

| | |
|---|---|
| Time Bounds - Ending | 100.0 |
| Time Step - Min | 0.0 |
| Time Step - Default | 0.05 |
| Time Step - Max | 0.1 |
| Error Tolerance - Absolute | 1.0E-9 |
| Error Tolerance - Relative | 1.0E-7 |
| Output Time Step | 1.0 |
| Use Symbolic Jacobian (T/F) | F |

##### 2.10.6.5. 2Photon Bleach w slow binding_1_1

| **Simulation Name: 2Photon Bleach w slow binding_1_1** |
|---|
| **Simulation Description: cloned from '2Photon Bleach w slow binding_1_1' owned by user temp<br>cloned from '2Photon Bleach w slow binding_1' owned by user temp<br>cloned from 'Copy of 2Photon Bleach w slow binding' owned by user temp<br>cloned from '2Photon Bleach w slower binding' owned by user temp<br>cloned from '2Photon Bleach w binding_1' owned by user temp** |

| **Overriden Parameters** | | |
|---|---|---|
| **Name** | **Actual Value** | **Default Value** |
| Fluor_diffusionRate | "1.0", "10.0" | 10.0 |
| Kf_BindF | 0.1 | 1.0 |
| Kr_BindF | 0.1 | 1.0 |
| Kf_BindD | 0.1 | 1.0 |
| Kr_BindD | 0.1 | 1.0 |
| kbleach_BleachFree | 10.0 | 1.0 |
| kbleach_BleachBound | 10.0 | 1.0 |
| Dark_diffusionRate | Fluor_diffusionRate | 10.0 |

| Geometry Setting | |
| --- | --- |
| Geometry Size (um) | (74.24, 74.24, 26.0) |
| Mesh Size (elements) | (256, 256, 35) |

| Advanced Settings | |
| --- | --- |
| Solver Name | Fully-Implicit Finite Volume, Regular Grid (Variable Time Step) |
| Time Bounds - Starting | 0.0 |
| Time Bounds - Ending | 100.0 |
| Time Step - Min | 0.0 |
| Time Step - Default | 0.05 |
| Time Step - Max | 0.1 |
| Error Tolerance - Absolute | 1.0E-9 |
| Error Tolerance - Relative | 1.0E-7 |
| Output Time Step | 1.0 |
| Use Symbolic Jacobian (T/F) | F |

##### 2.10.6.6. Widefield Bleach w slow binding_1_1

| **Simulation Name: Widefield Bleach w slow binding_1_1** |
| --- |
| **Simulation Description: cloned from 'Widefield Bleach w slow binding_1_1' owned by user temp<br>cloned from 'Widefield Bleach w slow binding_1' owned by user temp<br>cloned from 'Copy of Widefield Bleach w slow binding' owned by user temp<br>cloned from 'Widefield Bleach w slower binding' owned by user temp<br>cloned from 'Widefield Bleach w binding_1' owned by user temp** |

| Overriden Parameters | | |
| --- | --- | --- |
| **Name** | **Actual Value** | **Default Value** |

| | | |
|---|---|---|
| kbleach_BleachFree | 10.0 | 1.0 |
| Kf_BindD | 0.1 | 1.0 |
| kbleach_BleachBound | 10.0 | 1.0 |
| Kr_BindF | 0.1 | 1.0 |
| Kr_BindD | 0.1 | 1.0 |
| Fluor_diffusionRate | "1.0", "10.0" | 10.0 |
| Dark_diffusionRate | Fluor_diffusionRate | 10.0 |
| sigmaaxial | 30.0 | 1.5 |
| Kf_BindF | 0.1 | 1.0 |

| Geometry Setting | |
|---|---|
| Geometry Size (um) | (74.24, 74.24, 26.0) |
| Mesh Size (elements) | (256, 256, 35) |

| Advanced Settings | |
|---|---|
| Solver Name | Fully-Implicit Finite Volume, Regular Grid (Variable Time Step) |
| Time Bounds - Starting | 0.0 |
| Time Bounds - Ending | 100.0 |
| Time Step - Min | 0.0 |
| Time Step - Default | 0.05 |
| Time Step - Max | 0.1 |
| Error Tolerance - Absolute | 1.0E-9 |
| Error Tolerance - Relative | 1.0E-7 |
| Output Time Step | 1.0 |
| Use Symbolic Jacobian (T/F) | F |

#### 2.11. Application: ProjectZImage-based binding to liquid droplet, FRAP adjacent to droplet

| Application Name: ProjectZImage-based binding to liquid droplet, FRAP adjacent to droplet |
|---|
| Application Description: (copied from Image-based binding to liquid droplet, FRAP away from droplet) (copied from Image-based binding to liquid droplet) (copied from Image-based center circular bleach) (copied from Image-based center bleach) (copied from Spherical_Cell_Gaussian_Bleach) |

##### 2.11.1. Structure Mapping For ProjectZImage-based binding to liquid droplet, FRAP adjacent to droplet

| Structure Mapping | | | | |
|---|---|---|---|---|
| **Structure** | **Subdomain** | **Resolved (T/F)** | **Surf/Vol** | **VolFract** |
| EC | ec | F | | |
| NUC | Nucleus | F | | |
| CYT | cytosol | F | | |

##### 2.11.2. Reaction Mapping For ProjectZImage-based binding to liquid droplet, FRAP adjacent to droplet

| Reaction Mapping | | | |
|---|---|---|---|
| **Name** | **Type** | **Enabled (T/F)** | **Fast (T/F)** |
| BindD | Reaction | T | F |
| BindF | Reaction | T | F |
| BleachFree | Reaction | T | F |
| BleachBound | Reaction | T | F |

| Initial Conditions | | | | |
|---|---|---|---|---|
| **Species** | **Structure** | **Initial Conc.** | **Diffusion Const.** | **Fixed (T/F)** |

| Initial Conditions | | | | |
|---|---|---|---|---|
| **Species** | **Structure** | **Initial Conc.** | **Diffusion Const.** | **Fixed (T/F)** |
| s0 | CYT | 2.701562118716424 M | 10.0 $m^{2}.s^{1}$ | F |
| s1 | CYT | 0.0 M | 10.0 $m^{2}.s^{1}$ | F |
| s2 | CYT | (2.701562118716425 * ((((x - 23.0) ^ 2.0) + ((y - 39.0) ^ 2.0) + ((z - 13.0) ^ 2.0)) < 4.0)) M | (((((x - 23.0) ^ 2.0) + ((y - 39.0) ^ 2.0) + ((z - 13.0) ^ 2.0)) < 4.0) * BinderD) $m^{2}.s^{1}$ | F |
| s3 | CYT | (7.2984378812835775 * ((((x - 23.0) ^ 2.0) + ((y - 39.0) ^ 2.0) + ((z - 13.0) ^ 2.0)) < 4.0)) M | (((((x - 23.0) ^ 2.0) + ((y - 39.0) ^ 2.0) + ((z - 13.0) ^ 2.0)) < 4.0) * FBinderD) $m^{2}.s^{1}$ | F |
| s4 | CYT | (exp( - (((z - 10.0) ^ 2.0) / (2.0 * (sigmaaxial ^ 2.0)))) * ((((x - 17.0) ^ 2.0) + ((y - 39.0) ^ 2.0)) < (BleachRadius ^ 2.0))) M | 0.0 $m^{2}.s^{1}$ | F |
| s5 | CYT | 0.0 M | (((((x - 23.0) ^ 2.0) + ((y - 39.0) ^ 2.0) + ((z - 13.0) ^ 2.0)) < 4.0) * FBinderD) $m^{2}.s^{1}$ | F |

#### 2.11.3. Membrane Mapping For ProjectZImage-based binding to liquid droplet, FRAP adjacent to droplet

| Electrical Mapping - Membrane Potential | | | |
|---|---|---|---|
| **Membrane** | **Calculate V (T/F)** | **V initial** | **Specific Capacitance** |
| PM | F | 0.0 mV | 1.0 $pF.m^{2}$ |
| NM | F | 0.0 mV | 1.0 $pF.m^{2}$ |

| |
|---|
| Temperature: 300.0 K |

#### 2.11.4. Geometry: Site visit _Application0_20111127_695607844

| | |
|---|---|
| Size | (74.24, 74.24, 26.0) |
| Origin | (0.0, 0.0, 0.0) |

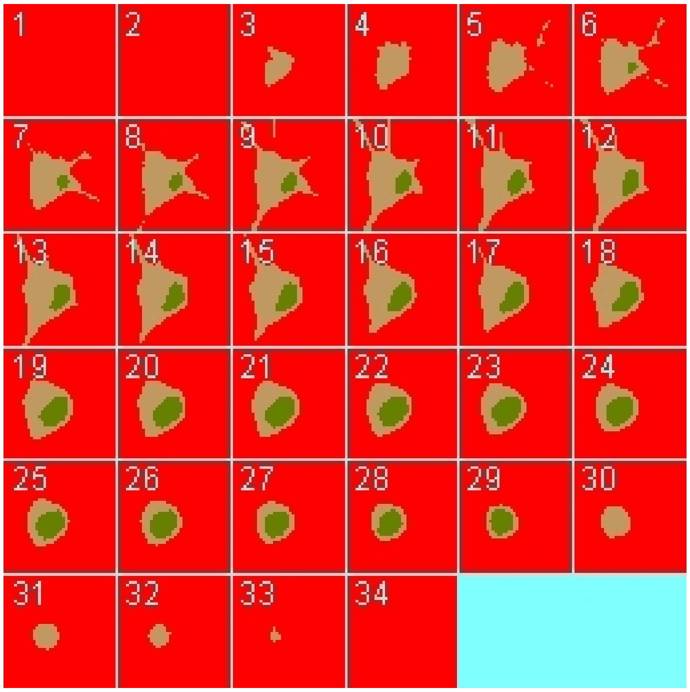

#### 2.11.5. Math Description: Copy of Image-based binding to liquid droplet, FRAP away from droplet_generated

##### 2.11.5.1. Constants

| Constant Name | Expression |
|---|---|
| _F_ | 96485.3321 |
| _F_nmol_ | 9.64853321E-5 |
| _K_GHK_ | 1.0E-9 |
| _N_pmol_ | 6.02214179E11 |
| _PI_ | 3.141592653589793 |
| _R_ | 8314.46261815 |
| _T_ | 300.0 |
| AreaPerUnitArea_NM | 1.0 |
| AreaPerUnitArea_PM | 1.0 |
| BinderD | 0.1 |

| Constant Name | Expression |
|---|---|
| BleachRadius | 2.0 |
| BleachX | 0.0 |
| Dark_diffusionRate | 10.0 |
| Dark_init_uM | 0.0 |
| DarkB_init_uM | 0.0 |
| duration | 1.0 |
| FBinderD | 0.1 |
| Fluor_diffusionRate | 10.0 |
| Fluor_init_uM | 2.701562118716424 |
| K_millivolts_per_volt | 1000.0 |
| kbleach_BleachBound | 1.0 |
| kbleach_BleachFree | 1.0 |
| Kf_BindD | 1.0 |
| Kf_BindF | 1.0 |
| KMOLE | 0.001660538783162726 |
| Kr_BindD | 1.0 |
| Kr_BindF | 1.0 |
| Kr_BleachBound | 0.0 |
| Kr_BleachFree | 0.0 |
| sigmaaxial | 1.5 |
| sigmalateral | 0.5 |
| start | 1.0 |
| UnitFactor_molecules_um_neg_3_uM_neg_1 | (1.0 * pow(KMOLE,-1.0)) |
| Voltage_NM | 0.0 |
| Voltage_PM | 0.0 |
| VolumePerUnitVolume_CYT | 1.0 |

| Constant Name | Expression |
| --- | --- |
| VolumePerUnitVolume_EC | 1.0 |
| VolumePerUnitVolume_NUC | 1.0 |

##### 2.11.5.2. Functions

| Function Name | Expression |
| --- | --- |
| Binder_diffusionRate | (((((x - 23.0) ^ 2.0) + ((y - 39.0) ^ 2.0) + ((z - 13.0) ^ 2.0)) < 4.0) * BinderD) |
| Binder_init_uM | (2.701562118716425 * ((((x - 23.0) ^ 2.0) + ((y - 39.0) ^ 2.0) + ((z - 13.0) ^ 2.0)) < 4.0)) |
| DarkB_diffusionRate | (((((x - 23.0) ^ 2.0) + ((y - 39.0) ^ 2.0) + ((z - 13.0) ^ 2.0)) < 4.0) * FBinderD) |
| FluorB_diffusionRate | (((((x - 23.0) ^ 2.0) + ((y - 39.0) ^ 2.0) + ((z - 13.0) ^ 2.0)) < 4.0) * FBinderD) |
| FluorB_init_uM | (7.2984378812835775 * ((((x - 23.0) ^ 2.0) + ((y - 39.0) ^ 2.0) + ((z - 13.0) ^ 2.0)) < 4.0)) |
| J_BindD | (((Kf_BindD * Dark) * Binder) - (Kr_BindD * DarkB)) |
| J_BindF | (((Kf_BindF * Fluor) * Binder) - (Kr_BindF * FluorB)) |
| J_BleachBound | ((Kf_BleachBound * FluorB) - (Kr_BleachBound * DarkB)) |
| J_BleachFree | ((Kf_BleachFree * Fluor) - (Kr_BleachFree * Dark)) |
| Kf_BleachBound | ((kbleach_BleachBound * Laser) * (t > start) * (t < (start + duration))) |
| Kf_BleachFree | ((kbleach_BleachFree * Laser) * (t > start) * (t < (start + duration))) |
| Laser_init_uM | (exp( - (((z - 10.0) ^ 2.0) / (2.0 * (sigmaaxial ^ 2.0)))) * ((((x - 17.0) ^ 2.0) + ((y - 39.0) ^ 2.0)) < (BleachRadius ^ 2.0))) |
| Size_CYT | (VolumePerUnitVolume_CYT * vcRegionVolume('cytosol')) |
| Size_EC | (VolumePerUnitVolume_EC * vcRegionVolume('ec')) |
| Size_NM | (AreaPerUnitArea_NM * vcRegionArea('Nucleus_cytosol_membrane')) |
| Size_NUC | (VolumePerUnitVolume_NUC * vcRegionVolume('Nucleus')) |
| Size_PM | (AreaPerUnitArea_PM * vcRegionArea('cytosol_ec_membrane')) |
| sobj_cytosol1_ec0_size | vcRegionArea('cytosol_ec_membrane') |
| sobj_Nucleus2_cytosol1_size | vcRegionArea('Nucleus_cytosol_membrane') |
| vobj_cytosol1_size | vcRegionVolume('cytosol') |

| Function Name | Expression |
| --- | --- |
| vobj_ec0_size | vcRegionVolume('ec') |
| vobj_Nucleus2_size | vcRegionVolume('Nucleus') |

#### 2.11.5.3. Volume Domains

### 2.11.5.3.1. ec

##### 2.11.5.3.2. cytosol

| PdeEquation Fluor | |
| --- | --- |
| Rate | ( - J_BindF - J_BleachFree) |
| Diffusion | Fluor_diffusionRate |
| Initial | Fluor_init_uM |

| PdeEquation Dark | |
| --- | --- |
| Rate | ( - J_BindD + J_BleachFree) |
| Diffusion | Dark_diffusionRate |
| Initial | Dark_init_uM |

| PdeEquation Binder | |
| --- | --- |
| Rate | ( - J_BindD - J_BindF) |
| Diffusion | Binder_diffusionRate |
| Initial | Binder_init_uM |

| PdeEquation FluorB | |
| --- | --- |
| Rate | (J_BindF - J_BleachBound) |
| Diffusion | FluorB_diffusionRate |
| Initial | FluorB_init_uM |

| OdeEquation Laser |
| --- |

| OdeEquation Laser | |
|---|---|
| Rate | 0.0 |
| Initial | Laser_init_uM |

| PdeEquation DarkB | |
|---|---|
| Rate | (J_BindD + J_BleachBound) |
| Diffusion | DarkB_diffusionRate |
| Initial | DarkB_init_uM |

##### 2.11.5.3.3. Nucleus

#### 2.11.5.4. Membrane Domains

##### 2.11.5.4.1. cytosol_ec_membrane

| JumpCondition Fluor | |
|---|---|
| InFlux | 0.0 |
| OutFlux | 0.0 |

| JumpCondition Dark | |
|---|---|
| InFlux | 0.0 |
| OutFlux | 0.0 |

| JumpCondition Binder | |
|---|---|
| InFlux | 0.0 |
| OutFlux | 0.0 |

| JumpCondition FluorB | |
|---|---|
| InFlux | 0.0 |
| OutFlux | 0.0 |

| JumpCondition DarkB | |
|---|---|
| InFlux | 0.0 |
| OutFlux | 0.0 |

##### 2.11.5.4.2. Nucleus_cytosol_membrane

| JumpCondition Fluor | |
|---|---|
| InFlux | 0.0 |
| OutFlux | 0.0 |

| JumpCondition Dark | |
|---|---|
| InFlux | 0.0 |
| OutFlux | 0.0 |

| JumpCondition Binder | |
|---|---|
| InFlux | 0.0 |
| OutFlux | 0.0 |

| JumpCondition FluorB | |
|---|---|
| InFlux | 0.0 |
| OutFlux | 0.0 |

| JumpCondition DarkB | |
|---|---|
| InFlux | 0.0 |
| OutFlux | 0.0 |

#### 2.11.6. Simulation(s)

##### 2.11.6.1. 2Photon Bleach w fast binding_1_1_1

| Simulation Name: 2Photon Bleach w fast binding_1_1_1 |
|---|

**Simulation  Description: cloned from '2Photon Bleach w fast binding_1_1_1' owned by user temp**
**cloned from '2Photon Bleach w fast binding_1_1' owned by user temp**
**cloned from '2Photon Bleach w fast binding_1' owned by user temp**
**cloned from '2Photon Bleach w binding_1' owned by user temp**

| **Overriden Parameters** | | |
|---|---|---|
| **Name** | **Actual Value** | **Default Value** |
| Fluor_diffusionRate | "1.0", "10.0" | 10.0 |
| Kf_BindF | 100.0 | 1.0 |
| Kr_BindF | 100.0 | 1.0 |
| Kf_BindD | 100.0 | 1.0 |
| Kr_BindD | 100.0 | 1.0 |
| kbleach_BleachFree | 10.0 | 1.0 |
| kbleach_BleachBound | 10.0 | 1.0 |
| Dark_diffusionRate | Fluor_diffusionRate | 10.0 |

| **Geometry Setting** | |
|---|---|
| Geometry Size (um) | (74.24, 74.24, 26.0) |
| Mesh Size (elements) | (257, 257, 35) |

| **Advanced Settings** | |
|---|---|
| Solver Name | Fully-Implicit Finite Volume, Regular Grid (Variable Time Step) |
| Time Bounds - Starting | 0.0 |
| Time Bounds - Ending | 100.0 |
| Time Step - Min | 0.0 |
| Time Step - Default | 0.05 |

| | |
|---|---|
| Time Step - Max | 0.1 |
| Error Tolerance - Absolute | 1.0E-9 |
| Error Tolerance - Relative | 1.0E-7 |
| Output Time Step | 1.0 |
| Use Symbolic Jacobian (T/F) | F |

#### 2.11.6.2. Widefield Bleach w fast  binding_1_1_1

| **Simulation Name: Widefield Bleach w fast binding_1_1_1** |
|---|
| **Simulation Description: cloned from 'Widefield Bleach w fast binding_1_1_1' owned by user temp<br>cloned from 'Widefield Bleach w fast binding_1_1_1' owned by user temp<br>cloned from 'Widefield Bleach w fast binding_1_1' owned by user temp<br>cloned from 'Widefield Bleach w fast binding_1' owned by user temp<br>cloned from 'Widefield Bleach w binding_1' owned by user temp** |

| **Overriden Parameters** | | |
|---|---|---|
| **Name** | **Actual Value** | **Default Value** |
| kbleach_BleachFree | 10.0 | 1.0 |
| Kf_BindD | 100.0 | 1.0 |
| kbleach_BleachBound | 10.0 | 1.0 |
| Kr_BindF | 100.0 | 1.0 |
| Kr_BindD | 100.0 | 1.0 |
| Fluor_diffusionRate | "3.0", "25.0" | 10.0 |
| Dark_diffusionRate | Fluor_diffusionRate | 10.0 |
| sigmaaxial | 30.0 | 1.5 |
| Kf_BindF | 100.0 | 1.0 |

| **Geometry Setting** | |
|---|---|
| Geometry Size (um) | (74.24, 74.24, 26.0) |

| | |
|---|---|
| Mesh Size (elements) | (256, 256, 35) |

| Advanced Settings | |
|---|---|
| Solver Name | Fully-Implicit Finite Volume, Regular Grid (Variable Time Step) |
| Time Bounds - Starting | 0.0 |
| Time Bounds - Ending | 100.0 |
| Time Step - Min | 0.0 |
| Time Step - Default | 0.05 |
| Time Step - Max | 0.1 |
| Error Tolerance - Absolute | 1.0E-9 |
| Error Tolerance - Relative | 1.0E-7 |
| Output Time Step | 1.0 |
| Use Symbolic Jacobian (T/F) | F |

##### 2.11.6.3. 2Photon Bleach w default binding_1_1_1

| Simulation Name: 2Photon Bleach w default binding_1_1_1 |
|---|
| Simulation Description: cloned from '2Photon Bleach w default binding_1_1_1' owned by user temp<br>cloned from '2Photon Bleach w default binding_1_1' owned by user temp<br>cloned from '2Photon Bleach w default binding_1' owned by user temp<br>cloned from '2Photon Bleach w slower binding' owned by user temp<br>cloned from '2Photon Bleach w binding_1' owned by user temp |

| Overriden Parameters | | |
|---|---|---|
| **Name** | **Actual Value** | **Default Value** |
| Fluor_diffusionRate | "1.0", "10.0" | 10.0 |
| kbleach_BleachFree | 10.0 | 1.0 |
| kbleach_BleachBound | 10.0 | 1.0 |

| | | |
|---|---|---|
| Dark_diffusionRate | Fluor_diffusionRate | 10.0 |

| Geometry Setting | |
|---|---|
| Geometry Size (um) | (74.24, 74.24, 26.0) |
| Mesh Size (elements) | (256, 256, 35) |

| Advanced Settings | |
|---|---|
| Solver Name | Fully-Implicit Finite Volume, Regular Grid (Variable Time Step) |
| Time Bounds - Starting | 0.0 |
| Time Bounds - Ending | 100.0 |
| Time Step - Min | 0.0 |
| Time Step - Default | 0.05 |
| Time Step - Max | 0.1 |
| Error Tolerance - Absolute | 1.0E-9 |
| Error Tolerance - Relative | 1.0E-7 |
| Output Time Step | 1.0 |
| Use Symbolic Jacobian (T/F) | F |

#### 2.11.6.4. Widefield Bleach w defalt binding_1_1_1

| Simulation Name: Widefield Bleach w defalt binding_1_1_1 |
|---|
| Simulation Description: cloned from 'Widefield Bleach w defalt binding_1_1_1' owned by user temp<br>cloned from 'Widefield Bleach w defalt binding_1_1' owned by user temp<br>cloned from 'Widefield Bleach w defalt binding_1' owned by user temp<br>cloned from 'Widefield Bleach w slower binding' owned by user temp<br>cloned from 'Widefield Bleach w binding_1' owned by user temp |

| Overriden Parameters |
|---|

| Name | Actual Value | Default Value |
|---|---|---|
| Fluor_diffusionRate | "1.0", "10.0" | 10.0 |
| kbleach_BleachFree | 10.0 | 1.0 |
| sigmaaxial | 30.0 | 1.5 |
| kbleach_BleachBound | 10.0 | 1.0 |
| Dark_diffusionRate | Fluor_diffusionRate | 10.0 |

| Geometry Setting | |
|---|---|
| Geometry Size (um) | (74.24, 74.24, 26.0) |
| Mesh Size (elements) | (256, 256, 35) |

| Advanced Settings | |
|---|---|
| Solver Name | Fully-Implicit Finite Volume, Regular Grid (Variable Time Step) |
| Time Bounds - Starting | 0.0 |
| Time Bounds - Ending | 100.0 |
| Time Step - Min | 0.0 |
| Time Step - Default | 0.05 |
| Time Step - Max | 0.1 |
| Error Tolerance - Absolute | 1.0E-9 |
| Error Tolerance - Relative | 1.0E-7 |
| Output Time Step | 1.0 |
| Use Symbolic Jacobian (T/F) | F |

##### 2.11.6.5. 2Photon Bleach w slow binding_1_1_1

| Simulation Name: 2Photon Bleach w slow binding_1_1_1 |
|---|

**Simulation Description: cloned from '2Photon Bleach w slow binding_1_1_1' owned by user temp**
**cloned from '2Photon Bleach w slow binding_1_1' owned by user temp**
**cloned from '2Photon Bleach w slow binding_1' owned by user temp**
**cloned from 'Copy of 2Photon Bleach w slow binding' owned by user temp**
**cloned from '2Photon Bleach w slower binding' owned by user temp**
**cloned from '2Photon Bleach w binding_1' owned by user temp**

| **Overriden Parameters** | | |
|---|---|---|
| **Name** | **Actual Value** | **Default Value** |
| Fluor_diffusionRate | "1.0", "10.0" | 10.0 |
| Kf_BindF | 0.1 | 1.0 |
| Kr_BindF | 0.1 | 1.0 |
| Kf_BindD | 0.1 | 1.0 |
| Kr_BindD | 0.1 | 1.0 |
| kbleach_BleachFree | 10.0 | 1.0 |
| kbleach_BleachBound | 10.0 | 1.0 |
| Dark_diffusionRate | Fluor_diffusionRate | 10.0 |

| **Geometry Setting** | |
|---|---|
| Geometry Size (um) | (74.24, 74.24, 26.0) |
| Mesh Size (elements) | (256, 256, 35) |

| **Advanced Settings** | |
|---|---|
| Solver Name | Fully-Implicit Finite Volume, Regular Grid (Variable Time Step) |
| Time Bounds - Starting | 0.0 |
| Time Bounds - Ending | 100.0 |
| Time Step - Min | 0.0 |

| | |
|---|---|
| Time Step - Default | 0.05 |
| Time Step - Max | 0.1 |
| Error Tolerance - Absolute | 1.0E-9 |
| Error Tolerance - Relative | 1.0E-7 |
| Output Time Step | 1.0 |
| Use Symbolic Jacobian (T/F) | F |

#### 2.11.6.6. Widefield Bleach w slow binding_1_1_1

| **Simulation Name: Widefield Bleach w slow binding_1_1_1** |
|---|
| **Simulation Description: cloned from 'Widefield Bleach w slow binding_1_1_1' owned by user temp<br>cloned from 'Widefield Bleach w slow binding_1_1_1' owned by user temp<br>cloned from 'Widefield Bleach w slow binding_1_1' owned by user temp<br>cloned from 'Widefield Bleach w slow binding_1' owned by user temp<br>cloned from 'Copy of Widefield Bleach w slow binding' owned by user temp<br>cloned from 'Widefield Bleach w slower binding' owned by user temp<br>cloned from 'Widefield Bleach w binding_1' owned by user temp** |

| **Overriden Parameters** | | |
|---|---|---|
| **Name** | **Actual Value** | **Default Value** |
| kbleach_BleachFree | 10.0 | 1.0 |
| Kf_BindD | 0.1 | 1.0 |
| kbleach_BleachBound | 10.0 | 1.0 |
| Kr_BindF | 0.1 | 1.0 |
| Kr_BindD | 0.1 | 1.0 |
| Fluor_diffusionRate | "3.0", "25.0" | 10.0 |
| Dark_diffusionRate | Fluor_diffusionRate | 10.0 |
| sigmaaxial | 30.0 | 1.5 |
| Kf_BindF | 0.1 | 1.0 |

| Geometry Setting | |
|---|---|
| Geometry Size (um) | (74.24, 74.24, 26.0) |
| Mesh Size (elements) | (256, 256, 35) |

| Advanced Settings | |
|---|---|
| Solver Name | Fully-Implicit Finite Volume, Regular Grid (Variable Time Step) |
| Time Bounds - Starting | 0.0 |
| Time Bounds - Ending | 100.0 |
| Time Step - Min | 0.0 |
| Time Step - Default | 0.05 |
| Time Step - Max | 0.1 |
| Error Tolerance - Absolute | 1.0E-9 |
| Error Tolerance - Relative | 1.0E-7 |
| Output Time Step | 1.0 |
| Use Symbolic Jacobian (T/F) | F |

#### 2.11.6.7. Widefield Bleach w fast  binding_long duration (FLIP)

| **Simulation Name: Widefield Bleach w fast binding_long duration (FLIP)** |
|---|
| **Simulation Description: cloned from 'Widefield Bleach w fast binding_long duration (FLIP)' owned by user temp<br>cloned from 'Widefield Bleach w fast binding_long duration (FLIP)' owned by user temp<br>cloned from 'Widefield Bleach w fast binding_1_1_1' owned by user temp<br>cloned from 'Widefield Bleach w fast binding_1_1' owned by user temp<br>cloned from 'Widefield Bleach w fast binding_1' owned by user temp<br>cloned from 'Widefield Bleach w binding_1' owned by user temp** |

| Overriden Parameters |
|---|

| Name | Actual Value | Default Value |
|---|---|---|
| kbleach_BleachFree | 10.0 | 1.0 |
| Kf_BindD | 100.0 | 1.0 |
| BleachRadius | 4.0 | 2.0 |
| kbleach_BleachBound | 10.0 | 1.0 |
| duration | 50.0 | 1.0 |
| Kr_BindF | 100.0 | 1.0 |
| Kr_BindD | 100.0 | 1.0 |
| Fluor_diffusionRate | "3.0", "25.0" | 10.0 |
| Dark_diffusionRate | Fluor_diffusionRate | 10.0 |
| sigmaaxial | 30.0 | 1.5 |
| Kf_BindF | 100.0 | 1.0 |

| Geometry Setting | |
|---|---|
| Geometry Size (um) | (74.24, 74.24, 26.0) |
| Mesh Size (elements) | (256, 256, 35) |

| Advanced Settings | |
|---|---|
| Solver Name | Fully-Implicit Finite Volume, Regular Grid (Variable Time Step) |
| Time Bounds - Starting | 0.0 |
| Time Bounds - Ending | 100.0 |
| Time Step - Min | 0.0 |
| Time Step - Default | 0.05 |
| Time Step - Max | 0.1 |
| Error Tolerance - Absolute | 1.0E-9 |

| | |
|---|---|
| Error Tolerance - Relative | 1.0E-7 |
| Output Time Step | 1.0 |
| Use Symbolic Jacobian (T/F) | F |

##### 2.11.6.8. Widefield Bleach w defalt binding_long duration (FLIP)

| **Simulation Name: Widefield Bleach w defalt binding_long duration (FLIP)** |
|---|
| **Simulation Description: cloned from 'Widefield Bleach w defalt binding_long duration (FLIP)' owned by user temp<br>cloned from 'Widefield Bleach w defalt binding_1_1_1' owned by user temp<br>cloned from 'Widefield Bleach w defalt binding_1_1' owned by user temp<br>cloned from 'Widefield Bleach w defalt binding_1' owned by user temp<br>cloned from 'Widefield Bleach w slower binding' owned by user temp<br>cloned from 'Widefield Bleach w binding_1' owned by user temp** |

| **Overriden Parameters** | | |
|---|---|---|
| **Name** | **Actual Value** | **Default Value** |
| BleachRadius | 4.0 | 2.0 |
| Fluor_diffusionRate | "1.0", "10.0" | 10.0 |
| kbleach_BleachFree | 10.0 | 1.0 |
| sigmaaxial | 30.0 | 1.5 |
| kbleach_BleachBound | 10.0 | 1.0 |
| duration | 50.0 | 1.0 |
| Dark_diffusionRate | Fluor_diffusionRate | 10.0 |

| **Geometry Setting** | |
|---|---|
| Geometry Size (um) | (74.24, 74.24, 26.0) |
| Mesh Size (elements) | (256, 256, 35) |

| **Advanced Settings** |
|---|

| | |
|---|---|
| Solver Name | Fully-Implicit Finite Volume, Regular Grid (Variable Time Step) |
| Time Bounds - Starting | 0.0 |
| Time Bounds - Ending | 100.0 |
| Time Step - Min | 0.0 |
| Time Step - Default | 0.05 |
| Time Step - Max | 0.1 |
| Error Tolerance - Absolute | 1.0E-9 |
| Error Tolerance - Relative | 1.0E-7 |
| Output Time Step | 1.0 |
| Use Symbolic Jacobian (T/F) | F |

##### 2.11.6.9. Widefield Bleach w slow binding_long duration (FLIP)

| **Simulation Name: Widefield Bleach w slow binding_long duration (FLIP)** |
|---|
| **Simulation Description: cloned from 'Widefield Bleach w slow binding_long duration (FLIP)' owned by user temp<br>cloned from 'Widefield Bleach w slow binding_long duration (FLIP)' owned by user temp<br>cloned from 'Widefield Bleach w slow binding_1_1_1' owned by user temp<br>cloned from 'Widefield Bleach w slow binding_1_1' owned by user temp<br>cloned from 'Widefield Bleach w slow binding_1' owned by user temp<br>cloned from 'Copy of Widefield Bleach w slow binding' owned by user temp<br>cloned from 'Widefield Bleach w slower binding' owned by user temp<br>cloned from 'Widefield Bleach w binding_1' owned by user temp** |

| **Overriden Parameters** | | |
|---|---|---|
| **Name** | **Actual Value** | **Default Value** |
| kbleach_BleachFree | 10.0 | 1.0 |
| Kf_BindD | 0.1 | 1.0 |
| BleachRadius | 4.0 | 2.0 |

| | | |
|---|---|---|
| kbleach_BleachBound | 10.0 | 1.0 |
| duration | 50.0 | 1.0 |
| Kr_BindF | 0.1 | 1.0 |
| Kr_BindD | 0.1 | 1.0 |
| Fluor_diffusionRate | "3.0", "25.0" | 10.0 |
| Dark_diffusionRate | Fluor_diffusionRate | 10.0 |
| sigmaaxial | 30.0 | 1.5 |
| Kf_BindF | 0.1 | 1.0 |

| **Geometry Setting** | |
|---|---|
| Geometry Size (um) | (74.24, 74.24, 26.0) |
| Mesh Size (elements) | (256, 256, 35) |

| **Advanced Settings** | |
|---|---|
| Solver Name | Fully-Implicit Finite Volume, Regular Grid (Variable Time Step) |
| Time Bounds - Starting | 0.0 |
| Time Bounds - Ending | 100.0 |
| Time Step - Min | 0.0 |
| Time Step - Default | 0.05 |
| Time Step - Max | 0.1 |
| Error Tolerance - Absolute | 1.0E-9 |
| Error Tolerance - Relative | 1.0E-7 |
| Output Time Step | 1.0 |
| Use Symbolic Jacobian (T/F) | F |

#### 2.12. Application: Convolved Image-based center circular bleach

| **Application Name: Convolved Image-based center circular bleach** |
|---|
| **Application Description: (copied from Image-based center circular bleach) (copied from Image-based center bleach) (copied from Spherical_Cell_Gaussian_Bleach)** |

##### 2.12.1. Structure Mapping For Convolved Image-based center circular bleach

| **Structure Mapping** | | | | |
|---|---|---|---|---|
| **Structure** | **Subdomain** | **Resolved (T/F)** | **Surf/Vol** | **VolFract** |
| EC | ec | F | | |
| NUC | Nucleus | F | | |
| CYT | cytosol | F | | |

##### 2.12.2. Reaction Mapping For Convolved Image-based center circular bleach

| **Reaction Mapping** | | | |
|---|---|---|---|
| **Name** | **Type** | **Enabled (T/F)** | **Fast (T/F)** |
| BindD | Reaction | T | F |
| BindF | Reaction | T | F |
| BleachFree | Reaction | T | F |
| BleachBound | Reaction | T | F |

| **Initial Conditions** | | | | |
|---|---|---|---|---|
| **Species** | **Structure** | **Initial Conc.** | **Diffusion Const.** | **Fixed (T/F)** |
| s0 | CYT | 2.701562118716424 M | 10.0 $m^2.s^1$ | F |
| s1 | CYT | 0.0 M | 10.0 $m^2.s^1$ | F |
| s2 | CYT | 2.701562118716425 M | 0.0 $m^2.s^1$ | F |

| Initial Conditions | | | | |
|---|---|---|---|---|
| **Species** | **Structure** | **Initial Conc.** | **Diffusion Const.** | **Fixed (T/F)** |
| s3 | CYT | 7.2984378812835775 M | 0.0 $m^{2}.s^{1}$ | F |
| s4 | CYT | (exp( - (((z - 13.0) ^ 2.0) / (2.0 * (sigmaaxial ^ 2.0)))) * ((((x - 23.0) ^ 2.0) + ((y - 39.0) ^ 2.0)) < 4.0)) M | 0.0 $m^{2}.s^{1}$ | F |
| s5 | CYT | 0.0 M | 0.0 $m^{2}.s^{1}$ | F |

#### 2.12.3. Membrane Mapping For Convolved Image-based center circular bleach

| Electrical Mapping - Membrane Potential | | | |
|---|---|---|---|
| **Membrane** | **Calculate V (T/F)** | **V initial** | **Specific Capacitance** |
| PM | F | 0.0 mV | 1.0 $pF.m^{2}$ |
| NM | F | 0.0 mV | 1.0 $pF.m^{2}$ |

| |
|---|
| Temperature: 300.0 K |

#### 2.12.4. Geometry: Site visit _Application0_20111127_695607844

| | |
|---|---|
| Size | (74.24, 74.24, 26.0) |
| Origin | (0.0, 0.0, 0.0) |

#### 2.12.5. Math Description: Copy of Image-based center circular bleach_generated

##### 2.12.5.1. Constants

| Constant Name | Expression |
|---|---|
| _F_ | 96485.3321 |
| _F_nmol_ | 9.64853321E-5 |
| _K_GHK_ | 1.0E-9 |

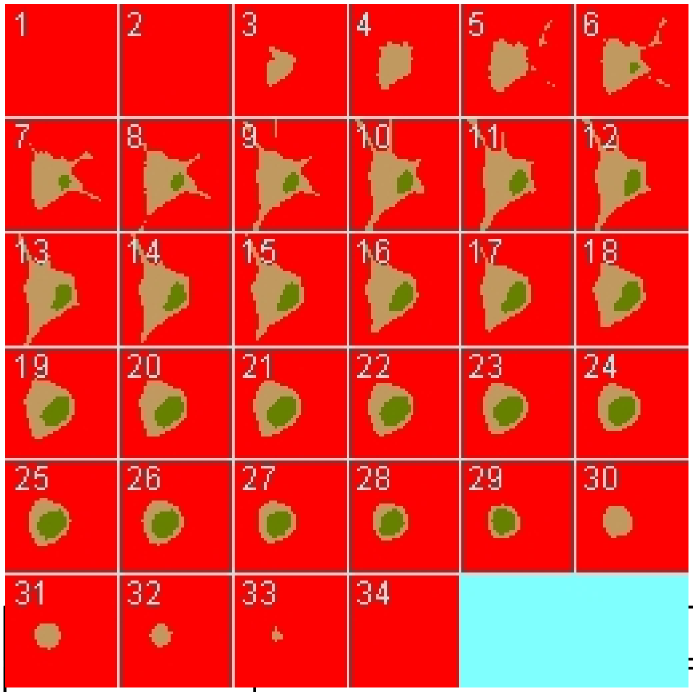

| Constant Name | Expression |
|---|---|
| _N_pmol_ | 6.02214179E11 |
| _PI_ | 3.141592653589793 |
| _R_ | 8314.46261815 |
| _T_ | 300.0 |
| AreaPerUnitArea_NM | 1.0 |
| AreaPerUnitArea_PM | 1.0 |
| Binder_init_uM | 2.701562118716425 |
| BleachRadius | 2.0 |
| Dark_diffusionRate | 10.0 |
| Dark_init_uM | 0.0 |
| DarkB_init_uM | 0.0 |
| duration | 1.0 |
| Fluor_diffusionRate | 10.0 |
| Fluor_init_uM | 2.701562118716424 |
| FluorB_init_uM | 7.2984378812835775 |
| K_millivolts_per_volt | 1000.0 |

| Constant Name | Expression |
| --- | --- |
| kbleach_BleachBound | 1.0 |
| kbleach_BleachFree | 1.0 |
| Kf_BindD | 1.0 |
| Kf_BindF | 1.0 |
| KMOLE | 0.001660538783162726 |
| Kr_BindD | 1.0 |
| Kr_BindF | 1.0 |
| Kr_BleachBound | 0.0 |
| Kr_BleachFree | 0.0 |
| sigmaaxial | 1.5 |
| sigmalateral | 0.5 |
| start | 1.0 |
| UnitFactor_molecules_um_neg_3_uM_neg_1 | (1.0 * pow(KMOLE,-1.0)) |
| Voltage_NM | 0.0 |
| Voltage_PM | 0.0 |
| VolumePerUnitVolume_CYT | 1.0 |
| VolumePerUnitVolume_EC | 1.0 |
| VolumePerUnitVolume_NUC | 1.0 |

#### 2.12.5.2. Functions

| Function Name | Expression |
| --- | --- |
| J_BindD | (((Kf_BindD * Dark) * Binder) - (Kr_BindD * DarkB)) |
| J_BindF | (((Kf_BindF * Fluor) * Binder) - (Kr_BindF * FluorB)) |
| J_BleachBound | ((Kf_BleachBound * FluorB) - (Kr_BleachBound * DarkB)) |
| J_BleachFree | ((Kf_BleachFree * Fluor) - (Kr_BleachFree * Dark)) |

| Function Name | Expression |
|---|---|
| Kf_BleachBound | ((kbleach_BleachBound * Laser) * (t > start) * (t < (start + duration))) |
| Kf_BleachFree | ((kbleach_BleachFree * Laser) * (t > start) * (t < (start + duration))) |
| Laser_init_uM | (exp( - (((z - 13.0) ^ 2.0) / (2.0 * (sigmaaxial ^ 2.0)))) * ((((x - 23.0) ^ 2.0) + ((y - 39.0) ^ 2.0)) < 4.0)) |
| Size_CYT | (VolumePerUnitVolume_CYT * vcRegionVolume('cytosol')) |
| Size_EC | (VolumePerUnitVolume_EC * vcRegionVolume('ec')) |
| Size_NM | (AreaPerUnitArea_NM * vcRegionArea('Nucleus_cytosol_membrane')) |
| Size_NUC | (VolumePerUnitVolume_NUC * vcRegionVolume('Nucleus')) |
| Size_PM | (AreaPerUnitArea_PM * vcRegionArea('cytosol_ec_membrane')) |
| sobj_cytosol1_ec0_size | vcRegionArea('cytosol_ec_membrane') |
| sobj_Nucleus2_cytosol1_size | vcRegionArea('Nucleus_cytosol_membrane') |
| vobj_cytosol1_size | vcRegionVolume('cytosol') |
| vobj_ec0_size | vcRegionVolume('ec') |
| vobj_Nucleus2_size | vcRegionVolume('Nucleus') |

#### 2.12.5.3. Volume Domains

### 2.12.5.3.1. ec

##### 2.12.5.3.2. cytosol

| PdeEquation Fluor | |
|---|---|
| Rate | ( - J_BindF - J_BleachFree) |
| Diffusion | Fluor_diffusionRate |
| Initial | Fluor_init_uM |

| PdeEquation Dark | |
|---|---|
| Rate | ( - J_BindD + J_BleachFree) |

| PdeEquation Dark | |
|---|---|
| Diffusion | Dark_diffusionRate |
| Initial | Dark_init_uM |

| OdeEquation Binder | |
|---|---|
| Rate | ( - J_BindD - J_BindF) |
| Initial | Binder_init_uM |

| OdeEquation FluorB | |
|---|---|
| Rate | (J_BindF - J_BleachBound) |
| Initial | FluorB_init_uM |

| OdeEquation Laser | |
|---|---|
| Rate | 0.0 |
| Initial | Laser_init_uM |

| OdeEquation DarkB | |
|---|---|
| Rate | (J_BindD + J_BleachBound) |
| Initial | DarkB_init_uM |

##### 2.12.5.3.3. Nucleus

#### 2.12.5.4. Membrane Domains

##### 2.12.5.4.1. cytosol_ec_membrane

| JumpCondition Fluor | |
|---|---|
| InFlux | 0.0 |
| OutFlux | 0.0 |

| JumpCondition Dark |
|---|

| JumpCondition Dark | |
|---|---|
| InFlux | 0.0 |
| OutFlux | 0.0 |

##### 2.12.5.4.2. Nucleus_cytosol_membrane

| JumpCondition Fluor | |
|---|---|
| InFlux | 0.0 |
| OutFlux | 0.0 |

| JumpCondition Dark | |
|---|---|
| InFlux | 0.0 |
| OutFlux | 0.0 |

#### 2.12.6. Simulation(s)

##### 2.12.6.1. 2Photon Bleach no binding_1

| |
|---|
| **Simulation Name: 2Photon Bleach no binding_1** |
| **Simulation Description: cloned from '2Photon Bleach no binding_1' owned by user temp<br>cloned from '2Photon Bleach_1' owned by user temp** |

| Overriden Parameters | | |
|---|---|---|
| **Name** | **Actual Value** | **Default Value** |
| Fluor_diffusionRate | "1.0", "10.0" | 10.0 |
| Binder_init_uM | 0.0 | 2.701562118716425 |
| kbleach_BleachFree | 10.0 | 1.0 |
| duration | "0.3", "3.0" | 1.0 |
| kbleach_BleachBound | 10.0 | 1.0 |
| Dark_diffusionRate | Fluor_diffusionRate | 10.0 |

| | | |
|---|---|---|
| Fluor_init_uM | 10.0 | 2.701562118716424 |
| FluorB_init_uM | 0.0 | 7.2984378812835775 |

| Geometry Setting | |
|---|---|
| Geometry Size (um) | (74.24, 74.24, 26.0) |
| Mesh Size (elements) | (256, 256, 90) |

| Advanced Settings | |
|---|---|
| Solver Name | Fully-Implicit Finite Volume, Regular Grid (Variable Time Step) |
| Time Bounds - Starting | 0.0 |
| Time Bounds - Ending | 20.0 |
| Time Step - Min | 0.0 |
| Time Step - Default | 0.05 |
| Time Step - Max | 0.1 |
| Error Tolerance - Absolute | 1.0E-9 |
| Error Tolerance - Relative | 1.0E-7 |
| Output Time Step | 0.5 |
| Use Symbolic Jacobian (T/F) | F |

#### 2.12.6.2. Widefield Bleach no binding_1

| Simulation Name: Widefield Bleach no binding_1 |
|---|
| Simulation Description: cloned from 'Widefield Bleach no binding_1' owned by user temp cloned from 'Widefield Bleach_1' owned by user temp |

| Overriden Parameters | | |
|---|---|---|
| **Name** | **Actual Value** | **Default Value** |

| | | |
|---|---|---|
| kbleach_BleachFree | 10.0 | 1.0 |
| kbleach_BleachBound | 10.0 | 1.0 |
| duration | "0.3", "3.0" | 1.0 |
| Fluor_init_uM | 10.0 | 2.701562118716424 |
| Fluor_diffusionRate | "1.0", "10.0" | 10.0 |
| Dark_diffusionRate | Fluor_diffusionRate | 10.0 |
| sigmaaxial | 30.0 | 1.5 |
| Binder_init_uM | 0.0 | 2.701562118716425 |
| FluorB_init_uM | 0.0 | 7.2984378812835775 |

| **Geometry Setting** | |
|---|---|
| Geometry Size (um) | (74.24, 74.24, 26.0) |
| Mesh Size (elements) | (256, 256, 90) |

| **Advanced Settings** | |
|---|---|
| Solver Name | Fully-Implicit Finite Volume, Regular Grid (Variable Time Step) |
| Time Bounds - Starting | 0.0 |
| Time Bounds - Ending | 20.0 |
| Time Step - Min | 0.0 |
| Time Step - Default | 0.05 |
| Time Step - Max | 0.1 |
| Error Tolerance - Absolute | 1.0E-9 |
| Error Tolerance - Relative | 1.0E-7 |
| Output Time Step | 0.5 |
| Use Symbolic Jacobian (T/F) | F |

#### 2.12.6.3. 2Photon Bleach w fast binding_2

| **Simulation Name: 2Photon Bleach w fast binding_2** |
|---|
| **Simulation Description: cloned from '2Photon Bleach w fast binding_2' owned by user temp cloned from '2Photon Bleach w binding_1' owned by user temp** |

| **Overriden Parameters** | | |
|---|---|---|
| **Name** | **Actual Value** | **Default Value** |
| Fluor_diffusionRate | "1.0", "10.0" | 10.0 |
| Kf_BindF | 100.0 | 1.0 |
| Kr_BindF | 100.0 | 1.0 |
| Kf_BindD | 100.0 | 1.0 |
| Kr_BindD | 100.0 | 1.0 |
| kbleach_BleachFree | 10.0 | 1.0 |
| kbleach_BleachBound | 10.0 | 1.0 |
| Dark_diffusionRate | Fluor_diffusionRate | 10.0 |

| **Geometry Setting** | |
|---|---|
| Geometry Size (um) | (74.24, 74.24, 26.0) |
| Mesh Size (elements) | (256, 256, 90) |

| **Advanced Settings** | |
|---|---|
| Solver Name | Fully-Implicit Finite Volume, Regular Grid (Variable Time Step) |
| Time Bounds - Starting | 0.0 |
| Time Bounds - Ending | 20.0 |
| Time Step - Min | 0.0 |

| | |
|---|---|
| Time Step - Default | 0.05 |
| Time Step - Max | 0.1 |
| Error Tolerance - Absolute | 1.0E-9 |
| Error Tolerance - Relative | 1.0E-7 |
| Output Time Step | 0.5 |
| Use Symbolic Jacobian (T/F) | F |

#### 2.12.6.4. Widefield Bleach w fast binding_2

| |
|---|
| **Simulation Name: Widefield Bleach w fast binding_2** |
| **Simulation Description: cloned from 'Widefield Bleach w fast binding_2' owned by user temp cloned from 'Widefield Bleach w binding_1' owned by user temp** |

| **Overriden Parameters** | | |
|---|---|---|
| **Name** | **Actual Value** | **Default Value** |
| kbleach_BleachFree | 10.0 | 1.0 |
| Kf_BindD | 100.0 | 1.0 |
| kbleach_BleachBound | 10.0 | 1.0 |
| duration | "0.3", "3.0" | 1.0 |
| Kr_BindF | 100.0 | 1.0 |
| Kr_BindD | 100.0 | 1.0 |
| Fluor_diffusionRate | "1.0", "10.0" | 10.0 |
| Dark_diffusionRate | Fluor_diffusionRate | 10.0 |
| sigmaaxial | 30.0 | 1.5 |
| Kf_BindF | 100.0 | 1.0 |

| **Geometry Setting** |
|---|

| | |
|---|---|
| Geometry Size (um) | (74.24, 74.24, 26.0) |
| Mesh Size (elements) | (256, 256, 90) |

| Advanced Settings | |
|---|---|
| Solver Name | Fully-Implicit Finite Volume, Regular Grid (Variable Time Step) |
| Time Bounds - Starting | 0.0 |
| Time Bounds - Ending | 20.0 |
| Time Step - Min | 0.0 |
| Time Step - Default | 0.05 |
| Time Step - Max | 0.1 |
| Error Tolerance - Absolute | 1.0E-9 |
| Error Tolerance - Relative | 1.0E-7 |
| Output Time Step | 0.5 |
| Use Symbolic Jacobian (T/F) | F |

#### 2.12.6.5. 2Photon Bleach w default binding_2

| **Simulation Name: 2Photon Bleach w default binding_2** |
|---|
| **Simulation Description: cloned from '2Photon Bleach w default binding_2' owned by user temp<br>cloned from '2Photon Bleach w slower binding' owned by user temp<br>cloned from '2Photon Bleach w binding_1' owned by user temp** |

| **Overriden Parameters** | | |
|---|---|---|
| **Name** | **Actual Value** | **Default Value** |
| Fluor_diffusionRate | "1.0", "10.0" | 10.0 |
| kbleach_BleachFree | 10.0 | 1.0 |
| kbleach_BleachBound | 10.0 | 1.0 |

| | | |
|---|---|---|
| Dark_diffusionRate | Fluor_diffusionRate | 10.0 |

| Geometry Setting | |
|---|---|
| Geometry Size (um) | (74.24, 74.24, 26.0) |
| Mesh Size (elements) | (256, 256, 90) |

| Advanced Settings | |
|---|---|
| Solver Name | Fully-Implicit Finite Volume, Regular Grid (Variable Time Step) |
| Time Bounds - Starting | 0.0 |
| Time Bounds - Ending | 20.0 |
| Time Step - Min | 0.0 |
| Time Step - Default | 0.05 |
| Time Step - Max | 0.1 |
| Error Tolerance - Absolute | 1.0E-9 |
| Error Tolerance - Relative | 1.0E-7 |
| Output Time Step | 0.5 |
| Use Symbolic Jacobian (T/F) | F |

#### 2.12.6.6. Widefield Bleach w defalt binding_2

| Simulation Name: Widefield Bleach w defalt binding_2 |
|---|
| Simulation Description: cloned from 'Widefield Bleach w defalt binding_2' owned by user temp<br>cloned from 'Widefield Bleach w slower binding' owned by user temp<br>cloned from 'Widefield Bleach w binding_1' owned by user temp |

| Overriden Parameters | | |
|---|---|---|
| **Name** | **Actual Value** | **Default Value** |

| | | |
|---|---|---|
| Fluor_diffusionRate | "1.0", "10.0" | 10.0 |
| sigmaaxial | 30.0 | 1.5 |
| kbleach_BleachFree | 10.0 | 1.0 |
| duration | "0.3", "3.0" | 1.0 |
| kbleach_BleachBound | 10.0 | 1.0 |
| Dark_diffusionRate | Fluor_diffusionRate | 10.0 |

| **Geometry Setting** | |
|---|---|
| Geometry Size (um) | (74.24, 74.24, 26.0) |
| Mesh Size (elements) | (256, 256, 90) |

| **Advanced Settings** | |
|---|---|
| Solver Name | Fully-Implicit Finite Volume, Regular Grid (Variable Time Step) |
| Time Bounds - Starting | 0.0 |
| Time Bounds - Ending | 20.0 |
| Time Step - Min | 0.0 |
| Time Step - Default | 0.05 |
| Time Step - Max | 0.1 |
| Error Tolerance - Absolute | 1.0E-9 |
| Error Tolerance - Relative | 1.0E-7 |
| Output Time Step | 0.5 |
| Use Symbolic Jacobian (T/F) | F |

#### 2.12.6.7. 2Photon Bleach w slow binding_2

| **Simulation Name: 2Photon Bleach w slow binding_2** |
|---|

**Simulation  Description: cloned from '2Photon Bleach w slow binding_2' owned by user temp**
**cloned from 'Copy of 2Photon Bleach w slow binding' owned by user temp**
**cloned from '2Photon Bleach w slower binding' owned by user temp**
**cloned from '2Photon Bleach w binding_1' owned by user temp**

| Overriden Parameters | | |
|---|---|---|
| **Name** | **Actual Value** | **Default Value** |
| Fluor_diffusionRate | "1.0", "10.0" | 10.0 |
| Kf_BindF | 0.1 | 1.0 |
| Kr_BindF | 0.1 | 1.0 |
| Kf_BindD | 0.1 | 1.0 |
| Kr_BindD | 0.1 | 1.0 |
| kbleach_BleachFree | 10.0 | 1.0 |
| kbleach_BleachBound | 10.0 | 1.0 |
| Dark_diffusionRate | Fluor_diffusionRate | 10.0 |

| Geometry Setting | |
|---|---|
| Geometry Size (um) | (74.24, 74.24, 26.0) |
| Mesh Size (elements) | (256, 256, 90) |

| Advanced Settings | |
|---|---|
| Solver Name | Fully-Implicit Finite Volume, Regular Grid (Variable Time Step) |
| Time Bounds - Starting | 0.0 |
| Time Bounds - Ending | 20.0 |
| Time Step - Min | 0.0 |
| Time Step - Default | 0.05 |

| | |
|---|---|
| Time Step - Max | 0.1 |
| Error Tolerance - Absolute | 1.0E-9 |
| Error Tolerance - Relative | 1.0E-7 |
| Output Time Step | 0.5 |
| Use Symbolic Jacobian (T/F) | F |

#### 2.12.6.8. Widefield Bleach w slow binding_2

| **Simulation Name: Widefield Bleach w slow binding_2** |
|---|
| **Simulation Description: cloned from 'Widefield Bleach w slow binding_2' owned by user temp<br>cloned from 'Copy of Widefield Bleach w slow binding' owned by user temp<br>cloned from 'Widefield Bleach w slower binding' owned by user temp<br>cloned from 'Widefield Bleach w binding_1' owned by user temp** |

| **Overriden Parameters** | | |
|---|---|---|
| **Name** | **Actual Value** | **Default Value** |
| kbleach_BleachFree | 10.0 | 1.0 |
| Kf_BindD | 0.1 | 1.0 |
| kbleach_BleachBound | 10.0 | 1.0 |
| duration | "0.3", "3.0" | 1.0 |
| Kr_BindF | 0.1 | 1.0 |
| Kr_BindD | 0.1 | 1.0 |
| Fluor_diffusionRate | "1.0", "10.0" | 10.0 |
| Dark_diffusionRate | Fluor_diffusionRate | 10.0 |
| sigmaaxial | 30.0 | 1.5 |
| Kf_BindF | 0.1 | 1.0 |

| **Geometry Setting** |
|---|

| | |
|---|---|
| Geometry Size (um) | (74.24, 74.24, 26.0) |
| Mesh Size (elements) | (256, 256, 90) |

| Advanced Settings | |
|---|---|
| Solver Name | Fully-Implicit Finite Volume, Regular Grid (Variable Time Step) |
| Time Bounds - Starting | 0.0 |
| Time Bounds - Ending | 20.0 |
| Time Step - Min | 0.0 |
| Time Step - Default | 0.05 |
| Time Step - Max | 0.1 |
| Error Tolerance - Absolute | 1.0E-9 |
| Error Tolerance - Relative | 1.0E-7 |
| Output Time Step | 0.5 |
| Use Symbolic Jacobian (T/F) | F |

#### 2.13. Application: ProjectZ Convolved Image-based center circular bleach

| **Application Name: ProjectZ Convolved Image-based center circular bleach** |
|---|
| **Application Description: (copied from Convolved Image-based center circular bleach) (copied from Image-based center circular bleach) (copied from Image-based center bleach) (copied from Spherical_Cell_Gaussian_Bleach)** |

##### 2.13.1. Structure Mapping For ProjectZ Convolved Image-based center circular bleach

| **Structure Mapping** | | | | |
|---|---|---|---|---|
| **Structure** | **Subdomain** | **Resolved (T/F)** | **Surf/Vol** | **VolFract** |
| EC | ec | F | | |
| NUC | Nucleus | F | | |

| Structure Mapping | | | | |
|---|---|---|---|---|
| **Structure** | **Subdomain** | **Resolved (T/F)** | **Surf/Vol** | **VolFract** |
| CYT | cytosol | F | | |

#### 2.13.2. Reaction Mapping For ProjectZ Convolved  Image-based center circular bleach

| Reaction Mapping | | | |
|---|---|---|---|
| **Name** | **Type** | **Enabled (T/F)** | **Fast (T/F)** |
| BindD | Reaction | T | F |
| BindF | Reaction | T | F |
| BleachFree | Reaction | T | F |
| BleachBound | Reaction | T | F |

| Initial Conditions | | | | |
|---|---|---|---|---|
| **Species** | **Structure** | **Initial Conc.** | **Diffusion Const.** | **Fixed (T/F)** |
| s0 | CYT | 2.701562118716424 M | 10.0 $m^2.s^1$ | F |
| s1 | CYT | 0.0 M | 10.0 $m^2.s^1$ | F |
| s2 | CYT | 2.701562118716425 M | 0.0 $m^2.s^1$ | F |
| s3 | CYT | 7.2984378812835775 M | 0.0 $m^2.s^1$ | F |
| s4 | CYT | (exp( - (((z - 13.0) ^ 2.0) / (2.0 * (sigmaaxial ^ 2.0)))) * ((((x - 23.0) ^ 2.0) + ((y - 39.0) ^ 2.0)) < 4.0)) M | 0.0 $m^2.s^1$ | F |
| s5 | CYT | 0.0 M | 0.0 $m^2.s^1$ | F |

#### 2.13.3. Membrane Mapping For ProjectZ Convolved  Image-based center circular bleach

| Electrical Mapping - Membrane Potential |
|---|

| Electrical Mapping - Membrane Potential | | | |
|---|---|---|---|
| **Membrane** | **Calculate V (T/F)** | **V initial** | **Specific Capacitance** |
| PM | F | 0.0 mV | 1.0 pF.m² |
| NM | F | 0.0 mV | 1.0 pF.m² |

| |
|---|
| Temperature: 300.0 K |

#### 2.13.4. Geometry: Site visit _Application0_20111127_695607844

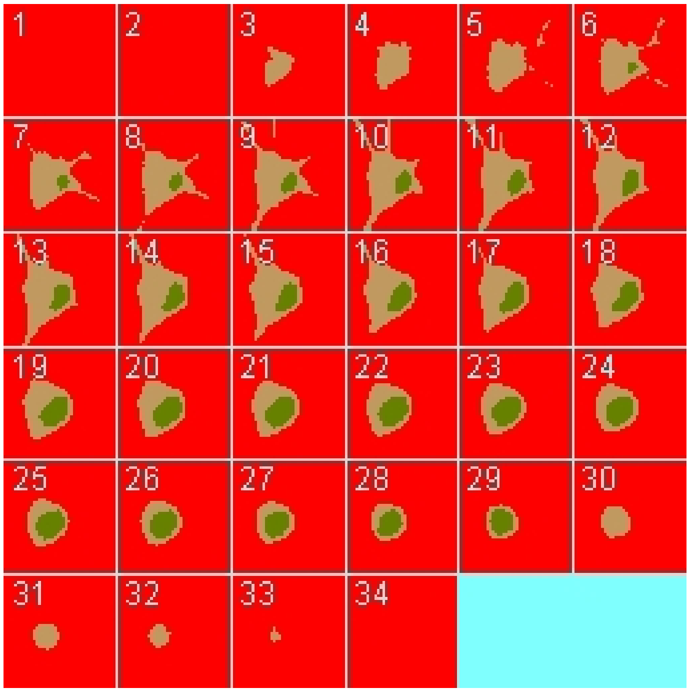

| | |
|---|---|
| Size | (74.24, 74.24, 26.0) |
| Origin | (0.0, 0.0, 0.0) |

#### 2.13.5. Math Description: Copy of Convolved  Image-based center circular bleach_generated

#### 2.13.5.1. Constants

| Constant Name | Expression |
| --- | --- |
| _F_ | 96485.3321 |
| _F_nmol_ | 9.64853321E-5 |
| _K_GHK_ | 1.0E-9 |
| _N_pmol_ | 6.02214179E11 |
| _PI_ | 3.141592653589793 |
| _R_ | 8314.46261815 |
| _T_ | 300.0 |
| AreaPerUnitArea_NM | 1.0 |
| AreaPerUnitArea_PM | 1.0 |
| Binder_init_uM | 2.701562118716425 |
| BleachRadius | 2.0 |
| Dark_diffusionRate | 10.0 |
| Dark_init_uM | 0.0 |
| DarkB_init_uM | 0.0 |
| duration | 1.0 |
| Fluor_diffusionRate | 10.0 |
| Fluor_init_uM | 2.701562118716424 |
| FluorB_init_uM | 7.2984378812835775 |
| K_millivolts_per_volt | 1000.0 |
| kbleach_BleachBound | 1.0 |
| kbleach_BleachFree | 1.0 |
| Kf_BindD | 1.0 |
| Kf_BindF | 1.0 |
| KMOLE | 0.001660538783162726 |
| Kr_BindD | 1.0 |
| Kr_BindF | 1.0 |

| Constant Name | Expression |
|---|---|
| Kr_BleachBound | 0.0 |
| Kr_BleachFree | 0.0 |
| sigmaaxial | 1.5 |
| sigmalateral | 0.5 |
| start | 1.0 |
| UnitFactor_molecules_um_neg_3_uM_neg_1 | (1.0 * pow(KMOLE,-1.0)) |
| Voltage_NM | 0.0 |
| Voltage_PM | 0.0 |
| VolumePerUnitVolume_CYT | 1.0 |
| VolumePerUnitVolume_EC | 1.0 |
| VolumePerUnitVolume_NUC | 1.0 |

#### 2.13.5.2. Functions

| Function Name | Expression |
|---|---|
| J_BindD | (((Kf_BindD * Dark) * Binder) - (Kr_BindD * DarkB)) |
| J_BindF | (((Kf_BindF * Fluor) * Binder) - (Kr_BindF * FluorB)) |
| J_BleachBound | ((Kf_BleachBound * FluorB) - (Kr_BleachBound * DarkB)) |
| J_BleachFree | ((Kf_BleachFree * Fluor) - (Kr_BleachFree * Dark)) |
| Kf_BleachBound | ((kbleach_BleachBound * Laser) * (t > start) * (t < (start + duration))) |
| Kf_BleachFree | ((kbleach_BleachFree * Laser) * (t > start) * (t < (start + duration))) |
| Laser_init_uM | (exp( - (((z - 13.0) ^ 2.0) / (2.0 * (sigmaaxial ^ 2.0)))) * ((((x - 23.0) ^ 2.0) + ((y - 39.0) ^ 2.0)) < 4.0)) |
| Size_CYT | (VolumePerUnitVolume_CYT * vcRegionVolume('cytosol')) |
| Size_EC | (VolumePerUnitVolume_EC * vcRegionVolume('ec')) |
| Size_NM | (AreaPerUnitArea_NM * vcRegionArea('Nucleus_cytosol_membrane')) |

| Function Name | Expression |
|---|---|
| Size_NUC | (VolumePerUnitVolume_NUC * vcRegionVolume('Nucleus')) |
| Size_PM | (AreaPerUnitArea_PM * vcRegionArea('cytosol_ec_membrane')) |
| sobj_cytosol1_ec0_size | vcRegionArea('cytosol_ec_membrane') |
| sobj_Nucleus2_cytosol1_size | vcRegionArea('Nucleus_cytosol_membrane') |
| vobj_cytosol1_size | vcRegionVolume('cytosol') |
| vobj_ec0_size | vcRegionVolume('ec') |
| vobj_Nucleus2_size | vcRegionVolume('Nucleus') |

##### 2.13.5.3. Volume Domains

#### 2.13.5.3.1. ec

###### 2.13.5.3.2. cytosol

| **PdeEquation Fluor** | |
|---|---|
| Rate | ( - J_BindF - J_BleachFree) |
| Diffusion | Fluor_diffusionRate |
| Initial | Fluor_init_uM |

| **PdeEquation Dark** | |
|---|---|
| Rate | ( - J_BindD + J_BleachFree) |
| Diffusion | Dark_diffusionRate |
| Initial | Dark_init_uM |

| **OdeEquation Binder** | |
|---|---|
| Rate | ( - J_BindD - J_BindF) |
| Initial | Binder_init_uM |

| OdeEquation FluorB | |
|---|---|
| Rate | (J_BindF - J_BleachBound) |
| Initial | FluorB_init_uM |

| OdeEquation Laser | |
|---|---|
| Rate | 0.0 |
| Initial | Laser_init_uM |

| OdeEquation DarkB | |
|---|---|
| Rate | (J_BindD + J_BleachBound) |
| Initial | DarkB_init_uM |

#### 2.13.5.3.3. Nucleus

#### 2.13.5.4. Membrane Domains

#### 2.13.5.4.1. cytosol_ec_membrane

| JumpCondition Fluor | |
|---|---|
| InFlux | 0.0 |
| OutFlux | 0.0 |

| JumpCondition Dark | |
|---|---|
| InFlux | 0.0 |
| OutFlux | 0.0 |

#### 2.13.5.4.2. Nucleus_cytosol_membrane

| JumpCondition Fluor | |
|---|---|
| InFlux | 0.0 |
| OutFlux | 0.0 |

| JumpCondition Dark | |
|---|---|
| InFlux | 0.0 |
| OutFlux | 0.0 |

#### 2.13.6. Simulation(s)

##### 2.13.6.1. 2Photon Bleach no binding_1_1

| **Simulation Name: 2Photon Bleach no binding_1_1** |
|---|
| **Simulation Description: cloned from '2Photon Bleach no binding_1_1' owned by user temp<br>cloned from '2Photon Bleach no binding_1' owned by user temp<br>cloned from '2Photon Bleach_1' owned by user temp** |

| Overriden Parameters | | |
|---|---|---|
| **Name** | **Actual Value** | **Default Value** |
| Fluor_diffusionRate | "1.0", "10.0" | 10.0 |
| Binder_init_uM | 0.0 | 2.701562118716425 |
| kbleach_BleachFree | 10.0 | 1.0 |
| duration | "0.3", "3.0" | 1.0 |
| kbleach_BleachBound | 10.0 | 1.0 |
| Dark_diffusionRate | Fluor_diffusionRate | 10.0 |
| Fluor_init_uM | 10.0 | 2.701562118716424 |
| FluorB_init_uM | 0.0 | 7.2984378812835775 |

| Geometry Setting | |
|---|---|
| Geometry Size (um) | (74.24, 74.24, 26.0) |
| Mesh Size (elements) | (256, 256, 90) |

| Advanced Settings | |
|---|---|
| Solver Name | Fully-Implicit Finite Volume, Regular Grid (Variable Time Step) |
| Time Bounds - Starting | 0.0 |
| Time Bounds - Ending | 20.0 |
| Time Step - Min | 0.0 |
| Time Step - Default | 0.05 |
| Time Step - Max | 0.1 |
| Error Tolerance - Absolute | 1.0E-9 |
| Error Tolerance - Relative | 1.0E-7 |
| Output Time Step | 0.5 |
| Use Symbolic Jacobian (T/F) | F |

#### 2.13.6.2. Widefield Bleach no binding_1_1

| Simulation Name: Widefield Bleach no binding_1_1 |
|---|
| Simulation Description: cloned from 'Widefield Bleach no binding_1_1' owned by user temp<br>cloned from 'Widefield Bleach no binding_1' owned by user temp<br>cloned from 'Widefield Bleach_1' owned by user temp |

| Overriden Parameters | | |
|---|---|---|
| **Name** | **Actual Value** | **Default Value** |
| kbleach_BleachFree | 10.0 | 1.0 |
| kbleach_BleachBound | 10.0 | 1.0 |
| duration | "0.3", "3.0" | 1.0 |
| Fluor_init_uM | 10.0 | 2.701562118716424 |
| Fluor_diffusionRate | "1.0", "10.0" | 10.0 |

| | | |
|---|---|---|
| Dark_diffusionRate | Fluor_diffusionRate | 10.0 |
| sigmaaxial | 30.0 | 1.5 |
| Binder_init_uM | 0.0 | 2.701562118716425 |
| FluorB_init_uM | 0.0 | 7.2984378812835775 |

| Geometry Setting | |
|---|---|
| Geometry Size (um) | (74.24, 74.24, 26.0) |
| Mesh Size (elements) | (256, 256, 90) |

| Advanced Settings | |
|---|---|
| Solver Name | Fully-Implicit Finite Volume, Regular Grid (Variable Time Step) |
| Time Bounds - Starting | 0.0 |
| Time Bounds - Ending | 20.0 |
| Time Step - Min | 0.0 |
| Time Step - Default | 0.05 |
| Time Step - Max | 0.1 |
| Error Tolerance - Absolute | 1.0E-9 |
| Error Tolerance - Relative | 1.0E-7 |
| Output Time Step | 0.5 |
| Use Symbolic Jacobian (T/F) | F |

##### 2.13.6.3. 2Photon Bleach w fast binding_2_1

| **Simulation Name: 2Photon Bleach w fast binding_2_1** |
|---|
| **Simulation Description: cloned from '2Photon Bleach w fast binding_2_1' owned by user temp<br>cloned from '2Photon Bleach w fast binding_2' owned by user temp<br>cloned from '2Photon Bleach w binding_1' owned by user temp** |

| Overriden Parameters | | |
|---|---|---|
| **Name** | **Actual Value** | **Default Value** |
| Fluor_diffusionRate | "1.0", "10.0" | 10.0 |
| Kf_BindF | 100.0 | 1.0 |
| Kr_BindF | 100.0 | 1.0 |
| Kf_BindD | 100.0 | 1.0 |
| Kr_BindD | 100.0 | 1.0 |
| kbleach_BleachFree | 10.0 | 1.0 |
| kbleach_BleachBound | 10.0 | 1.0 |
| Dark_diffusionRate | Fluor_diffusionRate | 10.0 |

| Geometry Setting | |
|---|---|
| Geometry Size (um) | (74.24, 74.24, 26.0) |
| Mesh Size (elements) | (256, 256, 90) |

| Advanced Settings | |
|---|---|
| Solver Name | Fully-Implicit Finite Volume, Regular Grid (Variable Time Step) |
| Time Bounds - Starting | 0.0 |
| Time Bounds - Ending | 20.0 |
| Time Step - Min | 0.0 |
| Time Step - Default | 0.05 |
| Time Step - Max | 0.1 |
| Error Tolerance - Absolute | 1.0E-9 |
| Error Tolerance - Relative | 1.0E-7 |

| | |
|---|---|
| Output Time Step | 0.5 |
| Use Symbolic Jacobian (T/F) | F |

##### 2.13.6.4. Widefield Bleach w fast binding_2_1

| **Simulation Name: Widefield Bleach w fast binding_2_1** |
|---|
| **Simulation Description: cloned from 'Widefield Bleach w fast binding_2_1' owned by user temp<br>cloned from 'Widefield Bleach w fast binding_2' owned by user temp<br>cloned from 'Widefield Bleach w binding_1' owned by user temp** |

| **Overriden Parameters** | | |
|---|---|---|
| **Name** | **Actual Value** | **Default Value** |
| kbleach_BleachFree | 10.0 | 1.0 |
| Kf_BindD | 100.0 | 1.0 |
| kbleach_BleachBound | 10.0 | 1.0 |
| duration | "0.3", "3.0" | 1.0 |
| Kr_BindF | 100.0 | 1.0 |
| Kr_BindD | 100.0 | 1.0 |
| Fluor_diffusionRate | "1.0", "10.0" | 10.0 |
| Dark_diffusionRate | Fluor_diffusionRate | 10.0 |
| sigmaaxial | 30.0 | 1.5 |
| Kf_BindF | 100.0 | 1.0 |

| **Geometry Setting** | |
|---|---|
| Geometry Size (um) | (74.24, 74.24, 26.0) |
| Mesh Size (elements) | (256, 256, 90) |

| **Advanced Settings** |
|---|

| | |
|---|---|
| Solver Name | Fully-Implicit Finite Volume, Regular Grid (Variable Time Step) |
| Time Bounds - Starting | 0.0 |
| Time Bounds - Ending | 20.0 |
| Time Step - Min | 0.0 |
| Time Step - Default | 0.05 |
| Time Step - Max | 0.1 |
| Error Tolerance - Absolute | 1.0E-9 |
| Error Tolerance - Relative | 1.0E-7 |
| Output Time Step | 0.5 |
| Use Symbolic Jacobian (T/F) | F |

#### 2.13.6.5. 2Photon Bleach w default binding_2_1

| **Simulation Name: 2Photon Bleach w default binding_2_1** |
|---|
| **Simulation Description: cloned from '2Photon Bleach w default binding_2_1' owned by user temp<br>cloned from '2Photon Bleach w default binding_2' owned by user temp<br>cloned from '2Photon Bleach w slower binding' owned by user temp<br>cloned from '2Photon Bleach w binding_1' owned by user temp** |

| **Overriden Parameters** | | |
|---|---|---|
| **Name** | **Actual Value** | **Default Value** |
| Fluor_diffusionRate | "1.0", "10.0" | 10.0 |
| kbleach_BleachFree | 10.0 | 1.0 |
| kbleach_BleachBound | 10.0 | 1.0 |
| Dark_diffusionRate | Fluor_diffusionRate | 10.0 |

| **Geometry Setting** | |
|---|---|
| Geometry Size (um) | (74.24, 74.24, 26.0) |

| | |
|---|---|
| Mesh Size (elements) | (256, 256, 90) |

| Advanced Settings | |
|---|---|
| Solver Name | Fully-Implicit Finite Volume, Regular Grid (Variable Time Step) |
| Time Bounds - Starting | 0.0 |
| Time Bounds - Ending | 20.0 |
| Time Step - Min | 0.0 |
| Time Step - Default | 0.05 |
| Time Step - Max | 0.1 |
| Error Tolerance - Absolute | 1.0E-9 |
| Error Tolerance - Relative | 1.0E-7 |
| Output Time Step | 0.5 |
| Use Symbolic Jacobian (T/F) | F |

#### 2.13.6.6. Widefield Bleach w defalt binding_2_1

| Simulation Name: Widefield Bleach w defalt binding_2_1 |
|---|
| Simulation Description: cloned from 'Widefield Bleach w defalt binding_2_1' owned by user temp<br>cloned from 'Widefield Bleach w defalt binding_2' owned by user temp<br>cloned from 'Widefield Bleach w slower binding' owned by user temp<br>cloned from 'Widefield Bleach w binding_1' owned by user temp |

| Overriden Parameters | | |
|---|---|---|
| **Name** | **Actual Value** | **Default Value** |
| Fluor_diffusionRate | "1.0", "10.0" | 10.0 |
| sigmaaxial | 30.0 | 1.5 |
| kbleach_BleachFree | 10.0 | 1.0 |
| duration | "0.3", "3.0" | 1.0 |

| | | |
|---|---|---|
| kbleach_BleachBound | 10.0 | 1.0 |
| Dark_diffusionRate | Fluor_diffusionRate | 10.0 |

| Geometry Setting | |
|---|---|
| Geometry Size (um) | (74.24, 74.24, 26.0) |
| Mesh Size (elements) | (256, 256, 90) |

| Advanced Settings | |
|---|---|
| Solver Name | Fully-Implicit Finite Volume, Regular Grid (Variable Time Step) |
| Time Bounds - Starting | 0.0 |
| Time Bounds - Ending | 20.0 |
| Time Step - Min | 0.0 |
| Time Step - Default | 0.05 |
| Time Step - Max | 0.1 |
| Error Tolerance - Absolute | 1.0E-9 |
| Error Tolerance - Relative | 1.0E-7 |
| Output Time Step | 0.5 |
| Use Symbolic Jacobian (T/F) | F |

#### 2.13.6.7. 2Photon Bleach w slow binding_2_1

| **Simulation Name: 2Photon Bleach w slow binding_2_1** |
|---|
| **Simulation Description: cloned from '2Photon Bleach w slow binding_2_1' owned by user temp<br>cloned from '2Photon Bleach w slow binding_2' owned by user temp<br>cloned from 'Copy of 2Photon Bleach w slow binding' owned by user temp<br>cloned from '2Photon Bleach w slower binding' owned by user temp<br>cloned from '2Photon Bleach w binding_1' owned by user temp** |

| Overriden Parameters | | |
|---|---|---|
| **Name** | **Actual Value** | **Default Value** |
| Fluor_diffusionRate | "1.0", "10.0" | 10.0 |
| Kf_BindF | 0.1 | 1.0 |
| Kr_BindF | 0.1 | 1.0 |
| Kf_BindD | 0.1 | 1.0 |
| Kr_BindD | 0.1 | 1.0 |
| kbleach_BleachFree | 10.0 | 1.0 |
| kbleach_BleachBound | 10.0 | 1.0 |
| Dark_diffusionRate | Fluor_diffusionRate | 10.0 |

| Geometry Setting | |
|---|---|
| Geometry Size (um) | (74.24, 74.24, 26.0) |
| Mesh Size (elements) | (256, 256, 90) |

| Advanced Settings | |
|---|---|
| Solver Name | Fully-Implicit Finite Volume, Regular Grid (Variable Time Step) |
| Time Bounds - Starting | 0.0 |
| Time Bounds - Ending | 20.0 |
| Time Step - Min | 0.0 |
| Time Step - Default | 0.05 |
| Time Step - Max | 0.1 |
| Error Tolerance - Absolute | 1.0E-9 |
| Error Tolerance - Relative | 1.0E-7 |

| | |
|---|---|
| Output Time Step | 0.5 |
| Use Symbolic Jacobian (T/F) | F |

#### 2.13.6.8. Widefield Bleach w slow binding_2_1

| **Simulation Name: Widefield Bleach w slow binding_2_1** |
|---|
| **Simulation Description: cloned from 'Widefield Bleach w slow binding_2_1' owned by user temp<br>cloned from 'Widefield Bleach w slow binding_2' owned by user temp<br>cloned from 'Copy of Widefield Bleach w slow binding' owned by user temp<br>cloned from 'Widefield Bleach w slower binding' owned by user temp<br>cloned from 'Widefield Bleach w binding_1' owned by user temp** |

| **Overriden Parameters** | | |
|---|---|---|
| **Name** | **Actual Value** | **Default Value** |
| kbleach_BleachFree | 10.0 | 1.0 |
| Kf_BindD | 0.1 | 1.0 |
| kbleach_BleachBound | 10.0 | 1.0 |
| duration | "0.3", "3.0" | 1.0 |
| Kr_BindF | 0.1 | 1.0 |
| Kr_BindD | 0.1 | 1.0 |
| Fluor_diffusionRate | "1.0", "10.0" | 10.0 |
| Dark_diffusionRate | Fluor_diffusionRate | 10.0 |
| sigmaaxial | 30.0 | 1.5 |
| Kf_BindF | 0.1 | 1.0 |

| **Geometry Setting** | |
|---|---|
| Geometry Size (um) | (74.24, 74.24, 26.0) |
| Mesh Size (elements) | (256, 256, 90) |

| Advanced Settings | |
|---|---|
| Solver Name | Fully-Implicit Finite Volume, Regular Grid (Variable Time Step) |
| Time Bounds - Starting | 0.0 |
| Time Bounds - Ending | 20.0 |
| Time Step - Min | 0.0 |
| Time Step - Default | 0.05 |
| Time Step - Max | 0.1 |
| Error Tolerance - Absolute | 1.0E-9 |
| Error Tolerance - Relative | 1.0E-7 |
| Output Time Step | 0.5 |
| Use Symbolic Jacobian (T/F) | F |

#### 2.14. Application: ProjectZ Convolved Image-based half bleach

| **Application Name: ProjectZ Convolved Image-based half bleach** |
|---|
| **Application Description: (copied from ProjectZ Convolved Image-based center circular bleach) (copied from Convolved Image-based center circular bleach) (copied from Image-based center circular bleach) (copied from Image-based center bleach) (copied from Spherical_Cell_Gaussian_Bleach)** |

##### 2.14.1. Structure Mapping For ProjectZ Convolved Image-based half bleach

| Structure Mapping | | | | |
|---|---|---|---|---|
| **Structure** | **Subdomain** | **Resolved (T/F)** | **Surf/Vol** | **VolFract** |
| EC | ec | F | | |
| NUC | Nucleus | F | | |
| CYT | cytosol | F | | |

##### 2.14.2. Reaction Mapping For ProjectZ Convolved Image-based half bleach

| Reaction Mapping | | | |
|---|---|---|---|
| **Name** | **Type** | **Enabled (T/F)** | **Fast (T/F)** |
| BindD | Reaction | T | F |
| BindF | Reaction | T | F |
| BleachFree | Reaction | T | F |
| BleachBound | Reaction | T | F |

| Initial Conditions | | | | |
|---|---|---|---|---|
| **Species** | **Structure** | **Initial Conc.** | **Diffusion Const.** | **Fixed (T/F)** |
| s0 | CYT | 2.701562118716424 M | 10.0 $m^2.s^1$ | F |
| s1 | CYT | 0.0 M | 10.0 $m^2.s^1$ | F |
| s2 | CYT | 2.701562118716425 M | 0.0 $m^2.s^1$ | F |
| s3 | CYT | 7.2984378812835775 M | 0.0 $m^2.s^1$ | F |
| s4 | CYT | (exp( - (((z - 13.0) ^ 2.0) / (2.0 * (sigmaaxial ^ 2.0)))) * (y < 39.0)) M | 0.0 $m^2.s^1$ | F |
| s5 | CYT | 0.0 M | 0.0 $m^2.s^1$ | F |

#### 2.14.3. Membrane Mapping For ProjectZ Convolved Image-based half bleach

| Electrical Mapping - Membrane Potential | | | |
|---|---|---|---|
| **Membrane** | **Calculate V (T/F)** | **V initial** | **Specific Capacitance** |
| PM | F | 0.0 mV | 1.0 $pF.m^2$ |
| NM | F | 0.0 mV | 1.0 $pF.m^2$ |

| |
|---|
| Temperature: 300.0 K |

##### 2.14.4. Geometry: Site visit _Application0_20111127_695607844

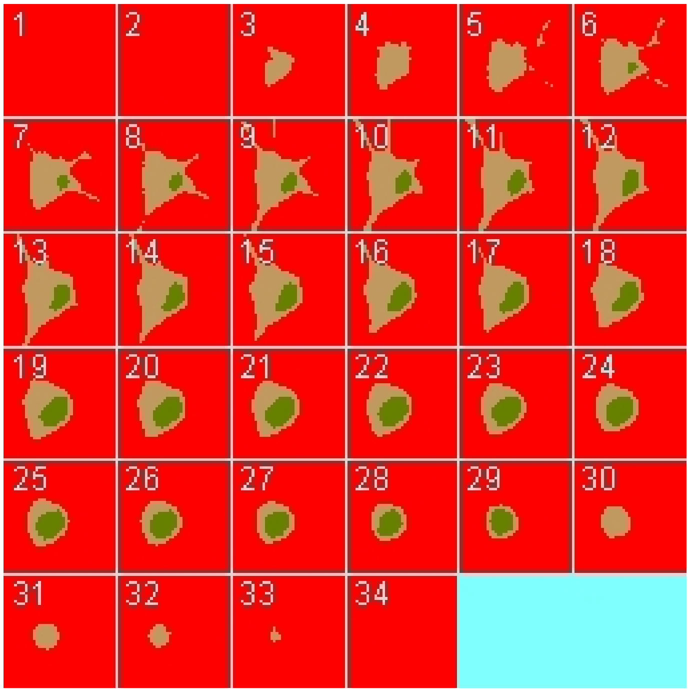

| | |
|---|---|
| Size | (74.24, 74.24, 26.0) |
| Origin | (0.0, 0.0, 0.0) |

##### 2.14.5. Math Description: Copy of ProjectZ Convolved  Image-based center circular bleach_generated

###### 2.14.5.1. Constants

| Constant Name | Expression |
|---|---|
| _F_ | 96485.3321 |
| _F_nmol_ | 9.64853321E-5 |
| _K_GHK_ | 1.0E-9 |
| _N_pmol_ | 6.02214179E11 |
| _PI_ | 3.141592653589793 |
| _R_ | 8314.46261815 |

| Constant Name | Expression |
|---|---|
| _T_ | 300.0 |
| AreaPerUnitArea_NM | 1.0 |
| AreaPerUnitArea_PM | 1.0 |
| Binder_init_uM | 2.701562118716425 |
| BleachRadius | 2.0 |
| Dark_diffusionRate | 10.0 |
| Dark_init_uM | 0.0 |
| DarkB_init_uM | 0.0 |
| duration | 1.0 |
| Fluor_diffusionRate | 10.0 |
| Fluor_init_uM | 2.701562118716424 |
| FluorB_init_uM | 7.2984378812835775 |
| K_millivolts_per_volt | 1000.0 |
| kbleach_BleachBound | 1.0 |
| kbleach_BleachFree | 1.0 |
| Kf_BindD | 1.0 |
| Kf_BindF | 1.0 |
| KMOLE | 0.001660538783162726 |
| Kr_BindD | 1.0 |
| Kr_BindF | 1.0 |
| Kr_BleachBound | 0.0 |
| Kr_BleachFree | 0.0 |
| sigmaaxial | 1.5 |
| sigmalateral | 0.5 |
| start | 1.0 |
| UnitFactor_molecules_um_neg_3_uM_neg_1 | (1.0 * pow(KMOLE,-1.0)) |
| Voltage_NM | 0.0 |

| Constant Name | Expression |
|---|---|
| Voltage_PM | 0.0 |
| VolumePerUnitVolume_CYT | 1.0 |
| VolumePerUnitVolume_EC | 1.0 |
| VolumePerUnitVolume_NUC | 1.0 |

##### 2.14.5.2. Functions

| Function Name | Expression |
|---|---|
| J_BindD | (((Kf_BindD * Dark) * Binder) - (Kr_BindD * DarkB)) |
| J_BindF | (((Kf_BindF * Fluor) * Binder) - (Kr_BindF * FluorB)) |
| J_BleachBound | ((Kf_BleachBound * FluorB) - (Kr_BleachBound * DarkB)) |
| J_BleachFree | ((Kf_BleachFree * Fluor) - (Kr_BleachFree * Dark)) |
| Kf_BleachBound | ((kbleach_BleachBound * Laser) * (t > start) * (t < (start + duration))) |
| Kf_BleachFree | ((kbleach_BleachFree * Laser) * (t > start) * (t < (start + duration))) |
| Laser_init_uM | (exp( - (((z - 13.0) ^ 2.0) / (2.0 * (sigmaaxial ^ 2.0)))) * (y < 39.0)) |
| Size_CYT | (VolumePerUnitVolume_CYT * vcRegionVolume('cytosol')) |
| Size_EC | (VolumePerUnitVolume_EC * vcRegionVolume('ec')) |
| Size_NM | (AreaPerUnitArea_NM * vcRegionArea('Nucleus_cytosol_membrane')) |
| Size_NUC | (VolumePerUnitVolume_NUC * vcRegionVolume('Nucleus')) |
| Size_PM | (AreaPerUnitArea_PM * vcRegionArea('cytosol_ec_membrane')) |
| sobj_cytosol1_ec0_size | vcRegionArea('cytosol_ec_membrane') |
| sobj_Nucleus2_cytosol1_size | vcRegionArea('Nucleus_cytosol_membrane') |
| vobj_cytosol1_size | vcRegionVolume('cytosol') |
| vobj_ec0_size | vcRegionVolume('ec') |
| vobj_Nucleus2_size | vcRegionVolume('Nucleus') |

#### 2.14.5.3. Volume Domains

### 2.14.5.3.1. ec

##### 2.14.5.3.2. cytosol

| PdeEquation Fluor | |
|---|---|
| Rate | ( - J_BindF - J_BleachFree) |
| Diffusion | Fluor_diffusionRate |
| Initial | Fluor_init_uM |

| PdeEquation Dark | |
|---|---|
| Rate | ( - J_BindD + J_BleachFree) |
| Diffusion | Dark_diffusionRate |
| Initial | Dark_init_uM |

| OdeEquation Binder | |
|---|---|
| Rate | ( - J_BindD - J_BindF) |
| Initial | Binder_init_uM |

| OdeEquation FluorB | |
|---|---|
| Rate | (J_BindF - J_BleachBound) |
| Initial | FluorB_init_uM |

| OdeEquation Laser | |
|---|---|
| Rate | 0.0 |
| Initial | Laser_init_uM |

| OdeEquation DarkB | |
|---|---|
| Rate | (J_BindD + J_BleachBound) |

| OdeEquation DarkB | |
|---|---|
| Initial | DarkB_init_uM |

##### 2.14.5.3.3. Nucleus

##### 2.14.5.4. Membrane Domains

##### 2.14.5.4.1. cytosol_ec_membrane

| JumpCondition Fluor | |
|---|---|
| InFlux | 0.0 |
| OutFlux | 0.0 |

| JumpCondition Dark | |
|---|---|
| InFlux | 0.0 |
| OutFlux | 0.0 |

##### 2.14.5.4.2. Nucleus_cytosol_membrane

| JumpCondition Fluor | |
|---|---|
| InFlux | 0.0 |
| OutFlux | 0.0 |

| JumpCondition Dark | |
|---|---|
| InFlux | 0.0 |
| OutFlux | 0.0 |

##### 2.14.6. Simulation(s)

##### 2.14.6.1. Widefield Bleach no binding_1_1_1

| |
|---|
| **Simulation Name: Widefield Bleach no binding_1_1_1** |
| **Simulation Description: cloned from 'Widefield Bleach no binding_1_1_1' owned by user temp** |

**cloned from 'Widefield Bleach no binding_1_1' owned by user temp**
**cloned from 'Widefield Bleach no binding_1' owned by user temp**
**cloned from 'Widefield Bleach_1' owned by user temp**

| **Overriden Parameters** | | |
|---|---|---|
| **Name** | **Actual Value** | **Default Value** |
| kbleach_BleachFree | 10.0 | 1.0 |
| kbleach_BleachBound | 10.0 | 1.0 |
| duration | "0.3", "3.0" | 1.0 |
| Fluor_init_uM | 10.0 | 2.701562118716424 |
| Fluor_diffusionRate | "1.0", "10.0" | 10.0 |
| Dark_diffusionRate | Fluor_diffusionRate | 10.0 |
| sigmaaxial | 30.0 | 1.5 |
| Binder_init_uM | 0.0 | 2.701562118716425 |
| FluorB_init_uM | 0.0 | 7.2984378812835775 |

| **Geometry Setting** | |
|---|---|
| Geometry Size (um) | (74.24, 74.24, 26.0) |
| Mesh Size (elements) | (256, 256, 90) |

| **Advanced Settings** | |
|---|---|
| Solver Name | Fully-Implicit Finite Volume, Regular Grid (Variable Time Step) |
| Time Bounds - Starting | 0.0 |
| Time Bounds - Ending | 20.0 |
| Time Step - Min | 0.0 |
| Time Step - Default | 0.05 |

| | |
|---|---|
| Time Step - Max | 0.1 |
| Error Tolerance - Absolute | 1.0E-9 |
| Error Tolerance - Relative | 1.0E-7 |
| Output Time Step | 0.5 |
| Use Symbolic Jacobian (T/F) | F |

##### 2.14.6.2. Widefield Bleach w fast binding_2_1_1

| **Simulation Name: Widefield Bleach w fast binding_2_1_1** |
|---|
| **Simulation Description: cloned from 'Widefield Bleach w fast binding_2_1_1' owned by user temp<br>cloned from 'Widefield Bleach w fast binding_2_1' owned by user temp<br>cloned from 'Widefield Bleach w fast binding_2' owned by user temp<br>cloned from 'Widefield Bleach w binding_1' owned by user temp** |

| **Overriden Parameters** | | |
|---|---|---|
| **Name** | **Actual Value** | **Default Value** |
| kbleach_BleachFree | 10.0 | 1.0 |
| Kf_BindD | 100.0 | 1.0 |
| kbleach_BleachBound | 10.0 | 1.0 |
| duration | "0.3", "3.0" | 1.0 |
| Kr_BindF | 100.0 | 1.0 |
| Kr_BindD | 100.0 | 1.0 |
| Fluor_diffusionRate | "1.0", "10.0" | 10.0 |
| Dark_diffusionRate | Fluor_diffusionRate | 10.0 |
| sigmaaxial | 30.0 | 1.5 |
| Kf_BindF | 100.0 | 1.0 |

| **Geometry Setting** |
|---|

| | |
|---|---|
| Geometry Size (um) | (74.24, 74.24, 26.0) |
| Mesh Size (elements) | (256, 256, 90) |

| **Advanced Settings** | |
|---|---|
| Solver Name | Fully-Implicit Finite Volume, Regular Grid (Variable Time Step) |
| Time Bounds - Starting | 0.0 |
| Time Bounds - Ending | 20.0 |
| Time Step - Min | 0.0 |
| Time Step - Default | 0.05 |
| Time Step - Max | 0.1 |
| Error Tolerance - Absolute | 1.0E-9 |
| Error Tolerance - Relative | 1.0E-7 |
| Output Time Step | 0.5 |
| Use Symbolic Jacobian (T/F) | F |

##### 2.14.6.3. Widefield Bleach w defalt binding_2_1_1

| **Simulation Name: Widefield Bleach w defalt binding_2_1_1** |
|---|
| **Simulation Description: cloned from 'Widefield Bleach w defalt binding_2_1_1' owned by user temp<br>cloned from 'Widefield Bleach w defalt binding_2_1' owned by user temp<br>cloned from 'Widefield Bleach w defalt binding_2' owned by user temp<br>cloned from 'Widefield Bleach w slower binding' owned by user temp<br>cloned from 'Widefield Bleach w binding_1' owned by user temp** |

| **Overriden Parameters** | | |
|---|---|---|
| **Name** | **Actual Value** | **Default Value** |
| Fluor_diffusionRate | "1.0", "10.0" | 10.0 |
| sigmaaxial | 30.0 | 1.5 |

| | | |
|---|---|---|
| kbleach_BleachFree | 10.0 | 1.0 |
| duration | "0.3", "3.0" | 1.0 |
| kbleach_BleachBound | 10.0 | 1.0 |
| Dark_diffusionRate | Fluor_diffusionRate | 10.0 |

| Geometry Setting | |
|---|---|
| Geometry Size (um) | (74.24, 74.24, 26.0) |
| Mesh Size (elements) | (256, 256, 90) |

| Advanced Settings | |
|---|---|
| Solver Name | Fully-Implicit Finite Volume, Regular Grid (Variable Time Step) |
| Time Bounds - Starting | 0.0 |
| Time Bounds - Ending | 20.0 |
| Time Step - Min | 0.0 |
| Time Step - Default | 0.05 |
| Time Step - Max | 0.1 |
| Error Tolerance - Absolute | 1.0E-9 |
| Error Tolerance - Relative | 1.0E-7 |
| Output Time Step | 0.5 |
| Use Symbolic Jacobian (T/F) | F |

##### 2.14.6.4. Widefield Bleach w slow binding_2_1_1

| Simulation Name: Widefield Bleach w slow binding_2_1_1 |
|---|
| **Simulation Description: cloned from 'Widefield Bleach w slow binding_2_1_1' owned by user temp**<br>**cloned from 'Widefield Bleach w slow binding_2_1' owned by user temp**<br>**cloned from 'Widefield Bleach w slow binding_2' owned by user temp**<br>**cloned from 'Copy of Widefield Bleach w slow binding' owned by user temp** |

**cloned from 'Widefield Bleach w slower binding' owned by user temp**
**cloned from 'Widefield Bleach w binding_1' owned by user temp**

| Overriden Parameters | | |
|---|---|---|
| **Name** | **Actual Value** | **Default Value** |
| kbleach_BleachFree | 10.0 | 1.0 |
| Kf_BindD | 0.1 | 1.0 |
| kbleach_BleachBound | 10.0 | 1.0 |
| duration | "0.3", "3.0" | 1.0 |
| Kr_BindF | 0.1 | 1.0 |
| Kr_BindD | 0.1 | 1.0 |
| Fluor_diffusionRate | "1.0", "10.0" | 10.0 |
| Dark_diffusionRate | Fluor_diffusionRate | 10.0 |
| sigmaaxial | 30.0 | 1.5 |
| Kf_BindF | 0.1 | 1.0 |

| Geometry Setting | |
|---|---|
| Geometry Size (um) | (74.24, 74.24, 26.0) |
| Mesh Size (elements) | (256, 256, 90) |

| Advanced Settings | |
|---|---|
| Solver Name | Fully-Implicit Finite Volume, Regular Grid (Variable Time Step) |
| Time Bounds - Starting | 0.0 |
| Time Bounds - Ending | 20.0 |
| Time Step - Min | 0.0 |
| Time Step - Default | 0.05 |

| | |
|---|---|
| Time Step - Max | 0.1 |
| Error Tolerance - Absolute | 1.0E-9 |
| Error Tolerance - Relative | 1.0E-7 |
| Output Time Step | 0.5 |
| Use Symbolic Jacobian (T/F) | F |

#### 2.15. Application: Non spatial to determine SS - vary Kd

| **Application Name: Non spatial to determine SS - vary Kd** |
|---|
| **Application Description: (copied from Non spatial to determine SS) (copied from Image-based center circular bleach) (copied from Image-based center bleach) (copied from Spherical_Cell_Gaussian_Bleach)** |

##### 2.15.1. Structure Mapping For Non spatial to determine SS - vary Kd

| **Structure Mapping** | | | | |
|---|---|---|---|---|
| **Structure** | **Subdomain** | **Resolved (T/F)** | **Surf/Vol** | **VolFract** |
| EC | Compartment | F | | |
| NUC | Compartment | F | | |
| CYT | Compartment | F | | |

##### 2.15.2. Reaction Mapping For Non spatial to determine SS - vary Kd

| **Reaction Mapping** | | | |
|---|---|---|---|
| **Name** | **Type** | **Enabled (T/F)** | **Fast (T/F)** |
| BindD | Reaction | T | F |
| BindF | Reaction | T | F |
| BleachFree | Reaction | T | F |
| BleachBound | Reaction | T | F |

| **Initial Conditions** |
|---|

| Initial Conditions | | | | |
|---|---|---|---|---|
| **Species** | **Structure** | **Initial Conc.** | **Diffusion Const.** | **Fixed (T/F)** |
| s0 | CYT | 10.0 M | 0.0 $m^2.s^1$ | F |
| s1 | CYT | 0.0 M | 0.0 $m^2.s^1$ | F |
| s2 | CYT | 10.0 M | 0.0 $m^2.s^1$ | F |
| s3 | CYT | 0.0 M | 0.0 $m^2.s^1$ | F |
| s4 | CYT | 0.0 M | 0.0 $m^2.s^1$ | F |
| s5 | CYT | 0.0 M | 0.0 $m^2.s^1$ | F |

#### 2.15.3. Membrane Mapping For Non spatial to determine SS - vary Kd

| Electrical Mapping - Membrane Potential | | | |
|---|---|---|---|
| **Membrane** | **Calculate V (T/F)** | **V initial** | **Specific Capacitance** |
| PM | F | 0.0 mV | 1.0 $pF.m^2$ |
| NM | F | 0.0 mV | 1.0 $pF.m^2$ |

| |
|---|
| Temperature: 300.0 K |

#### 2.15.4. Geometry: nonspatial267681593

Non spatial geometry.

#### 2.15.5. Math Description: Copy of Non spatial to determine SS_generated

##### 2.15.5.1. Constants

| Constant Name | Expression |
|---|---|
| _F_ | 96485.3321 |
| _F_nmol_ | 9.64853321E-5 |

| Constant Name | Expression |
|---|---|
| _K_GHK_ | 1.0E-9 |
| _N_pmol_ | 6.02214179E11 |
| _PI_ | 3.141592653589793 |
| _R_ | 8314.46261815 |
| _T_ | 300.0 |
| Binder_init_uM | 10.0 |
| BleachRadius | 2.0 |
| Dark_init_uM | 0.0 |
| DarkB_init_uM | 0.0 |
| duration | 1.0 |
| Fluor_init_uM | 10.0 |
| FluorB_init_uM | 0.0 |
| K_millivolts_per_volt | 1000.0 |
| kbleach_BleachBound | 1.0 |
| kbleach_BleachFree | 1.0 |
| Kf_BindD | 1.0 |
| Kf_BindF | 1.0 |
| KMOLE | 0.001660538783162726 |
| Kr_BindD | 1.0 |
| Kr_BindF | 1.0 |
| Kr_BleachBound | 0.0 |
| Kr_BleachFree | 0.0 |
| Laser_init_uM | 0.0 |
| sigmaaxial | 1.5 |
| sigmalateral | 0.5 |
| start | 1.0 |
| Voltage_NM | 0.0 |
| Voltage_PM | 0.0 |

##### 2.15.5.2. Functions

| Function Name | Expression |
| --- | --- |
| J_BindD | (((Kf_BindD * Dark) * Binder) - (Kr_BindD * DarkB)) |
| J_BindF | (((Kf_BindF * Fluor) * Binder) - (Kr_BindF * FluorB)) |
| J_BleachBound | ((Kf_BleachBound * FluorB) - (Kr_BleachBound * DarkB)) |
| J_BleachFree | ((Kf_BleachFree * Fluor) - (Kr_BleachFree * Dark)) |
| Kf_BleachBound | ((kbleach_BleachBound * Laser) * (t > start) * (t < (start + duration))) |
| Kf_BleachFree | ((kbleach_BleachFree * Laser) * (t > start) * (t < (start + duration))) |
| Size_CYT | (1.0 * 14891.899581611733) |
| Size_EC | (1.0 * 124712.10435961554) |
| Size_NM | (1.0 * 1406.7733692487282) |
| Size_NUC | (1.0 * 3697.013658772733) |
| Size_PM | (1.0 * 4738.640600365477) |

##### 2.15.5.3. Volume Domains

###### 2.15.5.3.1. Compartment

| OdeEquation Fluor | |
| --- | --- |
| Rate | ( - J_BindF - J_BleachFree) |
| Initial | Fluor_init_uM |

| OdeEquation Dark | |
| --- | --- |
| Rate | ( - J_BindD + J_BleachFree) |
| Initial | Dark_init_uM |

| OdeEquation Binder | |
| --- | --- |
| Rate | ( - J_BindD - J_BindF) |
| Initial | Binder_init_uM |

| OdeEquation FluorB | |
|---|---|
| Rate | (J_BindF - J_BleachBound) |
| Initial | FluorB_init_uM |

| OdeEquation Laser | |
|---|---|
| Rate | 0.0 |
| Initial | Laser_init_uM |

| OdeEquation DarkB | |
|---|---|
| Rate | (J_BindD + J_BleachBound) |
| Initial | DarkB_init_uM |

#### 2.15.6. Simulation(s)

### 2.15.6.1. Kd=1

| **Simulation Name: Kd=1** |
|---|
| **Simulation Description: cloned from 'Kd=1' owned by user temp<br>cloned from 'Simulation0' owned by user temp** |

| Advanced Settings | |
|---|---|
| Solver Name | Combined Stiff Solver (IDA/CVODE) |
| Time Bounds - Starting | 0.0 |
| Time Bounds - Ending | 20.0 |
| Time Step - Min | 1.0E-8 |
| Time Step - Default | 0.1 |
| Time Step - Max | 1.0 |
| Error Tolerance - Absolute | 1.0E-9 |

| | |
|---|---|
| Error Tolerance - Relative | 1.0E-9 |
| Keep Every | 1 |
| Keep At Most | 1000 |
| Use Symbolic Jacobian (T/F) | F |

## 2.15.6.2. Kd=10

| **Simulation Name: Kd=10** |
|---|
| **Simulation Description: cloned from ' Kd=10' owned by user temp<br>cloned from 'Simulation0' owned by user temp** |

| **Overriden Parameters** | | |
|---|---|---|
| **Name** | **Actual Value** | **Default Value** |
| Kr_BindF | 10.0 | 1.0 |
| Kr_BindD | 10.0 | 1.0 |

| **Advanced Settings** | |
|---|---|
| Solver Name | Combined Stiff Solver (IDA/CVODE) |
| Time Bounds - Starting | 0.0 |
| Time Bounds - Ending | 20.0 |
| Time Step - Min | 1.0E-8 |
| Time Step - Default | 0.1 |
| Time Step - Max | 1.0 |
| Error Tolerance - Absolute | 1.0E-9 |
| Error Tolerance - Relative | 1.0E-9 |
| Keep Every | 1 |
| Keep At Most | 1000 |

| | |
|---|---|
| Use Symbolic Jacobian (T/F) | F |

### 2.15.6.3. Kd=0_1

| |
|---|
| **Simulation Name: Kd=0_1** |
| **Simulation Description: cloned from 'Kd=0_1' owned by user temp<br>cloned from 'Simulation0' owned by user temp** |

| **Overriden Parameters** | | |
|---|---|---|
| **Name** | **Actual Value** | **Default Value** |
| Kr_BindF | 0.1 | 1.0 |
| Kr_BindD | 0.1 | 1.0 |

| **Advanced Settings** | |
|---|---|
| Solver Name | Combined Stiff Solver (IDA/CVODE) |
| Time Bounds - Starting | 0.0 |
| Time Bounds - Ending | 20.0 |
| Time Step - Min | 1.0E-8 |
| Time Step - Default | 0.1 |
| Time Step - Max | 1.0 |
| Error Tolerance - Absolute | 1.0E-9 |
| Error Tolerance - Relative | 1.0E-9 |
| Keep Every | 1 |
| Keep At Most | 1000 |
| Use Symbolic Jacobian (T/F) | F |

### 2.15.6.4. Kd=0_01

| |
|---|
| **Simulation Name: Kd=0_01** |

| |
|---|
| **Simulation Description: cloned from 'Kd=0_01' owned by user temp cloned from 'Simulation0' owned by user temp** |

| **Overriden Parameters** | | |
|---|---|---|
| **Name** | **Actual Value** | **Default Value** |
| Kr_BindF | 0.01 | 1.0 |
| Kr_BindD | 0.01 | 1.0 |

| **Advanced Settings** | |
|---|---|
| Solver Name | Combined Stiff Solver (IDA/CVODE) |
| Time Bounds - Starting | 0.0 |
| Time Bounds - Ending | 20.0 |
| Time Step - Min | 1.0E-8 |
| Time Step - Default | 0.1 |
| Time Step - Max | 1.0 |
| Error Tolerance - Absolute | 1.0E-9 |
| Error Tolerance - Relative | 1.0E-9 |
| Keep Every | 1 |
| Keep At Most | 1000 |
| Use Symbolic Jacobian (T/F) | F |

#### 2.16. Application: ProjectZ Convolved Image-based center circular bleach vary Kd

| **Application Name: ProjectZ Convolved Image-based center circular bleach vary Kd** |
|---|
| **Application Description: (copied from ProjectZ Convolved Image-based center circular bleach) (copied from Convolved Image-based center circular bleach) (copied from Image-based center circular bleach) (copied from Image-based center bleach) (copied from Spherical_Cell_Gaussian_Bleach)** |

#### 2.16.1. Structure Mapping For ProjectZ Convolved Image-based center circular bleach vary Kd

| Structure Mapping | | | | |
|---|---|---|---|---|
| **Structure** | **Subdomain** | **Resolved (T/F)** | **Surf/Vol** | **VolFract** |
| EC | ec | F | | |
| NUC | Nucleus | F | | |
| CYT | cytosol | F | | |

#### 2.16.2. Reaction Mapping For ProjectZ Convolved Image-based center circular bleach vary Kd

| Reaction Mapping | | | |
|---|---|---|---|
| **Name** | **Type** | **Enabled (T/F)** | **Fast (T/F)** |
| BindD | Reaction | T | F |
| BindF | Reaction | T | F |
| BleachFree | Reaction | T | F |
| BleachBound | Reaction | T | F |

| Initial Conditions | | | | |
|---|---|---|---|---|
| **Species** | **Structure** | **Initial Conc.** | **Diffusion Const.** | **Fixed (T/F)** |
| s0 | CYT | 2.701562118716424 M | 10.0 $m^2.s^1$ | F |
| s1 | CYT | 0.0 M | 10.0 $m^2.s^1$ | F |
| s2 | CYT | 2.701562118716425 M | 0.0 $m^2.s^1$ | F |
| s3 | CYT | 7.2984378812835775 M | 0.0 $m^2.s^1$ | F |
| s4 | CYT | (exp( - (((z - 13.0) ^ 2.0) / (2.0 * (sigmaaxial ^ 2.0)))) * ((((x - 22.0) ^ 2.0) + ((y - 32.0) ^ 2.0)) < 4.0)) | 0.0 $m^2.s^1$ | F |

| Initial Conditions | | | | |
|---|---|---|---|---|
| Species | Structure | Initial Conc. | Diffusion Const. | Fixed (T/F) |
| | | M | | |
| s5 | CYT | 0.0 M | 0.0 $m^2.s^1$ | F |

#### 2.16.3. Membrane Mapping For ProjectZ Convolved Image-based center circular bleach vary Kd

| Electrical Mapping - Membrane Potential | | | |
|---|---|---|---|
| Membrane | Calculate V (T/F) | V initial | Specific Capacitance |
| PM | F | 0.0 mV | 1.0 $pF.m^2$ |
| NM | F | 0.0 mV | 1.0 $pF.m^2$ |

| |
|---|
| Temperature: 300.0 K |

#### 2.16.4. Geometry: Site visit _Application0_20111127_695607844

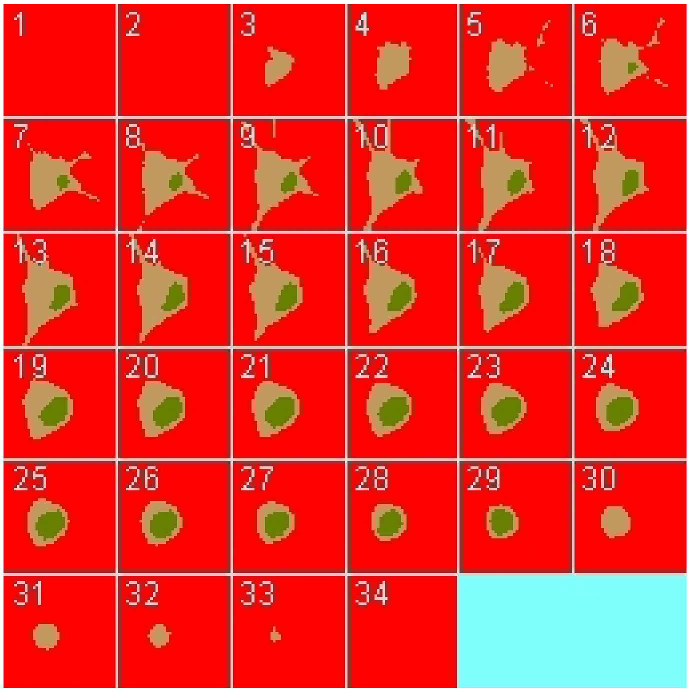

| | |
|---|---|
| Size | (74.24, 74.24, 26.0) |
| Origin | (0.0, 0.0, 0.0) |

#### 2.16.5. Math Description: Copy of ProjectZ Convolved  Image-based center circular bleach_generated

##### 2.16.5.1. Constants

| Constant Name | Expression |
|---|---|
| _F_ | 96485.3321 |
| _F_nmol_ | 9.64853321E-5 |
| _K_GHK_ | 1.0E-9 |
| _N_pmol_ | 6.02214179E11 |
| _PI_ | 3.141592653589793 |
| _R_ | 8314.46261815 |
| _T_ | 300.0 |
| AreaPerUnitArea_NM | 1.0 |
| AreaPerUnitArea_PM | 1.0 |
| Binder_init_uM | 2.701562118716425 |
| BleachRadius | 2.0 |
| Dark_diffusionRate | 10.0 |
| Dark_init_uM | 0.0 |
| DarkB_init_uM | 0.0 |
| duration | 1.0 |
| Fluor_diffusionRate | 10.0 |
| Fluor_init_uM | 2.701562118716424 |
| FluorB_init_uM | 7.2984378812835775 |
| K_millivolts_per_volt | 1000.0 |
| kbleach_BleachBound | 1.0 |
| kbleach_BleachFree | 1.0 |

| Constant Name | Expression |
|---|---|
| Kf_BindD | 1.0 |
| Kf_BindF | 1.0 |
| KMOLE | 0.001660538783162726 |
| Kr_BindD | 1.0 |
| Kr_BindF | 1.0 |
| Kr_BleachBound | 0.0 |
| Kr_BleachFree | 0.0 |
| sigmaaxial | 1.5 |
| sigmalateral | 0.5 |
| start | 1.0 |
| UnitFactor_molecules_um_neg_3_uM_neg_1 | (1.0 * pow(KMOLE,-1.0)) |
| Voltage_NM | 0.0 |
| Voltage_PM | 0.0 |
| VolumePerUnitVolume_CYT | 1.0 |
| VolumePerUnitVolume_EC | 1.0 |
| VolumePerUnitVolume_NUC | 1.0 |

#### 2.16.5.2. Functions

| Function Name | Expression |
|---|---|
| J_BindD | (((Kf_BindD * Dark) * Binder) - (Kr_BindD * DarkB)) |
| J_BindF | (((Kf_BindF * Fluor) * Binder) - (Kr_BindF * FluorB)) |
| J_BleachBound | ((Kf_BleachBound * FluorB) - (Kr_BleachBound * DarkB)) |
| J_BleachFree | ((Kf_BleachFree * Fluor) - (Kr_BleachFree * Dark)) |
| Kf_BleachBound | ((kbleach_BleachBound * Laser) * (t > start) * (t < (start + duration))) |
| Kf_BleachFree | ((kbleach_BleachFree * Laser) * (t > start) * (t < (start + duration))) |

| Function Name | Expression |
| --- | --- |
| Laser_init_uM | (exp( - (((z - 13.0) ^ 2.0) / (2.0 * (sigmaaxial ^ 2.0)))) * ((((x - 22.0) ^ 2.0) + ((y - 32.0) ^ 2.0)) < 4.0)) |
| Size_CYT | (VolumePerUnitVolume_CYT * vcRegionVolume('cytosol')) |
| Size_EC | (VolumePerUnitVolume_EC * vcRegionVolume('ec')) |
| Size_NM | (AreaPerUnitArea_NM * vcRegionArea('Nucleus_cytosol_membrane')) |
| Size_NUC | (VolumePerUnitVolume_NUC * vcRegionVolume('Nucleus')) |
| Size_PM | (AreaPerUnitArea_PM * vcRegionArea('cytosol_ec_membrane')) |
| sobj_cytosol1_ec0_size | vcRegionArea('cytosol_ec_membrane') |
| sobj_Nucleus2_cytosol1_size | vcRegionArea('Nucleus_cytosol_membrane') |
| vobj_cytosol1_size | vcRegionVolume('cytosol') |
| vobj_ec0_size | vcRegionVolume('ec') |
| vobj_Nucleus2_size | vcRegionVolume('Nucleus') |

#### 2.16.5.3. Volume Domains

### 2.16.5.3.1. ec

##### 2.16.5.3.2. cytosol

| PdeEquation Fluor | |
| --- | --- |
| Rate | ( - J_BindF - J_BleachFree) |
| Diffusion | Fluor_diffusionRate |
| Initial | Fluor_init_uM |

| PdeEquation Dark | |
| --- | --- |
| Rate | ( - J_BindD + J_BleachFree) |
| Diffusion | Dark_diffusionRate |
| Initial | Dark_init_uM |

| OdeEquation Binder | |
|---|---|
| Rate | ( - J_BindD - J_BindF) |
| Initial | Binder_init_uM |

| OdeEquation FluorB | |
|---|---|
| Rate | (J_BindF - J_BleachBound) |
| Initial | FluorB_init_uM |

| OdeEquation Laser | |
|---|---|
| Rate | 0.0 |
| Initial | Laser_init_uM |

| OdeEquation DarkB | |
|---|---|
| Rate | (J_BindD + J_BleachBound) |
| Initial | DarkB_init_uM |

#### 2.16.5.3.3. Nucleus

#### 2.16.5.4. Membrane Domains

#### 2.16.5.4.1. cytosol_ec_membrane

| JumpCondition Fluor | |
|---|---|
| InFlux | 0.0 |
| OutFlux | 0.0 |

| JumpCondition Dark | |
|---|---|
| InFlux | 0.0 |
| OutFlux | 0.0 |

##### 2.16.5.4.2. Nucleus_cytosol_membrane

| JumpCondition Fluor | |
|---|---|
| InFlux | 0.0 |
| OutFlux | 0.0 |

| JumpCondition Dark | |
|---|---|
| InFlux | 0.0 |
| OutFlux | 0.0 |

#### 2.16.6. Simulation(s)

##### 2.16.6.1. Widefield Bleach no binding_d=1

| **Simulation Name: Widefield Bleach no binding_d=1** |
|---|
| **Simulation Description: cloned from 'Widefield Bleach no binding_d=1' owned by user temp<br>cloned from 'Widefield Bleach no binding_1_1' owned by user temp<br>cloned from 'Widefield Bleach no binding_1' owned by user temp<br>cloned from 'Widefield Bleach_1' owned by user temp** |

| Overriden Parameters | | |
|---|---|---|
| **Name** | **Actual Value** | **Default Value** |
| kbleach_BleachFree | 10.0 | 1.0 |
| start | 9.0 | 1.0 |
| kbleach_BleachBound | 10.0 | 1.0 |
| Fluor_init_uM | 10.0 | 2.701562118716424 |
| Dark_diffusionRate | Fluor_diffusionRate | 10.0 |
| sigmaaxial | 30.0 | 1.5 |
| Fluor_diffusionRate | 25.0 | 10.0 |
| Binder_init_uM | 0.0 | 2.701562118716425 |

| | | |
|---|---|---|
| FluorB_init_uM | 0.0 | 7.2984378812835775 |

| Geometry Setting | |
|---|---|
| Geometry Size (um) | (74.24, 74.24, 26.0) |
| Mesh Size (elements) | (256, 256, 90) |

| Advanced Settings | |
|---|---|
| Solver Name | Fully-Implicit Finite Volume, Regular Grid (Variable Time Step) |
| Time Bounds - Starting | 0.0 |
| Time Bounds - Ending | 110.0 |
| Time Step - Min | 0.0 |
| Time Step - Default | 0.05 |
| Time Step - Max | 0.1 |
| Error Tolerance - Absolute | 1.0E-9 |
| Error Tolerance - Relative | 1.0E-7 |
| Output Time Step | 1.0 |
| Use Symbolic Jacobian (T/F) | F |

#### 2.16.6.2. Widefield Bleach Kd=1 d=1

| Simulation Name: Widefield Bleach Kd=1 d=1 |
|---|
| Simulation Description: cloned from 'Widefield Bleach Kd=1 d=1' owned by user temp<br>cloned from 'Widefield Bleach w fast binding_2_1' owned by user temp<br>cloned from 'Widefield Bleach w fast binding_2' owned by user temp<br>cloned from 'Widefield Bleach w binding_1' owned by user temp |

| Overriden Parameters | | |
|---|---|---|
| **Name** | **Actual Value** | **Default Value** |

| | | |
|---|---|---|
| Fluor_diffusionRate | 25.0 | 10.0 |
| sigmaaxial | 30.0 | 1.5 |
| kbleach_BleachFree | 10.0 | 1.0 |
| kbleach_BleachBound | 10.0 | 1.0 |
| Dark_diffusionRate | Fluor_diffusionRate | 10.0 |
| start | 9.0 | 1.0 |

| **Geometry Setting** | |
|---|---|
| Geometry Size (um) | (74.24, 74.24, 26.0) |
| Mesh Size (elements) | (256, 256, 90) |

| **Advanced Settings** | |
|---|---|
| Solver Name | Fully-Implicit Finite Volume, Regular Grid (Variable Time Step) |
| Time Bounds - Starting | 0.0 |
| Time Bounds - Ending | 110.0 |
| Time Step - Min | 0.0 |
| Time Step - Default | 0.05 |
| Time Step - Max | 0.1 |
| Error Tolerance - Absolute | 1.0E-9 |
| Error Tolerance - Relative | 1.0E-7 |
| Output Time Step | 1.0 |
| Use Symbolic Jacobian (T/F) | F |

##### 2.16.6.3. Widefield Bleach Kd=10 d=1

| **Simulation Name: Widefield Bleach Kd=10 d=1** |
|---|

**Simulation Description: cloned from 'Widefield Bleach Kd=10 d=1' owned by user temp**
**cloned from 'Widefield Bleach w defalt binding_2_1' owned by user temp**
**cloned from 'Widefield Bleach w defalt binding_2' owned by user temp**
**cloned from 'Widefield Bleach w slower binding' owned by user temp**
**cloned from 'Widefield Bleach w binding_1' owned by user temp**

| Overriden Parameters | | |
|---|---|---|
| **Name** | **Actual Value** | **Default Value** |
| kbleach_BleachFree | 10.0 | 1.0 |
| start | 9.0 | 1.0 |
| kbleach_BleachBound | 10.0 | 1.0 |
| Kr_BindF | 10.0 | 1.0 |
| Kr_BindD | 10.0 | 1.0 |
| Fluor_init_uM | 6.180339887498945 | 2.701562118716424 |
| Fluor_diffusionRate | 25.0 | 10.0 |
| Dark_diffusionRate | Fluor_diffusionRate | 10.0 |
| sigmaaxial | 30.0 | 1.5 |
| Binder_init_uM | 6.180339887498945 | 2.701562118716425 |
| FluorB_init_uM | 3.819660112501045 | 7.2984378812835775 |

| Geometry Setting | |
|---|---|
| Geometry Size (um) | (74.24, 74.24, 26.0) |
| Mesh Size (elements) | (256, 256, 90) |

| Advanced Settings | |
|---|---|
| Solver Name | Fully-Implicit Finite Volume, Regular Grid (Variable Time Step) |
| Time Bounds - Starting | 0.0 |

| | |
|---|---|
| Time Bounds - Ending | 110.0 |
| Time Step - Min | 0.0 |
| Time Step - Default | 0.05 |
| Time Step - Max | 0.1 |
| Error Tolerance - Absolute | 1.0E-9 |
| Error Tolerance - Relative | 1.0E-7 |
| Output Time Step | 1.0 |
| Use Symbolic Jacobian (T/F) | F |

#### 2.16.6.4. Widefield Bleach Kd=0_1 d=1

| **Simulation Name: Widefield Bleach Kd=0_1 d=1** |
|---|
| **Simulation Description: cloned from 'Widefield Bleach Kd=0_1 d=1' owned by user temp**<br>**cloned from 'Widefield Bleach w slow binding_2_1' owned by user temp**<br>**cloned from 'Widefield Bleach w slow binding_2' owned by user temp**<br>**cloned from 'Copy of Widefield Bleach w slow binding' owned by user temp**<br>**cloned from 'Widefield Bleach w slower binding' owned by user temp**<br>**cloned from 'Widefield Bleach w binding_1' owned by user temp** |

| **Overriden Parameters** | | |
|---|---|---|
| **Name** | **Actual Value** | **Default Value** |
| kbleach_BleachFree | 10.0 | 1.0 |
| start | 9.0 | 1.0 |
| kbleach_BleachBound | 10.0 | 1.0 |
| Kr_BindF | 0.1 | 1.0 |
| Kr_BindD | 0.1 | 1.0 |
| Fluor_init_uM | 0.95124921960895 | 2.701562118716424 |
| sigmaaxial | 30.0 | 1.5 |
| Dark_diffusionRate | Fluor_diffusionRate | 10.0 |

| | | |
|---|---|---|
| Fluor_diffusionRate | 25.0 | 10.0 |
| Binder_init_uM | 0.95124921960895 | 2.701562118716425 |
| FluorB_init_uM | 9.04875078039107 | 7.2984378812835775 |

| Geometry Setting | |
|---|---|
| Geometry Size (um) | (74.24, 74.24, 26.0) |
| Mesh Size (elements) | (256, 256, 90) |

| Advanced Settings | |
|---|---|
| Solver Name | Fully-Implicit Finite Volume, Regular Grid (Variable Time Step) |
| Time Bounds - Starting | 0.0 |
| Time Bounds - Ending | 110.0 |
| Time Step - Min | 0.0 |
| Time Step - Default | 0.05 |
| Time Step - Max | 0.1 |
| Error Tolerance - Absolute | 1.0E-9 |
| Error Tolerance - Relative | 1.0E-7 |
| Output Time Step | 1.0 |
| Use Symbolic Jacobian (T/F) | F |

##### 2.16.6.5. Widefield Bleach Kd=0_01 d=1

| **Simulation Name: Widefield Bleach Kd=0_01 d=1** |
|---|
| **Simulation Description: cloned from 'Widefield Bleach Kd=0_01 d=1' owned by user temp<br>cloned from 'Widefield Bleach w slow binding_2_1' owned by user temp<br>cloned from 'Widefield Bleach w slow binding_2' owned by user temp<br>cloned from 'Copy of Widefield Bleach w slow binding' owned by user temp<br>cloned from 'Widefield Bleach w slower binding' owned by user temp** |

**cloned from 'Widefield Bleach w binding_1' owned by user temp**

| Overriden Parameters | | |
|---|---|---|
| **Name** | **Actual Value** | **Default Value** |
| kbleach_BleachFree | 10.0 | 1.0 |
| start | 9.0 | 1.0 |
| kbleach_BleachBound | 10.0 | 1.0 |
| Kr_BindF | 0.01 | 1.0 |
| Kr_BindD | 0.01 | 1.0 |
| Fluor_init_uM | 0.31126729201739506 | 2.701562118716424 |
| Dark_diffusionRate | Fluor_diffusionRate | 10.0 |
| sigmaaxial | 30.0 | 1.5 |
| Binder_init_uM | 0.31126729201739506 | 2.701562118716425 |
| FluorB_init_uM | 9.688732707982611 | 7.2984378812835775 |

| Geometry Setting | |
|---|---|
| Geometry Size (um) | (74.24, 74.24, 26.0) |
| Mesh Size (elements) | (256, 256, 90) |

| Advanced Settings | |
|---|---|
| Solver Name | Fully-Implicit Finite Volume, Regular Grid (Variable Time Step) |
| Time Bounds - Starting | 0.0 |
| Time Bounds - Ending | 110.0 |
| Time Step - Min | 0.0 |
| Time Step - Default | 0.05 |

| | |
|---|---|
| Time Step - Max | 0.1 |
| Error Tolerance - Absolute | 1.0E-9 |
| Error Tolerance - Relative | 1.0E-7 |
| Output Time Step | 1.0 |
| Use Symbolic Jacobian (T/F) | F |

#### 2.16.6.6. Widefield Bleach no binding_d=3

| Simulation Name: Widefield Bleach no binding_d=3 |
|---|
| **Simulation Description: cloned from 'Widefield Bleach no binding_d=3' owned by user temp<br>cloned from 'Widefield Bleach no binding_d=1' owned by user temp<br>cloned from 'Widefield Bleach no binding_1_1' owned by user temp<br>cloned from 'Widefield Bleach no binding_1' owned by user temp<br>cloned from 'Widefield Bleach_1' owned by user temp** |

| Overriden Parameters | | |
|---|---|---|
| **Name** | **Actual Value** | **Default Value** |
| kbleach_BleachFree | 10.0 | 1.0 |
| start | 7.0 | 1.0 |
| kbleach_BleachBound | 10.0 | 1.0 |
| duration | 3.0 | 1.0 |
| Fluor_init_uM | 10.0 | 2.701562118716424 |
| Dark_diffusionRate | Fluor_diffusionRate | 10.0 |
| sigmaaxial | 30.0 | 1.5 |
| Fluor_diffusionRate | 25.0 | 10.0 |
| Binder_init_uM | 0.0 | 2.701562118716425 |
| FluorB_init_uM | 0.0 | 7.2984378812835775 |

| Geometry Setting |
|---|

| | |
|---|---|
| Geometry Size (um) | (74.24, 74.24, 26.0) |
| Mesh Size (elements) | (256, 256, 90) |

| **Advanced Settings** | |
|---|---|
| Solver Name | Fully-Implicit Finite Volume, Regular Grid (Variable Time Step) |
| Time Bounds - Starting | 0.0 |
| Time Bounds - Ending | 110.0 |
| Time Step - Min | 0.0 |
| Time Step - Default | 0.05 |
| Time Step - Max | 0.1 |
| Error Tolerance - Absolute | 1.0E-9 |
| Error Tolerance - Relative | 1.0E-7 |
| Output Time Step | 1.0 |
| Use Symbolic Jacobian (T/F) | F |

#### 2.16.6.7. Widefield Bleach Kd=1 d=3

| **Simulation Name: Widefield Bleach Kd=1 d=3** |
|---|
| **Simulation Description: cloned from 'Copy of Widefield Bleach Kd=1 d=1' owned by user temp<br>cloned from 'Widefield Bleach Kd=1 d=1' owned by user temp<br>cloned from 'Widefield Bleach w fast binding_2_1' owned by user temp<br>cloned from 'Widefield Bleach w fast binding_2' owned by user temp<br>cloned from 'Widefield Bleach w binding_1' owned by user temp** |

| **Overriden Parameters** | | |
|---|---|---|
| **Name** | **Actual Value** | **Default Value** |
| Fluor_diffusionRate | 25.0 | 10.0 |
| sigmaaxial | 30.0 | 1.5 |

| | | |
|---|---|---|
| kbleach_BleachFree | 10.0 | 1.0 |
| duration | 3.0 | 1.0 |
| kbleach_BleachBound | 10.0 | 1.0 |
| Dark_diffusionRate | Fluor_diffusionRate | 10.0 |
| start | 7.0 | 1.0 |

| **Geometry Setting** | |
|---|---|
| Geometry Size (um) | (74.24, 74.24, 26.0) |
| Mesh Size (elements) | (256, 256, 90) |

| **Advanced Settings** | |
|---|---|
| Solver Name | Fully-Implicit Finite Volume, Regular Grid (Variable Time Step) |
| Time Bounds - Starting | 0.0 |
| Time Bounds - Ending | 110.0 |
| Time Step - Min | 0.0 |
| Time Step - Default | 0.05 |
| Time Step - Max | 0.1 |
| Error Tolerance - Absolute | 1.0E-9 |
| Error Tolerance - Relative | 1.0E-7 |
| Output Time Step | 1.0 |
| Use Symbolic Jacobian (T/F) | F |

#### 2.16.6.8. Widefield Bleach Kd=10 d=3

| **Simulation Name: Widefield Bleach Kd=10 d=3** |
|---|
| **Simulation Description: cloned from 'Copy of Widefield Bleach Kd=10 d=1' owned by user temp cloned from 'Widefield Bleach Kd=10 d=1' owned by user temp** |

**cloned from 'Widefield Bleach w defalt binding_2_1' owned by user temp**
**cloned from 'Widefield Bleach w defalt binding_2' owned by user temp**
**cloned from 'Widefield Bleach w slower binding' owned by user temp**
**cloned from 'Widefield Bleach w binding_1' owned by user temp**

| Overriden Parameters | | |
|---|---|---|
| **Name** | **Actual Value** | **Default Value** |
| kbleach_BleachFree | 10.0 | 1.0 |
| start | 7.0 | 1.0 |
| kbleach_BleachBound | 10.0 | 1.0 |
| duration | 3.0 | 1.0 |
| Kr_BindF | 10.0 | 1.0 |
| Kr_BindD | 10.0 | 1.0 |
| Fluor_init_uM | 6.180339887498945 | 2.701562118716424 |
| Fluor_diffusionRate | 25.0 | 10.0 |
| Dark_diffusionRate | Fluor_diffusionRate | 10.0 |
| sigmaaxial | 30.0 | 1.5 |
| Binder_init_uM | 6.180339887498945 | 2.701562118716425 |
| FluorB_init_uM | 3.819660112501045 | 7.2984378812835775 |

| Geometry Setting | |
|---|---|
| Geometry Size (um) | (74.24, 74.24, 26.0) |
| Mesh Size (elements) | (256, 256, 90) |

| Advanced Settings | |
|---|---|
| Solver Name | Fully-Implicit Finite Volume, Regular Grid (Variable Time Step) |

| | |
|---|---|
| Time Bounds - Starting | 0.0 |
| Time Bounds - Ending | 110.0 |
| Time Step - Min | 0.0 |
| Time Step - Default | 0.05 |
| Time Step - Max | 0.1 |
| Error Tolerance - Absolute | 1.0E-9 |
| Error Tolerance - Relative | 1.0E-7 |
| Output Time Step | 1.0 |
| Use Symbolic Jacobian (T/F) | F |

#### 2.16.6.9. Widefield Bleach Kd=0_1 d=3

| |
|---|
| **Simulation Name: Widefield Bleach Kd=0_1 d=3** |
| **Simulation Description: cloned from 'Copy of Widefield Bleach Kd=0_1 d=1' owned by user temp<br>cloned from 'Widefield Bleach Kd=0_1 d=1' owned by user temp<br>cloned from 'Widefield Bleach w slow binding_2_1' owned by user temp<br>cloned from 'Widefield Bleach w slow binding_2' owned by user temp<br>cloned from 'Copy of Widefield Bleach w slow binding' owned by user temp<br>cloned from 'Widefield Bleach w slower binding' owned by user temp<br>cloned from 'Widefield Bleach w binding_1' owned by user temp** |

| **Overriden Parameters** | | |
|---|---|---|
| **Name** | **Actual Value** | **Default Value** |
| kbleach_BleachFree | 10.0 | 1.0 |
| start | 7.0 | 1.0 |
| kbleach_BleachBound | 10.0 | 1.0 |
| duration | 3.0 | 1.0 |
| Kr_BindF | 0.1 | 1.0 |
| Kr_BindD | 0.1 | 1.0 |

| | | |
|---|---|---|
| Fluor_init_uM | 0.95124921960895 | 2.701562118716424 |
| Fluor_diffusionRate | 25.0 | 10.0 |
| sigmaaxial | 30.0 | 1.5 |
| Dark_diffusionRate | Fluor_diffusionRate | 10.0 |
| Binder_init_uM | 0.95124921960895 | 2.701562118716425 |
| FluorB_init_uM | 9.04875078039107 | 7.2984378812835775 |

| Geometry Setting | |
|---|---|
| Geometry Size (um) | (74.24, 74.24, 26.0) |
| Mesh Size (elements) | (256, 256, 90) |

| Advanced Settings | |
|---|---|
| Solver Name | Fully-Implicit Finite Volume, Regular Grid (Variable Time Step) |
| Time Bounds - Starting | 0.0 |
| Time Bounds - Ending | 110.0 |
| Time Step - Min | 0.0 |
| Time Step - Default | 0.05 |
| Time Step - Max | 0.1 |
| Error Tolerance - Absolute | 1.0E-9 |
| Error Tolerance - Relative | 1.0E-7 |
| Output Time Step | 1.0 |
| Use Symbolic Jacobian (T/F) | F |

#### 2.16.6.10. Widefield Bleach Kd=0_01 d=3

| **Simulation Name: Widefield Bleach Kd=0_01 d=3** |
|---|

**Simulation Description: cloned from 'Copy of Widefield Bleach Kd=0_01 d=1' owned by user temp**
**cloned from 'Widefield Bleach Kd=0_01 d=1' owned by user temp**
**cloned from 'Widefield Bleach w slow binding_2_1' owned by user temp**
**cloned from 'Widefield Bleach w slow binding_2' owned by user temp**
**cloned from 'Copy of Widefield Bleach w slow binding' owned by user temp**
**cloned from 'Widefield Bleach w slower binding' owned by user temp**
**cloned from 'Widefield Bleach w binding_1' owned by user temp**

| **Overriden Parameters** | | |
|---|---|---|
| **Name** | **Actual Value** | **Default Value** |
| kbleach_BleachFree | 10.0 | 1.0 |
| start | 7.0 | 1.0 |
| kbleach_BleachBound | 10.0 | 1.0 |
| duration | 3.0 | 1.0 |
| Kr_BindF | 0.01 | 1.0 |
| Kr_BindD | 0.01 | 1.0 |
| Fluor_init_uM | 0.31126729201739506 | 2.701562118716424 |
| sigmaaxial | 30.0 | 1.5 |
| Dark_diffusionRate | Fluor_diffusionRate | 10.0 |
| Binder_init_uM | 0.31126729201739506 | 2.701562118716425 |
| FluorB_init_uM | 9.688732707982611 | 7.2984378812835775 |

| **Geometry Setting** | |
|---|---|
| Geometry Size (um) | (74.24, 74.24, 26.0) |
| Mesh Size (elements) | (256, 256, 90) |

| **Advanced Settings** | |
|---|---|
| Solver Name | Fully-Implicit Finite Volume, Regular Grid (Variable Time Step) |

| | |
|---|---|
| Time Bounds - Starting | 0.0 |
| Time Bounds - Ending | 110.0 |
| Time Step - Min | 0.0 |
| Time Step - Default | 0.05 |
| Time Step - Max | 0.1 |
| Error Tolerance - Absolute | 1.0E-9 |
| Error Tolerance - Relative | 1.0E-7 |
| Output Time Step | 1.0 |
| Use Symbolic Jacobian (T/F) | F |

#### 2.17. Application: ProjectZ Convolved Image-based half bleach vary Kd

| **Application Name: ProjectZ Convolved Image-based half bleach vary Kd** |
|---|
| **Application Description: (copied from ProjectZ Convolved Image-based half bleach) (copied from ProjectZ Convolved Image-based center circular bleach) (copied from Convolved Image-based center circular bleach) (copied from Image-based center circular bleach) (copied from Image-based center bleach) (copied from Spherical_Cell_Gaussian_Bleach)** |

##### 2.17.1. Structure Mapping For ProjectZ Convolved Image-based half bleach vary Kd

| **Structure Mapping** | | | | |
|---|---|---|---|---|
| **Structure** | **Subdomain** | **Resolved (T/F)** | **Surf/Vol** | **VolFract** |
| EC | ec | F | | |
| NUC | Nucleus | F | | |
| CYT | cytosol | F | | |

##### 2.17.2. Reaction Mapping For ProjectZ Convolved Image-based half bleach vary Kd

| **Reaction Mapping** | | | |
|---|---|---|---|
| **Name** | **Type** | **Enabled (T/F)** | **Fast (T/F)** |
| BindD | Reaction | T | F |

| Reaction Mapping | | | |
|---|---|---|---|
| **Name** | **Type** | **Enabled (T/F)** | **Fast (T/F)** |
| BindF | Reaction | T | F |
| BleachFree | Reaction | T | F |
| BleachBound | Reaction | T | F |

| Initial Conditions | | | | |
|---|---|---|---|---|
| **Species** | **Structure** | **Initial Conc.** | **Diffusion Const.** | **Fixed (T/F)** |
| s0 | CYT | 2.701562118716424 M | 10.0 $m^2.s^1$ | F |
| s1 | CYT | 0.0 M | 10.0 $m^2.s^1$ | F |
| s2 | CYT | 2.701562118716425 M | 0.0 $m^2.s^1$ | F |
| s3 | CYT | 7.2984378812835775 M | 0.0 $m^2.s^1$ | F |
| s4 | CYT | (exp( - (((z - 13.0) ^ 2.0) / (2.0 * (sigmaaxial ^ 2.0)))) * (y < 39.0)) M | 0.0 $m^2.s^1$ | F |
| s5 | CYT | 0.0 M | 0.0 $m^2.s^1$ | F |

#### 2.17.3. Membrane Mapping For ProjectZ Convolved Image-based half bleach vary Kd

| Electrical Mapping - Membrane Potential | | | |
|---|---|---|---|
| **Membrane** | **Calculate V (T/F)** | **V initial** | **Specific Capacitance** |
| PM | F | 0.0 mV | 1.0 $pF.m^2$ |
| NM | F | 0.0 mV | 1.0 $pF.m^2$ |

| |
|---|
| Temperature: 300.0 K |

#### 2.17.4. Geometry: Site visit _Application0_20111127_695607844

| | |
|---|---|
| Size | (74.24, 74.24, 26.0) |

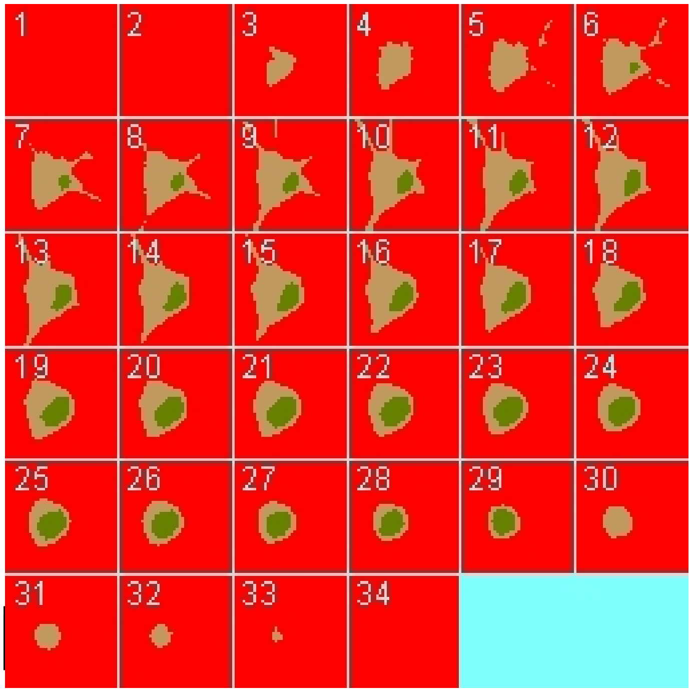

Origin (0.0, 0.0, 0.0)

##### 2.17.5. Math Description: Copy of ProjectZ Convolved Image-based half bleach_generated

###### 2.17.5.1. Constants

| Constant Name | Expression |
|---|---|
| _F_ | 96485.3321 |
| _F_nmol_ | 9.64853321E-5 |
| _K_GHK_ | 1.0E-9 |
| _N_pmol_ | 6.02214179E11 |
| _PI_ | 3.141592653589793 |
| _R_ | 8314.46261815 |
| _T_ | 300.0 |
| AreaPerUnitArea_NM | 1.0 |
| AreaPerUnitArea_PM | 1.0 |
| Binder_init_uM | 2.701562118716425 |

| Constant Name | Expression |
|---|---|
| BleachRadius | 2.0 |
| Dark_diffusionRate | 10.0 |
| Dark_init_uM | 0.0 |
| DarkB_init_uM | 0.0 |
| duration | 1.0 |
| Fluor_diffusionRate | 10.0 |
| Fluor_init_uM | 2.701562118716424 |
| FluorB_init_uM | 7.2984378812835775 |
| K_millivolts_per_volt | 1000.0 |
| kbleach_BleachBound | 1.0 |
| kbleach_BleachFree | 1.0 |
| Kf_BindD | 1.0 |
| Kf_BindF | 1.0 |
| KMOLE | 0.001660538783162726 |
| Kr_BindD | 1.0 |
| Kr_BindF | 1.0 |
| Kr_BleachBound | 0.0 |
| Kr_BleachFree | 0.0 |
| sigmaaxial | 1.5 |
| sigmalateral | 0.5 |
| start | 1.0 |
| UnitFactor_molecules_um_neg_3_uM_neg_1 | (1.0 * pow(KMOLE,-1.0)) |
| Voltage_NM | 0.0 |
| Voltage_PM | 0.0 |
| VolumePerUnitVolume_CYT | 1.0 |
| VolumePerUnitVolum | 1.0 |

| Constant Name | Expression |
| --- | --- |
| e_EC | |
| VolumePerUnitVolume_NUC | 1.0 |

#### 2.17.5.2. Functions

| Function Name | Expression |
| --- | --- |
| J_BindD | (((Kf_BindD * Dark) * Binder) - (Kr_BindD * DarkB)) |
| J_BindF | (((Kf_BindF * Fluor) * Binder) - (Kr_BindF * FluorB)) |
| J_BleachBound | ((Kf_BleachBound * FluorB) - (Kr_BleachBound * DarkB)) |
| J_BleachFree | ((Kf_BleachFree * Fluor) - (Kr_BleachFree * Dark)) |
| Kf_BleachBound | ((kbleach_BleachBound * Laser) * (t > start) * (t < (start + duration))) |
| Kf_BleachFree | ((kbleach_BleachFree * Laser) * (t > start) * (t < (start + duration))) |
| Laser_init_uM | (exp( - (((z - 13.0) ^ 2.0) / (2.0 * (sigmaaxial ^ 2.0)))) * (y < 39.0)) |
| Size_CYT | (VolumePerUnitVolume_CYT * vcRegionVolume('cytosol')) |
| Size_EC | (VolumePerUnitVolume_EC * vcRegionVolume('ec')) |
| Size_NM | (AreaPerUnitArea_NM * vcRegionArea('Nucleus_cytosol_membrane')) |
| Size_NUC | (VolumePerUnitVolume_NUC * vcRegionVolume('Nucleus')) |
| Size_PM | (AreaPerUnitArea_PM * vcRegionArea('cytosol_ec_membrane')) |
| sobj_cytosol1_ec0_size | vcRegionArea('cytosol_ec_membrane') |
| sobj_Nucleus2_cytosol1_size | vcRegionArea('Nucleus_cytosol_membrane') |
| vobj_cytosol1_size | vcRegionVolume('cytosol') |
| vobj_ec0_size | vcRegionVolume('ec') |
| vobj_Nucleus2_size | vcRegionVolume('Nucleus') |

#### 2.17.5.3. Volume Domains

### 2.17.5.3.1. ec

#### 2.17.5.3.2. cytosol

| PdeEquation Fluor | |
|---|---|
| Rate | ( - J_BindF - J_BleachFree) |
| Diffusion | Fluor_diffusionRate |
| Initial | Fluor_init_uM |

| PdeEquation Dark | |
|---|---|
| Rate | ( - J_BindD + J_BleachFree) |
| Diffusion | Dark_diffusionRate |
| Initial | Dark_init_uM |

| OdeEquation Binder | |
|---|---|
| Rate | ( - J_BindD - J_BindF) |
| Initial | Binder_init_uM |

| OdeEquation FluorB | |
|---|---|
| Rate | (J_BindF - J_BleachBound) |
| Initial | FluorB_init_uM |

| OdeEquation Laser | |
|---|---|
| Rate | 0.0 |
| Initial | Laser_init_uM |

| OdeEquation DarkB | |
|---|---|
| Rate | (J_BindD + J_BleachBound) |
| Initial | DarkB_init_uM |

#### 2.17.5.3.3. Nucleus

#### 2.17.5.4. Membrane Domains

##### 2.17.5.4.1. cytosol_ec_membrane

| JumpCondition Fluor | |
|---|---|
| InFlux | 0.0 |
| OutFlux | 0.0 |

| JumpCondition Dark | |
|---|---|
| InFlux | 0.0 |
| OutFlux | 0.0 |

##### 2.17.5.4.2. Nucleus_cytosol_membrane

| JumpCondition Fluor | |
|---|---|
| InFlux | 0.0 |
| OutFlux | 0.0 |

| JumpCondition Dark | |
|---|---|
| InFlux | 0.0 |
| OutFlux | 0.0 |

#### 2.17.6. Simulation(s)

##### 2.17.6.1. Widefield half cell Bleach no binding

| **Simulation Name: Widefield half cell Bleach no binding** |
|---|
| **Simulation Description: cloned from 'Widefield half cell Bleach no binding' owned by user temp<br>cloned from 'Widefield Bleach no binding_1_1_1' owned by user temp<br>cloned from 'Widefield Bleach no binding_1_1' owned by user temp<br>cloned from 'Widefield Bleach no binding_1' owned by user temp<br>cloned from 'Widefield Bleach_1' owned by user temp** |

| Overriden Parameters | | |
|---|---|---|
| **Name** | **Actual Value** | **Default Value** |
| Binder_init_uM | 0.0 | 2.701562118716425 |
| kbleach_BleachFree | 10.0 | 1.0 |
| sigmaaxial | 30.0 | 1.5 |
| kbleach_BleachBound | 10.0 | 1.0 |
| Dark_diffusionRate | Fluor_diffusionRate | 10.0 |
| start | 9.0 | 1.0 |
| FluorB_init_uM | 0.0 | 7.2984378812835775 |
| Fluor_init_uM | 10.0 | 2.701562118716424 |

| Geometry Setting | |
|---|---|
| Geometry Size (um) | (74.24, 74.24, 26.0) |
| Mesh Size (elements) | (256, 256, 90) |

| Advanced Settings | |
|---|---|
| Solver Name | Fully-Implicit Finite Volume, Regular Grid (Variable Time Step) |
| Time Bounds - Starting | 0.0 |
| Time Bounds - Ending | 110.0 |
| Time Step - Min | 0.0 |
| Time Step - Default | 0.05 |
| Time Step - Max | 0.1 |
| Error Tolerance - Absolute | 1.0E-9 |
| Error Tolerance - Relative | 1.0E-7 |

| | |
|---|---|
| Output Time Step | 1.0 |
| Use Symbolic Jacobian (T/F) | F |

##### 2.17.6.2. Widefield half cell Bleach Kd=1

| **Simulation Name: Widefield half cell Bleach Kd=1** |
|---|
| **Simulation Description: cloned from 'Widefield half cell Bleach Kd=1' owned by user temp<br>cloned from 'Widefield Bleach w fast binding_2_1_1' owned by user temp<br>cloned from 'Widefield Bleach w fast binding_2_1' owned by user temp<br>cloned from 'Widefield Bleach w fast binding_2' owned by user temp<br>cloned from 'Widefield Bleach w binding_1' owned by user temp** |

| **Overriden Parameters** | | |
|---|---|---|
| **Name** | **Actual Value** | **Default Value** |
| Fluor_diffusionRate | 25.0 | 10.0 |
| kbleach_BleachFree | 10.0 | 1.0 |
| sigmaaxial | 30.0 | 1.5 |
| kbleach_BleachBound | 10.0 | 1.0 |
| Dark_diffusionRate | Fluor_diffusionRate | 10.0 |
| start | 9.0 | 1.0 |

| **Geometry Setting** | |
|---|---|
| Geometry Size (um) | (74.24, 74.24, 26.0) |
| Mesh Size (elements) | (256, 256, 90) |

| **Advanced Settings** | |
|---|---|
| Solver Name | Fully-Implicit Finite Volume, Regular Grid (Variable Time Step) |
| Time Bounds - Starting | 0.0 |

| | |
|---|---|
| Time Bounds - Ending | 110.0 |
| Time Step - Min | 0.0 |
| Time Step - Default | 0.05 |
| Time Step - Max | 0.1 |
| Error Tolerance - Absolute | 1.0E-9 |
| Error Tolerance - Relative | 1.0E-7 |
| Output Time Step | 1.0 |
| Use Symbolic Jacobian (T/F) | F |

##### 2.17.6.3. Widefield half cell Bleach Kd=10

| **Simulation Name: Widefield half cell Bleach Kd=10** |
|---|
| **Simulation Description: cloned from ' Widefield half cell Bleach Kd=10' owned by user temp<br>cloned from 'Widefield Bleach w fast binding_2_1_1' owned by user temp<br>cloned from 'Widefield Bleach w fast binding_2_1' owned by user temp<br>cloned from 'Widefield Bleach w fast binding_2' owned by user temp<br>cloned from 'Widefield Bleach w binding_1' owned by user temp** |

| **Overriden Parameters** | | |
|---|---|---|
| **Name** | **Actual Value** | **Default Value** |
| kbleach_BleachFree | 10.0 | 1.0 |
| start | 9.0 | 1.0 |
| kbleach_BleachBound | 10.0 | 1.0 |
| Kr_BindF | 10.0 | 1.0 |
| Kr_BindD | 10.0 | 1.0 |
| Fluor_init_uM | 6.180339887498945 | 2.701562118716424 |
| Fluor_diffusionRate | 25.0 | 10.0 |
| Dark_diffusionRate | Fluor_diffusionRate | 10.0 |

| | | |
|---|---|---|
| sigmaaxial | 30.0 | 1.5 |
| Binder_init_uM | 6.180339887498945 | 2.701562118716425 |
| FluorB_init_uM | 3.819660112501045 | 7.2984378812835775 |

| Geometry Setting | |
|---|---|
| Geometry Size (um) | (74.24, 74.24, 26.0) |
| Mesh Size (elements) | (256, 256, 90) |

| Advanced Settings | |
|---|---|
| Solver Name | Fully-Implicit Finite Volume, Regular Grid (Variable Time Step) |
| Time Bounds - Starting | 0.0 |
| Time Bounds - Ending | 110.0 |
| Time Step - Min | 0.0 |
| Time Step - Default | 0.05 |
| Time Step - Max | 0.1 |
| Error Tolerance - Absolute | 1.0E-9 |
| Error Tolerance - Relative | 1.0E-7 |
| Output Time Step | 1.0 |
| Use Symbolic Jacobian (T/F) | F |

##### 2.17.6.4. Widefield half cell Bleach Kd=0_1

| Simulation Name: Widefield half cell Bleach Kd=0_1 |
|---|
| Simulation Description: cloned from 'Widefield half cell Bleach Kd=0_1' owned by user temp<br>cloned from 'Widefield Bleach w fast binding_2_1_1' owned by user temp<br>cloned from 'Widefield Bleach w fast binding_2_1' owned by user temp<br>cloned from 'Widefield Bleach w fast binding_2' owned by user temp<br>cloned from 'Widefield Bleach w binding_1' owned by user temp |

| Overriden Parameters | | |
|---|---|---|
| **Name** | **Actual Value** | **Default Value** |
| kbleach_BleachFree | 10.0 | 1.0 |
| start | 9.0 | 1.0 |
| kbleach_BleachBound | 10.0 | 1.0 |
| Kr_BindF | 0.1 | 1.0 |
| Kr_BindD | 0.1 | 1.0 |
| Fluor_init_uM | 0.95124921960895 | 2.701562118716424 |
| Fluor_diffusionRate | 25.0 | 10.0 |
| Dark_diffusionRate | Fluor_diffusionRate | 10.0 |
| sigmaaxial | 30.0 | 1.5 |
| Binder_init_uM | 0.95124921960895 | 2.701562118716425 |
| FluorB_init_uM | 9.04875078039107 | 7.2984378812835775 |

| Geometry Setting | |
|---|---|
| Geometry Size (um) | (74.24, 74.24, 26.0) |
| Mesh Size (elements) | (256, 256, 90) |

| Advanced Settings | |
|---|---|
| Solver Name | Fully-Implicit Finite Volume, Regular Grid (Variable Time Step) |
| Time Bounds - Starting | 0.0 |
| Time Bounds - Ending | 110.0 |
| Time Step - Min | 0.0 |
| Time Step - Default | 0.05 |

| | |
|---|---|
| Time Step - Max | 0.1 |
| Error Tolerance - Absolute | 1.0E-9 |
| Error Tolerance - Relative | 1.0E-7 |
| Output Time Step | 1.0 |
| Use Symbolic Jacobian (T/F) | F |

##### 2.17.6.5. Widefield half cell Bleach Kd=0_01

| **Simulation Name: Widefield half cell Bleach Kd=0_01** |
|---|
| **Simulation Description: cloned from 'Widefield half cell Bleach Kd=0_01' owned by user temp<br>cloned from 'Widefield Bleach w fast binding_2_1_1' owned by user temp<br>cloned from 'Widefield Bleach w fast binding_2_1' owned by user temp<br>cloned from 'Widefield Bleach w fast binding_2' owned by user temp<br>cloned from 'Widefield Bleach w binding_1' owned by user temp** |

| **Overriden Parameters** | | |
|---|---|---|
| **Name** | **Actual Value** | **Default Value** |
| kbleach_BleachFree | 10.0 | 1.0 |
| start | 9.0 | 1.0 |
| kbleach_BleachBound | 10.0 | 1.0 |
| Kr_BindF | 0.01 | 1.0 |
| Kr_BindD | 0.01 | 1.0 |
| Fluor_init_uM | 0.31126729201739506 | 2.701562118716424 |
| Fluor_diffusionRate | 25.0 | 10.0 |
| Dark_diffusionRate | Fluor_diffusionRate | 10.0 |
| sigmaaxial | 30.0 | 1.5 |
| Binder_init_uM | 0.31126729201739506 | 2.701562118716425 |
| FluorB_init_uM | 9.688732707982611 | 7.2984378812835775 |

| Geometry Setting | |
|---|---|
| Geometry Size (um) | (74.24, 74.24, 26.0) |
| Mesh Size (elements) | (256, 256, 90) |

| Advanced Settings | |
|---|---|
| Solver Name | Fully-Implicit Finite Volume, Regular Grid (Variable Time Step) |
| Time Bounds - Starting | 0.0 |
| Time Bounds - Ending | 110.0 |
| Time Step - Min | 0.0 |
| Time Step - Default | 0.05 |
| Time Step - Max | 0.1 |
| Error Tolerance - Absolute | 1.0E-9 |
| Error Tolerance - Relative | 1.0E-7 |
| Output Time Step | 1.0 |
| Use Symbolic Jacobian (T/F) | F |
