## Supplementary material for "Beyond analytic solution: analysis of FRAP experiments by spatial simulation of the forward problem": Full details of VCell BioModel "FRAP_Membrane_Rel"

### 1. Physiology For FRAP_Membrane_Rel

#### 1.1. General Info

| BioModel Name: FRAP_Membrane_Rel |
|---|
| Owner: les |

#### 1.2. Structures and Reactions Diagram

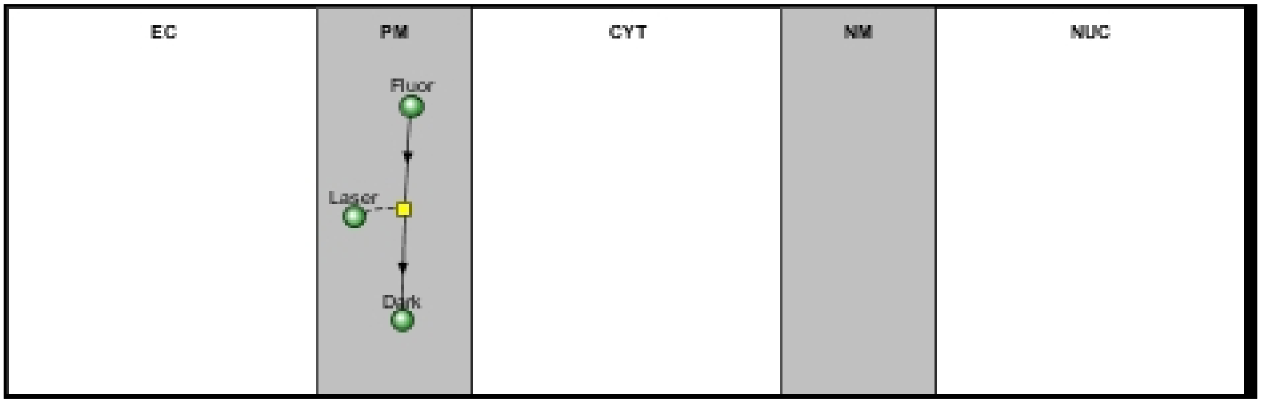

#### 1.3. Reaction(s) in PM

##### 1.3.1. Reaction r0

**Fluor** ⟷ **Dark**

| Modifiers List |
|---|
| Laser |

| Kinetics Parameters | | | |
|---|---|---|---|
| **Name** | **Expression** | **Role** | **Unit** |
| J | ((Kf * Fluor) - (Kr * Dark)) | reaction rate | molecules.m².s¹ |
| I | 0.0 | inward current density | pA.m² |
| netValence | 1.0 | net charge valence | 1 |

| Kinetics Parameters | | | |
|---|---|---|---|
| **Name** | **Expression** | **Role** | **Unit** |
| Kf | ((kbleach * Laser) * (t > start) * (t < (start + duration))) | forward rate constant | $s^{1}$ |
| Kr | 0.0 | reverse rate constant | $s^{1}$ |
| kbleach | 10.0 | user defined | $m^{2}.s^{1}.molecules^{1}$ |

### 2. Applications For FRAP_Membrane_Rel

#### 2.1. Application: Spherical_Cell_Gaussian_Bleach

| |
|---|
| **Application Name: Spherical_Cell_Gaussian_Bleach** |

##### 2.1.1. Structure Mapping For Spherical_Cell_Gaussian_Bleach

| **Structure Mapping** | | | | |
|---|---|---|---|---|
| **Structure** | **Subdomain** | **Resolved (T/F)** | **Surf/Vol** | **VolFract** |
| NUC | subdomain1 | F | | |
| CYT | subdomain1 | F | | |
| EC | subdomain0 | F | | |

##### 2.1.2. Reaction Mapping For Spherical_Cell_Gaussian_Bleach

| **Reaction Mapping** | | | |
|---|---|---|---|
| **Name** | **Type** | **Enabled (T/F)** | **Fast (T/F)** |
| r0 | Reaction | T | F |

| **Initial Conditions** | | | | |
|---|---|---|---|---|
| **Species** | **Structure** | **Initial Conc.** | **Diffusion Const.** | **Fixed (T/F)** |
| s0 | PM | 10.0 molecules.$m^2$ | 0.1 $m^2.s^1$ | F |
| s1 | PM | 0.0 molecules.$m^2$ | 0.1 $m^2.s^1$ | F |
| s2 | PM | exp( - ((((x - 11.0) ^ 2.0) / (2.0 * (sigmalateral ^ 2.0))) + (((y - 11.0) ^ 2.0) / (2.0 * (sigmalateral ^ 2.0))) + (((z - 1.0) ^ 2.0) / (2.0 * (sigmaaxial ^ 2.0))))) molecules.$m^2$ | 0.0 $m^2.s^1$ | F |

#### 2.1.3. Membrane Mapping For Spherical_Cell_Gaussian_Bleach

| Electrical Mapping - Membrane Potential | | | |
|---|---|---|---|
| **Membrane** | **Calculate V (T/F)** | **V initial** | **Specific Capacitance** |
| PM | F | 0.0 mV | 1.0 pF.m² |
| NM | F | 0.0 mV | 1.0 pF.m² |

| |
|---|
| Temperature: 300.0 K |

#### 2.1.4. Geometry: Geometry343793035

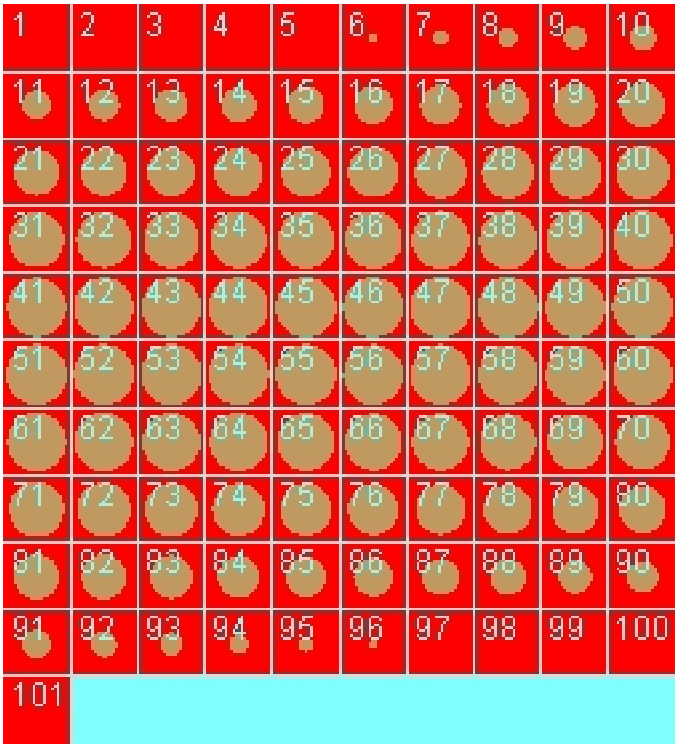

| | |
|---|---|
| Size | (22.0, 22.0, 22.0) |
| Origin | (0.0, 0.0, 0.0) |

#### 2.1.5. Math Description: Spherical_Cell_Gaussian_Bleach_generated

#### 2.1.5.1. Constants

| Constant Name | Expression |
|---|---|
| _F_ | 96485.3321 |
| _F_nmol_ | 9.64853321E-5 |
| _K_GHK_ | 1.0E-9 |
| _N_pmol_ | 6.02214179E11 |
| _PI_ | 3.141592653589793 |
| _R_ | 8314.46261815 |
| _T_ | 300.0 |
| AreaPerUnitArea_PM | 1.0 |
| AreaPerUnitVolume_NM | 1.2 |
| BleachRadius | 2.0 |
| Dark_diffusionRate | 0.1 |
| Dark_init_molecules_um_2 | 0.0 |
| duration | 1.0 |
| Fluor_diffusionRate | 0.1 |
| Fluor_init_molecules_um_2 | 10.0 |
| K_millivolts_per_volt | 1000.0 |
| kbleach | 10.0 |
| KMOLE | 0.001660538783162726 |
| Kr | 0.0 |
| netValence | 1.0 |
| sigmaaxial | 1.5 |
| sigmalateral | 0.5 |
| start | 1.0 |
| Voltage_NM | 0.0 |

| Constant Name | Expression |
| --- | --- |
| Voltage_PM | 0.0 |
| VolumePerUnitVolume_CYT | 1.0 |
| VolumePerUnitVolume_EC | 1.0 |
| VolumePerUnitVolume_NUC | 65.45 |

##### 2.1.5.2. Functions

| Function Name | Expression |
| --- | --- |
| J_r0 | ((Kf * Fluor) - (Kr * Dark)) |
| Kf | ((kbleach * Laser) * (t > start) * (t < (start + duration))) |
| Laser_init_molecules_um_2 | exp( - ((((x - 11.0) ^ 2.0) / (2.0 * (sigmalateral ^ 2.0))) + (((y - 11.0) ^ 2.0) / (2.0 * (sigmalateral ^ 2.0))) + (((z - 1.0) ^ 2.0) / (2.0 * (sigmaaxial ^ 2.0))))) |
| Size_CYT | (VolumePerUnitVolume_CYT * vcRegionVolume('subdomain1')) |
| Size_EC | (VolumePerUnitVolume_EC * vcRegionVolume('subdomain0')) |
| Size_NM | (AreaPerUnitVolume_NM * vcRegionVolume('subdomain1')) |
| Size_NUC | (VolumePerUnitVolume_NUC * vcRegionVolume('subdomain1')) |
| Size_PM | (AreaPerUnitArea_PM * vcRegionArea('subdomain0_subdomain1_membrane')) |
| sobj_subdomain11_subdomain00_size | vcRegionArea('subdomain0_subdomain1_membrane') |
| vobj_subdomain00_size | vcRegionVolume('subdomain0') |
| vobj_subdomain11_size | vcRegionVolume('subdomain1') |

##### 2.1.5.3. Volume Domains

###### 2.1.5.3.1. subdomain1

###### 2.1.5.3.2. subdomain0

#### 2.1.5.4. Membrane Domains

##### 2.1.5.4.1. subdomain0_subdomain1_membrane

| **PdeEquation Fluor** | |
|---|---|
| Rate | - J_r0 |
| Diffusion | Fluor_diffusionRate |
| Initial | Fluor_init_molecules_um_2 |

| **PdeEquation Dark** | |
|---|---|
| Rate | J_r0 |
| Diffusion | Dark_diffusionRate |
| Initial | Dark_init_molecules_um_2 |

| **OdeEquation Laser** | |
|---|---|
| Rate | 0.0 |
| Initial | Laser_init_molecules_um_2 |

#### 2.1.6. Simulation(s)

##### 2.1.6.1. Vary D and bleach Duration

| **Simulation Name: Vary D and bleach Duration** |
|---|

| **Overriden Parameters** | | |
|---|---|---|
| **Name** | **Actual Value** | **Default Value** |
| Fluor_diffusionRate | 0.1 to 1.0, 2 values | 0.1 |
| duration | "0.3", "1.0", "3.0" | 1.0 |
| Dark_diffusionRate | Fluor_diffusionRate | 0.1 |

| Geometry Setting | |
|---|---|
| Geometry Size (um) | (22.0, 22.0, 22.0) |
| Mesh Size (elements) | (51, 51, 51) |

| Advanced Settings | |
|---|---|
| Solver Name | Fully-Implicit Finite Volume, Regular Grid (Variable Time Step) |
| Time Bounds - Starting | 0.0 |
| Time Bounds - Ending | 20.0 |
| Time Step - Min | 0.0 |
| Time Step - Default | 0.05 |
| Time Step - Max | 0.1 |
| Error Tolerance - Absolute | 1.0E-9 |
| Error Tolerance - Relative | 1.0E-7 |
| Output Time Step | 0.5 |
| Use Symbolic Jacobian (T/F) | F |

#### 2.1.6.2. Widefield axial Vary D and bleach Duration

| Simulation Name: Widefield axial Vary D and bleach Duration |
|---|

| Overriden Parameters | | |
|---|---|---|
| **Name** | **Actual Value** | **Default Value** |
| Fluor_diffusionRate | 0.1 to 1.0, 2 values | 0.1 |
| sigmaaxial | 30.0 | 1.5 |
| duration | "0.1", "3.0", "1.0" | 1.0 |
| Dark_diffusionRate | Fluor_diffusionRate | 0.1 |

| Geometry Setting | |
|---|---|
| Geometry Size (um) | (22.0, 22.0, 22.0) |
| Mesh Size (elements) | (51, 51, 51) |

| Advanced Settings | |
|---|---|
| Solver Name | Fully-Implicit Finite Volume, Regular Grid (Variable Time Step) |
| Time Bounds - Starting | 0.0 |
| Time Bounds - Ending | 20.0 |
| Time Step - Min | 0.0 |
| Time Step - Default | 0.05 |
| Time Step - Max | 0.1 |
| Error Tolerance - Absolute | 1.0E-9 |
| Error Tolerance - Relative | 1.0E-7 |
| Output Time Step | 0.5 |
| Use Symbolic Jacobian (T/F) | F |

##### 2.1.6.3. Copy of Widefield axial Vary D and bleach Duration

| **Simulation Name: Copy of Widefield axial Vary D and bleach Duration** |
|---|
| **Simulation Description: cloned from 'Copy of Widefield axial Vary D and bleach Duration' owned by user temp** |

| Overriden Parameters | | |
|---|---|---|
| **Name** | **Actual Value** | **Default Value** |
| Fluor_diffusionRate | 0.1 to 1.0, 2 values | 0.1 |
| sigmaaxial | 30.0 | 1.5 |

| | | |
|---|---|---|
| duration | "0.1", "3.0", "1.0" | 1.0 |
| Dark_diffusionRate | Fluor_diffusionRate | 0.1 |

| Geometry Setting | |
|---|---|
| Geometry Size (um) | (22.0, 22.0, 22.0) |
| Mesh Size (elements) | (51, 51, 51) |

| Advanced Settings | |
|---|---|
| Solver Name | Fully-Implicit Finite Volume, Regular Grid (Variable Time Step) |
| Time Bounds - Starting | 0.0 |
| Time Bounds - Ending | 50.0 |
| Time Step - Min | 0.0 |
| Time Step - Default | 0.05 |
| Time Step - Max | 0.1 |
| Error Tolerance - Absolute | 1.0E-9 |
| Error Tolerance - Relative | 1.0E-7 |
| Output Time Step | 0.25 |
| Use Symbolic Jacobian (T/F) | F |

#### 2.2. Application: TIRF_center_bleach

| **Application Name: TIRF_center_bleach** |
|---|
| **Application Description: (copied from Spherical_Cell_Gaussian_Bleach)** |

##### 2.2.1. Structure Mapping For TIRF_center_bleach

| Structure Mapping | | | | |
|---|---|---|---|---|
| **Structure** | **Subdomain** | **Resolved (T/F)** | **Surf/Vol** | **VolFract** |

| Structure Mapping | | | | |
|---|---|---|---|---|
| **Structure** | **Subdomain** | **Resolved (T/F)** | **Surf/Vol** | **VolFract** |
| NUC | IC | F | | |
| CYT | IC | F | | |
| EC | EC | F | | |

#### 2.2.2. Reaction Mapping For TIRF_center_bleach

| Reaction Mapping | | | |
|---|---|---|---|
| **Name** | **Type** | **Enabled (T/F)** | **Fast (T/F)** |
| r0 | Reaction | T | F |

| Initial Conditions | | | | |
|---|---|---|---|---|
| **Species** | **Structure** | **Initial Conc.** | **Diffusion Const.** | **Fixed (T/F)** |
| s0 | PM | 10.0 molecules.$m^2$ | 0.1 $m^2.s^1$ | F |
| s1 | PM | 0.0 molecules.$m^2$ | 0.1 $m^2.s^1$ | F |
| s2 | PM | exp( - (((x ^ 2.0) / (2.0 * (sigmalateral ^ 2.0))) + ((y ^ 2.0) / (2.0 * (sigmalateral ^ 2.0))) + ((z ^ 2.0) / (2.0 * (sigmaaxial ^ 2.0))))) molecules.$m^2$ | 0.0 $m^2.s^1$ | F |

#### 2.2.3. Membrane Mapping For TIRF_center_bleach

| Electrical Mapping - Membrane Potential | | | |
|---|---|---|---|
| **Membrane** | **Calculate V (T/F)** | **V initial** | **Specific Capacitance** |
| PM | F | 0.0 mV | 1.0 pF.$m^2$ |
| NM | F | 0.0 mV | 1.0 pF.$m^2$ |

| |
|---|
| Temperature: 300.0 K |

#### 2.2.4. Geometry: Geometry7

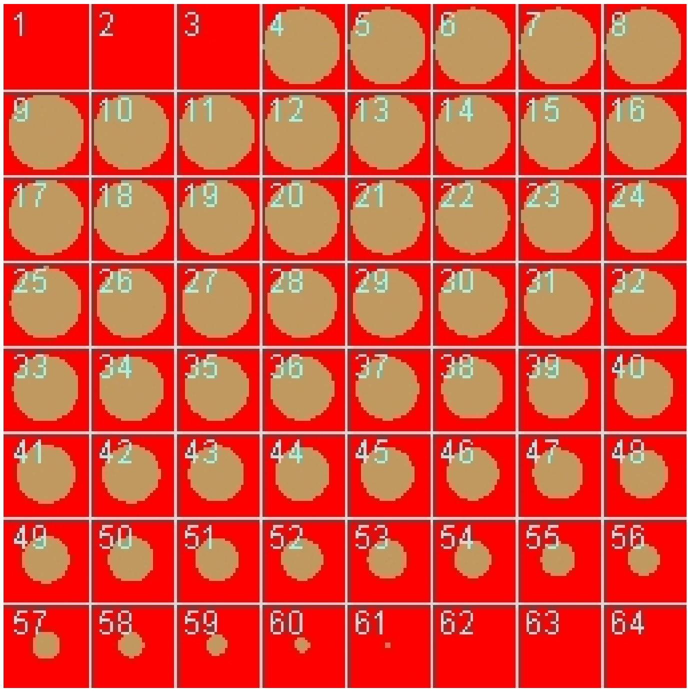

| | |
|---|---|
| Size | (22.0, 22.0, 11.0) |
| Origin | (-11.0, -11.0, -0.5) |

#### 2.2.5. Math Description: Copy of Spherical_Cell_Gaussian_Bleach_generated

##### 2.2.5.1. Constants

| Constant Name | Expression |
|---|---|
| _F_ | 96485.3321 |
| _F_nmol_ | 9.64853321E-5 |
| _K_GHK_ | 1.0E-9 |
| _N_pmol_ | 6.02214179E11 |
| _PI_ | 3.141592653589793 |

| Constant Name | Expression |
|---|---|
| _R_ | 8314.46261815 |
| _T_ | 300.0 |
| AreaPerUnitArea_PM | 1.0 |
| AreaPerUnitVolume_NM | 1.2 |
| BleachRadius | 2.0 |
| Dark_diffusionRate | 0.1 |
| Dark_init_molecules_um_2 | 0.0 |
| duration | 1.0 |
| Fluor_diffusionRate | 0.1 |
| Fluor_init_molecules_um_2 | 10.0 |
| K_millivolts_per_volt | 1000.0 |
| kbleach | 10.0 |
| KMOLE | 0.001660538783162726 |
| Kr | 0.0 |
| netValence | 1.0 |
| sigmaaxial | 1.5 |
| sigmalateral | 0.5 |
| start | 1.0 |
| Voltage_NM | 0.0 |
| Voltage_PM | 0.0 |
| VolumePerUnitVolume_CYT | 1.0 |
| VolumePerUnitVolume_EC | 1.0 |
| VolumePerUnitVolume_NUC | 65.45 |

##### 2.2.5.2. Functions

| Function Name | Expression |
|---|---|
| J_r0 | ((Kf * Fluor) - (Kr * Dark)) |
| Kf | ((kbleach * Laser) * (t > start) * (t < (start + duration))) |
| Laser_init_molecules_um_2 | exp( - (((x ^ 2.0) / (2.0 * (sigmalateral ^ 2.0))) + ((y ^ 2.0) / (2.0 * (sigmalateral ^ 2.0))) + ((z ^ 2.0) / (2.0 * (sigmaaxial ^ 2.0))))) |
| Size_CYT | (VolumePerUnitVolume_CYT * vcRegionVolume('IC')) |
| Size_EC | (VolumePerUnitVolume_EC * vcRegionVolume('EC')) |
| Size_NM | (AreaPerUnitVolume_NM * vcRegionVolume('IC')) |
| Size_NUC | (VolumePerUnitVolume_NUC * vcRegionVolume('IC')) |
| Size_PM | (AreaPerUnitArea_PM * vcRegionArea('EC_IC_membrane')) |
| sobj_IC1_EC0_size | vcRegionArea('EC_IC_membrane') |
| vobj_EC0_size | vcRegionVolume('EC') |
| vobj_IC1_size | vcRegionVolume('IC') |

##### 2.2.5.3. Volume Domains

#### 2.2.5.3.1. IC

#### 2.2.5.3.2. EC

##### 2.2.5.4. Membrane Domains

###### 2.2.5.4.1. EC_IC_membrane

| PdeEquation Fluor | |
|---|---|
| Rate | - J_r0 |
| Diffusion | Fluor_diffusionRate |
| Initial | Fluor_init_molecules_um_2 |

| PdeEquation Dark | |
|---|---|
| Rate | J_r0 |
| Diffusion | Dark_diffusionRate |
| Initial | Dark_init_molecules_um_2 |

| OdeEquation Laser | |
|---|---|
| Rate | 0.0 |
| Initial | Laser_init_molecules_um_2 |

#### 2.2.6. Simulation(s)

##### 2.2.6.1. Vary D and bleach duration

| Simulation Name: Vary D and bleach duration |
|---|

| Overriden Parameters | | |
|---|---|---|
| **Name** | **Actual Value** | **Default Value** |
| Fluor_diffusionRate | 0.1 to 1.0, 2 values | 0.1 |
| duration | "0.3", "1.0", "3.0" | 1.0 |
| Dark_diffusionRate | Fluor_diffusionRate | 0.1 |

| Geometry Setting | |
|---|---|
| Geometry Size (um) | (22.0, 22.0, 11.0) |
| Mesh Size (elements) | (51, 51, 26) |

| Advanced Settings | |
|---|---|
| Solver Name | Fully-Implicit Finite Volume, Regular Grid (Variable Time Step) |

| | |
|---|---|
| Time Bounds - Starting | 0.0 |
| Time Bounds - Ending | 20.0 |
| Time Step - Min | 0.0 |
| Time Step - Default | 0.05 |
| Time Step - Max | 0.1 |
| Error Tolerance - Absolute | 1.0E-9 |
| Error Tolerance - Relative | 1.0E-7 |
| Output Time Step | 0.5 |
| Use Symbolic Jacobian (T/F) | F |

##### 2.2.6.2. widefield axial laser D and bleach duration

| **Simulation Name: widefield axial laser D and bleach duration** |
|---|

| **Overriden Parameters** | | |
|---|---|---|
| **Name** | **Actual Value** | **Default Value** |
| Fluor_diffusionRate | 0.1 to 1.0, 2 values | 0.1 |
| sigmaaxial | 30.0 | 1.5 |
| duration | "0.3", "1.0", "3.0" | 1.0 |
| Dark_diffusionRate | Fluor_diffusionRate | 0.1 |

| **Geometry Setting** | |
|---|---|
| Geometry Size (um) | (22.0, 22.0, 11.0) |
| Mesh Size (elements) | (51, 51, 26) |

| **Advanced Settings** | |
|---|---|
| Solver Name | Fully-Implicit Finite Volume, Regular Grid (Variable Time Step) |

| | |
|---|---|
| Time Bounds - Starting | 0.0 |
| Time Bounds - Ending | 20.0 |
| Time Step - Min | 0.0 |
| Time Step - Default | 0.05 |
| Time Step - Max | 0.1 |
| Error Tolerance - Absolute | 1.0E-9 |
| Error Tolerance - Relative | 1.0E-7 |
| Output Time Step | 0.5 |
| Use Symbolic Jacobian (T/F) | F |

#### 2.3. Application: TIRF_off_center_bleach

| **Application Name: TIRF_off_center_bleach** |
|---|
| **Application Description: (copied from TIRF_center_bleach) (copied from Spherical_Cell_Gaussian_Bleach)** |

##### 2.3.1. Structure Mapping For TIRF_off_center_bleach

| **Structure Mapping** | | | | |
|---|---|---|---|---|
| **Structure** | **Subdomain** | **Resolved (T/F)** | **Surf/Vol** | **VolFract** |
| NUC | IC | F | | |
| CYT | IC | F | | |
| EC | EC | F | | |

##### 2.3.2. Reaction Mapping For TIRF_off_center_bleach

| **Reaction Mapping** | | | |
|---|---|---|---|
| **Name** | **Type** | **Enabled (T/F)** | **Fast (T/F)** |
| r0 | Reaction | T | F |

| **Initial Conditions** |
|---|

| Initial Conditions | | | | |
|---|---|---|---|---|
| **Species** | **Structure** | **Initial Conc.** | **Diffusion Const.** | **Fixed (T/F)** |
| s0 | PM | 10.0 molecules.m² | 0.1 m².s¹ | F |
| s1 | PM | 0.0 molecules.m² | 0.1 m².s¹ | F |
| s2 | PM | exp( - ((((x + 9.0) ^ 2.0) / (2.0 * (sigmalateral ^ 2.0))) + ((y ^ 2.0) / (2.0 * (sigmalateral ^ 2.0))) + ((z ^ 2.0) / (2.0 * (sigmaaxial ^ 2.0))))) molecules.m² | 0.0 m².s¹ | F |

#### 2.3.3. Membrane Mapping For TIRF_off_center_bleach

| Electrical Mapping - Membrane Potential | | | |
|---|---|---|---|
| **Membrane** | **Calculate V (T/F)** | **V initial** | **Specific Capacitance** |
| PM | F | 0.0 mV | 1.0 pF.m² |
| NM | F | 0.0 mV | 1.0 pF.m² |

| |
|---|
| Temperature: 300.0 K |

#### 2.3.4. Geometry: Geometry7

| | |
|---|---|
| Size | (22.0, 22.0, 11.0) |
| Origin | (-11.0, -11.0, -0.5) |

#### 2.3.5. Math Description: Copy of TIRF_center_bleach_generated

##### 2.3.5.1. Constants

| Constant Name | Expression |
|---|---|
| _F_ | 96485.3321 |

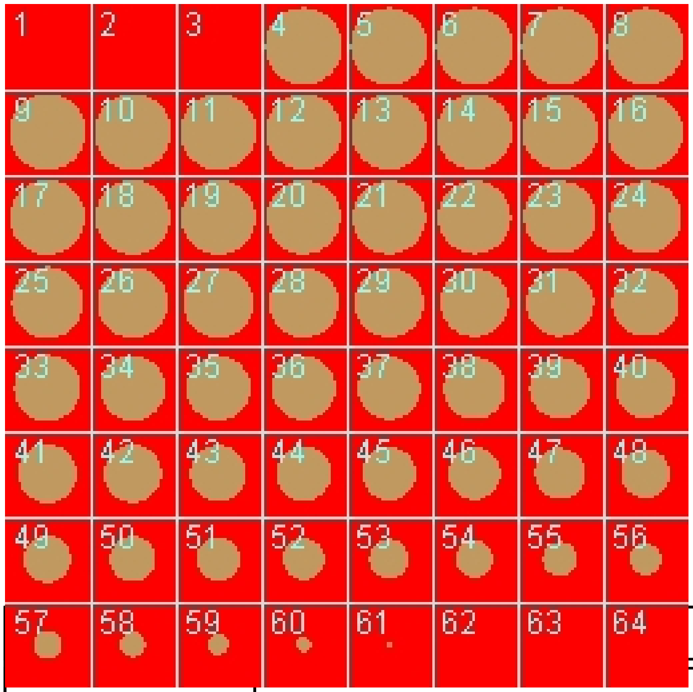

| Constant Name | Expression |
|---|---|
| _F_nmol_ | 9.64853321E-5 |
| _K_GHK_ | 1.0E-9 |
| _N_pmol_ | 6.02214179E11 |
| _PI_ | 3.141592653589793 |
| _R_ | 8314.46261815 |
| _T_ | 300.0 |
| AreaPerUnitArea_PM | 1.0 |
| AreaPerUnitVolume_NM | 1.2 |
| BleachRadius | 2.0 |
| Dark_diffusionRate | 0.1 |
| Dark_init_molecules_um_2 | 0.0 |
| duration | 1.0 |
| Fluor_diffusionRate | 0.1 |
| Fluor_init_molecules_um_2 | 10.0 |

| Constant Name | Expression |
| --- | --- |
| K_millivolts_per_volt | 1000.0 |
| kbleach | 10.0 |
| KMOLE | 0.001660538783162726 |
| Kr | 0.0 |
| netValence | 1.0 |
| sigmaaxial | 1.5 |
| sigmalateral | 0.5 |
| start | 1.0 |
| Voltage_NM | 0.0 |
| Voltage_PM | 0.0 |
| VolumePerUnitVolume_CYT | 1.0 |
| VolumePerUnitVolume_EC | 1.0 |
| VolumePerUnitVolume_NUC | 65.45 |

#### 2.3.5.2. Functions

| Function Name | Expression |
| --- | --- |
| J_r0 | ((Kf * Fluor) - (Kr * Dark)) |
| Kf | ((kbleach * Laser) * (t > start) * (t < (start + duration))) |
| Laser_init_molecules_um_2 | exp( - ((((x + 9.0) ^ 2.0) / (2.0 * (sigmalateral ^ 2.0))) + ((y ^ 2.0) / (2.0 * (sigmalateral ^ 2.0))) + ((z ^ 2.0) / (2.0 * (sigmaaxial ^ 2.0))))) |
| Size_CYT | (VolumePerUnitVolume_CYT * vcRegionVolume('IC')) |
| Size_EC | (VolumePerUnitVolume_EC * vcRegionVolume('EC')) |
| Size_NM | (AreaPerUnitVolume_NM * vcRegionVolume('IC')) |
| Size_NUC | (VolumePerUnitVolume_NUC * vcRegionVolume('IC')) |
| Size_PM | (AreaPerUnitArea_PM * vcRegionArea('EC_IC_membrane')) |
| sobj_IC1_EC0_size | vcRegionArea('EC_IC_membrane') |

| Function Name | Expression |
|---|---|
| vobj_EC0_size | vcRegionVolume('EC') |
| vobj_IC1_size | vcRegionVolume('IC') |

#### 2.3.5.3. Volume Domains

### 2.3.5.3.1. IC

### 2.3.5.3.2. EC

#### 2.3.5.4. Membrane Domains

##### 2.3.5.4.1. EC_IC_membrane

| PdeEquation Fluor | |
|---|---|
| Rate | - J_r0 |
| Diffusion | Fluor_diffusionRate |
| Initial | Fluor_init_molecules_um_2 |

| PdeEquation Dark | |
|---|---|
| Rate | J_r0 |
| Diffusion | Dark_diffusionRate |
| Initial | Dark_init_molecules_um_2 |

| OdeEquation Laser | |
|---|---|
| Rate | 0.0 |
| Initial | Laser_init_molecules_um_2 |

#### 2.3.6. Simulation(s)

##### 2.3.6.1. Vary D and bleach duration_1

| Simulation Name: Vary D and bleach duration_1 |
|---|

| Overriden Parameters | | |
|---|---|---|
| **Name** | **Actual Value** | **Default Value** |
| Fluor_diffusionRate | 0.1 to 1.0, 2 values | 0.1 |
| duration | "0.3", "1.0", "3.0" | 1.0 |
| Dark_diffusionRate | Fluor_diffusionRate | 0.1 |

| Geometry Setting | |
|---|---|
| Geometry Size (um) | (22.0, 22.0, 11.0) |
| Mesh Size (elements) | (51, 51, 26) |

| Advanced Settings | |
|---|---|
| Solver Name | Fully-Implicit Finite Volume, Regular Grid (Variable Time Step) |
| Time Bounds - Starting | 0.0 |
| Time Bounds - Ending | 20.0 |
| Time Step - Min | 0.0 |
| Time Step - Default | 0.05 |
| Time Step - Max | 0.1 |
| Error Tolerance - Absolute | 1.0E-9 |
| Error Tolerance - Relative | 1.0E-7 |
| Output Time Step | 0.5 |
| Use Symbolic Jacobian (T/F) | F |

#### 2.3.6.2. Widefield axial Vary D and bleach duration_1

**Simulation Name: Widefield axial Vary D and bleach duration_1**

| Overriden Parameters | | |
|---|---|---|
| **Name** | **Actual Value** | **Default Value** |
| Fluor_diffusionRate | 0.1 to 1.0, 2 values | 0.1 |
| sigmaaxial | 30.0 | 1.5 |
| duration | "0.3", "1.0", "3.0" | 1.0 |
| Dark_diffusionRate | Fluor_diffusionRate | 0.1 |

| Geometry Setting | |
|---|---|
| Geometry Size (um) | (22.0, 22.0, 11.0) |
| Mesh Size (elements) | (101, 101, 51) |

| Advanced Settings | |
|---|---|
| Solver Name | Fully-Implicit Finite Volume, Regular Grid (Variable Time Step) |
| Time Bounds - Starting | 0.0 |
| Time Bounds - Ending | 20.0 |
| Time Step - Min | 0.0 |
| Time Step - Default | 0.05 |
| Time Step - Max | 0.1 |
| Error Tolerance - Absolute | 1.0E-9 |
| Error Tolerance - Relative | 1.0E-7 |
| Output Time Step | 0.5 |
| Use Symbolic Jacobian (T/F) | F |

#### 2.4. Application: Image-based center bleach

| **Application Name: Image-based center bleach** |
|---|
| **Application Description: (copied from Spherical_Cell_Gaussian_Bleach)** |

##### 2.4.1. Structure Mapping For Image-based center bleach

| **Structure Mapping** | | | | |
|---|---|---|---|---|
| **Structure** | **Subdomain** | **Resolved (T/F)** | **Surf/Vol** | **VolFract** |
| NUC | Nucleus | F | | |
| CYT | cytosol | F | | |
| EC | ec | F | | |

##### 2.4.2. Reaction Mapping For Image-based center bleach

| **Reaction Mapping** | | | |
|---|---|---|---|
| **Name** | **Type** | **Enabled (T/F)** | **Fast (T/F)** |
| r0 | Reaction | T | F |

| **Initial Conditions** | | | | |
|---|---|---|---|---|
| **Species** | **Structure** | **Initial Conc.** | **Diffusion Const.** | **Fixed (T/F)** |
| s0 | PM | 10.0 molecules.$m^2$ | 0.1 $m^2.s^1$ | F |
| s1 | PM | 0.0 molecules.$m^2$ | 0.1 $m^2.s^1$ | F |
| s2 | PM | exp( - ((((x - 29.8) ^ 2.0) / (2.0 * (sigmalateral ^ 2.0))) + (((y - 40.2) ^ 2.0) / (2.0 * (sigmalateral ^ 2.0))) + (((z - 25.9) ^ 2.0) / (2.0 * (sigmaaxial ^ 2.0))))) molecules.$m^2$ | 0.0 $m^2.s^1$ | F |

#### 2.4.3. Membrane Mapping For Image-based center bleach

| Electrical Mapping - Membrane Potential | | | |
|---|---|---|---|
| **Membrane** | **Calculate V (T/F)** | **V initial** | **Specific Capacitance** |
| PM | F | 0.0 mV | 1.0 pF.m² |
| NM | F | 0.0 mV | 1.0 pF.m² |

| |
|---|
| Temperature: 300.0 K |

#### 2.4.4. Geometry: Site visit _Application0_20111127_695607844

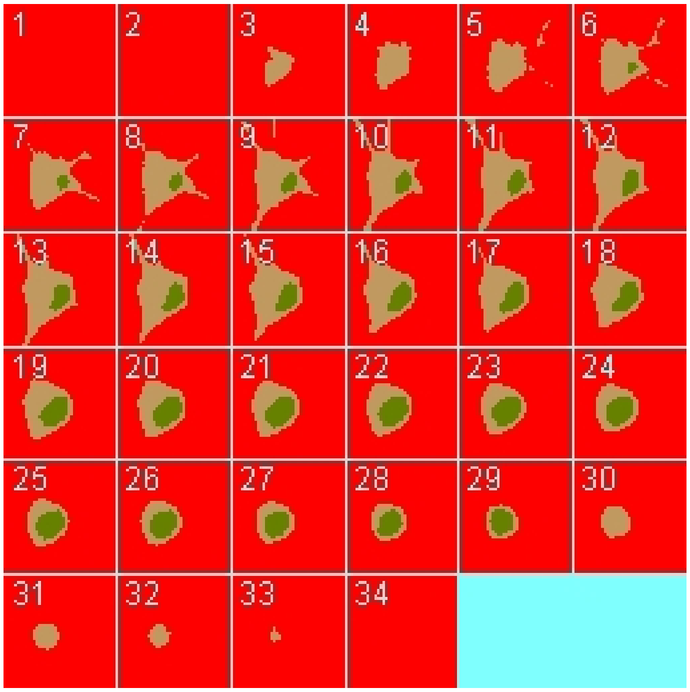

| | |
|---|---|
| Size | (74.24, 74.24, 26.0) |
| Origin | (0.0, 0.0, 0.0) |

#### 2.4.5. Math Description: Copy of Spherical_Cell_Gaussian_Bleach_generated

#### 2.4.5.1. Constants

| Constant Name | Expression |
|---|---|
| _F_ | 96485.3321 |
| _F_nmol_ | 9.64853321E-5 |
| _K_GHK_ | 1.0E-9 |
| _N_pmol_ | 6.02214179E11 |
| _PI_ | 3.141592653589793 |
| _R_ | 8314.46261815 |
| _T_ | 300.0 |
| AreaPerUnitArea_NM | 1.0 |
| AreaPerUnitArea_PM | 1.0 |
| BleachRadius | 2.0 |
| Dark_diffusionRate | 0.1 |
| Dark_init_molecules_um_2 | 0.0 |
| duration | 1.0 |
| Fluor_diffusionRate | 0.1 |
| Fluor_init_molecules_um_2 | 10.0 |
| K_millivolts_per_volt | 1000.0 |
| kbleach | 10.0 |
| KMOLE | 0.001660538783162726 |
| Kr | 0.0 |
| netValence | 1.0 |
| sigmaaxial | 1.5 |
| sigmalateral | 0.5 |
| start | 1.0 |
| Voltage_NM | 0.0 |
| Voltage_PM | 0.0 |

| Constant Name | Expression |
|---|---|
| VolumePerUnitVolume_CYT | 1.0 |
| VolumePerUnitVolume_EC | 1.0 |
| VolumePerUnitVolume_NUC | 1.0 |

#### 2.4.5.2. Functions

| Function Name | Expression |
|---|---|
| J_r0 | ((Kf * Fluor) - (Kr * Dark)) |
| Kf | ((kbleach * Laser) * (t > start) * (t < (start + duration))) |
| Laser_init_molecules_um_2 | exp( - ((((x - 29.8) ^ 2.0) / (2.0 * (sigmalateral ^ 2.0))) + (((y - 40.2) ^ 2.0) / (2.0 * (sigmalateral ^ 2.0))) + (((z - 25.9) ^ 2.0) / (2.0 * (sigmaaxial ^ 2.0))))) |
| Size_CYT | (VolumePerUnitVolume_CYT * vcRegionVolume('cytosol')) |
| Size_EC | (VolumePerUnitVolume_EC * vcRegionVolume('ec')) |
| Size_NM | (AreaPerUnitArea_NM * vcRegionArea('Nucleus_cytosol_membrane')) |
| Size_NUC | (VolumePerUnitVolume_NUC * vcRegionVolume('Nucleus')) |
| Size_PM | (AreaPerUnitArea_PM * vcRegionArea('cytosol_ec_membrane')) |
| sobj_cytosol1_ec0_size | vcRegionArea('cytosol_ec_membrane') |
| sobj_Nucleus2_cytosol1_size | vcRegionArea('Nucleus_cytosol_membrane') |
| vobj_cytosol1_size | vcRegionVolume('cytosol') |
| vobj_ec0_size | vcRegionVolume('ec') |
| vobj_Nucleus2_size | vcRegionVolume('Nucleus') |

#### 2.4.5.3. Volume Domains

### 2.4.5.3.1. ec

###### 2.4.5.3.2. cytosol

###### 2.4.5.3.3. Nucleus

##### 2.4.5.4. Membrane Domains

###### 2.4.5.4.1. cytosol_ec_membrane

| PdeEquation Fluor | |
|---|---|
| Rate | - J_r0 |
| Diffusion | Fluor_diffusionRate |
| Initial | Fluor_init_molecules_um_2 |

| PdeEquation Dark | |
|---|---|
| Rate | J_r0 |
| Diffusion | Dark_diffusionRate |
| Initial | Dark_init_molecules_um_2 |

| OdeEquation Laser | |
|---|---|
| Rate | 0.0 |
| Initial | Laser_init_molecules_um_2 |

###### 2.4.5.4.2. Nucleus_cytosol_membrane

##### 2.4.6. Simulation(s)

###### 2.4.6.1. Vary D and bleach Duration_1

| Simulation Name: Vary D and bleach Duration_1 |
|---|

| Overriden Parameters |
|---|

| Name | Actual Value | Default Value |
|---|---|---|
| Fluor_diffusionRate | 0.1 to 1.0, 2 values | 0.1 |
| duration | "0.3", "1.0", "3.0" | 1.0 |
| Dark_diffusionRate | Fluor_diffusionRate | 0.1 |

| Geometry Setting | |
|---|---|
| Geometry Size (um) | (74.24, 74.24, 26.0) |
| Mesh Size (elements) | (201, 201, 71) |

| Advanced Settings | |
|---|---|
| Solver Name | Fully-Implicit Finite Volume, Regular Grid (Variable Time Step) |
| Time Bounds - Starting | 0.0 |
| Time Bounds - Ending | 20.0 |
| Time Step - Min | 0.0 |
| Time Step - Default | 0.05 |
| Time Step - Max | 0.1 |
| Error Tolerance - Absolute | 1.0E-9 |
| Error Tolerance - Relative | 1.0E-7 |
| Output Time Step | 0.5 |
| Use Symbolic Jacobian (T/F) | F |

#### 2.4.6.2. Widefield axial Vary D and bleach Duration_1

| Simulation Name: Widefield axial Vary D and bleach Duration_1 |
|---|

| Overriden Parameters | | |
|---|---|---|
| **Name** | **Actual Value** | **Default Value** |

| | | |
|---|---|---|
| Fluor_diffusionRate | 0.1 to 1.0, 2 values | 0.1 |
| sigmaaxial | 30.0 | 1.5 |
| duration | "0.1", "3.0", "1.0" | 1.0 |
| Dark_diffusionRate | Fluor_diffusionRate | 0.1 |

| Geometry Setting | |
|---|---|
| Geometry Size (um) | (74.24, 74.24, 26.0) |
| Mesh Size (elements) | (201, 201, 71) |

| Advanced Settings | |
|---|---|
| Solver Name | Fully-Implicit Finite Volume, Regular Grid (Variable Time Step) |
| Time Bounds - Starting | 0.0 |
| Time Bounds - Ending | 20.0 |
| Time Step - Min | 0.0 |
| Time Step - Default | 0.05 |
| Time Step - Max | 0.1 |
| Error Tolerance - Absolute | 1.0E-9 |
| Error Tolerance - Relative | 1.0E-7 |
| Output Time Step | 0.5 |
| Use Symbolic Jacobian (T/F) | F |

##### 2.4.6.3. Copy of Widefield axial Vary D and bleach Duration_1

| **Simulation Name: Copy of Widefield axial Vary D and bleach Duration_1** |
|---|
| **Simulation Description: cloned from 'Copy of Widefield axial Vary D and bleach Duration_1' owned by user temp** |

| Overriden Parameters | | |
|---|---|---|
| **Name** | **Actual Value** | **Default Value** |
| Fluor_diffusionRate | 2.0 | 0.1 |
| sigmalateral | 1.5 | 0.5 |
| sigmaaxial | 30.0 | 1.5 |
| Dark_diffusionRate | Fluor_diffusionRate | 0.1 |

| Geometry Setting | |
|---|---|
| Geometry Size (um) | (74.24, 74.24, 26.0) |
| Mesh Size (elements) | (401, 401, 141) |

| Advanced Settings | |
|---|---|
| Solver Name | Fully-Implicit Finite Volume, Regular Grid (Variable Time Step) |
| Time Bounds - Starting | 0.0 |
| Time Bounds - Ending | 20.0 |
| Time Step - Min | 0.0 |
| Time Step - Default | 0.05 |
| Time Step - Max | 0.1 |
| Error Tolerance - Absolute | 1.0E-9 |
| Error Tolerance - Relative | 1.0E-7 |
| Output Time Step | 1.0 |
| Use Symbolic Jacobian (T/F) | F |

#### 2.4.6.4. Copy of Copy of Widefield axial Vary D and bleach Duration_1

| Simulation Name: Copy of Copy of Widefield axial Vary D and bleach Duration_1 |
|---|

**Simulation Description: cloned from 'Copy of Copy of Widefield axial Vary D and bleach Duration_1' owned by user temp**
**cloned from 'Copy of Widefield axial Vary D and bleach Duration_1' owned by user temp**

| Overriden Parameters | | |
|---|---|---|
| **Name** | **Actual Value** | **Default Value** |
| Fluor_diffusionRate | 2.0 | 0.1 |
| sigmalateral | 1.5 | 0.5 |
| sigmaaxial | 30.0 | 1.5 |
| Dark_diffusionRate | Fluor_diffusionRate | 0.1 |

| Geometry Setting | |
|---|---|
| Geometry Size (um) | (74.24, 74.24, 26.0) |
| Mesh Size (elements) | (801, 801, 281) |

| Advanced Settings | |
|---|---|
| Solver Name | Fully-Implicit Finite Volume, Regular Grid (Variable Time Step) |
| Time Bounds - Starting | 0.0 |
| Time Bounds - Ending | 20.0 |
| Time Step - Min | 0.0 |
| Time Step - Default | 0.05 |
| Time Step - Max | 0.1 |
| Error Tolerance - Absolute | 1.0E-9 |
| Error Tolerance - Relative | 1.0E-7 |
| Output Time Step | 1.0 |
| Use Symbolic Jacobian (T/F) | F |

#### 2.5. Application: Image-based process bleach

| **Application Name: Image-based process bleach** |
| --- |
| **Application Description: (copied from Image-based center bleach) (copied from Spherical_Cell_Gaussian_Bleach)** |

##### 2.5.1. Structure Mapping For Image-based process bleach

| **Structure Mapping** | | | | |
| --- | --- | --- | --- | --- |
| **Structure** | **Subdomain** | **Resolved (T/F)** | **Surf/Vol** | **VolFract** |
| NUC | Nucleus | F | | |
| CYT | cytosol | F | | |
| EC | ec | F | | |

##### 2.5.2. Reaction Mapping For Image-based process bleach

| **Reaction Mapping** | | | |
| --- | --- | --- | --- |
| **Name** | **Type** | **Enabled (T/F)** | **Fast (T/F)** |
| r0 | Reaction | T | F |

| **Initial Conditions** | | | | |
| --- | --- | --- | --- | --- |
| **Species** | **Structure** | **Initial Conc.** | **Diffusion Const.** | **Fixed (T/F)** |
| s0 | PM | 10.0 molecules.$m^2$ | 0.1 $m^2.s^1$ | F |
| s1 | PM | 0.0 molecules.$m^2$ | 0.1 $m^2.s^1$ | F |
| s2 | PM | exp( - ((((x - 28.5) ^ 2.0) / (2.0 * (sigmalateral ^ 2.0))) + (((y - 7.0) ^ 2.0) / (2.0 * (sigmalateral ^ 2.0))) + (((z - 8.0) ^ 2.0) / (2.0 * (sigmaaxial ^ 2.0))))) molecules.$m^2$ | 0.0 $m^2.s^1$ | F |

#### 2.5.3. Membrane Mapping For Image-based process bleach

| Electrical Mapping - Membrane Potential | | | |
|---|---|---|---|
| **Membrane** | **Calculate V (T/F)** | **V initial** | **Specific Capacitance** |
| PM | F | 0.0 mV | 1.0 pF.m² |
| NM | F | 0.0 mV | 1.0 pF.m² |

| |
|---|
| Temperature: 300.0 K |

#### 2.5.4. Geometry: Site visit _Application0_20111127_695607844

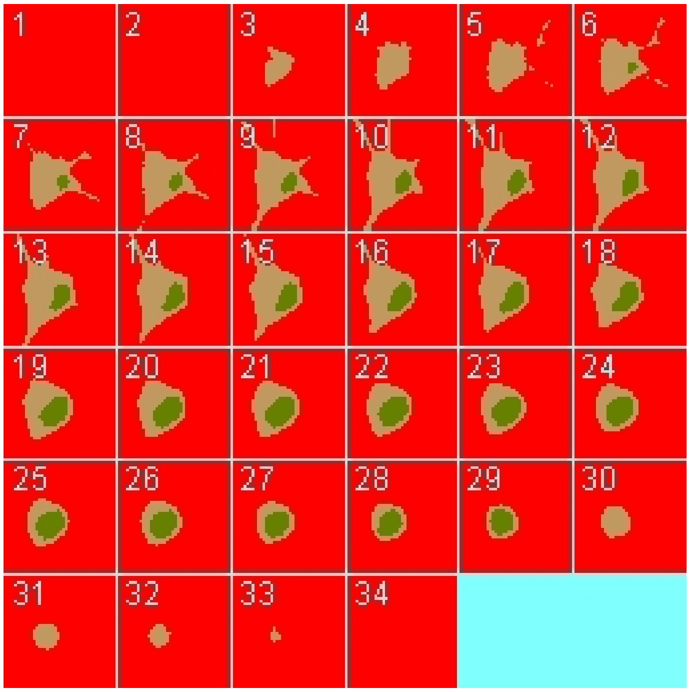

| | |
|---|---|
| Size | (74.24, 74.24, 26.0) |
| Origin | (0.0, 0.0, 0.0) |

#### 2.5.5. Math Description: Copy of Image-based center bleach_generated

##### 2.5.5.1. Constants

| Constant Name | Expression |
|---|---|
| _F_ | 96485.3321 |
| _F_nmol_ | 9.64853321E-5 |
| _K_GHK_ | 1.0E-9 |
| _N_pmol_ | 6.02214179E11 |
| _PI_ | 3.141592653589793 |
| _R_ | 8314.46261815 |
| _T_ | 300.0 |
| AreaPerUnitArea_NM | 1.0 |
| AreaPerUnitArea_PM | 1.0 |
| BleachRadius | 2.0 |
| Dark_diffusionRate | 0.1 |
| Dark_init_molecules_um_2 | 0.0 |
| duration | 1.0 |
| Fluor_diffusionRate | 0.1 |
| Fluor_init_molecules_um_2 | 10.0 |
| K_millivolts_per_volt | 1000.0 |
| kbleach | 10.0 |
| KMOLE | 0.001660538783162726 |
| Kr | 0.0 |
| netValence | 1.0 |
| sigmaaxial | 1.5 |
| sigmalateral | 0.5 |
| start | 1.0 |
| Voltage_NM | 0.0 |
| Voltage_PM | 0.0 |

| Constant Name | Expression |
|---|---|
| VolumePerUnitVolume_CYT | 1.0 |
| VolumePerUnitVolume_EC | 1.0 |
| VolumePerUnitVolume_NUC | 1.0 |

#### 2.5.5.2. Functions

| Function Name | Expression |
|---|---|
| J_r0 | ((Kf * Fluor) - (Kr * Dark)) |
| Kf | ((kbleach * Laser) * (t > start) * (t < (start + duration))) |
| Laser_init_molecules_um_2 | exp( - ((((x - 28.5) ^ 2.0) / (2.0 * (sigmalateral ^ 2.0))) + (((y - 7.0) ^ 2.0) / (2.0 * (sigmalateral ^ 2.0))) + (((z - 8.0) ^ 2.0) / (2.0 * (sigmaaxial ^ 2.0))))) |
| Size_CYT | (VolumePerUnitVolume_CYT * vcRegionVolume('cytosol')) |
| Size_EC | (VolumePerUnitVolume_EC * vcRegionVolume('ec')) |
| Size_NM | (AreaPerUnitArea_NM * vcRegionArea('Nucleus_cytosol_membrane')) |
| Size_NUC | (VolumePerUnitVolume_NUC * vcRegionVolume('Nucleus')) |
| Size_PM | (AreaPerUnitArea_PM * vcRegionArea('cytosol_ec_membrane')) |
| sobj_cytosol1_ec0_size | vcRegionArea('cytosol_ec_membrane') |
| sobj_Nucleus2_cytosol1_size | vcRegionArea('Nucleus_cytosol_membrane') |
| vobj_cytosol1_size | vcRegionVolume('cytosol') |
| vobj_ec0_size | vcRegionVolume('ec') |
| vobj_Nucleus2_size | vcRegionVolume('Nucleus') |

#### 2.5.5.3. Volume Domains

### 2.5.5.3.1. ec

#### 2.5.5.3.2. cytosol

#### 2.5.5.3.3. Nucleus

#### 2.5.5.4. Membrane Domains

#### 2.5.5.4.1. cytosol_ec_membrane

| PdeEquation Fluor | |
|---|---|
| Rate | - J_r0 |
| Diffusion | Fluor_diffusionRate |
| Initial | Fluor_init_molecules_um_2 |

| PdeEquation Dark | |
|---|---|
| Rate | J_r0 |
| Diffusion | Dark_diffusionRate |
| Initial | Dark_init_molecules_um_2 |

| OdeEquation Laser | |
|---|---|
| Rate | 0.0 |
| Initial | Laser_init_molecules_um_2 |

#### 2.5.5.4.2. Nucleus_cytosol_membrane

#### 2.5.6. Simulation(s)

#### 2.5.6.1. Vary D and bleach Duration_1_1

| Simulation Name: Vary D and bleach Duration_1_1 |
|---|

| Overriden Parameters |
|---|

| Name | Actual Value | Default Value |
|---|---|---|
| Fluor_diffusionRate | 0.1 to 1.0, 2 values | 0.1 |
| duration | "0.3", "1.0", "3.0" | 1.0 |
| Dark_diffusionRate | Fluor_diffusionRate | 0.1 |

| Geometry Setting | |
|---|---|
| Geometry Size (um) | (74.24, 74.24, 26.0) |
| Mesh Size (elements) | (201, 201, 71) |

| Advanced Settings | |
|---|---|
| Solver Name | Fully-Implicit Finite Volume, Regular Grid (Variable Time Step) |
| Time Bounds - Starting | 0.0 |
| Time Bounds - Ending | 20.0 |
| Time Step - Min | 0.0 |
| Time Step - Default | 0.05 |
| Time Step - Max | 0.1 |
| Error Tolerance - Absolute | 1.0E-9 |
| Error Tolerance - Relative | 1.0E-7 |
| Output Time Step | 0.5 |
| Use Symbolic Jacobian (T/F) | F |

#### 2.5.6.2. Widefield axial Vary D and bleach Duration_1_1

| Simulation Name: Widefield axial Vary D and bleach Duration_1_1 |
|---|

| Overriden Parameters | | |
|---|---|---|
| **Name** | **Actual Value** | **Default Value** |

| | | |
|---|---|---|
| Fluor_diffusionRate | 0.1 to 1.0, 2 values | 0.1 |
| sigmaaxial | 30.0 | 1.5 |
| duration | "0.1", "3.0", "1.0" | 1.0 |
| Dark_diffusionRate | Fluor_diffusionRate | 0.1 |

| **Geometry Setting** | |
|---|---|
| Geometry Size (um) | (74.24, 74.24, 26.0) |
| Mesh Size (elements) | (201, 201, 71) |

| **Advanced Settings** | |
|---|---|
| Solver Name | Fully-Implicit Finite Volume, Regular Grid (Variable Time Step) |
| Time Bounds - Starting | 0.0 |
| Time Bounds - Ending | 20.0 |
| Time Step - Min | 0.0 |
| Time Step - Default | 0.05 |
| Time Step - Max | 0.1 |
| Error Tolerance - Absolute | 1.0E-9 |
| Error Tolerance - Relative | 1.0E-7 |
| Output Time Step | 0.5 |
| Use Symbolic Jacobian (T/F) | F |

#### 2.6. Application: CircularBleach Center Image based

| **Application Name: CircularBleach Center Image based** |
|---|
| **Application Description: (copied from Image-based center bleach) (copied from Spherical_Cell_Gaussian_Bleach)** |

#### 2.6.1. Structure Mapping For CircularBleach Center Image based

| Structure Mapping | | | | |
|---|---|---|---|---|
| **Structure** | **Subdomain** | **Resolved (T/F)** | **Surf/Vol** | **VolFract** |
| NUC | Nucleus | F | | |
| CYT | cytosol | F | | |
| EC | ec | F | | |

#### 2.6.2. Reaction Mapping For CircularBleach Center Image based

| Reaction Mapping | | | |
|---|---|---|---|
| **Name** | **Type** | **Enabled (T/F)** | **Fast (T/F)** |
| r0 | Reaction | T | F |

| Initial Conditions | | | | |
|---|---|---|---|---|
| **Species** | **Structure** | **Initial Conc.** | **Diffusion Const.** | **Fixed (T/F)** |
| s0 | PM | 10.0 molecules.m² | 0.1 m².s¹ | F |
| s1 | PM | 0.0 molecules.m² | 0.1 m².s¹ | F |
| s2 | PM | (exp( - (((z - 25.9) ^ 2.0) / (2.0 * (sigmaaxial ^ 2.0)))) * ((((x - 29.8) ^ 2.0) + ((y - 40.2) ^ 2.0)) < 4.0)) molecules.m² | 0.0 m².s¹ | F |

#### 2.6.3. Membrane Mapping For CircularBleach Center Image based

| Electrical Mapping - Membrane Potential | | | |
|---|---|---|---|
| **Membrane** | **Calculate V (T/F)** | **V initial** | **Specific Capacitance** |
| PM | F | 0.0 mV | 1.0 pF.m² |
| NM | F | 0.0 mV | 1.0 pF.m² |

| |
|---|
| Temperature: 300.0 K |

#### 2.6.4. Geometry: Site visit _Application0_20111127_695607844

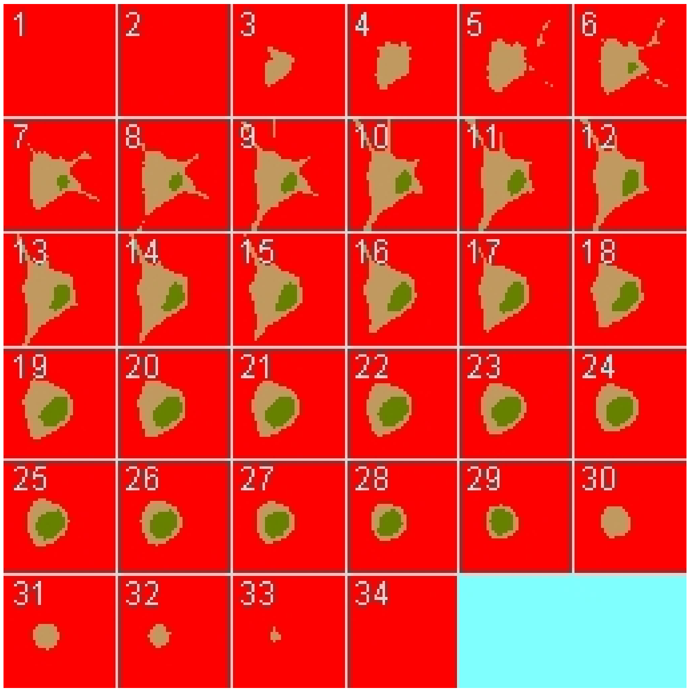

| | |
|---|---|
| Size | (74.24, 74.24, 26.0) |
| Origin | (0.0, 0.0, 0.0) |

#### 2.6.5. Math Description: Copy of Image-based center bleach_generated

##### 2.6.5.1. Constants

| **Constant Name** | **Expression** |
|---|---|
| _F_ | 96485.3321 |
| _F_nmol_ | 9.64853321E-5 |
| _K_GHK_ | 1.0E-9 |
| _N_pmol_ | 6.02214179E11 |
| _PI_ | 3.141592653589793 |

| Constant Name | Expression |
| --- | --- |
| _R_ | 8314.46261815 |
| _T_ | 300.0 |
| AreaPerUnitArea_NM | 1.0 |
| AreaPerUnitArea_PM | 1.0 |
| BleachRadius | 2.0 |
| Dark_diffusionRate | 0.1 |
| Dark_init_molecules_um_2 | 0.0 |
| duration | 1.0 |
| Fluor_diffusionRate | 0.1 |
| Fluor_init_molecules_um_2 | 10.0 |
| K_millivolts_per_volt | 1000.0 |
| kbleach | 10.0 |
| KMOLE | 0.001660538783162726 |
| Kr | 0.0 |
| netValence | 1.0 |
| sigmaaxial | 1.5 |
| sigmalateral | 0.5 |
| start | 1.0 |
| Voltage_NM | 0.0 |
| Voltage_PM | 0.0 |
| VolumePerUnitVolume_CYT | 1.0 |
| VolumePerUnitVolume_EC | 1.0 |
| VolumePerUnitVolume_NUC | 1.0 |

#### 2.6.5.2. Functions

| Function Name | Expression |
|---|---|
| J_r0 | ((Kf * Fluor) - (Kr * Dark)) |
| Kf | ((kbleach * Laser) * (t > start) * (t < (start + duration))) |
| Laser_init_molecules_um_2 | (exp( - (((z - 25.9) ^ 2.0) / (2.0 * (sigmaaxial ^ 2.0)))) * ((((x - 29.8) ^ 2.0) + ((y - 40.2) ^ 2.0)) < 4.0)) |
| Size_CYT | (VolumePerUnitVolume_CYT * vcRegionVolume('cytosol')) |
| Size_EC | (VolumePerUnitVolume_EC * vcRegionVolume('ec')) |
| Size_NM | (AreaPerUnitArea_NM * vcRegionArea('Nucleus_cytosol_membrane')) |
| Size_NUC | (VolumePerUnitVolume_NUC * vcRegionVolume('Nucleus')) |
| Size_PM | (AreaPerUnitArea_PM * vcRegionArea('cytosol_ec_membrane')) |
| sobj_cytosol1_ec0_size | vcRegionArea('cytosol_ec_membrane') |
| sobj_Nucleus2_cytosol1_size | vcRegionArea('Nucleus_cytosol_membrane') |
| vobj_cytosol1_size | vcRegionVolume('cytosol') |
| vobj_ec0_size | vcRegionVolume('ec') |
| vobj_Nucleus2_size | vcRegionVolume('Nucleus') |

#### 2.6.5.3. Volume Domains

### 2.6.5.3.1. ec

##### 2.6.5.3.2. cytosol

##### 2.6.5.3.3. Nucleus

#### 2.6.5.4. Membrane Domains

##### 2.6.5.4.1. cytosol_ec_membrane

| PdeEquation Fluor |
|---|

| PdeEquation Fluor | |
|---|---|
| Rate | - J_r0 |
| Diffusion | Fluor_diffusionRate |
| Initial | Fluor_init_molecules_um_2 |

| PdeEquation Dark | |
|---|---|
| Rate | J_r0 |
| Diffusion | Dark_diffusionRate |
| Initial | Dark_init_molecules_um_2 |

| OdeEquation Laser | |
|---|---|
| Rate | 0.0 |
| Initial | Laser_init_molecules_um_2 |

##### 2.6.5.4.2. Nucleus_cytosol_membrane

#### 2.6.6. Simulation(s)

##### 2.6.6.1. Widefield axial Vary D and bleach Duration_1_2

| |
|---|
| **Simulation Name: Widefield axial Vary D and bleach Duration_1_2** |
| **Simulation Description: cloned from 'Widefield axial Vary D and bleach Duration_1_2' owned by user temp** |

| **Overriden Parameters** | | |
|---|---|---|
| **Name** | **Actual Value** | **Default Value** |
| Fluor_diffusionRate | 0.1 to 1.0, 2 values | 0.1 |
| sigmaaxial | 30.0 | 1.5 |
| duration | "0.1", "3.0", "1.0" | 1.0 |
| Dark_diffusionRate | Fluor_diffusionRate | 0.1 |

| Geometry Setting | |
|---|---|
| Geometry Size (um) | (74.24, 74.24, 26.0) |
| Mesh Size (elements) | (201, 201, 71) |

| Advanced Settings | |
|---|---|
| Solver Name | Fully-Implicit Finite Volume, Regular Grid (Variable Time Step) |
| Time Bounds - Starting | 0.0 |
| Time Bounds - Ending | 20.0 |
| Time Step - Min | 0.0 |
| Time Step - Default | 0.05 |
| Time Step - Max | 0.1 |
| Error Tolerance - Absolute | 1.0E-9 |
| Error Tolerance - Relative | 1.0E-7 |
| Output Time Step | 0.5 |
| Use Symbolic Jacobian (T/F) | F |

##### 2.6.6.2. Copy of Widefield axial Vary D and bleach Duration_1_1

| **Simulation Name: Copy of Widefield axial Vary D and bleach Duration_1_1** |
|---|
| **Simulation Description: cloned from 'Copy of Widefield axial Vary D and bleach Duration_1_1' owned by user temp<br>cloned from 'Copy of Widefield axial Vary D and bleach Duration_1' owned by user temp** |

| Overriden Parameters | | |
|---|---|---|
| **Name** | **Actual Value** | **Default Value** |
| Fluor_diffusionRate | 2.0 | 0.1 |

| | | |
|---|---|---|
| sigmalateral | 1.5 | 0.5 |
| sigmaaxial | 30.0 | 1.5 |
| Dark_diffusionRate | Fluor_diffusionRate | 0.1 |

| **Geometry Setting** | |
|---|---|
| Geometry Size (um) | (74.24, 74.24, 26.0) |
| Mesh Size (elements) | (401, 401, 141) |

| **Advanced Settings** | |
|---|---|
| Solver Name | Fully-Implicit Finite Volume, Regular Grid (Variable Time Step) |
| Time Bounds - Starting | 0.0 |
| Time Bounds - Ending | 20.0 |
| Time Step - Min | 0.0 |
| Time Step - Default | 0.05 |
| Time Step - Max | 0.1 |
| Error Tolerance - Absolute | 1.0E-9 |
| Error Tolerance - Relative | 1.0E-7 |
| Output Time Step | 1.0 |
| Use Symbolic Jacobian (T/F) | F |

#### 2.6.6.3. Copy of Copy of Widefield axial Vary D and bleach Duration_1_1 1

| **Simulation Name: Copy of Copy of Widefield axial Vary D and bleach Duration_1_1 1** |
|---|
| **Simulation Description: cloned from 'Copy of Copy of Widefield axial Vary D and bleach Duration_1_1 1' owned by user temp<br>cloned from 'Copy of Widefield axial Vary D and bleach Duration_1_1' owned by user temp<br>cloned from 'Copy of Widefield axial Vary D and bleach Duration_1' owned by user temp** |

| Overriden Parameters | | |
|---|---|---|
| **Name** | **Actual Value** | **Default Value** |
| Fluor_diffusionRate | 2.0 | 0.1 |
| sigmalateral | 1.5 | 0.5 |
| sigmaaxial | 30.0 | 1.5 |
| Dark_diffusionRate | Fluor_diffusionRate | 0.1 |

| Geometry Setting | |
|---|---|
| Geometry Size (um) | (74.24, 74.24, 26.0) |
| Mesh Size (elements) | (257, 257, 37) |

| Advanced Settings | |
|---|---|
| Solver Name | Fully-Implicit Finite Volume, Regular Grid (Variable Time Step) |
| Time Bounds - Starting | 0.0 |
| Time Bounds - Ending | 20.0 |
| Time Step - Min | 0.0 |
| Time Step - Default | 0.05 |
| Time Step - Max | 0.1 |
| Error Tolerance - Absolute | 1.0E-9 |
| Error Tolerance - Relative | 1.0E-7 |
| Output Time Step | 1.0 |
| Use Symbolic Jacobian (T/F) | F |

#### 2.6.6.4. Copy of Widefield axial Vary D and bleach Duration_1_2

| **Simulation Name: Copy of Widefield axial Vary D and bleach Duration_1_2** |
|---|

**Simulation Description: cloned from 'Copy of Widefield axial Vary D and bleach Duration_1_2' owned by user temp**
**cloned from 'Widefield axial Vary D and bleach Duration_1_2' owned by user temp**

| Overriden Parameters | | |
|---|---|---|
| **Name** | **Actual Value** | **Default Value** |
| Fluor_diffusionRate | 0.1 to 1.0, 2 values | 0.1 |
| sigmaaxial | 30.0 | 1.5 |
| duration | "0.1", "3.0", "1.0" | 1.0 |
| Dark_diffusionRate | Fluor_diffusionRate | 0.1 |

| Geometry Setting | |
|---|---|
| Geometry Size (um) | (74.24, 74.24, 26.0) |
| Mesh Size (elements) | (201, 201, 71) |

| Advanced Settings | |
|---|---|
| Solver Name | Fully-Implicit Finite Volume, Regular Grid (Variable Time Step) |
| Time Bounds - Starting | 0.0 |
| Time Bounds - Ending | 40.0 |
| Time Step - Min | 0.0 |
| Time Step - Default | 0.05 |
| Time Step - Max | 0.1 |
| Error Tolerance - Absolute | 1.0E-9 |
| Error Tolerance - Relative | 1.0E-7 |
| Output Time Step | 0.4 |
| Use Symbolic Jacobian (T/F) | F |

#### 2.7. Application: Spherical_Cell_Circular_Bleach

| Application Name: Spherical_Cell_Circular_Bleach |
|---|
| **Application Description: (copied from Spherical_Cell_Gaussian_Bleach)** |

##### 2.7.1. Structure Mapping For Spherical_Cell_Circular_Bleach

| Structure Mapping | | | | |
|---|---|---|---|---|
| **Structure** | **Subdomain** | **Resolved (T/F)** | **Surf/Vol** | **VolFract** |
| NUC | subdomain1 | F | | |
| CYT | subdomain1 | F | | |
| EC | subdomain0 | F | | |

##### 2.7.2. Reaction Mapping For Spherical_Cell_Circular_Bleach

| Reaction Mapping | | | |
|---|---|---|---|
| **Name** | **Type** | **Enabled (T/F)** | **Fast (T/F)** |
| r0 | Reaction | T | F |

| Initial Conditions | | | | |
|---|---|---|---|---|
| **Species** | **Structure** | **Initial Conc.** | **Diffusion Const.** | **Fixed (T/F)** |
| s0 | PM | 10.0 molecules.$m^2$ | 0.1 $m^2.s^1$ | F |
| s1 | PM | 0.0 molecules.$m^2$ | 0.1 $m^2.s^1$ | F |
| s2 | PM | (exp( - (((z - 1.0) ^ 2.0) / (2.0 * (sigmaaxial ^ 2.0)))) * ((((x - 11.0) ^ 2.0) + ((y - 11.0) ^ 2.0)) < (BleachRadius ^ 2.0))) molecules.$m^2$ | 0.0 $m^2.s^1$ | F |

##### 2.7.3. Membrane Mapping For Spherical_Cell_Circular_Bleach

| Electrical Mapping - Membrane Potential | | | |
|---|---|---|---|
| **Membrane** | **Calculate V (T/F)** | **V initial** | **Specific Capacitance** |
| PM | F | 0.0 mV | 1.0 pF.m² |
| NM | F | 0.0 mV | 1.0 pF.m² |

| |
|---|
| Temperature: 300.0 K |

##### 2.7.4. Geometry: Geometry343793035

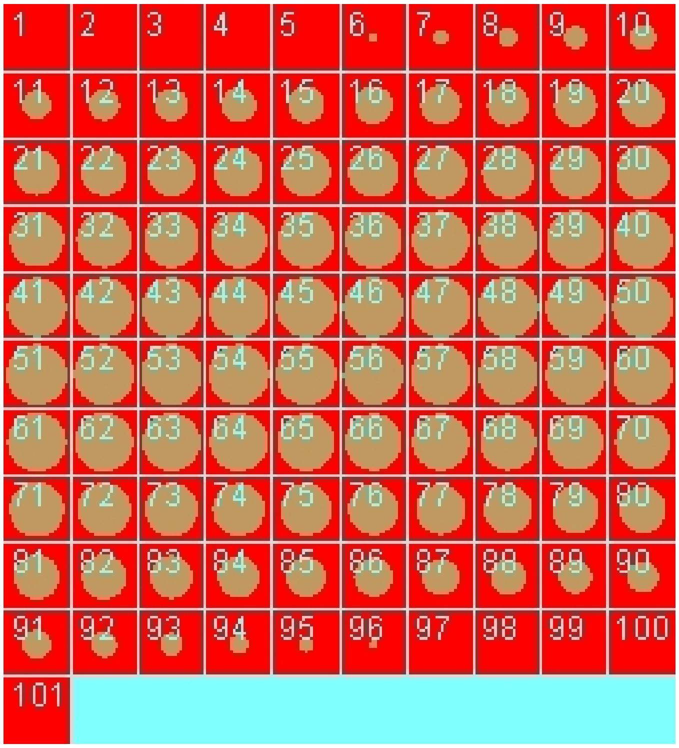

| | |
|---|---|
| Size | (22.0, 22.0, 22.0) |
| Origin | (0.0, 0.0, 0.0) |

##### 2.7.5. Math Description: Copy of Spherical_Cell_Gaussian_Bleach_generated

#### 2.7.5.1. Constants

| Constant Name | Expression |
|---|---|
| _F_ | 96485.3321 |
| _F_nmol_ | 9.64853321E-5 |
| _K_GHK_ | 1.0E-9 |
| _N_pmol_ | 6.02214179E11 |
| _PI_ | 3.141592653589793 |
| _R_ | 8314.46261815 |
| _T_ | 300.0 |
| AreaPerUnitArea_PM | 1.0 |
| AreaPerUnitVolume_NM | 1.2 |
| BleachRadius | 2.0 |
| Dark_diffusionRate | 0.1 |
| Dark_init_molecules_um_2 | 0.0 |
| duration | 1.0 |
| Fluor_diffusionRate | 0.1 |
| Fluor_init_molecules_um_2 | 10.0 |
| K_millivolts_per_volt | 1000.0 |
| kbleach | 10.0 |
| KMOLE | 0.001660538783162726 |
| Kr | 0.0 |
| netValence | 1.0 |
| sigmaaxial | 1.5 |
| sigmalateral | 0.5 |
| start | 1.0 |
| Voltage_NM | 0.0 |

| Constant Name | Expression |
|---|---|
| Voltage_PM | 0.0 |
| VolumePerUnitVolume_CYT | 1.0 |
| VolumePerUnitVolume_EC | 1.0 |
| VolumePerUnitVolume_NUC | 65.45 |

##### 2.7.5.2. Functions

| Function Name | Expression |
|---|---|
| J_r0 | ((Kf * Fluor) - (Kr * Dark)) |
| Kf | ((kbleach * Laser) * (t > start) * (t < (start + duration))) |
| Laser_init_molecules_um_2 | (exp( - (((z - 1.0) ^ 2.0) / (2.0 * (sigmaaxial ^ 2.0)))) * ((((x - 11.0) ^ 2.0) + ((y - 11.0) ^ 2.0)) < (BleachRadius ^ 2.0))) |
| Size_CYT | (VolumePerUnitVolume_CYT * vcRegionVolume('subdomain1')) |
| Size_EC | (VolumePerUnitVolume_EC * vcRegionVolume('subdomain0')) |
| Size_NM | (AreaPerUnitVolume_NM * vcRegionVolume('subdomain1')) |
| Size_NUC | (VolumePerUnitVolume_NUC * vcRegionVolume('subdomain1')) |
| Size_PM | (AreaPerUnitArea_PM * vcRegionArea('subdomain0_subdomain1_membrane')) |
| sobj_subdomain11_subdomain00_size | vcRegionArea('subdomain0_subdomain1_membrane') |
| vobj_subdomain00_size | vcRegionVolume('subdomain0') |
| vobj_subdomain11_size | vcRegionVolume('subdomain1') |

##### 2.7.5.3. Volume Domains

###### 2.7.5.3.1. subdomain1

###### 2.7.5.3.2. subdomain0

#### 2.7.5.4. Membrane Domains

##### 2.7.5.4.1. subdomain0_subdomain1_membrane

| PdeEquation Fluor | |
|---|---|
| Rate | - J_r0 |
| Diffusion | Fluor_diffusionRate |
| Initial | Fluor_init_molecules_um_2 |

| PdeEquation Dark | |
|---|---|
| Rate | J_r0 |
| Diffusion | Dark_diffusionRate |
| Initial | Dark_init_molecules_um_2 |

| OdeEquation Laser | |
|---|---|
| Rate | 0.0 |
| Initial | Laser_init_molecules_um_2 |

#### 2.7.6. Simulation(s)

##### 2.7.6.1. Vary D and bleach Duration_2

| Simulation Name: Vary D and bleach Duration_2 |
|---|
| **Simulation Description: cloned from 'Vary D and bleach Duration_2' owned by user temp** |

| Overriden Parameters | | |
|---|---|---|
| **Name** | **Actual Value** | **Default Value** |
| Fluor_diffusionRate | 0.1 to 1.0, 2 values | 0.1 |
| duration | "0.3", "1.0", "3.0" | 1.0 |
| Dark_diffusionRate | Fluor_diffusionRate | 0.1 |

| Geometry Setting | |
|---|---|
| Geometry Size (um) | (22.0, 22.0, 22.0) |
| Mesh Size (elements) | (51, 51, 51) |

| Advanced Settings | |
|---|---|
| Solver Name | Fully-Implicit Finite Volume, Regular Grid (Variable Time Step) |
| Time Bounds - Starting | 0.0 |
| Time Bounds - Ending | 20.0 |
| Time Step - Min | 0.0 |
| Time Step - Default | 0.05 |
| Time Step - Max | 0.1 |
| Error Tolerance - Absolute | 1.0E-9 |
| Error Tolerance - Relative | 1.0E-7 |
| Output Time Step | 0.5 |
| Use Symbolic Jacobian (T/F) | F |

#### 2.7.6.2. Widefield axial Vary D and bleach Duration_2

| |
|---|
| **Simulation Name: Widefield axial Vary D and bleach Duration_2** |
| **Simulation Description: cloned from 'Widefield axial Vary D and bleach Duration_2' owned by user temp** |

| Overriden Parameters | | |
|---|---|---|
| **Name** | **Actual Value** | **Default Value** |
| Fluor_diffusionRate | 0.1 to 1.0, 2 values | 0.1 |
| sigmaaxial | 30.0 | 1.5 |
| duration | "0.1", "3.0", "1.0" | 1.0 |

| | | |
|---|---|---|
| Dark_diffusionRate | Fluor_diffusionRate | 0.1 |

| Geometry Setting | |
|---|---|
| Geometry Size (um) | (22.0, 22.0, 22.0) |
| Mesh Size (elements) | (51, 51, 51) |

| Advanced Settings | |
|---|---|
| Solver Name | Fully-Implicit Finite Volume, Regular Grid (Variable Time Step) |
| Time Bounds - Starting | 0.0 |
| Time Bounds - Ending | 20.0 |
| Time Step - Min | 0.0 |
| Time Step - Default | 0.05 |
| Time Step - Max | 0.1 |
| Error Tolerance - Absolute | 1.0E-9 |
| Error Tolerance - Relative | 1.0E-7 |
| Output Time Step | 0.5 |
| Use Symbolic Jacobian (T/F) | F |

##### 2.7.6.3. longer Widefield axial Vary D and bleach Duration_2

| **Simulation Name: longer Widefield axial Vary D and bleach Duration_2** |
|---|
| **Simulation Description: cloned from 'Copy of Widefield axial Vary D and bleach Duration_2' owned by user temp<br>cloned from 'Widefield axial Vary D and bleach Duration_2' owned by user temp** |

| **Overriden Parameters** | | |
|---|---|---|
| **Name** | **Actual Value** | **Default Value** |

| | | |
|---|---|---|
| Fluor_diffusionRate | 0.1 to 1.0, 2 values | 0.1 |
| sigmaaxial | 30.0 | 1.5 |
| duration | "0.1", "3.0", "1.0" | 1.0 |
| Dark_diffusionRate | Fluor_diffusionRate | 0.1 |

| **Geometry Setting** | |
|---|---|
| Geometry Size (um) | (22.0, 22.0, 22.0) |
| Mesh Size (elements) | (51, 51, 51) |

| **Advanced Settings** | |
|---|---|
| Solver Name | Fully-Implicit Finite Volume, Regular Grid (Variable Time Step) |
| Time Bounds - Starting | 0.0 |
| Time Bounds - Ending | 50.0 |
| Time Step - Min | 0.0 |
| Time Step - Default | 0.05 |
| Time Step - Max | 0.1 |
| Error Tolerance - Absolute | 1.0E-9 |
| Error Tolerance - Relative | 1.0E-7 |
| Output Time Step | 0.25 |
| Use Symbolic Jacobian (T/F) | F |

##### 2.7.6.4. BleachR= 4 longer Widefield axial Vary D and bleach Duration_2

| **Simulation Name: BleachR= 4 longer Widefield axial Vary D and bleach Duration_2** |
|---|
| **Simulation Description: cloned from 'BleachR= 4 longer Widefield axial Vary D and bleach Duration_2' owned by user temp<br>cloned from 'Copy of Widefield axial Vary D and bleach Duration_2' owned by user temp<br>cloned from 'Widefield axial Vary D and bleach Duration_2' owned by user temp** |

| Overriden Parameters | | |
|---|---|---|
| **Name** | **Actual Value** | **Default Value** |
| BleachRadius | 4.0 | 2.0 |
| Fluor_diffusionRate | 0.1 to 1.0, 2 values | 0.1 |
| sigmaaxial | 30.0 | 1.5 |
| duration | "0.1", "3.0", "1.0" | 1.0 |
| Dark_diffusionRate | Fluor_diffusionRate | 0.1 |

| Geometry Setting | |
|---|---|
| Geometry Size (um) | (22.0, 22.0, 22.0) |
| Mesh Size (elements) | (51, 51, 51) |

| Advanced Settings | |
|---|---|
| Solver Name | Fully-Implicit Finite Volume, Regular Grid (Variable Time Step) |
| Time Bounds - Starting | 0.0 |
| Time Bounds - Ending | 50.0 |
| Time Step - Min | 0.0 |
| Time Step - Default | 0.05 |
| Time Step - Max | 0.1 |
| Error Tolerance - Absolute | 1.0E-9 |
| Error Tolerance - Relative | 1.0E-7 |
| Output Time Step | 0.25 |
| Use Symbolic Jacobian (T/F) | F |

#### 2.7.6.5. Long run for BleachR= 4 longer Widefield axial Vary D and bleach Duration_2

| **Simulation Name: Long run for BleachR= 4 longer Widefield axial Vary D and bleach Duration_2** |
|---|
| **Simulation Description: cloned from 'Ling run for BleachR= 4 longer Widefield axial Vary D and bleach Duration_2' owned by user temp<br>cloned from 'BleachR= 4 longer Widefield axial Vary D and bleach Duration_2' owned by user temp<br>cloned from 'Copy of Widefield axial Vary D and bleach Duration_2' owned by user temp<br>cloned from 'Widefield axial Vary D and bleach Duration_2' owned by user temp** |

| **Overriden Parameters** | | |
|---|---|---|
| **Name** | **Actual Value** | **Default Value** |
| BleachRadius | 4.0 | 2.0 |
| Fluor_diffusionRate | 0.1 to 1.0, 2 values | 0.1 |
| sigmaaxial | 30.0 | 1.5 |
| duration | "0.1", "3.0", "1.0" | 1.0 |
| Dark_diffusionRate | Fluor_diffusionRate | 0.1 |

| **Geometry Setting** | |
|---|---|
| Geometry Size (um) | (22.0, 22.0, 22.0) |
| Mesh Size (elements) | (51, 51, 51) |

| **Advanced Settings** | |
|---|---|
| Solver Name | Fully-Implicit Finite Volume, Regular Grid (Variable Time Step) |
| Time Bounds - Starting | 0.0 |
| Time Bounds - Ending | 200.0 |
| Time Step - Min | 0.0 |
| Time Step - Default | 0.05 |

| | |
|---|---|
| Time Step - Max | 0.1 |
| Error Tolerance - Absolute | 1.0E-9 |
| Error Tolerance - Relative | 1.0E-7 |
| Output Time Step | 0.25 |
| Use Symbolic Jacobian (T/F) | F |

##### 2.7.6.6. BleachR=1 Widefield axial Vary D and bleach Duration long run

| |
|---|
| **Simulation Name: BleachR=1 Widefield axial Vary D and bleach Duration long run** |
| **Simulation Description: cloned from 'BleachR=1 Widefield axial Vary D and bleach Duration long run' owned by user temp<br>cloned from 'Copy of Widefield axial Vary D and bleach Duration_2' owned by user temp<br>cloned from 'Widefield axial Vary D and bleach Duration_2' owned by user temp** |

| **Overriden Parameters** | | |
|---|---|---|
| **Name** | **Actual Value** | **Default Value** |
| BleachRadius | 1.0 | 2.0 |
| Fluor_diffusionRate | 0.1 to 1.0, 2 values | 0.1 |
| sigmaaxial | 30.0 | 1.5 |
| duration | "0.1", "3.0", "1.0" | 1.0 |
| Dark_diffusionRate | Fluor_diffusionRate | 0.1 |

| **Geometry Setting** | |
|---|---|
| Geometry Size (um) | (22.0, 22.0, 22.0) |
| Mesh Size (elements) | (51, 51, 51) |

| **Advanced Settings** | |
|---|---|
| Solver Name | Fully-Implicit Finite Volume, Regular Grid (Variable Time Step) |

| | |
|---|---|
| Time Bounds - Starting | 0.0 |
| Time Bounds - Ending | 200.0 |
| Time Step - Min | 0.0 |
| Time Step - Default | 0.05 |
| Time Step - Max | 0.1 |
| Error Tolerance - Absolute | 1.0E-9 |
| Error Tolerance - Relative | 1.0E-7 |
| Output Time Step | 0.25 |
| Use Symbolic Jacobian (T/F) | F |

#### 2.7.6.7. Finer mesh BleachR=1 Widefield axial Vary D and bleach Duration

| **Simulation Name: Finer mesh BleachR=1 Widefield axial Vary D and bleach Duration** |
|---|
| **Simulation Description: cloned from 'Finer mesh BleachR=1 Widefield axial Vary D and bleach Duration' owned by user temp<br>cloned from 'BleachR=1 Widefield axial Vary D and bleach Duration long run' owned by user temp<br>cloned from 'Copy of Widefield axial Vary D and bleach Duration_2' owned by user temp<br>cloned from 'Widefield axial Vary D and bleach Duration_2' owned by user temp** |

| **Overriden Parameters** | | |
|---|---|---|
| **Name** | **Actual Value** | **Default Value** |
| BleachRadius | 1.0 | 2.0 |
| Fluor_diffusionRate | 0.1 to 1.0, 2 values | 0.1 |
| sigmaaxial | 30.0 | 1.5 |
| duration | "0.1", "3.0", "1.0" | 1.0 |
| Dark_diffusionRate | Fluor_diffusionRate | 0.1 |

| **Geometry Setting** | |
|---|---|
| Geometry Size (um) | (22.0, 22.0, 22.0) |

| | |
|---|---|
| Mesh Size (elements) | (151, 151, 151) |

| Advanced Settings | |
|---|---|
| Solver Name | Fully-Implicit Finite Volume, Regular Grid (Variable Time Step) |
| Time Bounds - Starting | 0.0 |
| Time Bounds - Ending | 100.0 |
| Time Step - Min | 0.0 |
| Time Step - Default | 0.05 |
| Time Step - Max | 0.1 |
| Error Tolerance - Absolute | 1.0E-9 |
| Error Tolerance - Relative | 1.0E-7 |
| Output Time Step | 0.25 |
| Use Symbolic Jacobian (T/F) | F |

##### 2.7.6.8. Copy of Finer mesh BleachR=1 Widefield axial Vary D and bleach Duration

| Simulation Name: Copy of Finer mesh BleachR=1 Widefield axial Vary D and bleach Duration |
|---|
| Simulation Description: cloned from 'Copy of Finer mesh BleachR=1 Widefield axial Vary D and bleach Duration' owned by user temp<br>cloned from 'Finer mesh BleachR=1 Widefield axial Vary D and bleach Duration' owned by user temp<br>cloned from 'BleachR=1 Widefield axial Vary D and bleach Duration long run' owned by user temp<br>cloned from 'Copy of Widefield axial Vary D and bleach Duration_2' owned by user temp<br>cloned from 'Widefield axial Vary D and bleach Duration_2' owned by user temp |

| Overriden Parameters | | |
|---|---|---|
| **Name** | **Actual Value** | **Default Value** |
| BleachRadius | 1.0 | 2.0 |
| Fluor_diffusionRate | 0.1 to 1.0, 2 values | 0.1 |

| | | |
|---|---|---|
| sigmaaxial | 30.0 | 1.5 |
| duration | "0.1", "3.0", "1.0" | 1.0 |
| Dark_diffusionRate | Fluor_diffusionRate | 0.1 |

| Geometry Setting | |
|---|---|
| Geometry Size (um) | (22.0, 22.0, 22.0) |
| Mesh Size (elements) | (151, 151, 151) |

| Advanced Settings | |
|---|---|
| Solver Name | Fully-Implicit Finite Volume, Regular Grid (Variable Time Step) |
| Time Bounds - Starting | 0.0 |
| Time Bounds - Ending | 20.0 |
| Time Step - Min | 0.0 |
| Time Step - Default | 0.05 |
| Time Step - Max | 0.1 |
| Error Tolerance - Absolute | 1.0E-9 |
| Error Tolerance - Relative | 1.0E-7 |
| Output Time Step | 0.1 |
| Use Symbolic Jacobian (T/F) | F |

#### 2.7.6.9. BleachR= 4 longer Widefield axial Vary D and bleach Duration 1s finer mesh

| **Simulation Name: BleachR= 4 longer Widefield axial Vary D and bleach Duration 1s finer mesh** |
|---|
| **Simulation Description: cloned from 'BleachR= 4 longer Widefield axial Vary D and bleach Duration 1s finer mesh' owned by user temp<br>cloned from 'BleachR= 4 longer Widefield axial Vary D and bleach Duration_2' owned by user temp<br>cloned from 'Copy of Widefield axial Vary D and bleach Duration_2' owned by user temp<br>cloned from 'Widefield axial Vary D and bleach Duration_2' owned by user temp** |

| Overriden Parameters | | |
| --- | --- | --- |
| **Name** | **Actual Value** | **Default Value** |
| BleachRadius | 4.0 | 2.0 |
| Fluor_diffusionRate | 0.1 to 1.0, 2 values | 0.1 |
| sigmaaxial | 30.0 | 1.5 |
| Dark_diffusionRate | Fluor_diffusionRate | 0.1 |

| Geometry Setting | |
| --- | --- |
| Geometry Size (um) | (22.0, 22.0, 22.0) |
| Mesh Size (elements) | (151, 151, 151) |

| Advanced Settings | |
| --- | --- |
| Solver Name | Fully-Implicit Finite Volume, Regular Grid (Variable Time Step) |
| Time Bounds - Starting | 0.0 |
| Time Bounds - Ending | 50.0 |
| Time Step - Min | 0.0 |
| Time Step - Default | 0.05 |
| Time Step - Max | 0.1 |
| Error Tolerance - Absolute | 1.0E-9 |
| Error Tolerance - Relative | 1.0E-7 |
| Output Time Step | 0.1 |
| Use Symbolic Jacobian (T/F) | F |

#### 2.8. Application: Image-based process circle bleach

| **Application Name: Image-based process circle bleach** |
| --- |

| Application Description: (copied from Image-based process bleach) (copied from Image-based center bleach) (copied from Spherical_Cell_Gaussian_Bleach) |
|---|

#### 2.8.1. Structure Mapping For Image-based process circle bleach

| Structure Mapping | | | | |
|---|---|---|---|---|
| **Structure** | **Subdomain** | **Resolved (T/F)** | **Surf/Vol** | **VolFract** |
| NUC | Nucleus | F | | |
| CYT | cytosol | F | | |
| EC | ec | F | | |

#### 2.8.2. Reaction Mapping For Image-based process circle bleach

| Reaction Mapping | | | |
|---|---|---|---|
| **Name** | **Type** | **Enabled (T/F)** | **Fast (T/F)** |
| r0 | Reaction | T | F |

| Initial Conditions | | | | |
|---|---|---|---|---|
| **Species** | **Structure** | **Initial Conc.** | **Diffusion Const.** | **Fixed (T/F)** |
| s0 | PM | 10.0 molecules.m² | 0.1 m².s¹ | F |
| s1 | PM | 0.0 molecules.m² | 0.1 m².s¹ | F |
| s2 | PM | (exp( - (((z - 8.0) ^ 2.0) / (2.0 * (sigmaaxial ^ 2.0)))) * ((((x - 28.5) ^ 2.0) + ((y - 7.0) ^ 2.0)) < 4.0)) molecules.m² | 0.0 m².s¹ | F |

#### 2.8.3. Membrane Mapping For Image-based process circle bleach

| Electrical Mapping - Membrane Potential | | | |
|---|---|---|---|
| **Membrane** | **Calculate V (T/F)** | **V initial** | **Specific Capacitance** |
| PM | F | 0.0 mV | 1.0 pF.m² |

| Electrical Mapping - Membrane Potential | | | |
|---|---|---|---|
| **Membrane** | **Calculate V (T/F)** | **V initial** | **Specific Capacitance** |
| NM | F | 0.0 mV | 1.0 pF.m² |

| |
|---|
| Temperature: 300.0 K |

#### 2.8.4. Geometry: Site visit _Application0_20111127_695607844

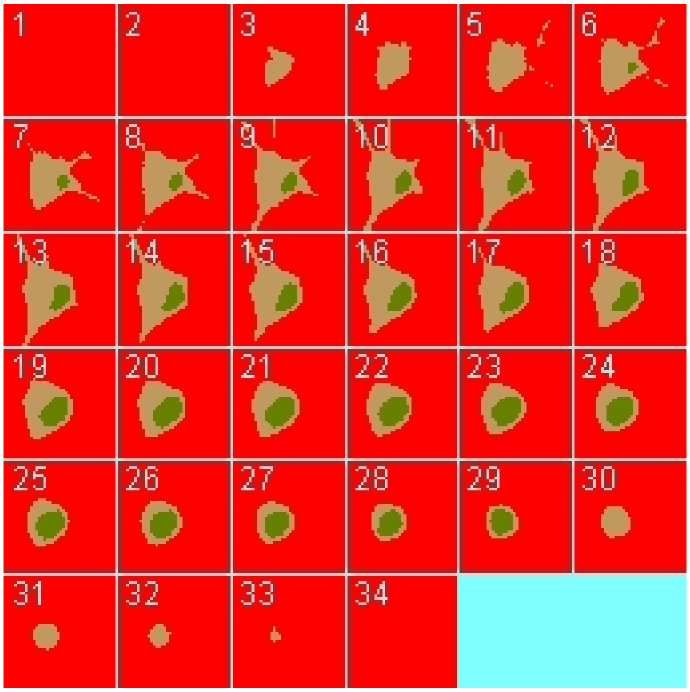

| | |
|---|---|
| Size | (74.24, 74.24, 26.0) |
| Origin | (0.0, 0.0, 0.0) |

#### 2.8.5. Math Description: Copy of Image-based process bleach_generated

##### 2.8.5.1. Constants

| Constant Name | Expression |
|---|---|
| _F_ | 96485.3321 |

| Constant Name | Expression |
| --- | --- |
| _F_nmol_ | 9.64853321E-5 |
| _K_GHK_ | 1.0E-9 |
| _N_pmol_ | 6.02214179E11 |
| _PI_ | 3.141592653589793 |
| _R_ | 8314.46261815 |
| _T_ | 300.0 |
| AreaPerUnitArea_NM | 1.0 |
| AreaPerUnitArea_PM | 1.0 |
| BleachRadius | 2.0 |
| Dark_diffusionRate | 0.1 |
| Dark_init_molecules_um_2 | 0.0 |
| duration | 1.0 |
| Fluor_diffusionRate | 0.1 |
| Fluor_init_molecules_um_2 | 10.0 |
| K_millivolts_per_volt | 1000.0 |
| kbleach | 10.0 |
| KMOLE | 0.001660538783162726 |
| Kr | 0.0 |
| netValence | 1.0 |
| sigmaaxial | 1.5 |
| sigmalateral | 0.5 |
| start | 1.0 |
| Voltage_NM | 0.0 |
| Voltage_PM | 0.0 |
| VolumePerUnitVolume_CYT | 1.0 |
| VolumePerUnitVolum | 1.0 |

| Constant Name | Expression |
|---|---|
| e_EC | |
| VolumePerUnitVolume_NUC | 1.0 |

##### 2.8.5.2. Functions

| Function Name | Expression |
|---|---|
| J_r0 | ((Kf * Fluor) - (Kr * Dark)) |
| Kf | ((kbleach * Laser) * (t > start) * (t < (start + duration))) |
| Laser_init_molecules_um_2 | (exp( - (((z - 8.0) ^ 2.0) / (2.0 * (sigmaaxial ^ 2.0)))) * ((((x - 28.5) ^ 2.0) + ((y - 7.0) ^ 2.0)) < 4.0)) |
| Size_CYT | (VolumePerUnitVolume_CYT * vcRegionVolume('cytosol')) |
| Size_EC | (VolumePerUnitVolume_EC * vcRegionVolume('ec')) |
| Size_NM | (AreaPerUnitArea_NM * vcRegionArea('Nucleus_cytosol_membrane')) |
| Size_NUC | (VolumePerUnitVolume_NUC * vcRegionVolume('Nucleus')) |
| Size_PM | (AreaPerUnitArea_PM * vcRegionArea('cytosol_ec_membrane')) |
| sobj_cytosol1_ec0_size | vcRegionArea('cytosol_ec_membrane') |
| sobj_Nucleus2_cytosol1_size | vcRegionArea('Nucleus_cytosol_membrane') |
| vobj_cytosol1_size | vcRegionVolume('cytosol') |
| vobj_ec0_size | vcRegionVolume('ec') |
| vobj_Nucleus2_size | vcRegionVolume('Nucleus') |

##### 2.8.5.3. Volume Domains

#### 2.8.5.3.1. ec

###### 2.8.5.3.2. cytosol

###### 2.8.5.3.3. Nucleus

#### 2.8.5.4. Membrane Domains

##### 2.8.5.4.1. cytosol_ec_membrane

| PdeEquation Fluor | |
|---|---|
| Rate | - J_r0 |
| Diffusion | Fluor_diffusionRate |
| Initial | Fluor_init_molecules_um_2 |

| PdeEquation Dark | |
|---|---|
| Rate | J_r0 |
| Diffusion | Dark_diffusionRate |
| Initial | Dark_init_molecules_um_2 |

| OdeEquation Laser | |
|---|---|
| Rate | 0.0 |
| Initial | Laser_init_molecules_um_2 |

##### 2.8.5.4.2. Nucleus_cytosol_membrane

#### 2.8.6. Simulation(s)

##### 2.8.6.1. Vary D and bleach Duration_1_1_1

| Simulation Name: Vary D and bleach Duration_1_1_1 |
|---|
| Simulation Description: cloned from 'Vary D and bleach Duration_1_1_1' owned by user temp |

| Overriden Parameters | | |
|---|---|---|
| **Name** | **Actual Value** | **Default Value** |
| Fluor_diffusionRate | 0.1 to 1.0, 2 values | 0.1 |

| | | |
|---|---|---|
| duration | "0.3", "1.0", "3.0" | 1.0 |
| Dark_diffusionRate | Fluor_diffusionRate | 0.1 |

| **Geometry Setting** | |
|---|---|
| Geometry Size (um) | (74.24, 74.24, 26.0) |
| Mesh Size (elements) | (201, 201, 71) |

| **Advanced Settings** | |
|---|---|
| Solver Name | Fully-Implicit Finite Volume, Regular Grid (Variable Time Step) |
| Time Bounds - Starting | 0.0 |
| Time Bounds - Ending | 20.0 |
| Time Step - Min | 0.0 |
| Time Step - Default | 0.05 |
| Time Step - Max | 0.1 |
| Error Tolerance - Absolute | 1.0E-9 |
| Error Tolerance - Relative | 1.0E-7 |
| Output Time Step | 0.5 |
| Use Symbolic Jacobian (T/F) | F |

#### 2.8.6.2. Widefield axial Vary D and bleach Duration_1_1_1

| **Simulation Name: Widefield axial Vary D and bleach Duration_1_1_1** |
|---|
| **Simulation Description: cloned from 'Widefield axial Vary D and bleach Duration_1_1_1' owned by user temp** |

| **Overriden Parameters** | | |
|---|---|---|
| **Name** | **Actual Value** | **Default Value** |

| | | |
|---|---|---|
| Fluor_diffusionRate | 0.1 to 1.0, 2 values | 0.1 |
| sigmaaxial | 30.0 | 1.5 |
| duration | "0.1", "3.0", "1.0" | 1.0 |
| Dark_diffusionRate | Fluor_diffusionRate | 0.1 |

| **Geometry Setting** | |
|---|---|
| Geometry Size (um) | (74.24, 74.24, 26.0) |
| Mesh Size (elements) | (401, 401, 141) |

| **Advanced Settings** | |
|---|---|
| Solver Name | Fully-Implicit Finite Volume, Regular Grid (Variable Time Step) |
| Time Bounds - Starting | 0.0 |
| Time Bounds - Ending | 20.0 |
| Time Step - Min | 0.0 |
| Time Step - Default | 0.05 |
| Time Step - Max | 0.1 |
| Error Tolerance - Absolute | 1.0E-9 |
| Error Tolerance - Relative | 1.0E-7 |
| Output Time Step | 0.5 |
| Use Symbolic Jacobian (T/F) | F |

#### 2.9. Application: 200 um planar membrane

| **Application Name: 200 um planar membrane** |
|---|
| **Application Description: (copied from large planar membrane)** |

##### 2.9.1. Structure Mapping For 200 um planar membrane

| Structure Mapping | | | | |
|---|---|---|---|---|
| **Structure** | **Subdomain** | **Resolved (T/F)** | **Surf/Vol** | **VolFract** |
| NUC | subdomain1 | F | | |
| CYT | subdomain1 | F | | |
| EC | subdomain0 | F | | |

#### 2.9.2. Reaction Mapping For 200 um  planar membrane

| Reaction Mapping | | | |
|---|---|---|---|
| **Name** | **Type** | **Enabled (T/F)** | **Fast (T/F)** |
| r0 | Reaction | T | F |

| Initial Conditions | | | | |
|---|---|---|---|---|
| **Species** | **Structure** | **Initial Conc.** | **Diffusion Const.** | **Fixed (T/F)** |
| s0 | PM | 10.0 molecules.$m^2$ | 0.1 $m^2.s^1$ | F |
| s1 | PM | 0.0 molecules.$m^2$ | 0.1 $m^2.s^1$ | F |
| s2 | PM | (exp( - (((z - 1.0) ^ 2.0) / (2.0 * (sigmaaxial ^ 2.0)))) * ((((x - 100.0) ^ 2.0) + ((y - 100.0) ^ 2.0)) < (BleachRadius ^ 2.0))) molecules.$m^2$ | 0.0 $m^2.s^1$ | F |

#### 2.9.3. Membrane Mapping For 200 um  planar membrane

| Electrical Mapping - Membrane Potential | | | |
|---|---|---|---|
| **Membrane** | **Calculate V (T/F)** | **V initial** | **Specific Capacitance** |
| PM | F | 0.0 mV | 1.0 pF.$m^2$ |
| NM | F | 0.0 mV | 1.0 pF.$m^2$ |

| |
|---|
| Temperature: 300.0 K |

#### 2.9.4. Geometry: Geometry1704059799

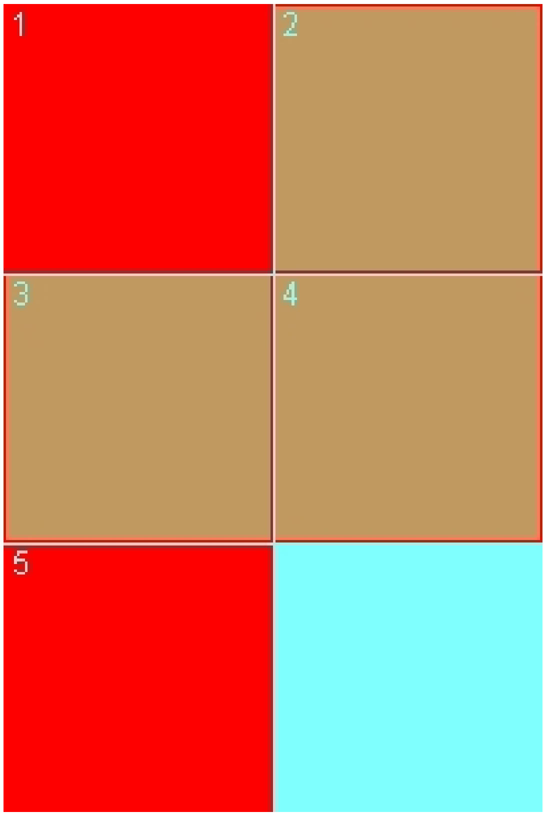

| | |
|---|---|
| Size | (202.0, 202.0, 2.0) |
| Origin | (-1.0, -1.0, -0.5) |

#### 2.9.5. Math Description: Copy of large planar membrane_generated

##### 2.9.5.1. Constants

| Constant Name | Expression |
|---|---|
| _F_ | 96485.3321 |
| _F_nmol_ | 9.64853321E-5 |
| _K_GHK_ | 1.0E-9 |

| Constant Name | Expression |
|---|---|
| _N_pmol_ | 6.02214179E11 |
| _PI_ | 3.141592653589793 |
| _R_ | 8314.46261815 |
| _T_ | 300.0 |
| AreaPerUnitArea_PM | 1.0 |
| AreaPerUnitVolume_NM | 1.0 |
| BleachRadius | 2.0 |
| Dark_diffusionRate | 0.1 |
| Dark_init_molecules_um_2 | 0.0 |
| duration | 1.0 |
| Fluor_diffusionRate | 0.1 |
| Fluor_init_molecules_um_2 | 10.0 |
| K_millivolts_per_volt | 1000.0 |
| kbleach | 10.0 |
| KMOLE | 0.001660538783162726 |
| Kr | 0.0 |
| netValence | 1.0 |
| sigmaaxial | 1.5 |
| sigmalateral | 0.5 |
| start | 1.0 |
| Voltage_NM | 0.0 |
| Voltage_PM | 0.0 |
| VolumePerUnitVolume_CYT | 1.0 |
| VolumePerUnitVolume_EC | 1.0 |
| VolumePerUnitVolum | 0.5 |

| Constant Name | Expression |
| --- | --- |
| e_NUC | |

2.9.5.2. Functions

| Function Name | Expression |
| --- | --- |
| J_r0 | ((Kf * Fluor) - (Kr * Dark)) |
| Kf | ((kbleach * Laser) * (t > start) * (t < (start + duration))) |
| Laser_init_molecules_um_2 | (exp( - (((z - 1.0) ^ 2.0) / (2.0 * (sigmaaxial ^ 2.0)))) * ((((x - 100.0) ^ 2.0) + ((y - 100.0) ^ 2.0)) < (BleachRadius ^ 2.0))) |
| Size_CYT | (VolumePerUnitVolume_CYT * vcRegionVolume('subdomain1')) |
| Size_EC | (VolumePerUnitVolume_EC * vcRegionVolume('subdomain0')) |
| Size_NM | (AreaPerUnitVolume_NM * vcRegionVolume('subdomain1')) |
| Size_NUC | (VolumePerUnitVolume_NUC * vcRegionVolume('subdomain1')) |
| Size_PM | (AreaPerUnitArea_PM * vcRegionArea('subdomain0_subdomain1_membrane')) |
| sobj_subdomain11_subdomain00_size | vcRegionArea('subdomain0_subdomain1_membrane') |
| vobj_subdomain00_size | vcRegionVolume('subdomain0') |
| vobj_subdomain11_size | vcRegionVolume('subdomain1') |

2.9.5.3. Volume Domains

2.9.5.3.1. subdomain1

2.9.5.3.2. subdomain0

2.9.5.4. Membrane Domains

2.9.5.4.1. subdomain0_subdomain1_membrane

| PdeEquation Fluor |
| --- |

| PdeEquation Fluor | |
|---|---|
| Rate | - J_r0 |
| Diffusion | Fluor_diffusionRate |
| Initial | Fluor_init_molecules_um_2 |

| PdeEquation Dark | |
|---|---|
| Rate | J_r0 |
| Diffusion | Dark_diffusionRate |
| Initial | Dark_init_molecules_um_2 |

| OdeEquation Laser | |
|---|---|
| Rate | 0.0 |
| Initial | Laser_init_molecules_um_2 |

#### 2.9.6. Simulation(s)

##### 2.9.6.1. Vary Bleach Radius, FluorD=1

| Simulation Name: Vary Bleach Radius, FluorD=1 |
|---|
| **Simulation Description: cloned from 'Simulation0_1' owned by user temp<br>cloned from 'Simulation0_1' owned by user temp<br>cloned from 'Simulation0' owned by user temp** |

| Overriden Parameters | | |
|---|---|---|
| **Name** | **Actual Value** | **Default Value** |
| BleachRadius | "1.0", "2.0", "4.0" | 2.0 |
| Fluor_diffusionRate | 1.0 | 0.1 |
| Dark_diffusionRate | 1.0 | 0.1 |

| Geometry Setting |
|---|

| | |
|---|---|
| Geometry Size (um) | (202.0, 202.0, 2.0) |
| Mesh Size (elements) | (302, 302, 3) |

| Advanced Settings | |
|---|---|
| Solver Name | Fully-Implicit Finite Volume, Regular Grid (Variable Time Step) |
| Time Bounds - Starting | 0.0 |
| Time Bounds - Ending | 100.0 |
| Time Step - Min | 0.0 |
| Time Step - Default | 0.05 |
| Time Step - Max | 0.1 |
| Error Tolerance - Absolute | 1.0E-9 |
| Error Tolerance - Relative | 1.0E-7 |
| Output Time Step | 0.5 |
| Use Symbolic Jacobian (T/F) | F |

#### 2.10. Application: 100um planar membrane, value BC

| **Application Name: 100um planar membrane, value BC** |
|---|
| **Application Description: (copied from 200 um planar membrane) (copied from large planar membrane)** |

##### 2.10.1. Structure Mapping For 100um planar membrane, value BC

| Structure Mapping | | | | |
|---|---|---|---|---|
| **Structure** | **Subdomain** | **Resolved (T/F)** | **Surf/Vol** | **VolFract** |
| NUC | subdomain1 | F | | |
| CYT | subdomain1 | F | | |
| EC | subdomain0 | F | | |

##### 2.10.2. Reaction Mapping For 100um planar membrane, value BC

| Reaction Mapping | | | |
|---|---|---|---|
| **Name** | **Type** | **Enabled (T/F)** | **Fast (T/F)** |
| r0 | Reaction | T | F |

| Initial Conditions | | | | |
|---|---|---|---|---|
| **Species** | **Structure** | **Initial Conc.** | **Diffusion Const.** | **Fixed (T/F)** |
| s0 | PM | 10.0 molecules.$m^{2}$ | 0.1 $m^{2}.s^{1}$ | F |
| s1 | PM | 0.0 molecules.$m^{2}$ | 0.1 $m^{2}.s^{1}$ | F |
| s2 | PM | (exp( - (((z - 1.0) ^ 2.0) / (2.0 * (sigmaaxial ^ 2.0)))) * ((((x - 50.0) ^ 2.0) + ((y - 50.0) ^ 2.0)) < (BleachRadius ^ 2.0))) molecules.$m^{2}$ | 0.0 $m^{2}.s^{1}$ | F |

##### 2.10.3. Membrane Mapping For 100um planar membrane, value BC

| Electrical Mapping - Membrane Potential | | | |
|---|---|---|---|
| **Membrane** | **Calculate V (T/F)** | **V initial** | **Specific Capacitance** |
| PM | F | 0.0 mV | 1.0 pF.$m^{2}$ |
| NM | F | 0.0 mV | 1.0 pF.$m^{2}$ |

| |
|---|
| Temperature: 300.0 K |

##### 2.10.4. Geometry: Geometry1704059799

| | |
|---|---|
| Size | (102.0, 102.0, 2.0) |
| Origin | (-1.0, -1.0, -0.5) |

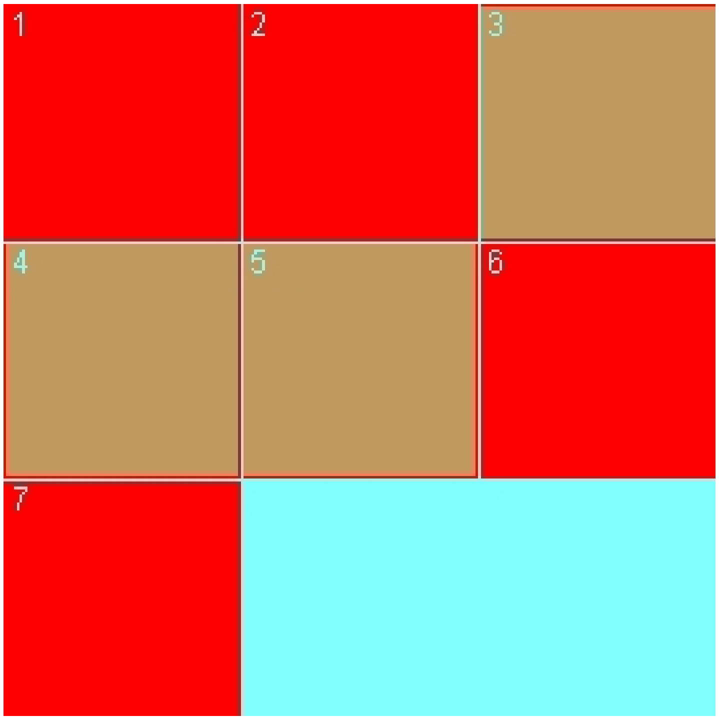

#### 2.10.5. Math Description: Copy of 200 um  planar membrane_generated

##### 2.10.5.1. Constants

| Constant Name | Expression |
| --- | --- |
| _F_ | 96485.3321 |
| _F_nmol_ | 9.64853321E-5 |
| _K_GHK_ | 1.0E-9 |
| _N_pmol_ | 6.02214179E11 |
| _PI_ | 3.141592653589793 |
| _R_ | 8314.46261815 |
| _T_ | 300.0 |
| AreaPerUnitArea_PM | 1.0 |
| AreaPerUnitVolume_NM | 1.0 |
| BleachRadius | 2.0 |

| Constant Name | Expression |
| --- | --- |
| Dark_diffusionRate | 0.1 |
| Dark_init_molecules_um_2 | 0.0 |
| duration | 1.0 |
| Fluor_diffusionRate | 0.1 |
| Fluor_init_molecules_um_2 | 10.0 |
| K_millivolts_per_volt | 1000.0 |
| kbleach | 10.0 |
| KMOLE | 0.001660538783162726 |
| Kr | 0.0 |
| netValence | 1.0 |
| sigmaaxial | 1.5 |
| sigmalateral | 0.5 |
| start | 1.0 |
| Voltage_NM | 0.0 |
| Voltage_PM | 0.0 |
| VolumePerUnitVolume_CYT | 1.0 |
| VolumePerUnitVolume_EC | 1.0 |
| VolumePerUnitVolume_NUC | 0.5 |

#### 2.10.5.2. Functions

| Function Name | Expression |
| --- | --- |
| J_r0 | ((Kf * Fluor) - (Kr * Dark)) |
| Kf | ((kbleach * Laser) * (t > start) * (t < (start + duration))) |
| Laser_init_molecules_um_2 | (exp( - (((z - 1.0) ^ 2.0) / (2.0 * (sigmaaxial ^ 2.0)))) * ((((x - 50.0) ^ 2.0) + ((y - 50.0) ^ 2.0)) < (BleachRadius ^ 2.0))) |

| Function Name | Expression |
| --- | --- |
| Size_CYT | (VolumePerUnitVolume_CYT * vcRegionVolume('subdomain1')) |
| Size_EC | (VolumePerUnitVolume_EC * vcRegionVolume('subdomain0')) |
| Size_NM | (AreaPerUnitVolume_NM * vcRegionVolume('subdomain1')) |
| Size_NUC | (VolumePerUnitVolume_NUC * vcRegionVolume('subdomain1')) |
| Size_PM | (AreaPerUnitArea_PM * vcRegionArea('subdomain0_subdomain1_membrane')) |
| sobj_subdomain11_subdomain00_size | vcRegionArea('subdomain0_subdomain1_membrane') |
| vobj_subdomain00_size | vcRegionVolume('subdomain0') |
| vobj_subdomain11_size | vcRegionVolume('subdomain1') |

#### 2.10.5.3. Volume Domains

##### 2.10.5.3.1. subdomain1

##### 2.10.5.3.2. subdomain0

#### 2.10.5.4. Membrane Domains

##### 2.10.5.4.1. subdomain0_subdomain1_membrane

| PdeEquation Fluor | |
| --- | --- |
| Rate | - J_r0 |
| Diffusion | Fluor_diffusionRate |
| Initial | Fluor_init_molecules_um_2 |

| PdeEquation Dark | |
| --- | --- |
| Rate | J_r0 |
| Diffusion | Dark_diffusionRate |
| Initial | Dark_init_molecules_um_2 |

| OdeEquation Laser | |
|---|---|
| Rate | 0.0 |
| Initial | Laser_init_molecules_um_2 |

#### 2.10.6. Simulation(s)

##### 2.10.6.1. Vary Bleach Radius, FluorD=1_1

| **Simulation Name: Vary Bleach Radius, FluorD=1_1** |
|---|
| **Simulation Description: cloned from 'Vary Bleach Radius, FluorD=1_1' owned by user temp<br>cloned from 'Simulation0_1' owned by user temp<br>cloned from 'Simulation0_1' owned by user temp<br>cloned from 'Simulation0' owned by user temp** |

| Overriden Parameters | | |
|---|---|---|
| **Name** | **Actual Value** | **Default Value** |
| BleachRadius | "1.0", "2.0", "4.0" | 2.0 |
| Fluor_diffusionRate | 1.0 | 0.1 |
| Dark_diffusionRate | 1.0 | 0.1 |

| Geometry Setting | |
|---|---|
| Geometry Size (um) | (102.0, 102.0, 2.0) |
| Mesh Size (elements) | (402, 402, 3) |

| Advanced Settings | |
|---|---|
| Solver Name | Fully-Implicit Finite Volume, Regular Grid (Variable Time Step) |
| Time Bounds - Starting | 0.0 |
| Time Bounds - Ending | 100.0 |

| | |
|---|---|
| Time Step - Min | 0.0 |
| Time Step - Default | 0.05 |
| Time Step - Max | 0.1 |
| Error Tolerance - Absolute | 1.0E-9 |
| Error Tolerance - Relative | 1.0E-7 |
| Output Time Step | 0.5 |
| Use Symbolic Jacobian (T/F) | F |
